# Biological Factors and Self-Perception of Stress Predict Human Freeze-Like Responses in the Context of Self-Defence Training and Personal Experience with Violence

**DOI:** 10.1101/2021.11.02.466879

**Authors:** Peter Lenart, Michal Vít, Klára Marečková, Jan Novák, Filip Zlámal, Michal Mikl, Zdenko Reguli, Martin Bugala, Jitka Čihounková, Pavel Přecechtěl, Vojtěch Malčík, Tomáš Vojtíšek, Jan Kučera, Jana Fialová Kučerová, Veronika Hajnová, Marie Tomandlová, Radek Šíp, Lucie Ráčková, Markéta Grulichová, Josef Tomandl, Milan Brázdil, Julie Bienertova-Vasku

## Abstract

Many animals react to threatening stimuli such as a predator attacks by freezing. However, little experimental research investigated freeze response in humans. Here, we have employed practices commonly used in self-defence training to create two unique scenarios simulating armed physical attacks. Sixty healthy men volunteers divided into three groups of twenty (untrained, trained but unexperienced, trained and experienced) underwent these scenarios accompanied by measurement of biochemical, physiological, and psychological markers of stress. All participants also underwent an fMRI session during which they observed neutral and negative images from the International Affective Picture System (IAPS). Our results show that scenarios simulating physical attacks can induce a freeze-like response in men. In addition, we demonstrate that while electrodermal activity (EDA), subjective stress perception, and brain activity in fMRI predict freeze-like response in men, their effect on freeze-like response is entirely dependent on the level of training and experience of a given individual.

## Introduction

It is well established that animals react to danger by employing one of three main strategies: fight, flight, or freeze.^1–3^ While processes aroused by the feeling of danger, fear, or anxiety are more complex in humans, they lead to very similar reactions.^4^ From the human perspective, such reactions are relevant for a wide range of real-life situations and essential for various professions such as police officers, firefighters, etc., where dangerous high-stress situations are common. Of the three reactions to danger, freeze – sometimes conflated with tonic immobility^5^ – stands out, since it is harmful in almost any imaginable situation a modern-day person might encounter. If, for example, a police officer is attacked by an armed criminal, or if the pilots of a commercial plane realize that the engines have stopped working and only have a limited time to react until the plane crashes, freezing is the worst possible reaction. Thus, in professional and personal self-defence training, it is especially important to reinforce the ability to switch between distinct defensive behaviors, such as freezing and active fight-or-flight responses.^6^ However, while researchers have studied freeze response in animals for decades, studies of freeze response in humans are scarce. No published study to date has explored the human freeze response in reaction to a convincing simulation of a physical attack.^7,8^

Thus far, studies of humans have reported freeze-like behavior, assessed by postural control, skin conductance, and heart rate, in response to images of mutilation^9,10^, images of angry faces^11^, unpleasant images provided by the International Affective Picture System^12^ or footage showing the aftermath of fatal car accidents.^13^ Moreover, it was shown that humans might experience a freeze-like response operationalized as a feeling of immobility after breathing CO_2_-enriched air.^5^ Most interestingly, it was also demonstrated that the approaching motion of threatening stimuli (images of spiders or snakes) could induce a freeze-like response, operationalized as a slowing of reaction times.^14^ This observation seems to be more closely connected to the origins of the concept of freeze than viewing images of angry faces. Furthermore, an individual’s vulnerability to freezing after being subject to such threatening stimuli depends on their cognitive vulnerability to anxiety.^15^ Studies where participants observed images of an attacker aiming a firearm at them were presumably closest to real-life conditions of interpersonal violence.^6,16^ However, these simulations were still far from convincing.

While the above-mentioned studies all contribute to research into human freeze response, they all share the same limitations. The feeling of danger these studies induced was relatively mild and thus hardly comparable with high-stress situations where freezing would be expected to occur in real life, such as a physical attack. Therefore, in our study, 60 male volunteers underwent two specially devised scenarios simulating armed physical attacks to study human freeze response in a more realistic setting.

We hypothesized that freeze-like responses in humans may be affected by stress levels, and we thus measured biochemical, physiological, and psychological markers of stress. We chose cortisol and osteocalcin as biochemical stress markers since both are well-known stress-related hormones^17–20^ and because cortisol levels were previously associated with freezing in rodents and primates.^21,22^ As physiological stress markers we measured heart rate variability (HRV) and Electrodermal activity (EDA).^23,24^ Both have been utilized, with varying results, in previous studies of human freeze-like response.^7,25^ As psychological markers of stress we recorded subjective ratings of physical and psychological stress reported by participants and measured anxiety using the State-Trait Anxiety Inventory questionnaire which was previously connected to freeze-like response in humans.^26^ In addition, we hypothesized that since parts of the limbic system are involved in freeze response in animals,^7,27–31^ its activation while regulating the response to negative affective images presented during the fMRI task will predict freeze-like response in scenarios simulating physical attacks.

## Methods

### Study subjects

We recruited 60 healthy men and divided them into three groups. The first group (A) consisted of 20 healthy male controls with no previous experience with martial arts and combat sports or self-defence training. The second group (B) included 20 students of the Special Education of Security Bodies study program at the Faculty of Sports Studies at Masaryk University. These individuals were specially educated to serve in first responder professions and well trained in martial arts and combat sports as well as in personal self-defence. The third group (C) consisted of 20 members of the Rapid response unit of the Police of the Czech Republic from the South Moravian headquarters. This SWAT team is designed to intervene against particularly dangerous and armed perpetrators. These participants have extensive professional self-defence training (including hand-to-hand combat skills, arresting techniques, and weapons training) as well as real-life experience with situations involving physical violence.

Ethical approval for the study was obtained from the Research Ethics Committee of Masaryk University under Ref. No.: EKV-2018-106-R1 on 3 June 2019 for project MUNI/A/1403/2018. Written informed consent was obtained from all study participants. Participants could end the scenario or leave the study altogether at any given time and were repeatedly made aware of this fact. Special safety precautions were taken in scenarios with participants from group A.

### Study chronology

All participants visited the university three times. During their first visit they were introduced to the study and signed the informed consent form. They also provided a biological sample of blood and saliva, filled in the Trait part of the State-Trait Anxiety Inventory questionnaire (STAI-T), and underwent structural and functional magnetic resonance imaging (MRI).

During their second visit, all subjects participated in our low-anxiety scenario (LA) – simulating a physical attack – while wearing a set of sensors measuring physiological markers of stress (heart rate, galvanic skin response) and camera glasses. Several static cameras also recorded the scenario itself from different angles. Immediately before and after the scenario, participants provided blood and saliva samples, filled out the State part of the State-Trait Anxiety Inventory questionnaire (STAI-S), and reported a subjective rating of physical and psychological stress on a scale from 1 to 10.

During the third visit, all subjects participated in our high-anxiety scenario (HA) which simulated a physical attack. Again, several cameras recorded the scenario itself from different angles and each participant was equipped with camera glasses. All participants were equipped with the same set of sensors used during the low-anxiety scenario. They provided blood and saliva samples, filled out the STAI-S questionnaire, and reported a subjective rating of physical and psychological stress on a scale from 1 to 10.

### Scenarios

Scenarios simulating a physical attack are common in personal self-defence training, police officer training, etc. However, such scenarios were, to the best of our knowledge, never before used to induce stress to study freeze response in humans. Therefore, both of the scenarios we employed were developed uniquely for this study. The intensity of the LA scenario was similar to scenarios commonly used for self-defence training. The HA scenario differed from classical self-defence training in that it more prominently featured an element of surprise. In both scenarios attackers modified the physical attack intensity according to the skill levels of individual participants. In general, physical attacks were thus less intense in the case of untrained individuals.

Physical attack intensity was also modified according to participant response. For example, when participants froze, the attackers significantly decreased the speed and frequency of active attacks and interacted more verbally, e.g., by encouraging participants to action by phrases such as “Defend yourself!” or “Get out!”. In case the participants did not begin to actively defend themselves, the supervising researcher ended the scenario.

Prior to the beginning of the scenarios, all participants were informed that each scenario would involve the staging of an attempted crime, including a verbal or physical attack by a person equipped with a protective suit. The participants were instructed to react as they would in real life, which means that reasonable physical contact with the attacker was permitted. All participants were also repeatedly instructed that they could end the scenario prematurely at any time by verbally stating “stop” or using any other words or sentences indicating that they intended to stop the scenario. Participants from inexperienced group A wore protective equipment (safety helmet) to minimize the risk of accidental injury. After the scenario, all participants took part in a psychological debriefing. Immediately before and after each of the scenarios and during the debriefing sessions, participants reported their subjective feelings of physical and mental stress on a scale from 1 to 10.

### Low anxiety scenario

In this scenario participants entered a gym used for martial arts and combat sports practice and self-defence training, already expecting a potential unspecified verbal or/and physical attacks. The synopsis of the low anxiety scenario was simple. Participants were instructed to enter the venue, position themselves on an “X” marked on the floor, and observe the scene. No specific task was set. Thirty seconds after the participant entered the scene, a mock attacker dressed in a protective suit (High Gear Impact Reduction suit by Blauer Tactical Systems Inc.) and armed with a fake rubber knife appeared on the opposite side of the gym. For the first ten seconds, the attacker passively observed the participant. Then, the attacker began to communicate with the participant. The communication escalated into a verbal conflict and the attacker slowly began to approach the participant. The intensity of the verbal attack increased and resulted in a simulated physical attack using the fake rubber knife. The participants could react to the situation in any way (e.g., stop the scenario, escape, or resist the attacker). After the scenario, all participants took part in a psychological debriefing session.

### High anxiety scenario

The high-anxiety scenario was more complex and task-specific. Participants were instructed to walk through a complicated path divided into different rooms and corridors (total length of approximately 500 meters). They were asked to enter a particular room, locate a piece of paper with their code number, and bring it back to the starting location. However, these instructions were only a distraction. In reality, the participant went down a flight of stairs to the basement and entered a room specifically designed for the scenario (equipped with protective equipment including carpets and mats). After a few steps, loud dog barking sounds were played behind the participants, and, immediately afterward, a mock attacker appeared on the other side of the room. The attacker, dressed in a protective suit (High Gear Impact Reduction suit by Blauer Tactical Systems Inc.), attacked each participant verbally and then physically like in the low-anxiety scenario. The spatial conditions were the same as in the low anxiety scenario (distance between the attacker and participant, room dimensions). However, compared to the low-anxiety scenario, the attack simulation happened much faster, giving the participants little time to plan their defence. In addition, the attacker used a special electric plastic knife (Shocknife™ SK-2 by Setcan Corporation) which imparts a very mild electric shock upon contact and produces strong visual and audio effects while in use. By increasing the intensity of the verbal attack, the attacker shortened the distance and started to simulate a physical attack with a plastic knife. The participants could react freely to the situation in any way (e. g. stop the scenario, escape, or resist the attacker). Participants from the inexperienced group A wore protective equipment (safety helmets) to minimize the risk of accidental injury. After the scenario, all participants took part in a psychological debriefing session.

### Psychological debriefing

After both scenarios, participants were taken to a quiet room where they took part in a psychological debriefing session. The interview was led by an experienced police psychologist and the head of the Department of Gymnastics and Combatives, responsible for implementing the scenarios.

The primary goal of the psychological debriefing session was to ensure that no mental or physical harm was done to the participants during the course of the scenario and that they returned to a resting mental state similar to that in which they came to the research facility. The debriefing was conducted in the form of a semistructured interview, in which the psychologist asked prepared questions and evaluated the participant’s answers.

The secondary goal of the psychological debriefing session was to ascertain additional information about the subjective perception of the course of the scenarios by the participants (how they perceived the threat, how they evaluated their decision to solve the scenario, etc.).

The psychological debriefing session was recorded (audio) for documentation and further analysis purposes. After confirming that all ethical and safety procedures were followed, the participants were allowed to leave the research facility.

### Freeze-like response evaluation

Participants’ performance was assessed using videorecordings of the the scenarios by a panel of six experts. Five of them were martial arts and combat sports and self-defence training experts (see Supplementary Data 1 for qualifications) while the sixth was a forensic pathologist with 22 years of experience, who estimated the likelihood of survival for each participant in each scenario. The freeze-like response, operationalized as a passive reaction to attack, was evaluated on a scale from 0 to 4, where 0 corresponded to a completely passive reaction and 4 to a very active reaction. Precise criteria for the evaluation are listed in Supplementary Data 1. The experts evaluated the video recordings simultaneously in one room using the same video recordings. Importantly, expert evaluators sat behind different computers and could not consult each other or share their scores between themselves while scoring participants.

### MRI and fMRI data acquisition

MRI was acquired using a 3T Siemens Prisma scanner with a 64-channel head-neck coil. The MRI protocol included T1 MPRAGE structural images and a mild visual stress fMRI task where participants observed neutral and negative images from the International Affective Picture System (IAPS) and were instructed to either observe or regulate their negative affect induced by the stimuli (Supplementary Data 2). High-resolution T1 MPRAGE images were obtained using 0.8 mm isotropic voxels covering the whole head. The fMRI acquisition details were as follows: TR = 0.7 s; 3 echoes, TE = 16 ms, 38 ms, and 59 ms; voxel size 3 × 3 × 3 mm; whole-brain coverage; MB factor 6; PAT factor 2; FA 47°, total of 2006 scans.

### MRI and fMRI data processing

All MRI data were processed in SPM12, build 6225 (Wellcome Trust Centre for Neuroimaging at University College London, UK; http://www.fil.ion.ucl.ac.uk/spm/), running under Matlab 8.4. R2014b. The functional images were processed as follows: the time series were realigned to the first volume to correct for head motion, transformed into standard stereotactic space (MNI), and spatially smoothed with a Gaussian filter (FWHM = 5 mm). The T1-weighted anatomical high-resolution data were registered to the mean functional image of each subject and spatially normalized into MNI space. Task data were processed with a general linear model as implemented in SPM12, modeling the three conditions and contrasting them. T-statistics maps for individual contrasts were parceled with an AAL atlas. To decrease the number of comparisons, we chose six brain regions from the limbic system (hippocampus, parahippocampus, amygdala, thalamus, insula, cingulum) corresponding to 16 AAL defined regions (6 AAL regions for cingulum 2 ALL for each of the remaining regions; the activity of each brain region was obtained by averaging the responses over corresponding AALs, see Supplementary Data 3). Pearson’s correlations between the activity of these six regions in different contrasts and freeze-like scores in LA and HA scenarios were then calculated.

### Biomarkers

Blood and saliva were collected at five time points (baseline and immediately before and after each of the two scenarios). Saliva was kept on ice, centrifuged, and mixed with Protease Inhibitor Cocktail (#P2714, MilliporeSigma). Plasma was extracted from EDTA-whole blood, and both saliva and plasma samples were processed and stored at -80 °C within 90 minutes.

To assess salivary cortisol, thawed and centrifuged samples were run in duplicates using Human Cortisol (Saliva) ELISA kit (#RCD005R, BioVendor LM, Czech Republic) according to the manufacturer’s instructions.

The quantitative measurement of undercarboxylated osteocalcin in plasma samples was performed in duplicate using the Human ucOC ELISA Kit (#MBS700581, MyBioSource, CA, USA) according to the manufacturer’s instructions.

### HRV and EDA

Physiological data of electrodermal activity (EDA) and heart rate variability (HRV) were recorded during both scenarios. EDA signal was measured using the E4 Wristband device by Empatica Inc. Company. The device was placed on each participant’s right wrist. For EDA signal acquisition, disposable pre-gelled ECG Ag/Cl electrodes were used. Electrodes placed on the first phalanx of the index finger and middle finger were connected to an E4 Wristband by E4 EDA lead wires extension. Data was saved using the E4 Manager software and cloud.

HRV signal was measured using Polar Vantage V with the heart rate sensor Polar H10 device by Polar Electro Inc. The sensor was placed on each participant’s chest. Data was saved using the Polar Flow software and cloud.

### Statistical analysis

All descriptive statistics of continuous variables are represented by mean and standard deviation. Intergroup comparisons (i.e. between performance groups A, B, and C) were performed using parametric (ANOVA, linear model using generalized least squares – in the case of unequal variances) or nonparametric tests (Kruskal-Wallis test). Normality was verified using statistical tests of normality (Shapiro-Wilk, Anderson-Darling, Pearson tests) and graphically (histogram, Q-Q plot). In case data did not follow normality, we log transformed the data and verified the normality (of these transformed data) again. Dependence between continuous variables was studied using pairwise Pearson’s correlation coefficient.

The dynamic of cortisol and osteocalcin with regard to performance group membership was studied with the linear model using generalized least squares (R package `nlme`)^32^. This type of model facilitates the modelling of mean values of the response variable depending on different kinds of independent variables and accounting for various types of correlations in the data (time, spatial) and heteroscedasticity. In addition, *post hoc* tests were implemented using multiple comparison tests with p-value adjustment using the Tukey method (R package `emmeans`)^33^.

Linear regression models were used to study the effect of BOLD activity (treated as independent variable) on freeze like response (treated as dependent variable), separately for LA and HA scenarios.

Moreover, four group of variables (anthropometry, cortisol and osteocalcin levels, subjective perception of stress and STAI levels) were incorporated as potential confounders.

Because of relatively high correlation levels between variables in each group first FA (in each group) was conducted. Extracted factors were treated as independent variables in the models.

Statistical analysis was performed in statistical software R (version 3.5.1) using Rstudio software (version 1.2.5019).^34^ All results of statistical testing with p-values less than 0.05 were considered significant.

## Results

### Inter- and intra-group comparisons of freeze-like response

As expected, participants in the untrained A group exhibited a more intense freeze-like response in both scenarios. On the other hand, freeze-like response was least pronounced among professionals in group C (Figure 1). However, more surprisingly, while the intensity of freeze-like response in trained group B and professional group C participants increased in the HA scenario compared to LA, untrained group A participants froze less in the more intense HA scenario.

**Figure 1.**
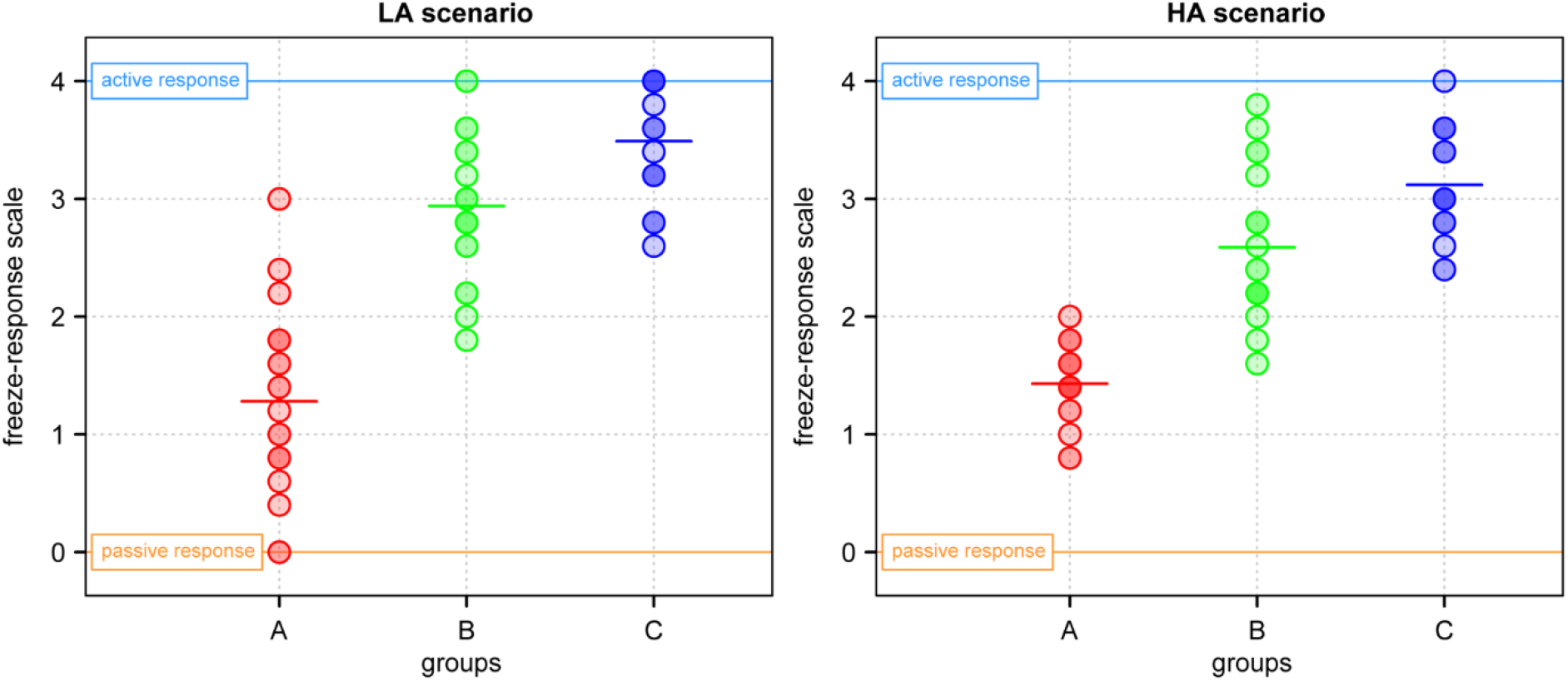
Freeze-like response in low anxiety (LA) and high anxiety (HA) scenarios. A higher score reflects less intense freezing and conversely. Group A consisted of 20 healthy males without previous experience in martial arts, combat sports, or self-defence training. Group B included 20 males trained in martial arts, combat sports, and personal self-defence, but without real-life experience with violence. Group C consisted of 20 members of the Rapid Response Unit of the Police of the Czech Republic from the South Moravian headquarters. These males were both trained and had experience with real-life violence.

The increase in average score in the HA scenario compared to the LA scenario in group 1 stems from the fact that this was the only group where several participants scored 1 point or lower. Interestingly, all participants who scored 1 or lower on the freeze-like response in the LA scenario improved their score in HA, implying that participants who experienced the worst freeze-like reaction also learned the most from this experience. On the other hand, only three participants from trained group B and only two participants from professional group C increased their freeze-like score in the HA scenario compared to the LA scenario.

When merged together, all three groups exhibited a strong significant correlation between freeze-like response in LA and HA scenarios (r = 0.80, p < 0.001). However, this was likely caused mostly by the intergroup differences in freeze-like response. There was no significant correlation between freeze-like response in LA and HA scenarios in groups A and B and a weak correlation in group C (r = 0.40, p = 0.077; r = 0.21, p = 0.374; r = 0.60, p = 0.005; respectively).

Interestingly, while the intensity of the freeze-like response did not affect the likelihood of survival in the LA scenario (p = 0.738), it significantly affected survival in the HA scenario (p = 0.014). Unsurprisingly, the likelihood of survival was also modified by group membership (p < 0.001) in both scenarios.

### State-Trait Anxiety Inventory questionnaire

STAI questionnaire scores significantly differed between groups at all time points (Table 1). Surprisingly, baseline STAI-S values (time points 2 and 4) were highest in group B, suggesting that group B experienced the highest anticipatory stress levels.

**Table 1.** Average group scores from the State-Trait Anxiety Inventory questionnaire. 1 – before MRI. 2 – before LA scenario. 3 – after LA scenario. 4 – before HA scenario. 5 – after HA scenario. Group A consisted of 20 healthy males without previous experience in martial arts, combat sports, or self-defence training. Group B included 20 males trained in martial arts, combat sports, and personal self-defence, but without real-life experience with violence. Finally, group C consisted of 20 members of the Rapid Response Unit of the Police of the Czech Republic from the South Moravian headquarters. These males were both trained and had experience with real-life violence.

| Questionnaire<br>(time point) | Groups (average score ± SD) | | | p-value | transformation | test |
| --- | --- | --- | --- | --- | --- | --- |
|  | A | B | C |  |  |  |
| STAI-T (1) | 36.45 ± 8.13 | 34.75 ± 7.12 | 27.55 ± 4.20 | <0.001 | ln | ANOVA |
| STAI-S (2) | 32.84 ± 6.66 | 39.40 ± 9.25 | 30.35 ± 5.00 | 0.001 | ln | ANOVA |
| STAI-S (3) | 43.55 ± 8.88 | 42.75 ± 9.14 | 32.05 ± 7.10 | <0.001 | id | ANOVA |
| STAI-S (4) | 36.45 ± 8.90 | 37.70 ± 10.12 | 30.80 ± 5.22 | 0.005 | id | MEM |
| STAI-S (5) | 40.90 ± 9.62 | 40.26 ± 12.37 | 32.65 ± 5.88 | 0.015 | id | ANOVA |

After adjustment for multiple comparisons, professionals from group C had significantly lower STAI-T scores than either groups A or B (p = <0.001; p = 0.001). No significant difference between groups A and B was established (p = 0.725). Group C also had significantly lower STAI-S score than group B at every time point (p = 0.001; p = <0.001; p = 0.030; p = 0.042 for time points 2 to 5). Group C also had a significantly lower STAI-S score than untrained group A after both scenarios (p = <0.001 and p = 0.024 for LA and HA scenarios). However, STAI-S scores for groups C and A measured before the scenarios were not significantly different (p = 0.521 and 0.052). STAI-S scores measured before the LA scenario differed significantly between the A and B groups (p = 0.030), but at every other time point, the STAI-S scores of groups A and B were virtually the same (p = 0.952; p = 0.910; p = 0.977 for time points 3 to 5).

Across all groups, STAI-T scores measured before MRI sessions showed a weak but significant correlation with STAI-S scores across all time points (Supplementary Data 4). Across individual groups, the professional group C showed the highest correlation between STAI-T and STAI-S scores (r= 0.50, p = 0.025; r= 0.66, p = 0.002; r= 0.58, p = 0.008; r= 0.55, p = 0.012; for time points 2 to 5) (Supplementary Data 4). Trained group B also showed a consistent correlation between STAI-T and STAI-S scores across all time points (Supplementary Data 4). However, the untrained group A exhibited a very different trend. Not only were the correlations between STAI-T and STAI-S significant only after each of the scenarios (time points 3 and 5), but the correlation between STAI-T and STAI-S was negative (r= -0.53, p = 0.016; r= -0.47, p = 0.034; for time points 3 and 5) (Supplementary Data 4). These results thus suggest that STAI-T scores have a very different meaning in differently trained and experienced individuals.

While STAI-S scores were affected by the LA and HA scenarios and significant correlations between STAI-T and STAI-S scores were established, no significant correlation between STAI-T or STAI-S scores and freeze-like response was observed in HA scenarios. Accordingly, no significant correlation was found for groups B and C in the LA scenario. However, STAI-S measured before the LA scenario showed a significant negative correlation with freeze-like response in untrained group A (r= -0.56, p = 0.014).

### Cortisol and osteocalcin levels

In this study, cortisol and osteocalcin were measured for two related reasons. First, cortisol and osteocalcin were measured immediately before and after each scenario to assess if the scenarios were as stressful as intended for all groups despite differences in training and experience with real-life violence. In this regard, our results show that both LA and in particular the HA scenario were stressful even for the most trained and experienced participants from group C (Figure 2). Furthermore, it seems that osteocalcin is a more reliable marker of stress of the two, as, unlike in the case of cortisol, its baseline levels remained relatively stable during all three baseline samplings (before MRI, before LA scenario, and before HA scenario). In addition, osteocalcin levels were consistent with the STAI questionnaire data showing that group B exhibited the highest anticipatory stress: their osteocalcin levels were consistently higher than groups A and C.

**Figure 2.**
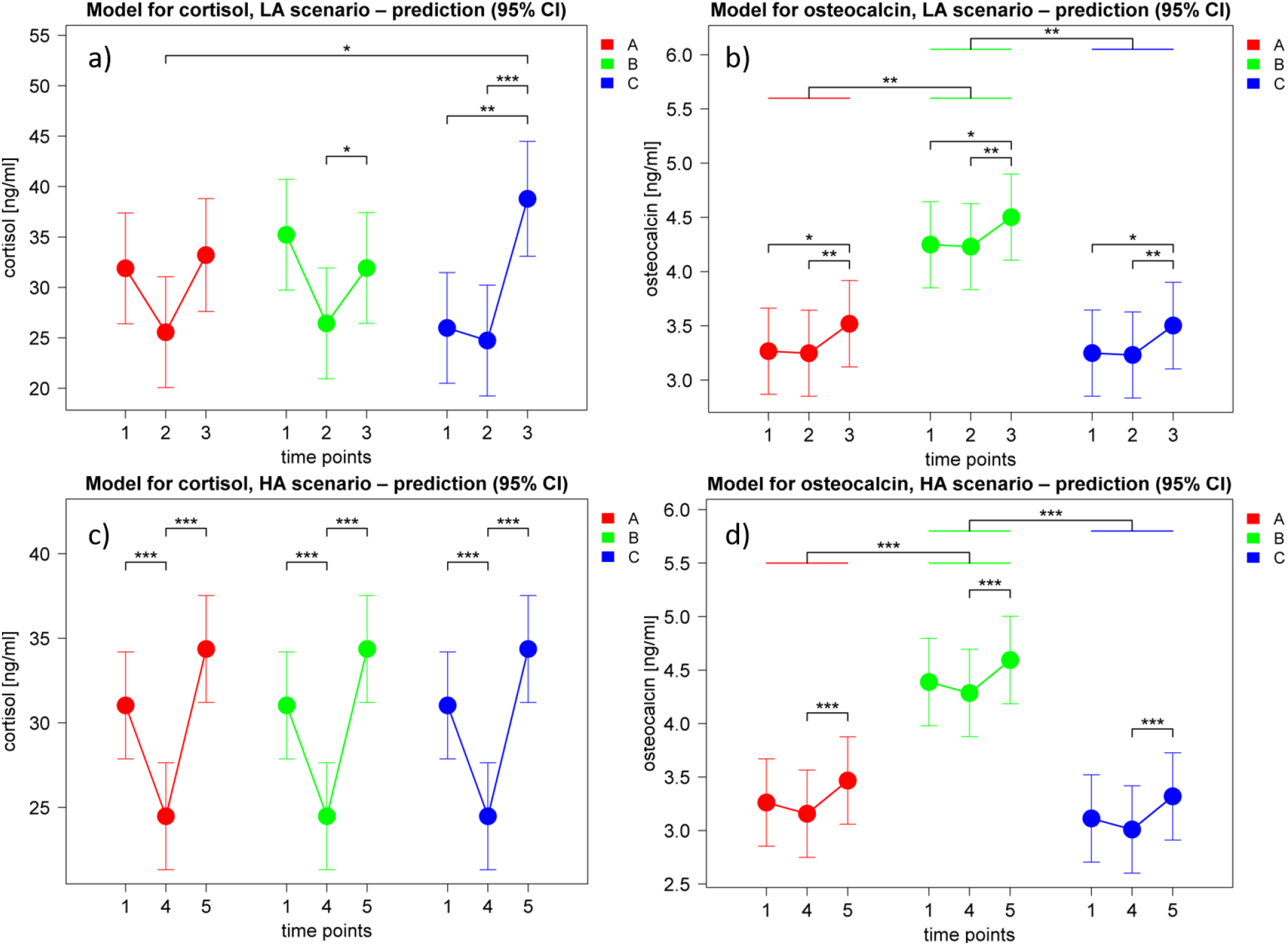
Change in cortisol and osteocalcin levels in LA and HA scenarios. Cortisol and osteocalcin were measured at five time points. 1 – before MRI. 2 – before LA scenario. 3 – after LA scenario. 4 – before HA scenario. 5 – after HA scenario. Group A consisted of 20 healthy males without previous experience in martial arts, combat sports, or self-defence training. Group B included 20 males trained in martial arts, combat sports, and personal self-defence, but without real-life experience with violence. Finally, group C consisted of 20 members of the Rapid Response Unit of the Police of the Czech Republic from the South Moravian headquarters. These males were both trained and had experience with real-life violence

While cortisol and osteocalcin levels confirmed what the STAI-S scores indicated, i.e. that both LA and HA scenarios were stressful, and osteocalcin levels even affirmed the higher basal stress levels of participants from group B, no correlation between STAI-S scores and cortisol and osteocalcin levels (Supplementary Data 4).

The second reason for measuring cortisol and osteocalcin was to assess whether their levels predict subsequent freeze-like responses in LA and HA scenarios. In short, they do not. Adding freeze-like response score, BMI, or age to regression models predicting osteocalcin or cortisol levels did not significantly improve the model over models predicting osteocalcin or cortisol levels from a time point and group affiliation (Supplementary Data 4).

### Subjective stress perception

In both scenarios, subjectively perceived physical and mental stress significantly rose after the scenario and then significantly decreased before the start of the debriefing session for all three groups (in all cases p < 0.001; Figure 3). Furthermore, in all comparisons, self-perceived physical and mental stress levels after the debriefing were no longer significantly different from self-perceived physical and mental stress levels before the scenario (Figure 3). This observation constitutes further evidence suggesting that our scenarios were stressful as intended for all groups despite their different levels of training and experience with real-life violence. In addition, these findings also show that the stress induced by the scenarios was transient and short-term as intended and that participants calmed down to pre-scenario levels even before the start of the debriefing session.

**Figure 3.**
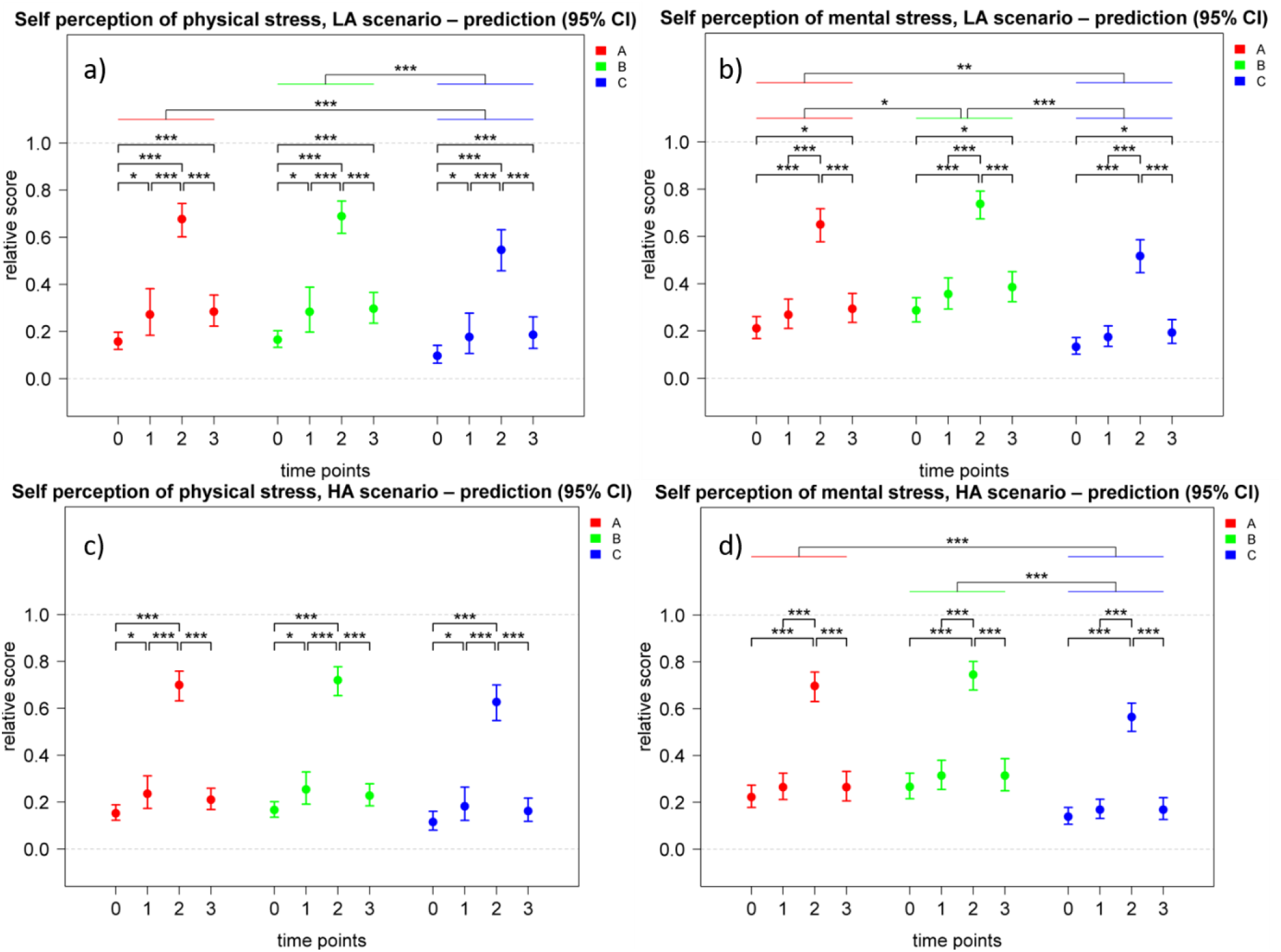
Development of self-perceived physical and mental stress in LA and HA scenarios. Participants were asked to rate their personal, physical, and mental stress levels at four time points. 0 – before MRI (day without scenario). 1 – before scenario. 2 – after scenario. 3 – immediately before debriefing. The subjective rating of physical and mental stress levels were linearly transformed to scale 0-1. Group A consisted of 20 healthy males without previous experience in martial arts, combat sports, or self-defence training. Group B included 20 males trained in martial arts, combat sports, and personal self-defence, but without real-life experience with violence. Finally, group C consisted of 20 members of the Rapid Response Unit of the Police of the Czech Republic from the South Moravian headquarters. These males were both trained and had experience with real-life violence.

As a freeze-like response predictors, the self-perception of physical and mental stress was of limited usefulness. In all groups merged together, no significant correlation with freeze-like response was established and only physical stress after the LA and mental stress after the HA scenario showed a weak but significant negative correlation with freeze-like response scores in LA scenarios (r = -0.30, p = 0.019; r = -0.28, p = 0.032). In the untrained and inexperienced group A, no significant correlation was observed between stress perception and freeze-like response in the LA scenario. In the HA scenario, group A showed a significant correlation with both mental and physical subjective stress before the scenario and freeze-like score (r = 0.45, p = 0.049; r = 0.53, p = 0.017, for physical and mental stress, respectively), meaning that the more stressed the inexperienced participants felt, the less they were freezing in the HA scenario. In the trained group B, the perception of physical and mental stress negatively correlated with the freeze-like response score in the LA scenario (r = -0.51, p = 0.021; r= -0.56, p = 0.011; respectively); no significant correlation was established for the HA scenario. No significant correlation was observed between freeze-like response and the subjective perception of stress in either scenario for the trained and experienced group C.

### Electrodermal activity

In addition to the subjective stress perception and biochemical markers of stress levels, we also measured heart-rate variability (HRV) and electrodermal activity (EDA) as physiological markers of stress^23,24^. Unfortunately, HRV data were full of artifacts and thus unsuitable for further analysis (Supplementary Data 5), likely due to the dynamic and contact nature of the employed scenarios. Fortunately, EDA data fared much better, with 47 LA and 49 HA records available for analysis.

Results of marginal models show that in the LA scenario (Supplementary Data 6) all studied EDA-related variables were significantly affected by predetermined time points in the scenarios, thus suggesting that the skin conductivity of participants reacted to events in scenarios. In addition, almost every variable also had a different time development in different groups (group : timepoint), showing that, as expected, the skin conductivity of trained and untrained individuals differs. Results of marginal models in the HA scenario (Supplementary Data 7) show that all studied EDA related variables except the response latency of the first significant SCR wrw (CDA.Latency) were significantly affected by predetermined time points in the scenarios, suggesting that the skin conductivity of participants reacted to events in scenarios. In addition, mean tonic activity (CDA.tonic) significantly differed between groups. Furthermore, the area (i.e. time integral) of the phasic driver wrw (CDA.ISCR) and the sum of SCR-amplitudes of significant SCR wrw (reconvolved with corresponding phasic driver peaks) (CDA.AmpSum) had different time development in different groups (group: timepoint).

Interestingly, EDA correlations with freeze-like response differed between groups as well as between scenarios. In the LA scenario, CDA.tonic upon verbal contact with the attacker negatively correlated with freeze-like response score in group A (r = -0.53, p = 0.034). In addition, CDA.tonic in group A also showed non-significant but suggestive negative correlations upon scenario start (−0.46, p = 0.074) and visual contact with the attacker (−0.49, p = 0.054). On the other hand, groups B and C did not indicate a correlation between CDA.tonic and freeze-like response. Still, instead, both showed negative correlations between non-specific responses in EDA (CDA.nSCR) and freeze-like response score. However, the timing of these correlations was different. In group B, CDA.nSCR negatively correlated with the freeze-like score upon visual contact with the attacker (r = -0.53, p = 0.034), while the negative correlation between CDA.nSCR became significant only after verbal contact (r = -0.56, p = 0.028).

No significant correlations were observed between EDA and freeze-like responses in any of the groups in the HA scenario. This finding may have resulted from the faster pace of the scenario as all significant correlations during the LA scenario were in time points corresponding to visual and verbal contact with the attacker and both of these parts happened much faster in the HA scenario (see descriptive statistics in Supplementary Data 6 and 7). However, a non-significant but suggestive pattern of correlation with CDC.tonic and freeze-like response scores was present in group B across all parts of the scenario, i.e. after the start of the measurement (r = -0.50, p = 0.096), at the start of the scenario (r = -0.49, p = 0.104), on visual contact (r = -0.49, p = 0.104), on verbal contact (r = -0.49, p = 0.104), on physical contact (r = -0.56, p = 0.061), and at the end of the scenario (r = -0.57, p = 0.055).

### Brain activity during the fMRI task correlates with freeze-like response

Changes in blood oxygenation level-dependent (BOLD) signal in all contrasts during the fMRI task where participants viewed neutral or negative IAPS images were associated with freeze-like response with corelaction distinctly different between groups an almost non-existent in merged sample. Since the contrast regulate > negative reflects the ability to regulate response to negative affective stimuli, we report it as the most relevant predictor of one’s ability not to freeze in threatening situations below, and provide results for the remaining contrasts in the Supplementary Data 3.

In untrained participants from group A, BOLD signal in all analyzed anatomical parts of the brain (hippocampus, parahippocampus, amygdala, thalamus, insula, cingulum) during the regulate > negative contrast negatively correlated with the subsequent score on the freeze-like response scale in the HA scenario (Figure 4). In other words, Higher BOLD response in the limbic system was associated with lower freeze-like score and thus more freezing behavior. However, there were no correlations between change in BOLD signals and a freeze-like response in the LA scenario. The brain region with the strongest association with freeze-like response was cingulum (r = -0.66 p = 0.002), suggesting that higher cingulum activation in untrained men results in more freezing.

**Figure 4.**
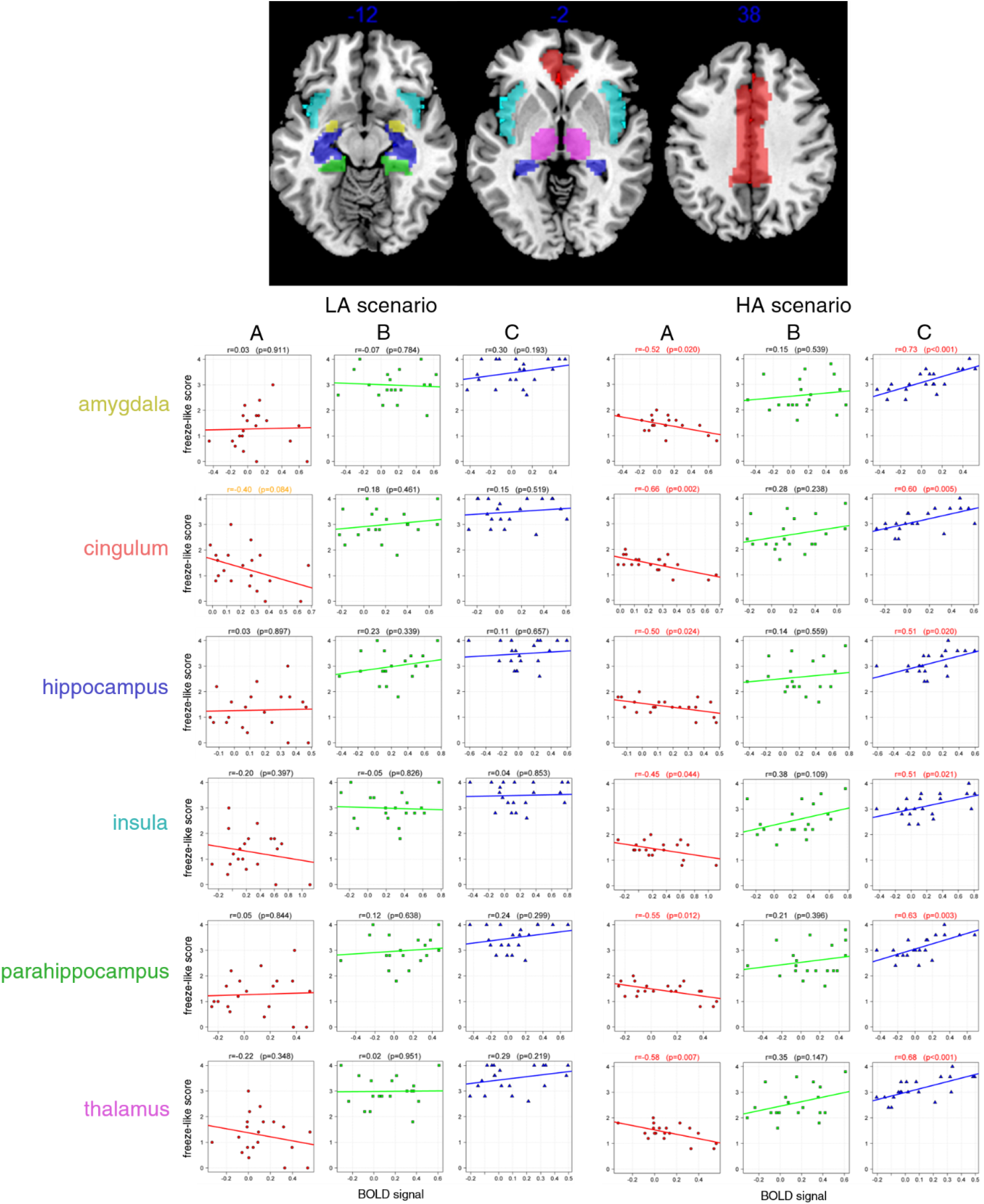
Brain activity correlates with freeze-like response in training and experience-dependent manner. Top part shows the AAL-based regions of interest on axial slices at z=-12, z=-2 and z=38. The graphs show Pearson’s correlations between the BOLD signal in the analyzed brain regions and subsequent freeze-like response in either LA or HA scenario. Significant correlations are highlighted in red and trends are highlighted in orange. Group A consisted of 20 healthy males without previous experience in martial arts, combat sports, or self-defence training. Group B included 20 males trained in martial arts, combat sports, and personal self-defence, but without real-life experience with violence. Finally, group C consisted of 20 members of the Rapid Response Unit of the Police of the Czech Republic from the South Moravian headquarters. These males were both trained and had experience with real-life violence.

The situation was dramatically different in trained group B. There were no significant correlations of BOLD signal in regulate > negative contrast with a freeze-like response in the HA or LA scenario (Figure 4).

The situation was again different in the trained and experienced group C, where BOLD activity in regulate > negative contrast in all analyzed brain regions correlated significantly with a higher freeze-like score (less freezing) in the HA scenario (Figure 4). From the analyzed brain regions, the BOLD signal in the amygdala showed the strongest correlation with the freeze-like response in the HA scenario (r = 0.73 with p <0.001). Notably, professional group C was the only one to show a significant correlation between BOLD signal and freeze-like response in the LA scenario. Specifically, increased thalamic activity showed a moderately strong (r = 0.45) and significant (p = 0.045) correlation with freeze-like response in the LA scenario.

When the BOLD activity was analysed for all groups combined, there was only one significant (p = 0.041) but very weak correlation (r = 0.27) between the BOLD signal in the insula during neutral-negative contrast and freeze-like response.

### Complex model

To assess the relative importance of the factors affecting the freeze-like response, we have decided to construct series of linear regression models. Because most of the measured variables were correlated, we have first used factor analysis (with varimax rotation) on four groups of analyzed variables (State-Trait Anxiety Inventory scores, cortisol and osteocalcin levels, subjective stress perception, BOLD activity in limbic system in regulate > negative contrast). The resulting factors were used in combination with anthropometric data (age, height, weight) to create linear regression models predicting freeze-like response in LA and HA scenarios (Supplementary data 8).

In the LA scenario, only results from State-Trait Anxiety Inventory questionnaires and age of participants significantly improved the regression model compared to its basic version containing just information about the group participants belonged to. Therefore, the freeze-like response in the LA scenario can be predicted from the subject’s group, score from the State-Trait Inventory questionnaires, and age (Table 2). (effects for Stai-Trait and age are specific for each group - due to significant interaction between these variables and group)

**Table 2.**
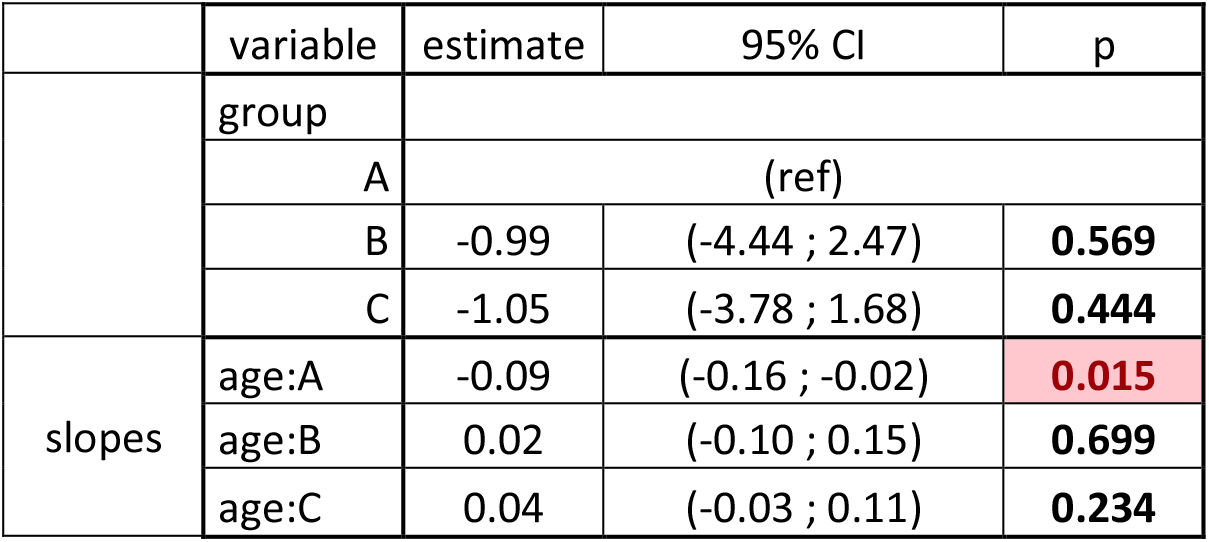
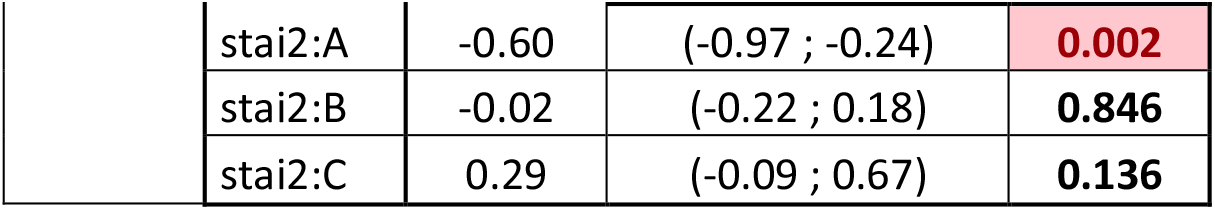
Best linear regression model for the LA scenario. Effects for Stai-Trait (Factor stai2) and age are specific for each group - due to significant interaction between these variables and group

|  | variable | estimate | 95% CI | p |
| --- | --- | --- | --- | --- |
|  | group |  |  |  |
|  | A | (ref) |  |  |
|  | B | -0.99 | (-4.44 ; 2.47) | **0.569** |
|  | C | -1.05 | (-3.78 ; 1.68) | **0.444** |
| slopes | age:A | -0.09 | (-0.16 ; -0.02) | **0.015** |
|  | age:B | 0.02 | (-0.10 ; 0.15) | **0.699** |
|  | age:C | 0.04 | (-0.03 ; 0.11) | **0.234** |
|  | stai2:A | -0.60 | (-0.97 ; -0.24) | **0.002** |
|  | stai2:B | -0.02 | (-0.22 ; 0.18) | **0.846** |
|  | stai2:C | 0.29 | (-0.09 ; 0.67) | **0.136** |

In the HA scenario, neither results from State-Trait Anxiety Inventory questionnaires or age of the participants significantly improved the basic model. However, the model was significantly improved by including information about the levels of cortisol and osteocalcin and information about the BOLD activity in the limbic system in regulate > negative contrast. Therefore, freeze-like response in the HA scenario can be predicted from the individual’s group, levels of cortisol and osteocalcin, and limbic system activity during the task in fMRI (Table 3).

**Table 3.** Best linear regression model for the HA scenario. Effects for cortisol and osteocalcin levels (factors ko.HA1-3) and limbic system activity (Factor AAL1) are specific for each group - due to significant interaction between these variables and group

|  | variable | estimate | 95% CI | p |
| --- | --- | --- | --- | --- |
|  | group |  |  |  |
|  | A | (ref) |  |  |
|  | B | 1.18 | (0.90 ; 1.46) | **<0.001** |
|  | C | 1.54 | (1.26 ; 1.83) | **<0.001** |
| slopes | ko.HA1:A | 0.10 | (-0.10 ; 0.30) | **0.331** |
|  | ko.HA1:B | -0.20 | (-0.38 ; -0.02) | **0.030** |
|  | ko.HA1:C | -0.12 | (-0.36 ; 0.13) | **0.341** |
|  | ko.HA2:A | 0.13 | (-0.06 ; 0.32) | **0.182** |
|  | ko.HA2:B | 0.33 | (0.13 ; 0.52) | **0.001** |
|  | ko.HA2:C | 0.09 | (-0.05 ; 0.23) | **0.221** |
|  | ko.HA3:A | 0.05 | (-0.13 ; 0.22) | **0.595** |
|  | ko.HA3:B | 0.27 | (0.12 ; 0.43) | **0.001** |
|  | ko.HA3:C | -0.15 | (-0.35 ; 0.06) | **0.155** |
|  | AAL1:A | -0.18 | (-0.37 ; 0.02) | **0.072** |
|  | AAL1:B | 0.15 | (-0.02 ; 0.32) | **0.091** |
|  | AAL1:C | 0.22 | (0.07 ; 0.37) | **0.006** |

## Discussion

This is the first study to show that scenarios simulating armed physical attacks can elicit a freeze-like response in humans. The intensity and tendency to freeze are profoundly affected by personal training and experience. Interestingly, our HA scenario, which featured an element of surprise, elicited a freeze response even in highly trained and experienced professionals who performed without any signs of freezing in our more conventional LA scenario. We also show that feelings of anxiety assessed by STAI-S and cortisol and osteocalcin levels are at best of limited use in predicting freeze-like response in male subjects. On the other hand, we show that the subjective perception of stress and EDA correlates with freeze-like response. Nevertheless, these correlations are entirely dependent on the level of training and experience as well as on scenario intensity. Furthermore, our results suggest that BOLD signal levels in several brain regions while subjects observed neutral and negative images from the IAPS in fMRI predict freeze-like responses in subsequent scenarios. However, we also show that with respect to freezing tendency, the activity of these regions means very different things in untrained males, trained males, and males with both training and real-life experience with situations involving physical violence.

Our results show that while highly trained and experienced professionals perform perfectly in normal training routines (LA scenario), they may still freeze in scenarios featuring an element of surprise (HA scenario). This observation is in accordance with previous findings that the shooting accuracy of trained police officers changes from 90% in training to only 50% during real life-threatening situations.^35,36^ However, beyond merely demonstrating that even trained individuals experience freeze-like response, our results also show the tremendous effectiveness of training in preventing freeze response. In our study, training was by far the most important single factor affecting the likelihood and intensity of freezing. Even a highly trained professional sometimes experienced a freeze, but untrained individuals experienced it more often and more intensely. This also suggests that it may be possible to develop training strategies further reducing the likelihood and intensity of freeze response, e.g. by incorporating an element of surprise into training routines.

Our results also show that freezing, particularly in untrained individuals, may be so intense as to prevent any meaningful defence. Individuals with a score of 1 or lower on the freeze-like response scale reacted at best by taking a small step back and extending their hands slightly to an attack, which in real life would prove deadly in real life. This suggests that in some cases the absence of evidence of defensive behaviour in victims of violent crimes may result from freezing.

While the role of stress hormones in freeze response seems logical and the associations between freezing and basal and stress-induced cortisol levels were previously reported in primates and rodents,^21,22^ we found no connection between basal or stress-induced levels of cortisol or osteocalcin and freeze-like response. A possible interpretation might be that cortisol and osteocalcin levels in humans may correspond to the intensity of the reaction rather than to its type (fight, flight, freeze), which occurs depending on context. Our results agree with such interpretations as they show that persons experiencing the same perceived threat and showing the same overall response in terms of cortisol and osteocalcin levels react very differently based on previous training and experience.

An unexpected element in our study was the increased stress levels before scenarios in trained group B judged by the STAI-S scores and osteocalcin levels. This might be explained by the fact that, unlike participants from group A and C, participants from group B were enrolled in a study program taught by some of the researchers involved in the study. Therefore, they may have felt more pressured to perform well in the scenarios.

The neuroimaging part of this study showed that the activation of the same brain areas, while viewing the same IAPS pictures, has an entirely different connection to freeze response based on training and experience levels. While the activation of all studied parts of the limbic system (hippocampus, parahippocampus, amygdala, thalamus, insula, cingulum) in the regulate > negative contrast correlated strongly with more freezing in the untrained group A, the activation of the same parts of the brain in the trained and experienced group C correlated with less freezing. Interestingly, there were no correlations between brain activity in fMRI and freezing in group B – which had training but lacked real-world experience with violence. This may be due to the fact that the group B, is in terms of experience somewhere in the middle of groups A and C which show opposite trends. Additionaly group B also had the highest heterogeneity in freeze-like respone in the HA scenario (figure 1). There are undoubtedly many ways to interpret our findings that indicate that the activity of the same brain regions has a different effect based on an individual’s level of experience and training. Neverthless, given that all analysed brain regions are connected to reaction to threatening stimuli^37–43^ one plausible interpretation is that the activation of the limbic system while viewing images from IAPS reflects a higher tendency to recognize threats. Thus, in untrained individuals the activation of the limbic system correlates with a higher tendency to freeze in the face of physical violence as these untrained individuals do not have the resources to react differently. However, the higher propensity for limbic system activation leads to less freezing in trained individuals as the stronger recognition of the threat strengthens the behavioral patterns learned for such situations.

At first sight, it may seem that our results showing the tremendous importance of self-defense training in determining an individual’s propensity to freeze are in conflict with results of animal studies showing that freezing is not subject to habituation^7^. However, this is a false comparison as training in self-defense is very different from habituation. Habituation, is a decreased responsiveness to a stimulus with repeated presentation.^44^ Self-defense training is a process of active learning how to respond better when someone attacks you accompanied by training to increase physical fitness. Therefore, parallel to self-defense training in animal models is not a mouse that is confronted with a cat multiple times. The parallel of self-defense training in mice would be a mouse that learns how to effectively defend against a cat and significantly improves its strength and stamina in order to be able to do so in practice.

The main limitation of this study is that it was conducted solely on males; it is thus difficult to assess how well its findings might be applicable to females. Initially, we intended to continue the study by recruiting the same number and groups of female participants for altered scenarios (including female attackers). However, the ongoing COVID-19 pandemic delayed and complicated the recruitment to the point where we instead decided to publish the male half of the study. Nevertheless, even though we cannot be sure whether the frequency of freezing in a threatening situation differs between males and females, it seems unlikely that the frequency would be lower in females given that up to 70% of female rape victims report a significant feeling of immobility during sexual attacks.^45^

Overall, we have shown that training and experience are crucial in the context of human freeze-like response. Focusing entirely on biochemical, biological, or psychological factors or brain activity without taking training and experience into account can lead to a loss of most relevant information, making the results uninformative. Therefore, future research into human fight, flight, and freeze response should always consider the role of training and experience. In addition, by showing that freezing is common in untrained men and can also be induced in professionals by simple scenarios simulating verbal and physical attacks, our study suggests that it is reasonable to expect an absence of active defence in a significant percentage of victims of all types of crimes. This finding has a potentially far-reaching implication in various fields ranging from forensic medicine to policy-making.

## Supporting information

fMRI raw data

## Acknowledgments

The project was supported by the project MUNI/A/1403/2018 Evaluation of self-defence scenario training and stress tolerance among students of study programme SESB and ASESB by Masaryk University, Grant Agency of Masaryk University, Faculty of Sports Studies. CETOCOEN PLUS (CZ.02.1.01/0.0/0.0/15_003/0000469) project of the Ministry of Education, Youth and Sports of the Czech Republic. The project was also supported by CETOCOEN EXCELLENCE Teaming 2 project supported by Horizon2020 (857560) and the Ministry of Education, Youth and Sports of the Czech Republic (02.1.01/0.0/0.0/18_046/0015975) as well as by the RECETOX research infrastructure (LM2018121). We also acknowledge the MAFIL core facility supported by the Czech-BioImaging large RI project (LM2018129, funded by MEYS CR) for their support in obtaining the scientific data presented in this paper. In addition, V.H. was supported by the mathematical and statistical modelling grant of Masaryk University (MUNI/A/1615/2020).

## Author contributions

P.L. identified the research question, helped with the study design, wrote the first draft of the manuscript, and interpreted the results. M.V. designed and supervised the scenarios, including EDA and HRV measurements and methodology of freeze-like response evaluation, and recruited study participants. K.M. designed the MRI protocol, contributed to data analysis. J.N. helped with design scenarios, including EDA and HRV measurements, and methodology of freeze-like response evaluation, and recruited study participants. F.Z. analyzed the data. M.M. processed fMRI data, including statistical parametric mapping-based analysis and exported fMRI features for complex statistics. Z.R. lead the debriefing. M.B., J.Č., P.P., V.M., and Z.R., evaluated the freeze-like response from the video recordings. T.V. assessed survival scores using video recordings. J.T., J.K., J.F.K., M.T., performed the measurement of osteocalcin/cortisol. V.H. helped with data management and analysed HRV data, R.Š., L.R., M.G., contributed by data acquisition. M.B. provided feedback on MRI studies and critically revised the manuscript. J. B.V. supervised the workflow of the project and controlled and co-designed its critical steps.. All authors co-wrote the manuscript. P. L. and M.V. contributed to the manuscript in equal measure.

## Data availability

Anonymized data are either presented with the manuscript, or present in supplementary data further information is available upon reasonable request. Unfortunately, to protect the identity of the study participants, we are unable to share most video recordings. Illustrative recordings of the study scenarios were provided to the reviewers upon obtaining a written agreement from the affected participants.

