## Supplementary data 2 for "Biological Factors and Self-Perception of Stress Predict Human Freeze-Like Responses in the Context of Self-Defence Training and Personal Experience with Violence"

Selection of IAPS images

We chose images from the IAPS database based on their affective ratings (valence and arousal). The rating process of the database is described elsewhere (Bradley & Lang, 2007, pp. 30–35). In an entire IAPS dataset, the valence ranges from 1,31 to 8,34 and arousal from 1,72 to 7,35. For the stressful images, we chose images that were low on valence (mean = 2,90; SD = 0,61), high in arousal (mean = 6,19; SD = 0,62), and contained motives of physical violence, e.g., attacking animal, attacking human, knives, guns and others. For the neutral images, we chose those with medium values for arousal (mean = 5,04; SD = 1,02) and valence (mean = 4,58; SD = 0,76) which were resembling pictures chosen for stressful task, e.g. calm animals, people engaging in non-threatening activities, knives used in non-threatening context.

Participants were given instructions to look at the pictures. While looking at subset of the stressful pictures (22) they were task to observe the pictures and do not regulate the elicited emotion. While looking at the other subset of stressful pictures (22), they were asked to regulate their emotions elicited by these pictures. In between arousing pictures, the neutral pictures (22 pictures) emerged. This subset did not require any regulation.

#### Selection of negative IAPS pictures for observe task

| | Negative<br>to observe | valence mean | valence SD | arousal mean | arousal SD |
|---|---|---|---|---|---|
| **animals** | 1302 | 4.21 | -1.78 | 6 | -1.87 |
| | 1300 | 3.55 | -1.78 | 6.79 | -1.84 |
| | 1932 | 3.85 | -2.11 | 6.47 | -2.2 |
| | 1220 | 3.47 | -1.82 | 5.57 | -2.34 |
| | 1052 | 3.5 | -1.87 | 6.52 | -2.23 |
| **guns** | 3500 | 2.21 | -1.34 | 6.99 | -2.19 |
| | 6250.1 | 2.63 | -1.74 | 6.92 | -1.92 |
| | 6211 | 3.62 | -2.07 | 5.9 | -2.22 |
| | 6560 | 2.16 | -1.41 | 6.53 | -2.42 |
| | 6230 | 2.37 | -1.57 | 7.35 | -2.01 |
| | 6242 | 2.69 | -1.59 | 5.43 | -2.36 |
| **physical violence** | 6360 | 2.23 | -1.73 | 6.33 | -2.51 |
| | 6315 | 2.31 | -1.69 | 6.38 | -2.39 |
| | 6836 | 3.46 | -1.61 | 5.47 | -1.91 |
| | 9426 | 3.08 | -1.51 | 5.28 | -2.02 |
| | 9594 | 3.76 | -1.7 | 5.17 | -2.17 |
| | 9419 | 2.55 | -1.33 | 5.19 | -2.08 |
| | 6312 | 2.48 | -1.52 | 6.37 | -2.3 |
| **knive** | 6300 | 2.59 | -1.66 | 6.61 | -1.97 |
| | 6350 | 1.9 | -1.29 | 7.29 | -1.87 |
| | 6510 | 2.46 | -1.58 | 6.96 | -2.09 |
| | 6550 | 2.73 | -2.38 | 7.09 | -1.98 |

| | | |
|---|---|---|
| **mean** | 2.900454545 | 6.300454545 |
| **SD** | 0.645477112 | 0.682891428 |

#### Selection of negative IAPS pictures for the regulate task

| | Negative<br><br>to regulate | valence mean | valence SD | arousal mean | arousal SD |
|---|---|---|---|---|---|
| **animals** | 1301 | 3.7 | -1.66 | 5.77 | -2.18 |
| | 1525 | 3.09 | -1.72 | 6.51 | -2.25 |
| | 1930 | 3.79 | -1.92 | 6.42 | -2.07 |
| | 1205 | 3.65 | -1.76 | 5.79 | -2.18 |
| | 1050 | 3.46 | -2.15 | 6.87 | -1.68 |
| **guns** | 3530 | 1.8 | -1.32 | 6.82 | -2.09 |
| | 6244 | 3.09 | -1.78 | 5.68 | -2.51 |
| | 2811 | 2.17 | -1.38 | 6.9 | -2.22 |
| | 6243 | 2.33 | -1.49 | 5.99 | -2.23 |
| | 6571 | 2.85 | -2.05 | 5.59 | -2.5 |
| **physical violence** | 6840 | 3.63 | -2.04 | 5.95 | -1.9 |
| | 9452 | 3.19 | -1.98 | 5.14 | -2.3 |
| | 9042 | 3.15 | -1.89 | 5.78 | -2.48 |
| | 9592 | 3.34 | -1.75 | 5.23 | -2.09 |
| | 2683 | 2.62 | -1.78 | 6.21 | -2.15 |
| | 6821 | 2.38 | -1.72 | 6.29 | -2.02 |
| | 8230 | 2.95 | -1.88 | 5.91 | -2.15 |
| | 9423 | 2.61 | -1.51 | 5.66 | -2.15 |
| | 9427 | 2.89 | -1.47 | 5.5 | -2.09 |
| **knive** | 4664.2 | 2.79 | -1.77 | 6.13 | -2.29 |
| | 6313 | 1.98 | -1.38 | 6.94 | -2.23 |
| | 6540 | 2.19 | -1.56 | 6.83 | -2.14 |

| | | | |
|---|---|---|---|
| **mean** | 2.893181818 | | 6.086818182 |
| **SD** | 0.568925513 | | 0.537345211 |
| **all_negative_mean** | 2.896818182 | | 6.193636364 |
| **all_negative_SD** | 0.607201313 | | 0.61011832 |

### Selection of neutral IAPS pictures

| Neutral | valence mean | valence SD | arousal mean | arousal SD |
|---|---|---|---|---|
| 2362 | 6.74 | -1.34 | 4.6 | -2.09 |
| 1500 | 7.24 | -1.88 | 4.12 | -2.5 |
| 1030 | 4.3 | -2.35 | 5.46 | -2.43 |
| 1240 | 4.22 | -1.94 | 4.92 | -2.17 |
| 6250.2 | 6.32 | -1.7 | 5.13 | -2.06 |
| 6800 | 4.01 | -2.06 | 4.87 | -2.57 |
| 2383 | 4.72 | -1.36 | 3.41 | -1.83 |
| 2780 | 4.77 | -1.76 | 4.86 | -2.05 |
| 2704 | 4.85 | -1.89 | 5.3 | -2.16 |
| 4000 | 4.82 | -1.66 | 3.97 | -2.15 |
| 3210 | 4.49 | -1.91 | 5.39 | -1.91 |
| 2690 | 4.78 | -1.43 | 4.02 | -2.07 |
| 4233 | 4.56 | -1.86 | 3.96 | -2.15 |
| 2700 | 3.19 | -1.56 | 4.77 | -1.97 |
| 1510 | 7.01 | -2.07 | 4.28 | -2.47 |
| 5622 | 6.33 | -1.78 | 5.34 | -1.96 |
| 1230 | 4.61 | -1.74 | 4.03 | -2.41 |
| 9913 | 4.38 | -1.89 | 4.42 | -2.14 |
| 8010 | 4.38 | -1.86 | 4.12 | -2.08 |
| 4664.1 | 5.63 | -2.61 | 6.63 | -1.87 |
| 9401 | 4.53 | -1.31 | 3.88 | -1.98 |
| 2396 | 4.91 | -1.05 | 3.34 | -1.83 |
| **mean** | 5.035909091 | | 4.582727273 | |
| **SD** | 1.025979174 | | 0.758402519 | |
