## Supplementary data 5 for "Biological Factors and Self-Perception of Stress Predict Human Freeze-Like Responses in the Context of Self-Defence Training and Personal Experience with Violence"

### Heart rate variability for low anxiety scenario

The file contains plots of time-series of heart rate and RR intervals in the low anxiety scenario. In each plot, the time series line consists of the black and red parts. The black parts denote valid data. The red parts denote automatically detected artefacts. To detect artifact, we used kernel smoothing with bandwidth 10 s and Gaussian kernel. In case the relative difference between the smooth line and raw data was larger than 20%, we called the raw data value an artifact. The vertical blue lines in each plot denotes events in the scenario.

| participant ID | identified problem with the recording |
|---|---|
| STR01 | - |
| STR02 | RR intervals recording failed after approximately 1.5 minutes |
| STR03 | - |
| STR04 | - |
| STR05 | - |
| STR06 | participant ID was not assigned |
| STR07 | - |
| STR08 | - |
| STR09 | - |
| STR10 | - |
| STR11 | - |
| STR12 | - |
| STR13 | - |
| STR14 | participant ID was not assigned |
| STR15 | - |
| STR16 | - |
| STR17 | - |
| STR18 | - |
| STR19 | - |
| STR20 | recording missing |
| STR21 | - |
| STR22 | - |
| STR23 | - |
| STR24 | - |
| STR25 | - |
| STR26 | - |
| STR27 | - |
| STR28 | - |
| STR29 | - |
| STR30 | - |
| STR31 | - |
| STR32 | - |
| STR33 | - |
| STR34 | - |
| STR35 | - |
| STR36 | - |
| STR37 | - |

| | |
|---|---|
| STR38 | RR intervals recording failed after approximately 2 minutes |
| STR39 | - |
| STR40 | - |
| STR41 | - |
| STR42 | - |
| STR43 | - |
| STR44 | - |
| STR45 | - |
| STR46 | - |
| STR47 | - |
| STR48 | - |
| STR49 | - |
| STR50 | - |
| STR51 | - |
| STR52 | - |
| STR53 | - |
| STR54 | - |
| STR55 | - |
| STR56 | - |
| STR57 | - |
| STR58 | - |
| STR59 | - |
| STR60 | - |
| STR61 | recording missing |
| STR62 | - |

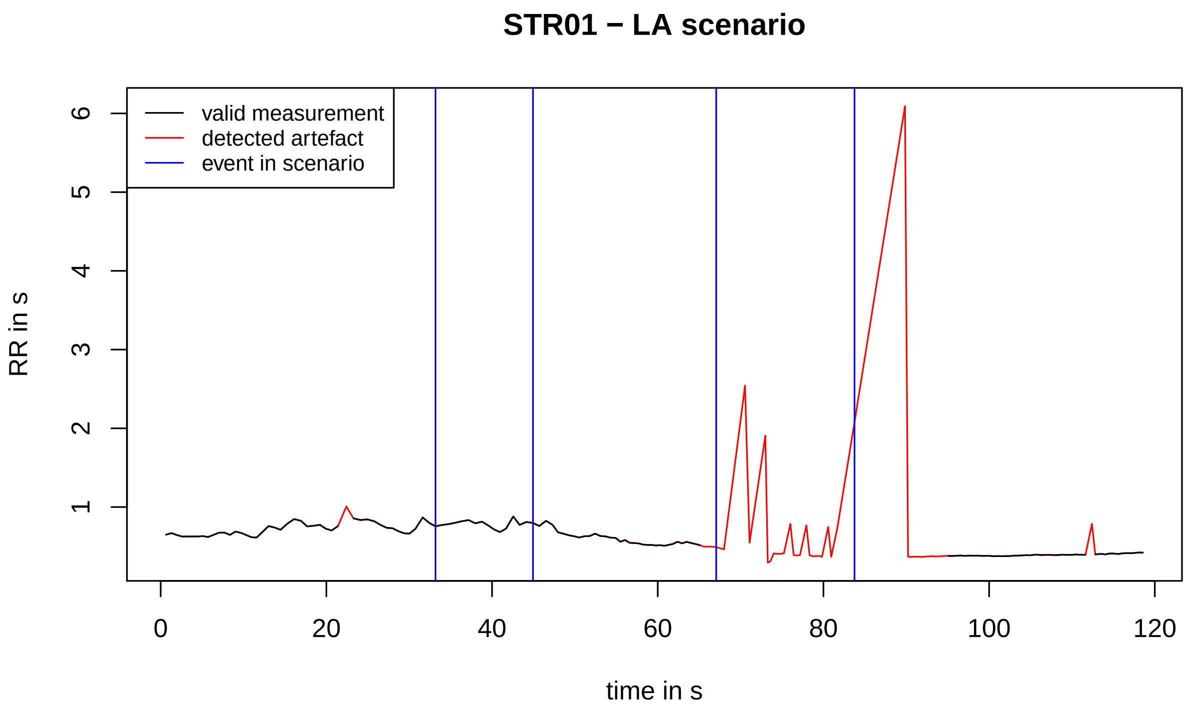
STR01 − LA scenario
valid measurement
detected artefact
event in scenario
RR in s
time in s
0
20
40
60
80
100
120
1
2
3
4
5
6

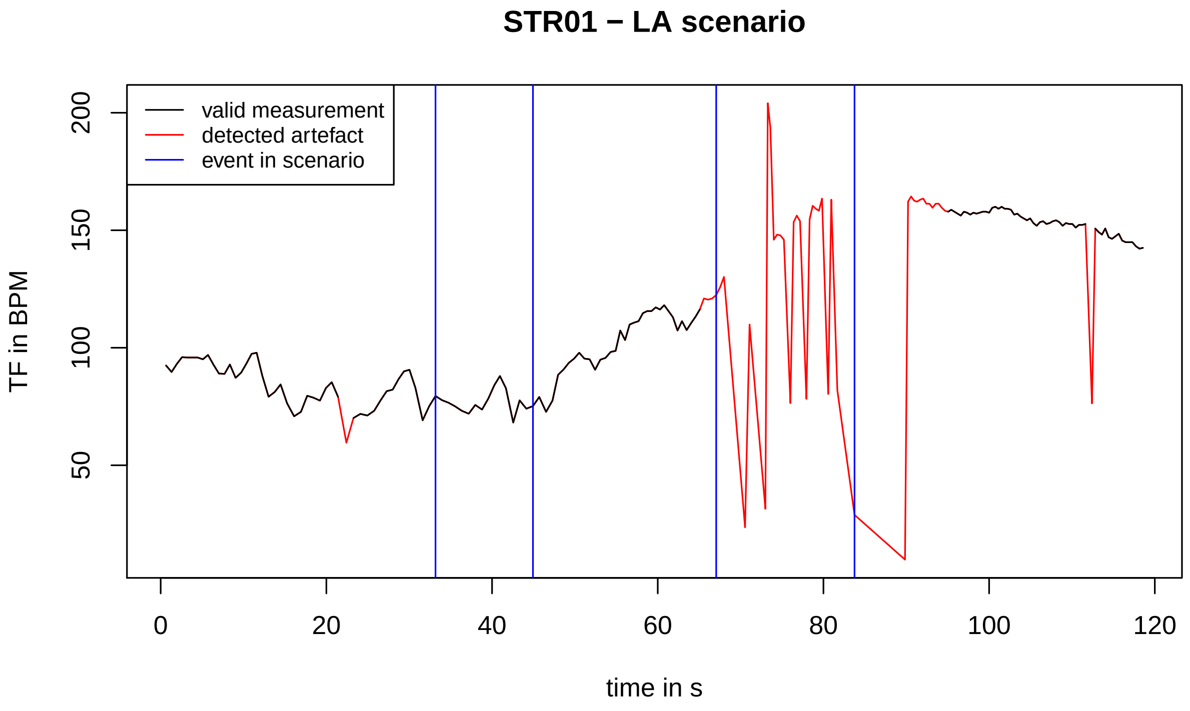
STR01 − LA scenario
valid measurement
detected artefact
event in scenario
TF in BPM
time in s
0
20
40
60
80
100
120
50
100
150
200

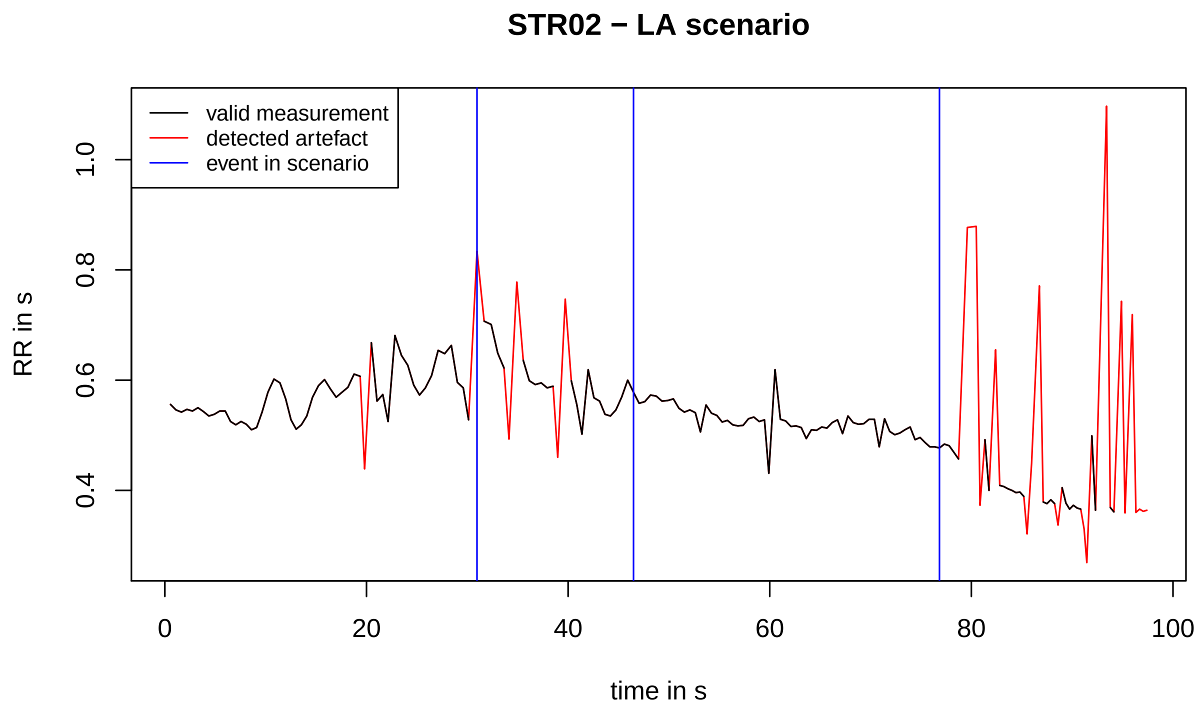
STR02 − LA scenario
valid measurement
detected artefact
event in scenario
RR in s
0.4
0.6
0.8
1.0
0
20
40
60
80
100
time in s

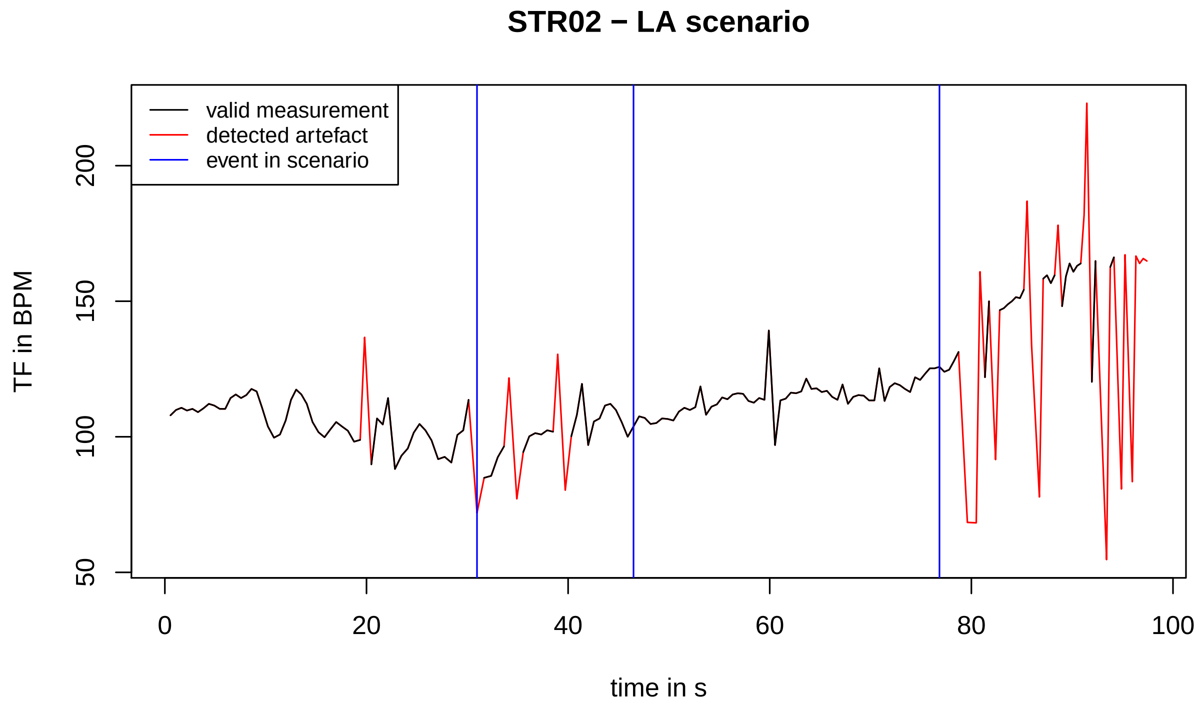
STR02 − LA scenario
valid measurement
detected artefact
event in scenario
TF in BPM
50
100
150
200
0
20
40
60
80
100
time in s

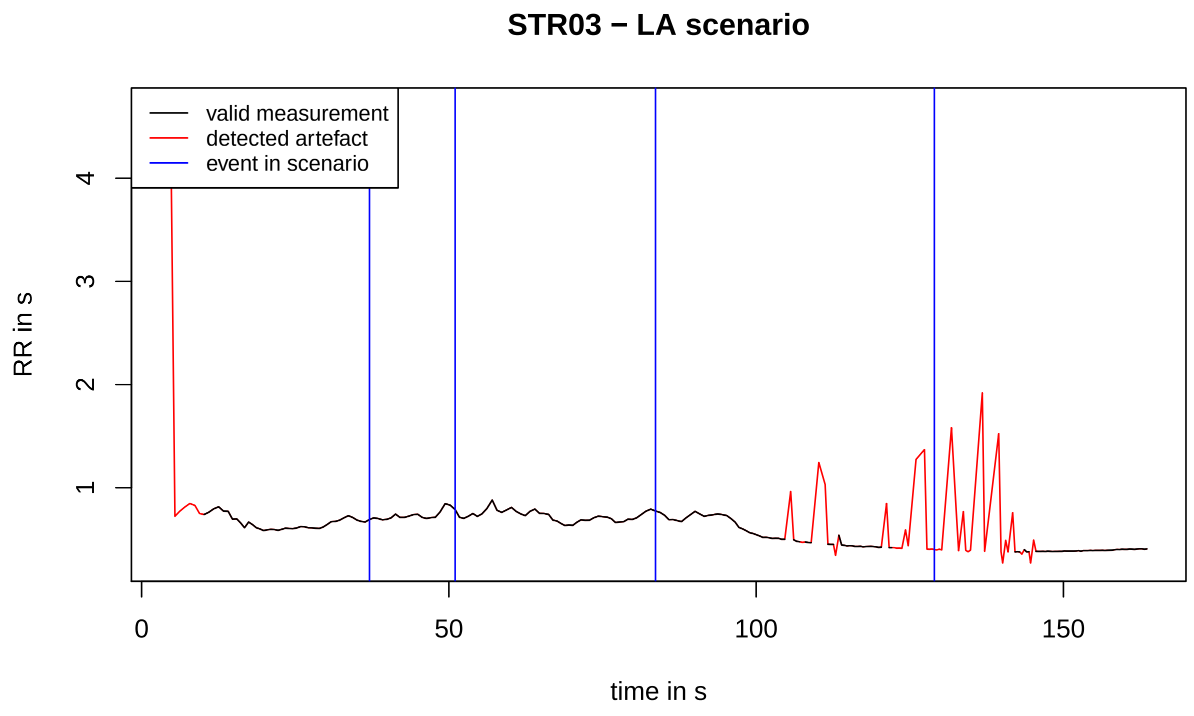
STR03 − LA scenario
valid measurement
detected artefact
event in scenario
RR in s
4
3
2
1
0
50
100
150
time in s

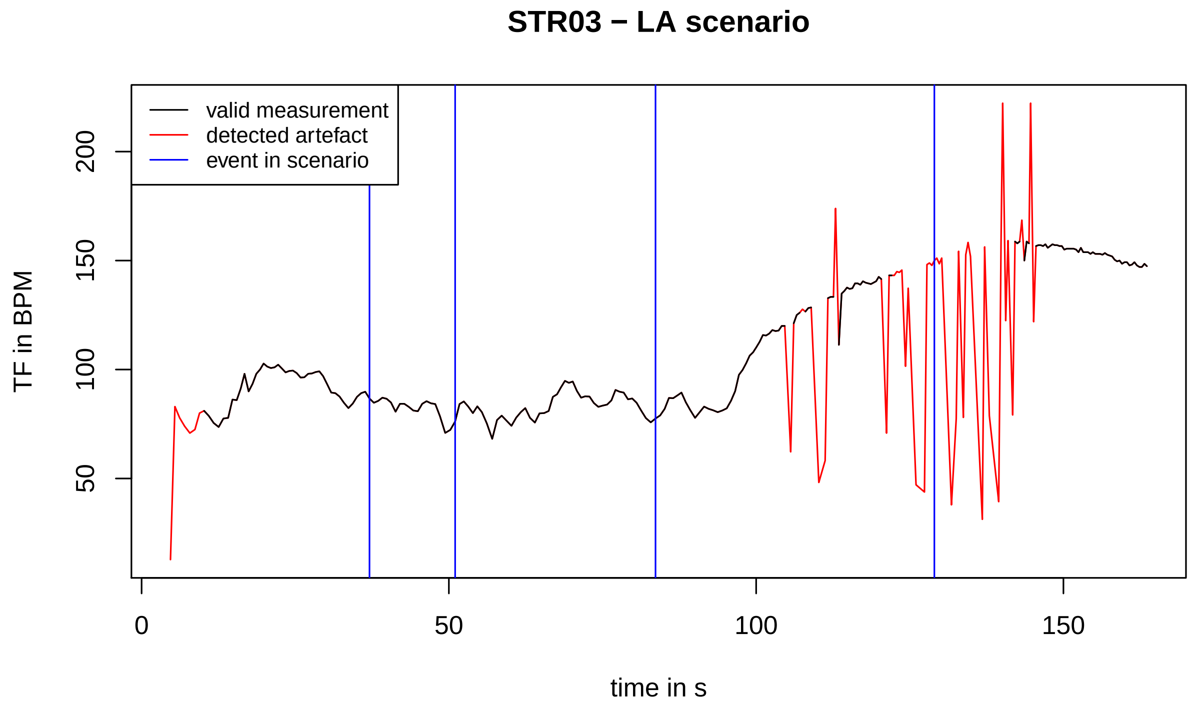
STR03 − LA scenario
valid measurement
detected artefact
event in scenario
TF in BPM
200
150
100
50
0
50
100
150
time in s

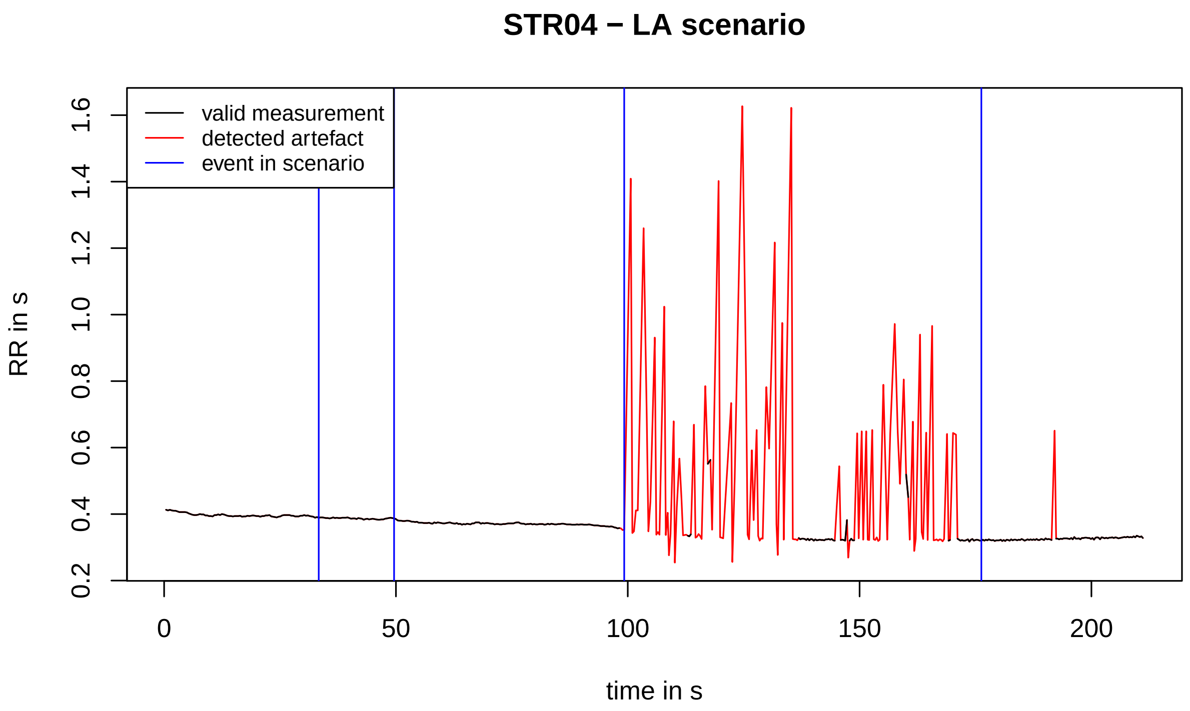
STR04 − LA scenario
valid measurement
detected artefact
event in scenario
RR in s
0.2
0.4
0.6
0.8
1.0
1.2
1.4
1.6
0
50
100
150
200
time in s

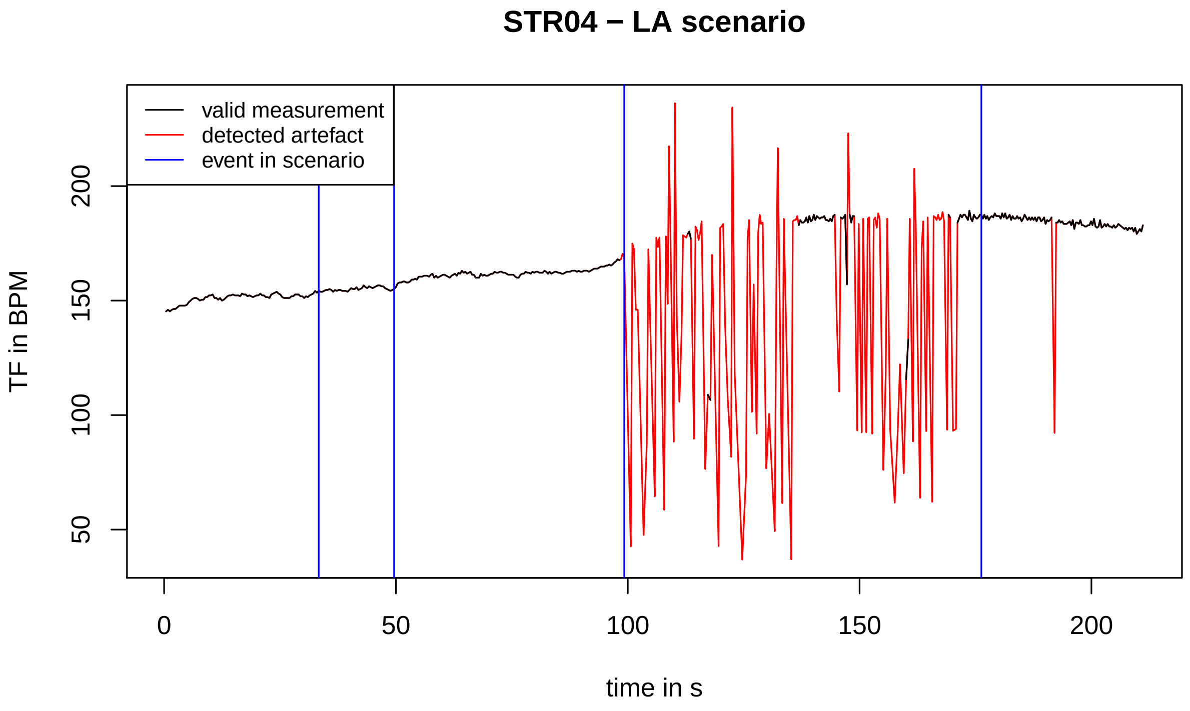
STR04 − LA scenario
valid measurement
detected artefact
event in scenario
TF in BPM
50
100
150
200
0
50
100
150
200
time in s

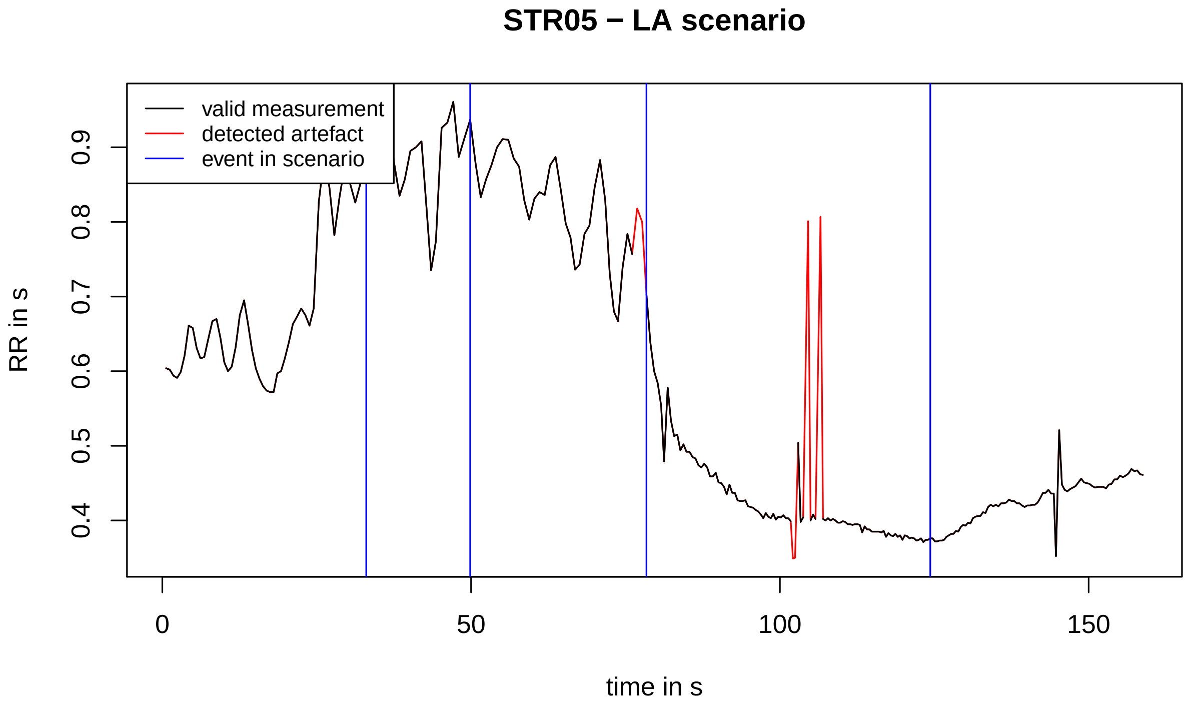
STR05 − LA scenario
valid measurement
detected artefact
event in scenario
RR in s
0.4
0.5
0.6
0.7
0.8
0.9
0
50
100
150
time in s

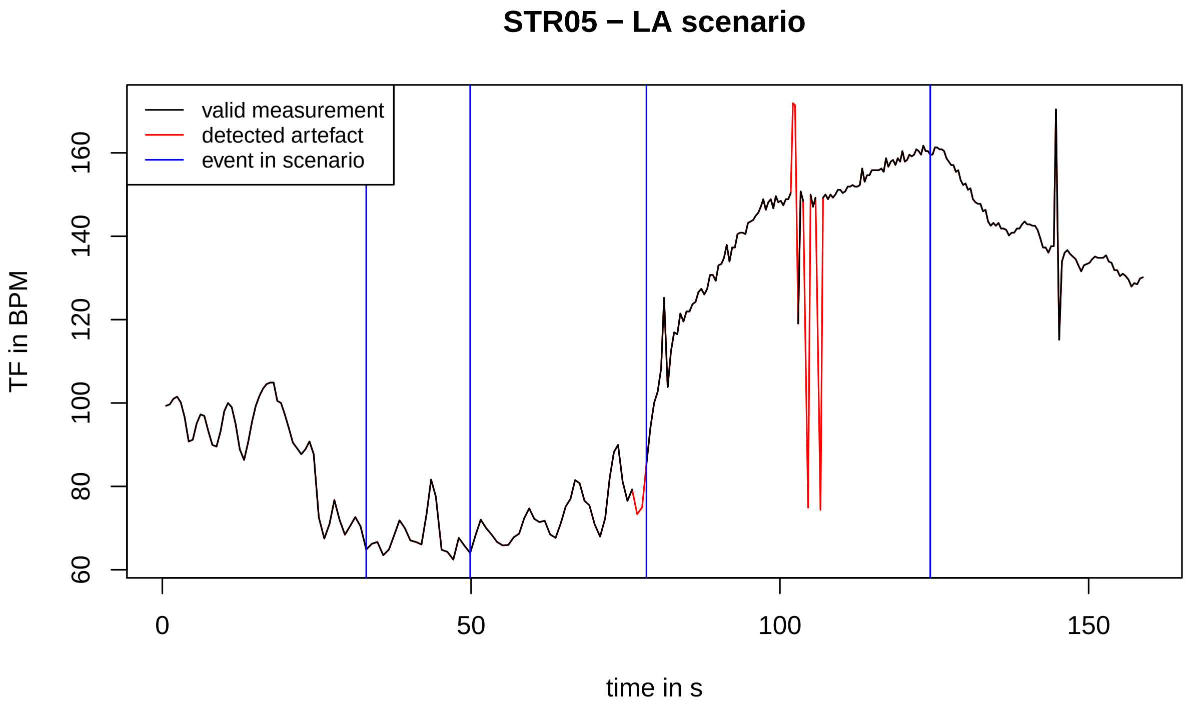
STR05 − LA scenario
valid measurement
detected artefact
event in scenario
TF in BPM
60
80
100
120
140
160
0
50
100
150
time in s

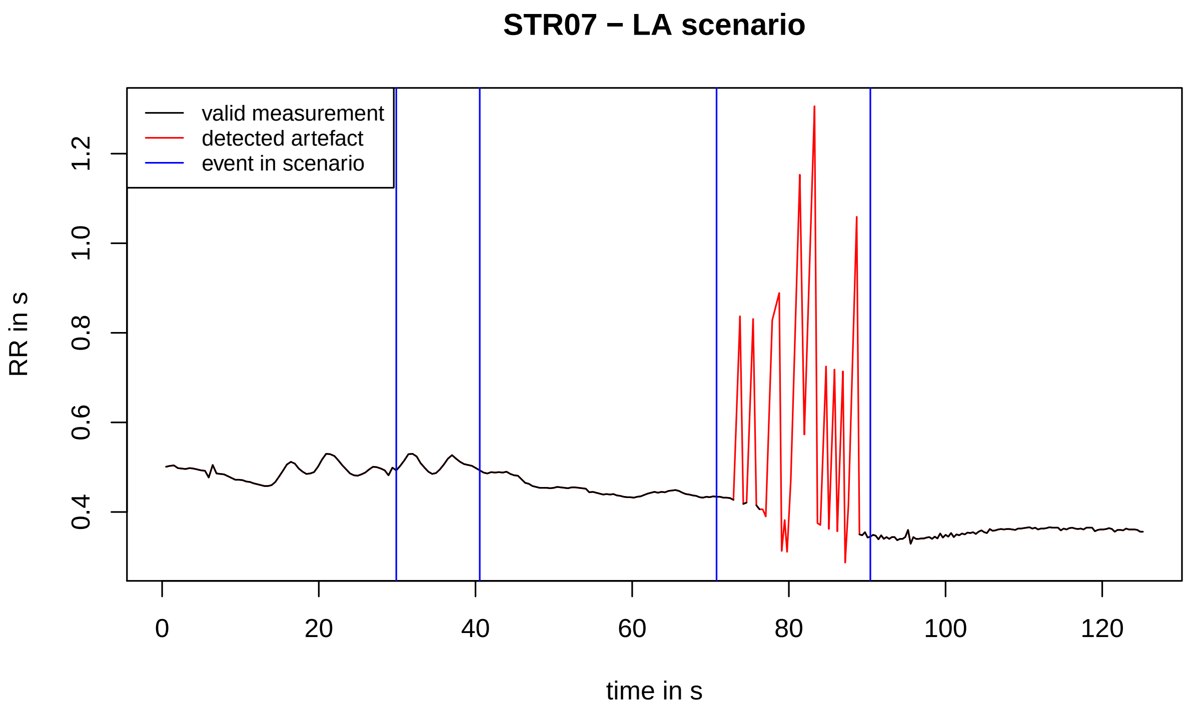
STR07 − LA scenario
valid measurement
detected artefact
event in scenario
RR in s
0.4
0.6
0.8
1.0
1.2
0
20
40
60
80
100
120
time in s

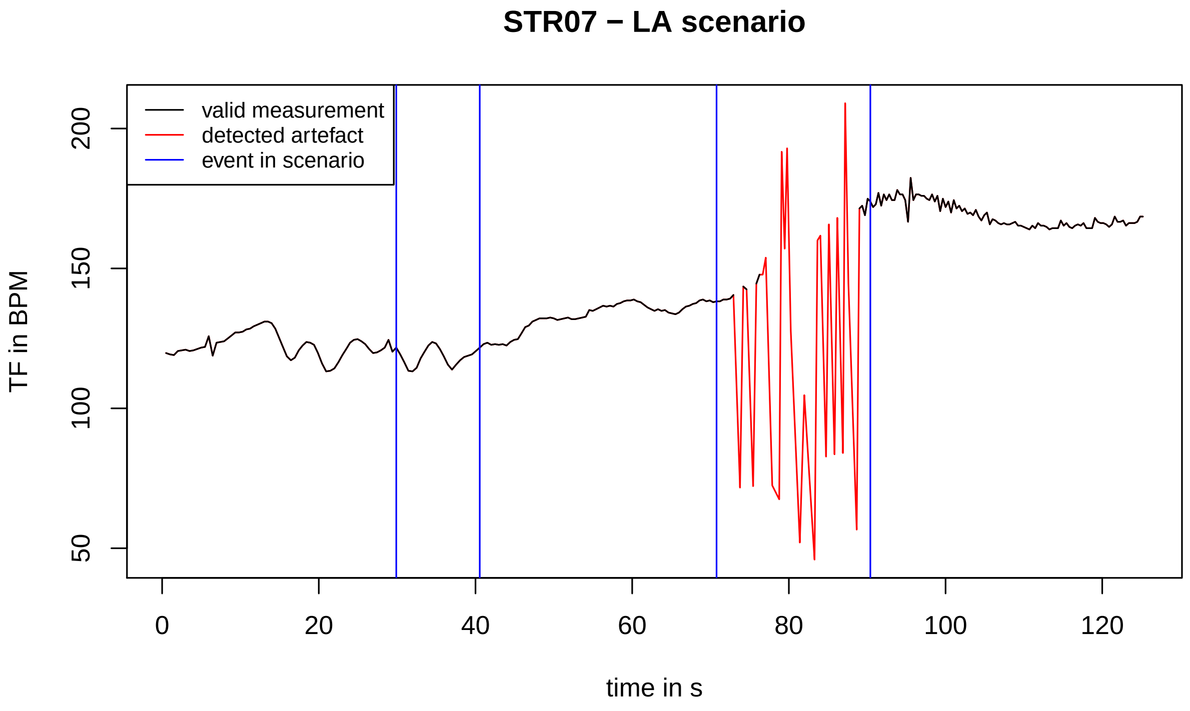
STR07 − LA scenario
valid measurement
detected artefact
event in scenario
TF in BPM
50
100
150
200
0
20
40
60
80
100
120
time in s

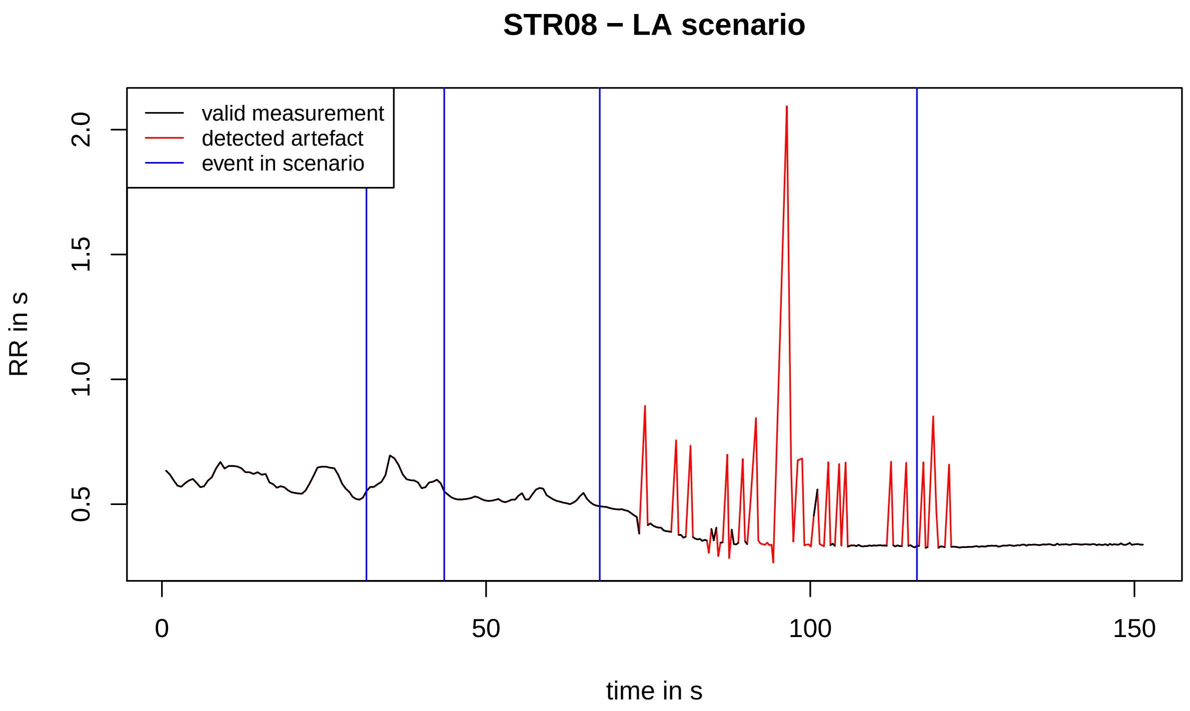
STR08 − LA scenario
valid measurement
detected artefact
event in scenario
RR in s
0.5
1.0
1.5
2.0
0
50
100
150
time in s

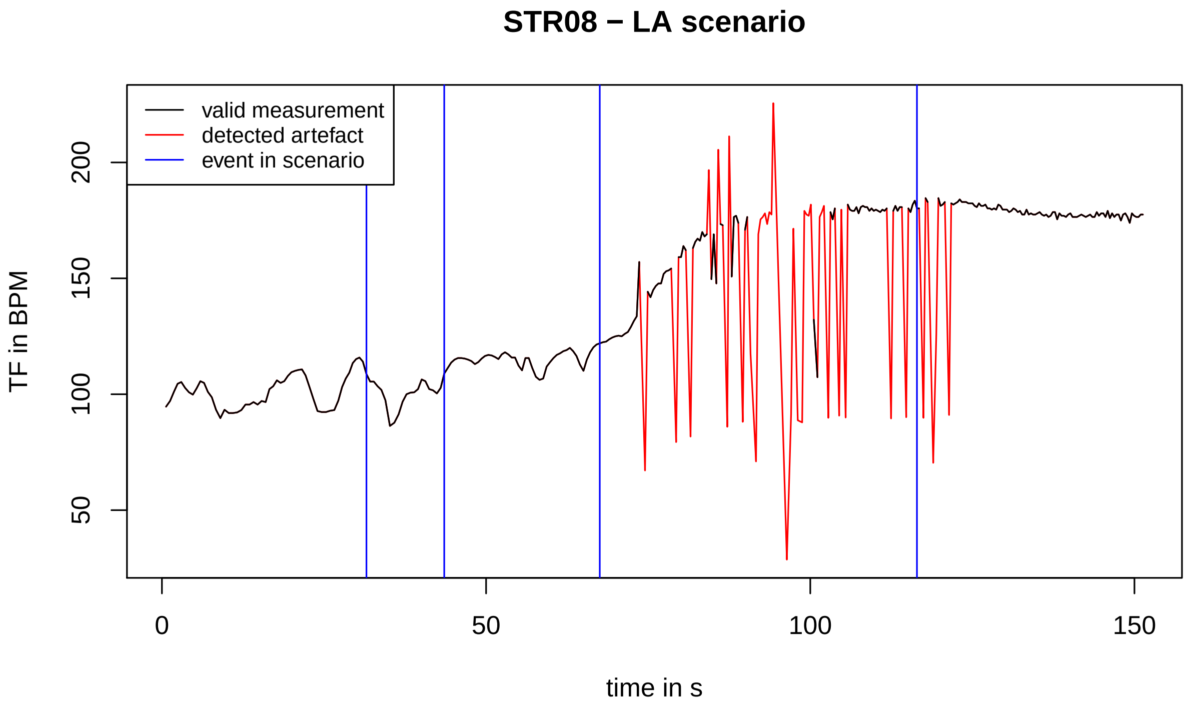
STR08 − LA scenario
valid measurement
detected artefact
event in scenario
TF in BPM
50
100
150
200
0
50
100
150
time in s

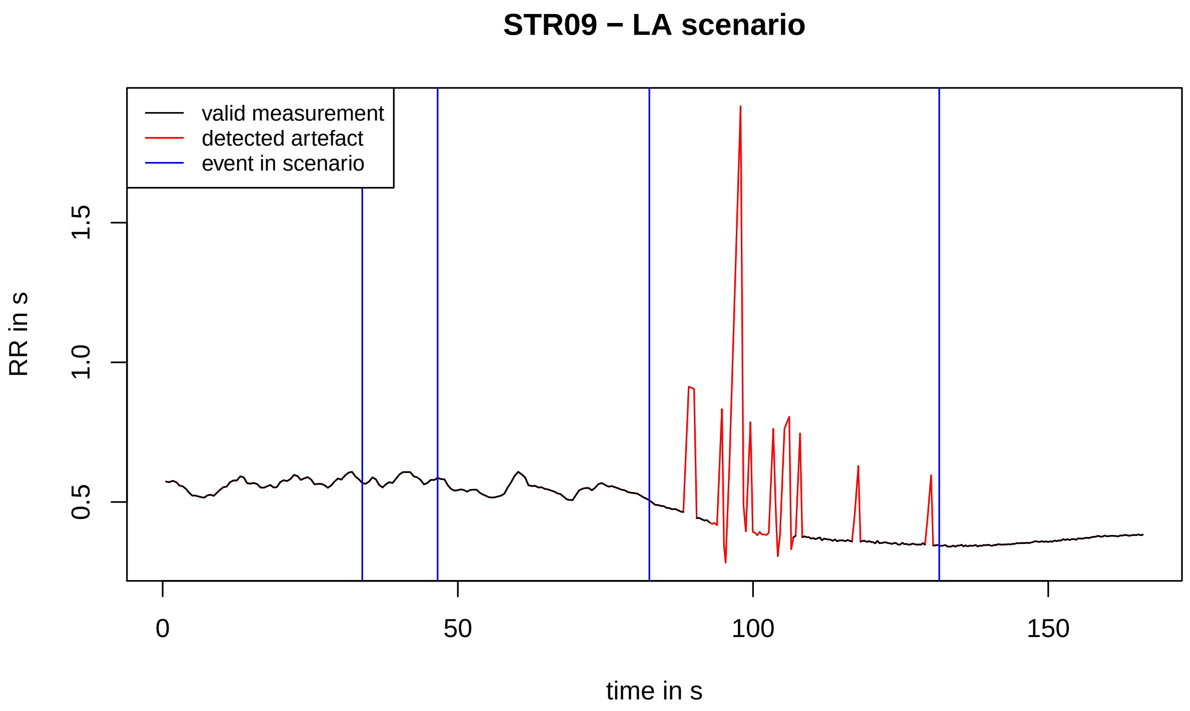
STR09 − LA scenario
valid measurement
detected artefact
event in scenario
RR in s
0.5
1.0
1.5
0
50
100
150
time in s

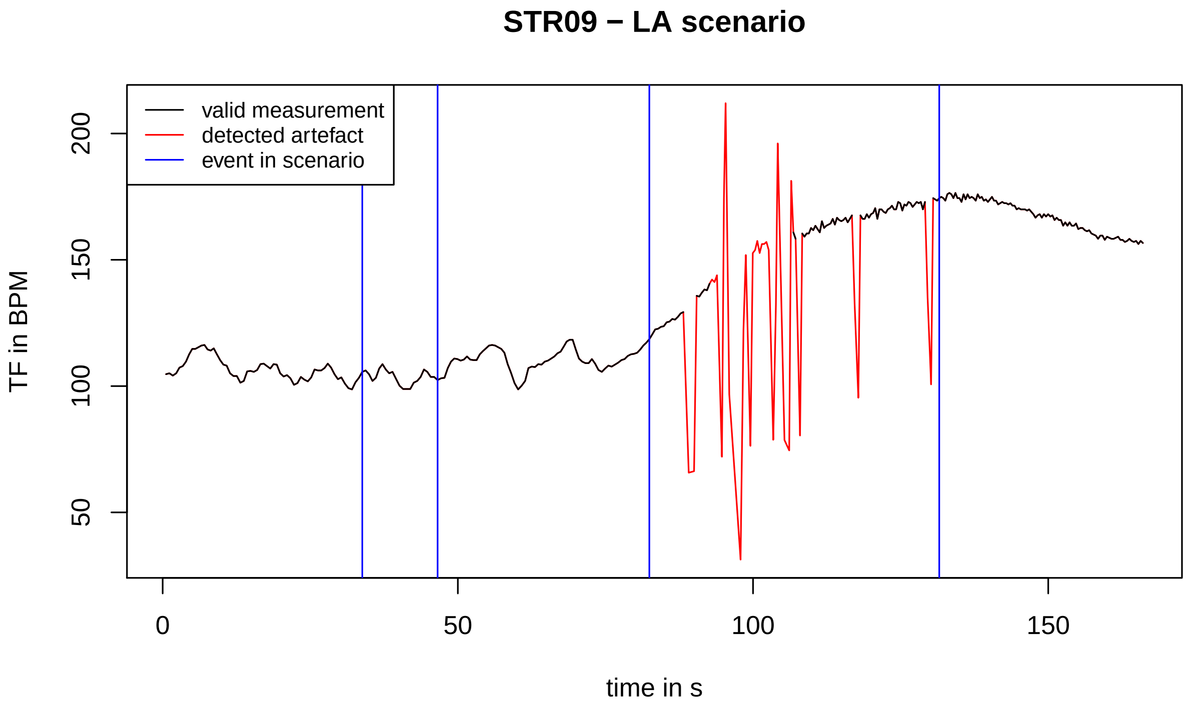
STR09 − LA scenario
valid measurement
detected artefact
event in scenario
TF in BPM
50
100
150
200
0
50
100
150
time in s

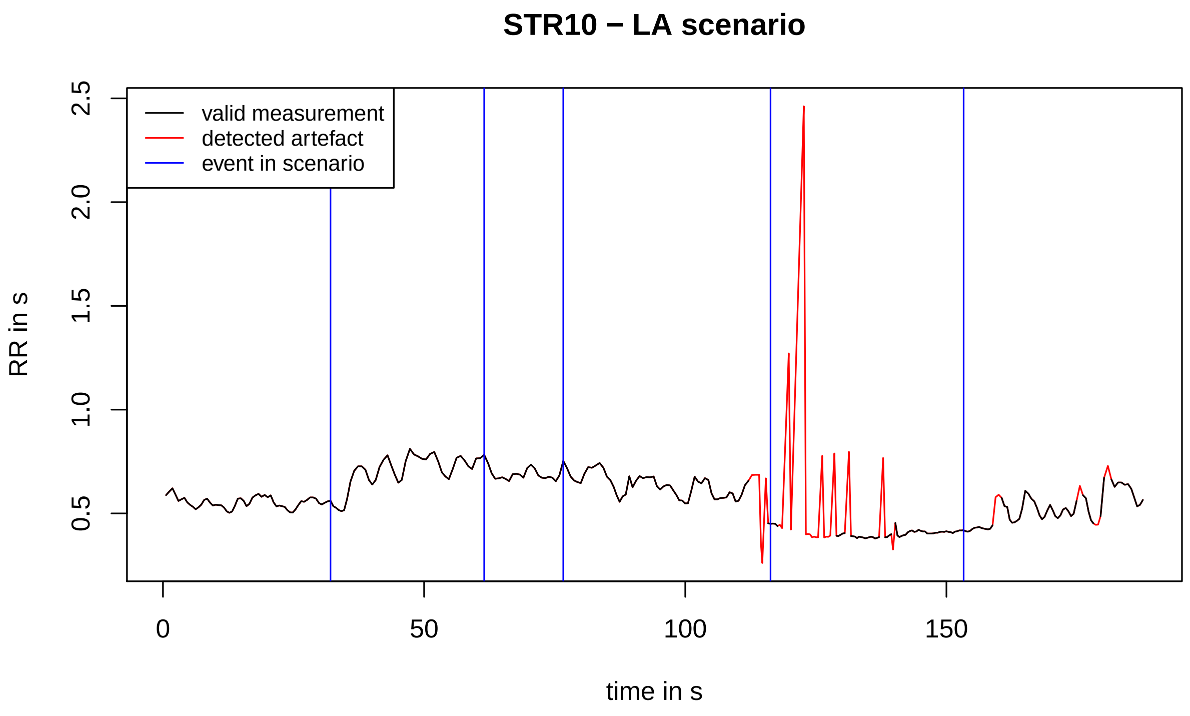
STR10 − LA scenario
valid measurement
detected artefact
event in scenario
RR in s
0.5
1.0
1.5
2.0
2.5
0
50
100
150
time in s

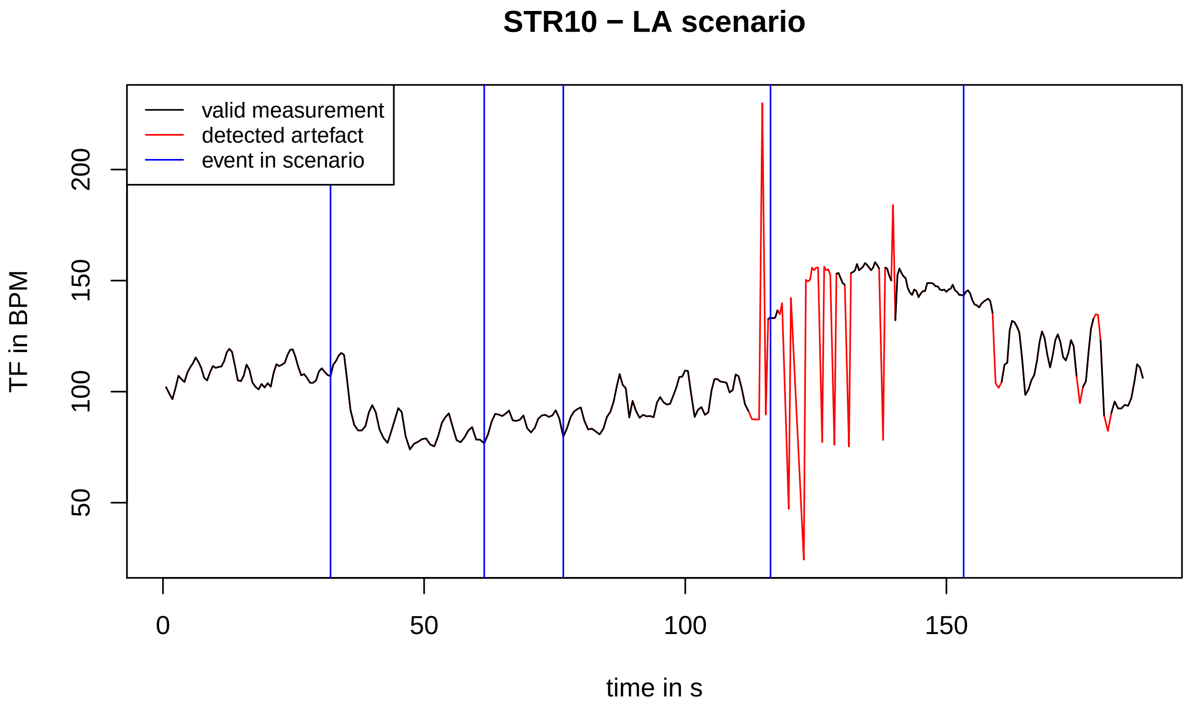
STR10 − LA scenario
valid measurement
detected artefact
event in scenario
TF in BPM
50
100
150
200
0
50
100
150
time in s

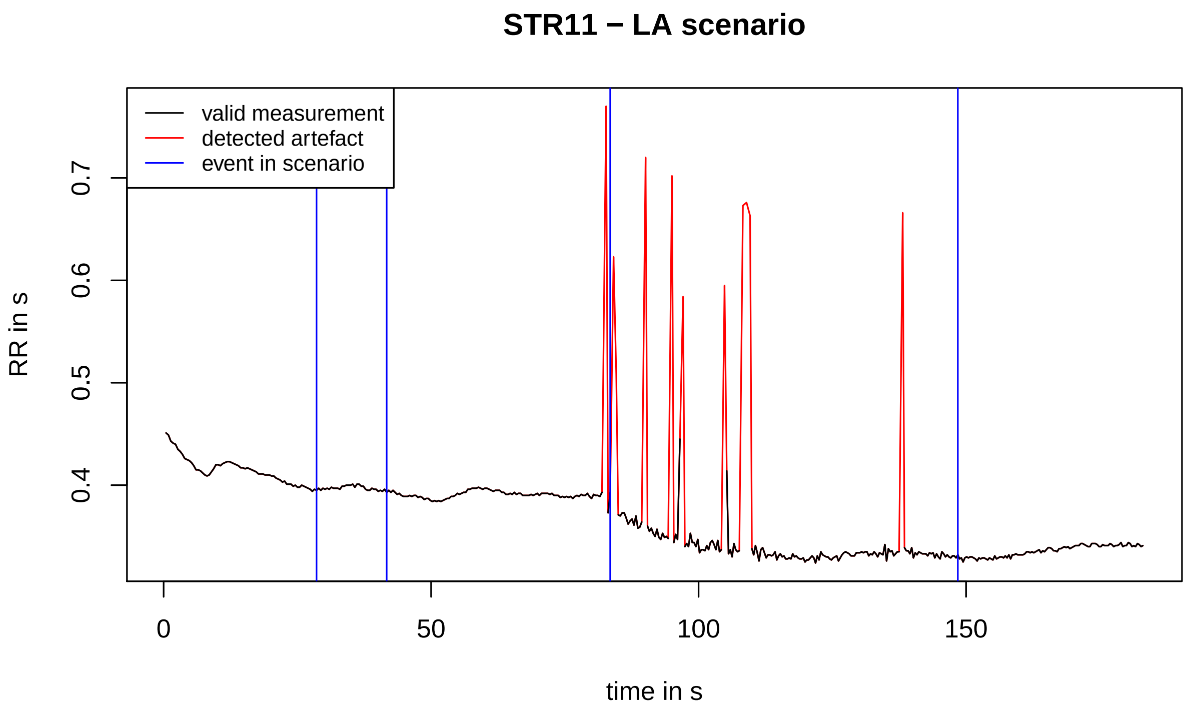
STR11 − LA scenario
valid measurement
detected artefact
event in scenario
RR in s
0.4
0.5
0.6
0.7
0
50
100
150
time in s

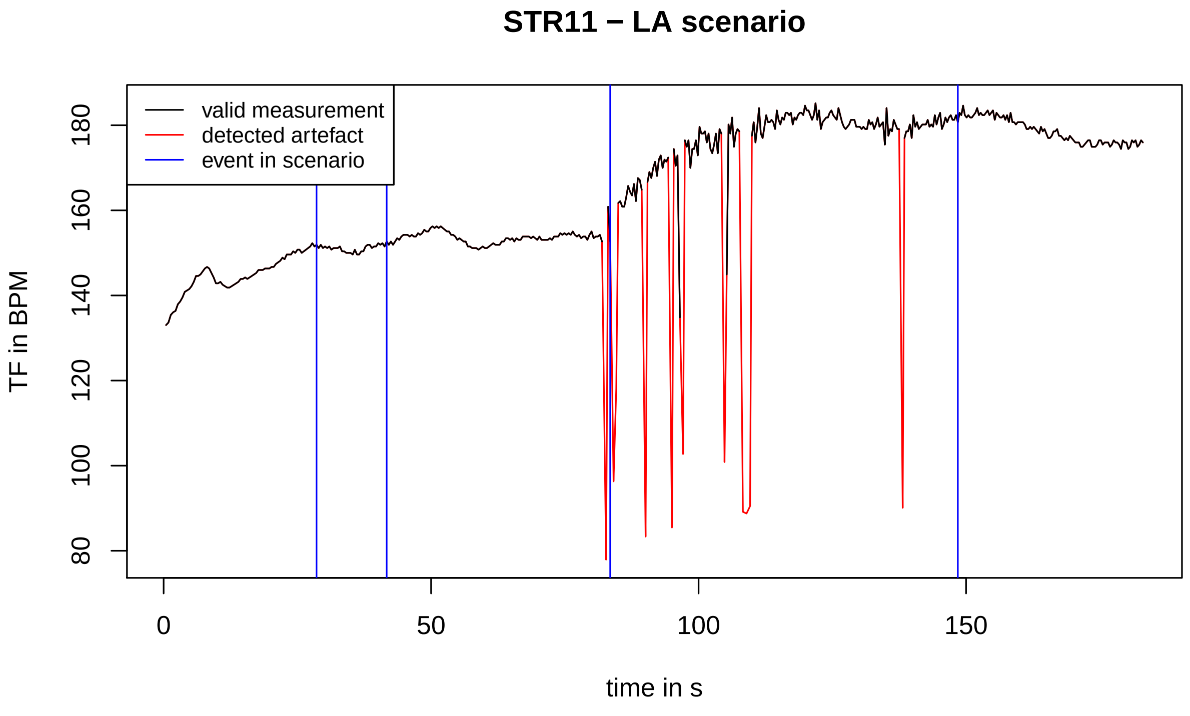
STR11 − LA scenario
valid measurement
detected artefact
event in scenario
TF in BPM
80
100
120
140
160
180
0
50
100
150
time in s

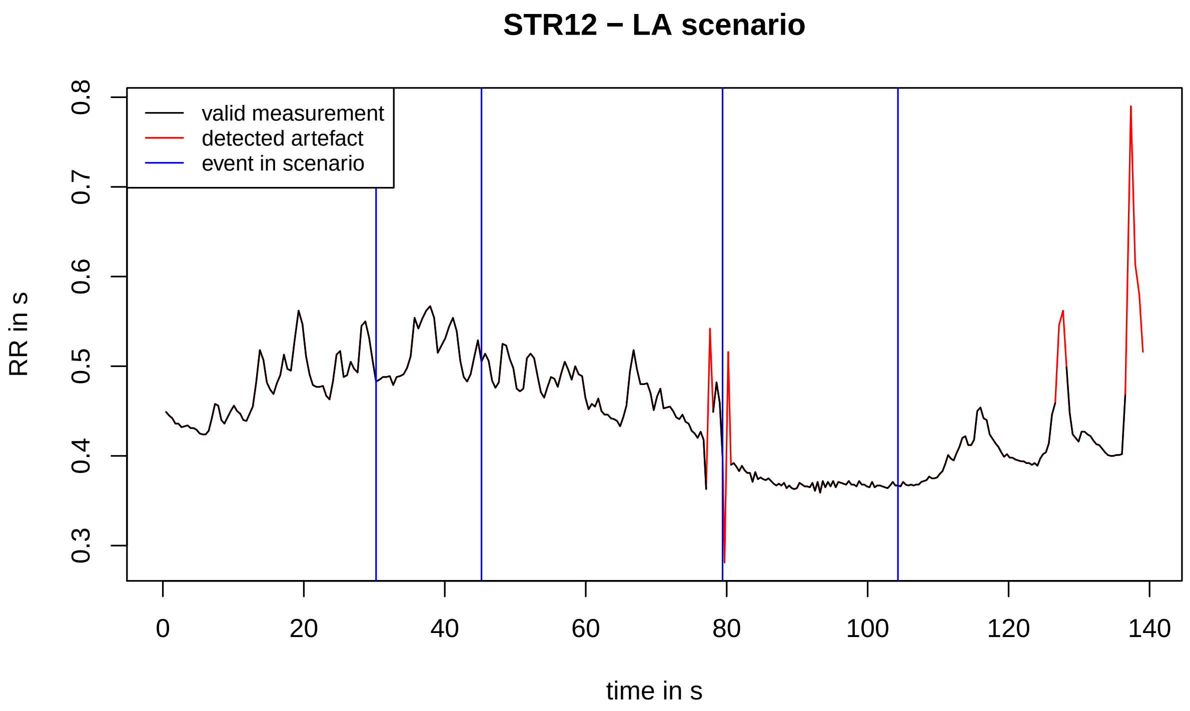
STR12 − LA scenario
valid measurement
detected artefact
event in scenario
RR in s
0.3
0.4
0.5
0.6
0.7
0.8
0
20
40
60
80
100
120
140
time in s

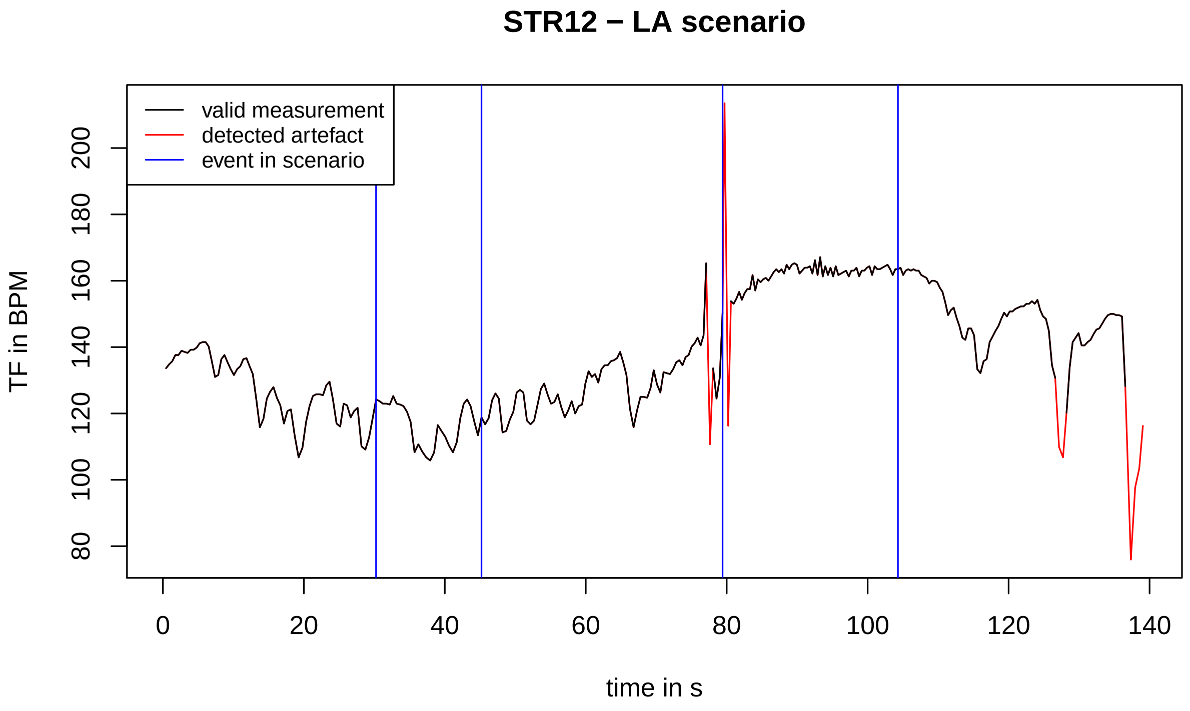
STR12 − LA scenario
valid measurement
detected artefact
event in scenario
TF in BPM
80
100
120
140
160
180
200
0
20
40
60
80
100
120
140
time in s

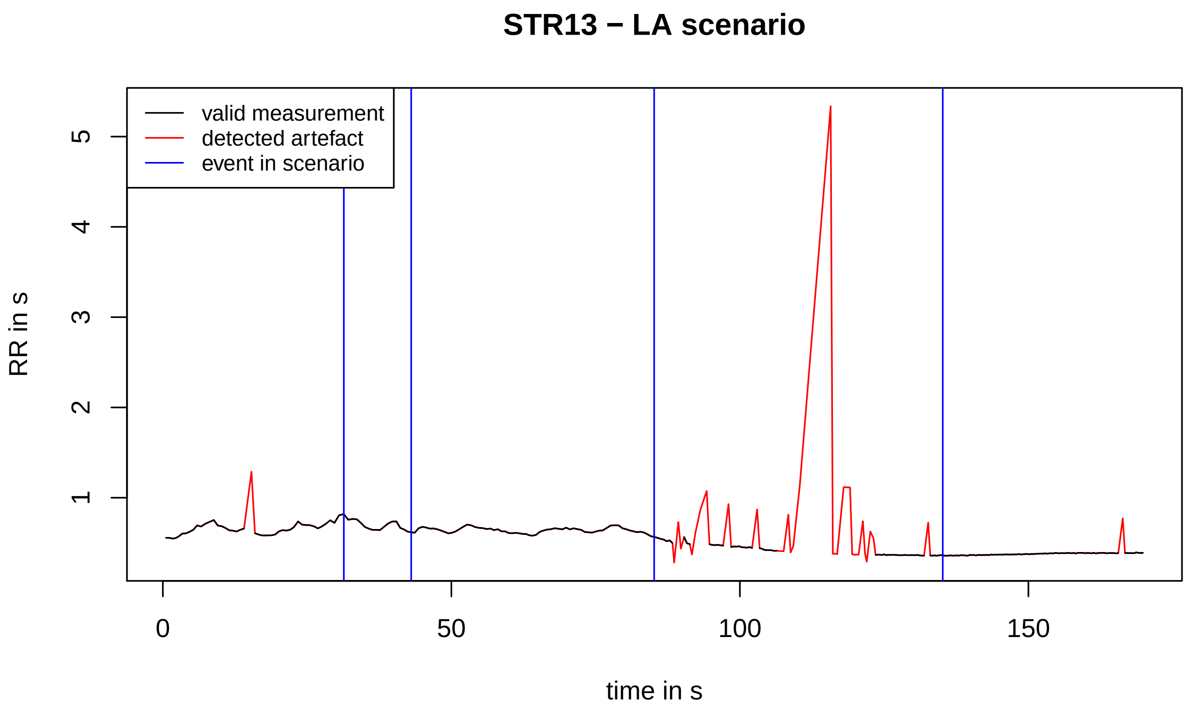
STR13 − LA scenario
valid measurement
detected artefact
event in scenario
RR in s
time in s
1
2
3
4
5
0
50
100
150

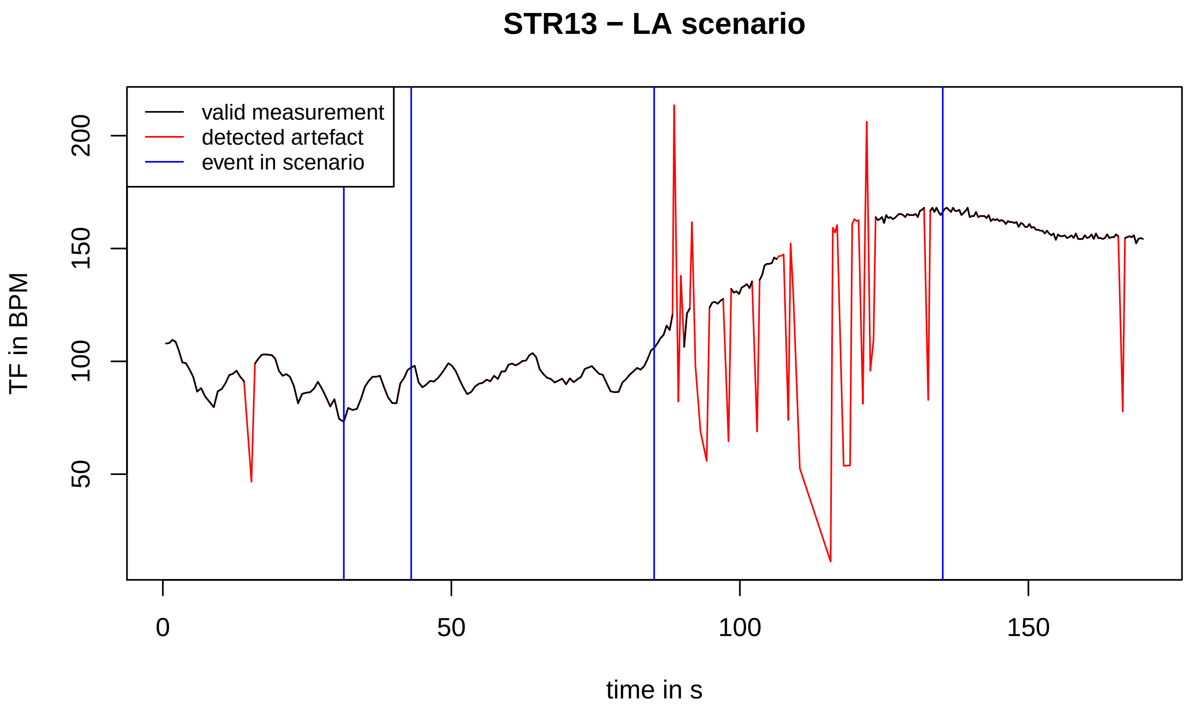
STR13 − LA scenario
valid measurement
detected artefact
event in scenario
TF in BPM
time in s
50
100
150
200
0
50
100
150

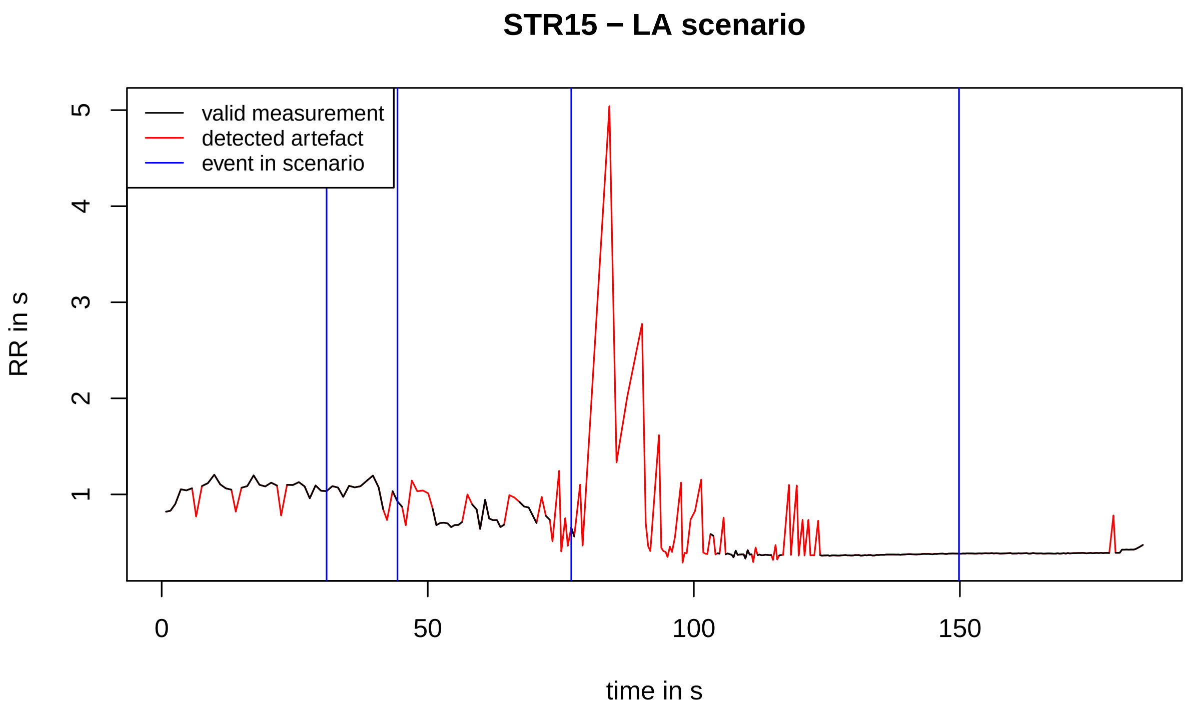
STR15 − LA scenario
valid measurement
detected artefact
event in scenario
RR in s
1
2
3
4
5
0
50
100
150
time in s

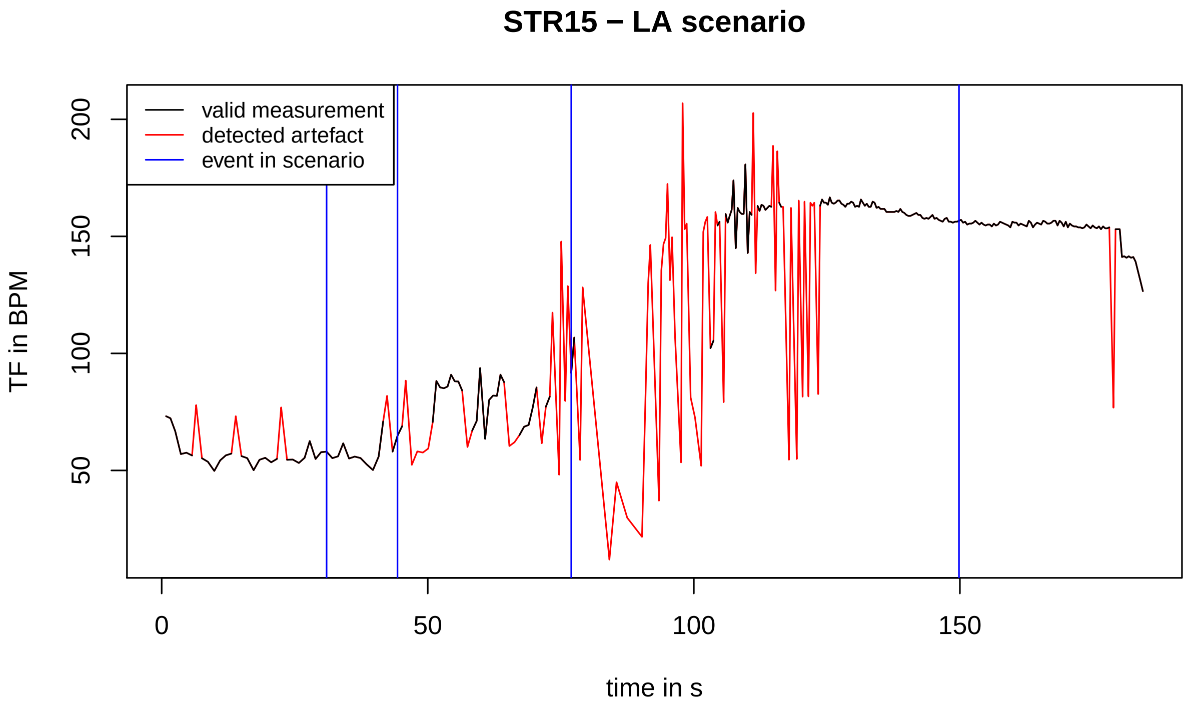
STR15 − LA scenario
valid measurement
detected artefact
event in scenario
TF in BPM
50
100
150
200
0
50
100
150
time in s

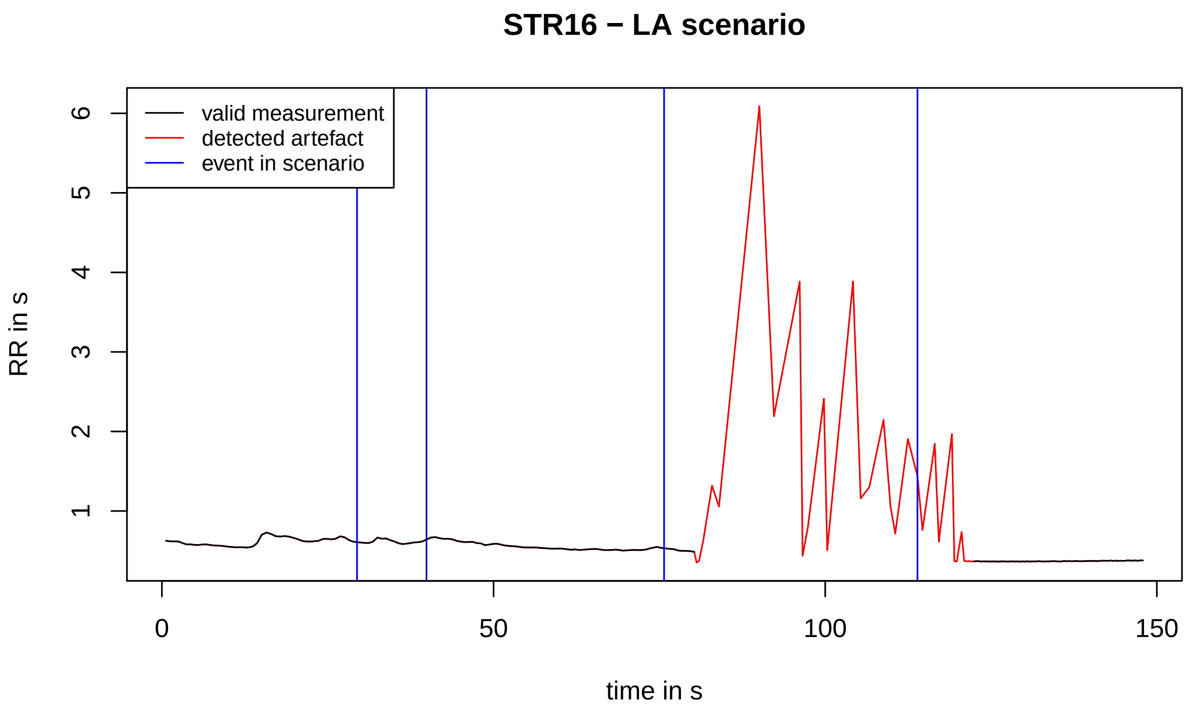
STR16 − LA scenario
valid measurement
detected artefact
event in scenario
RR in s
1
2
3
4
5
6
0
50
100
150
time in s

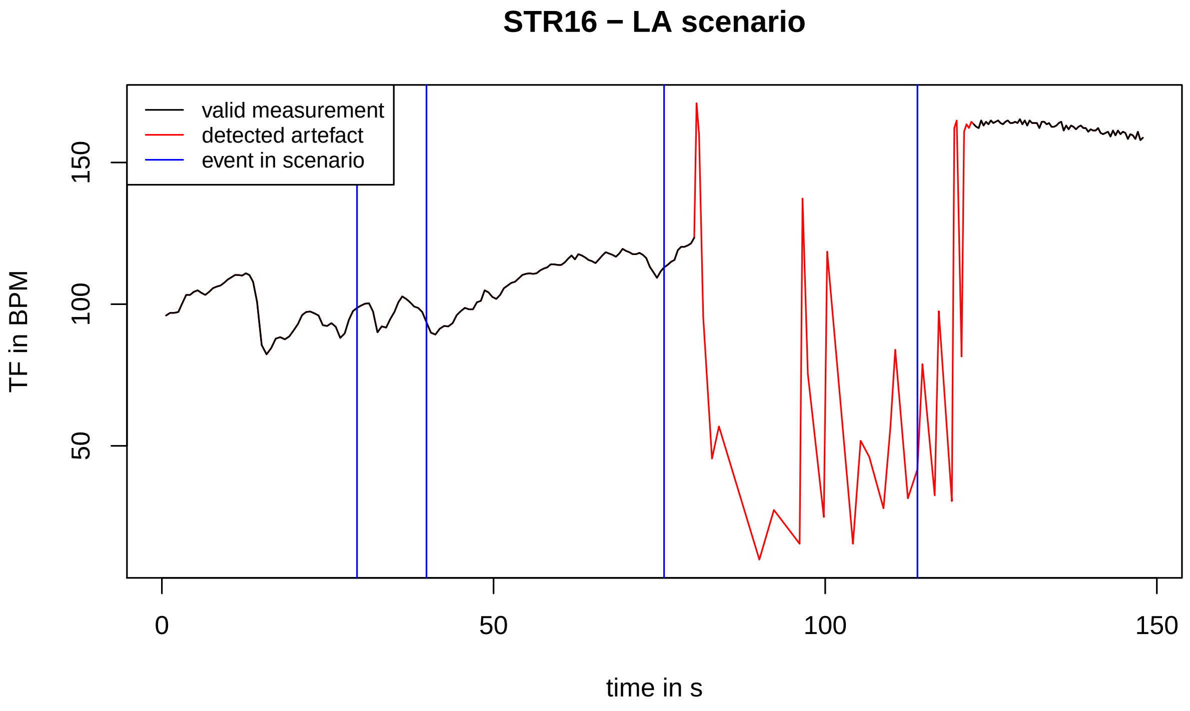
STR16 − LA scenario
valid measurement
detected artefact
event in scenario
TF in BPM
50
100
150
0
50
100
150
time in s

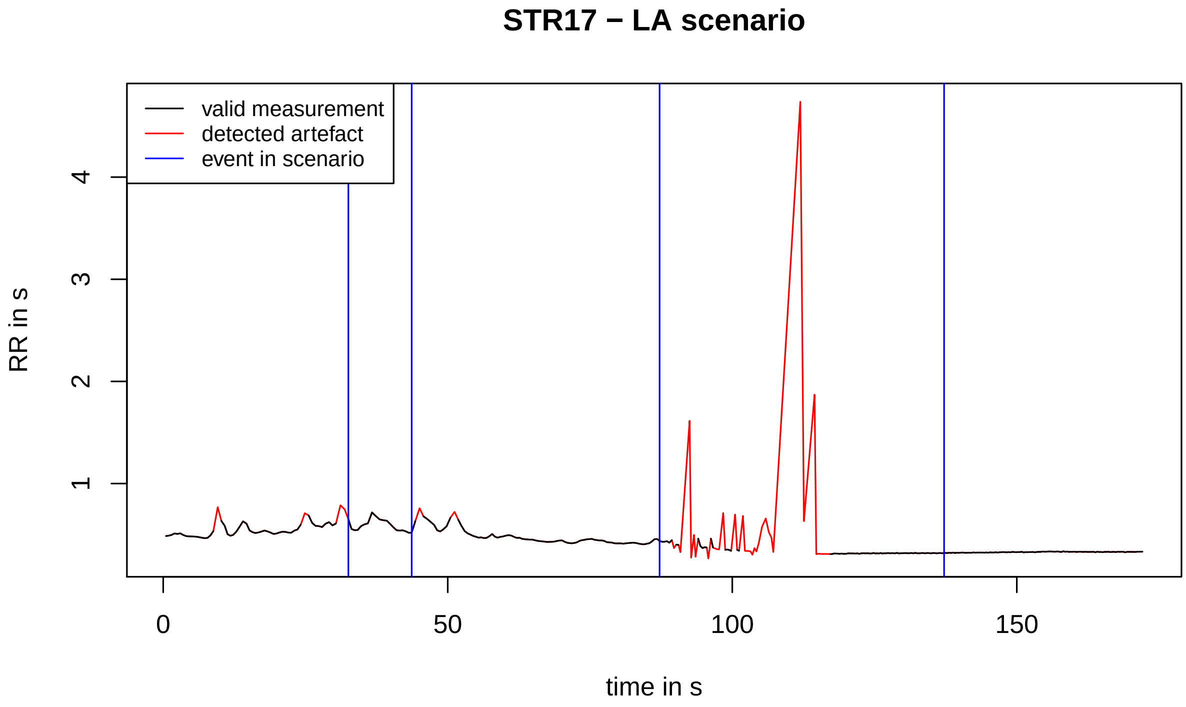
STR17 − LA scenario
valid measurement
detected artefact
event in scenario
RR in s
1
2
3
4
0
50
100
150
time in s

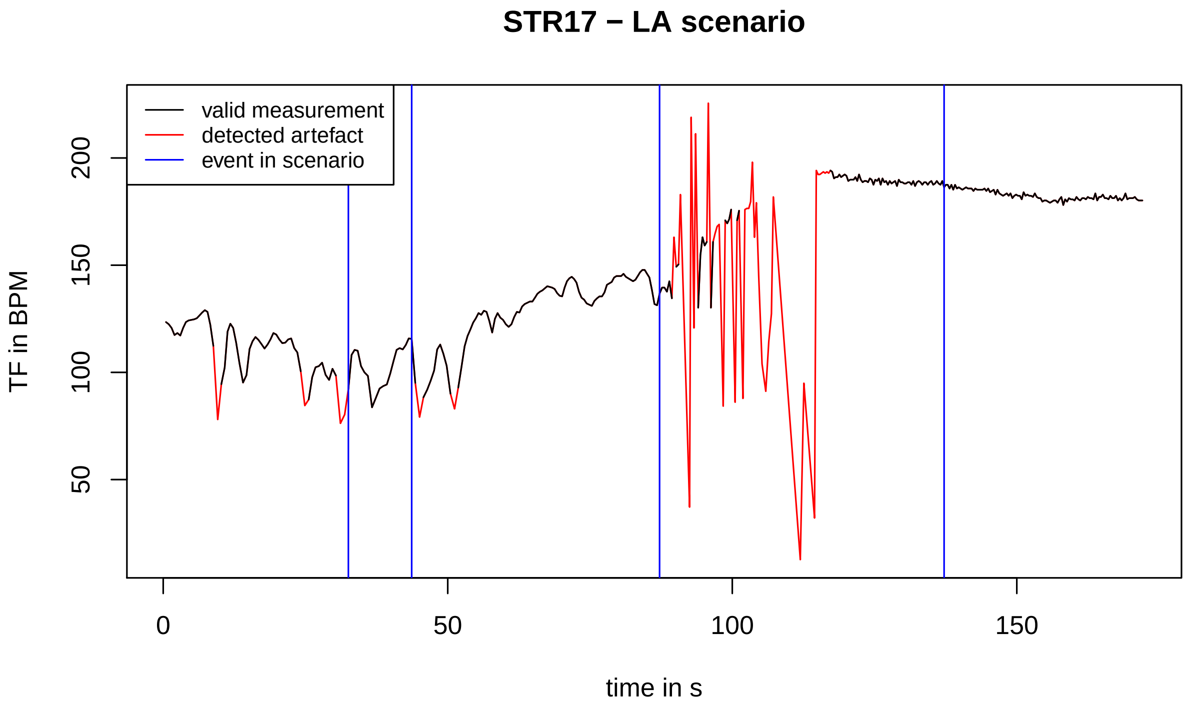
STR17 − LA scenario
valid measurement
detected artefact
event in scenario
TF in BPM
50
100
150
200
0
50
100
150
time in s

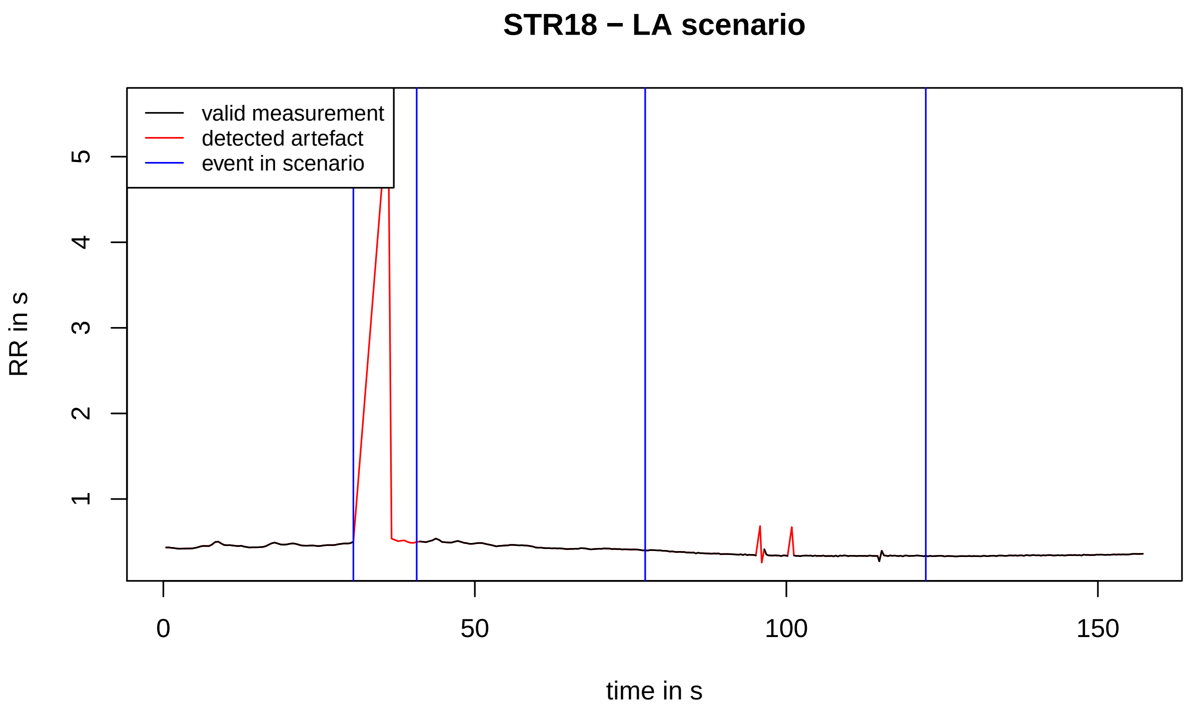
STR18 − LA scenario
valid measurement
detected artefact
event in scenario
RR in s
1
2
3
4
5
0
50
100
150
time in s

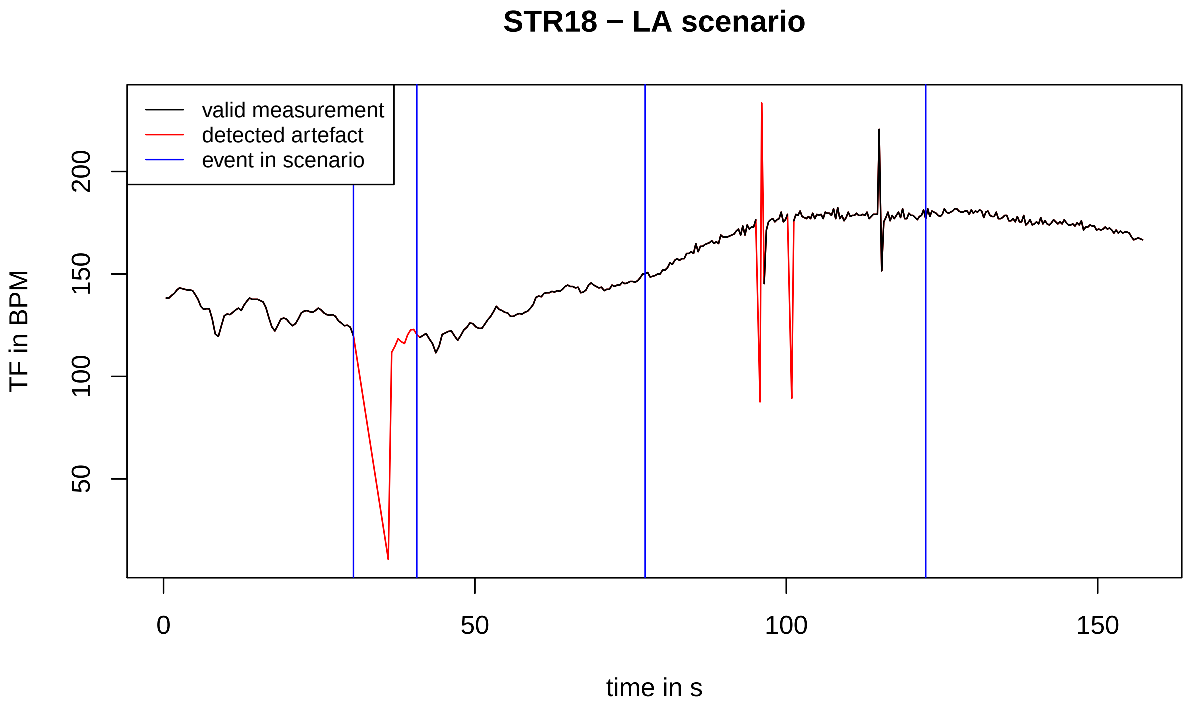
STR18 − LA scenario
valid measurement
detected artefact
event in scenario
TF in BPM
50
100
150
200
0
50
100
150
time in s

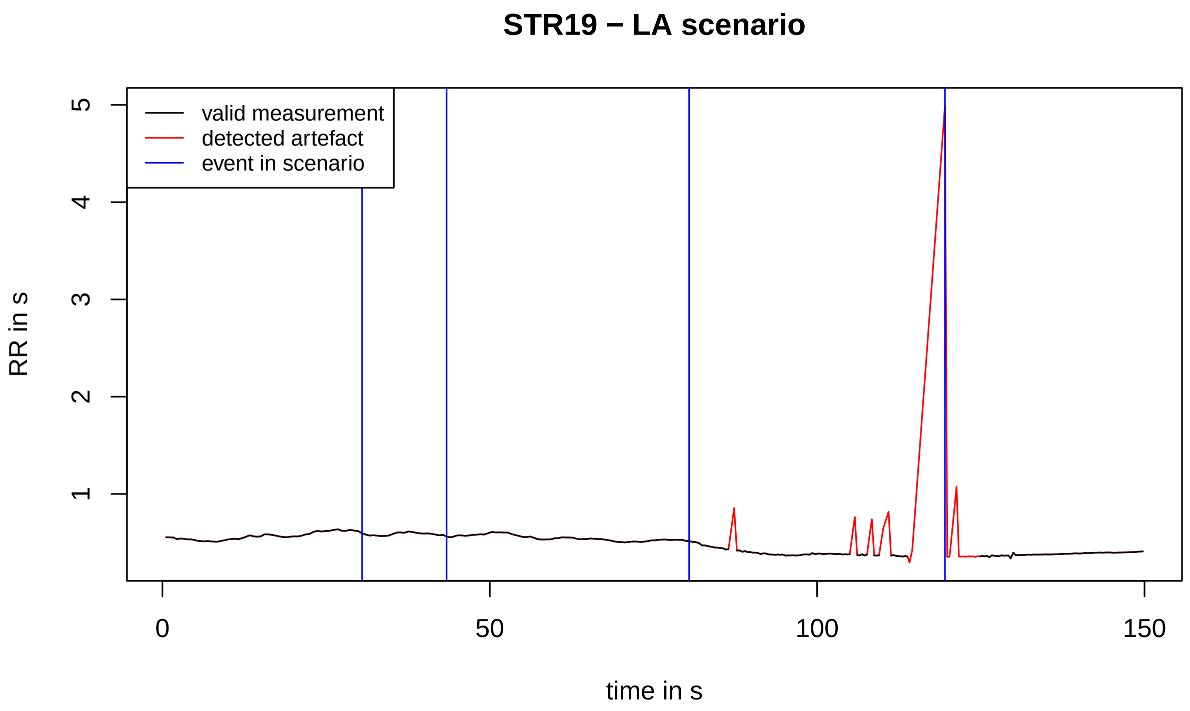
STR19 − LA scenario
valid measurement
detected artefact
event in scenario
RR in s
1
2
3
4
5
0
50
100
150
time in s

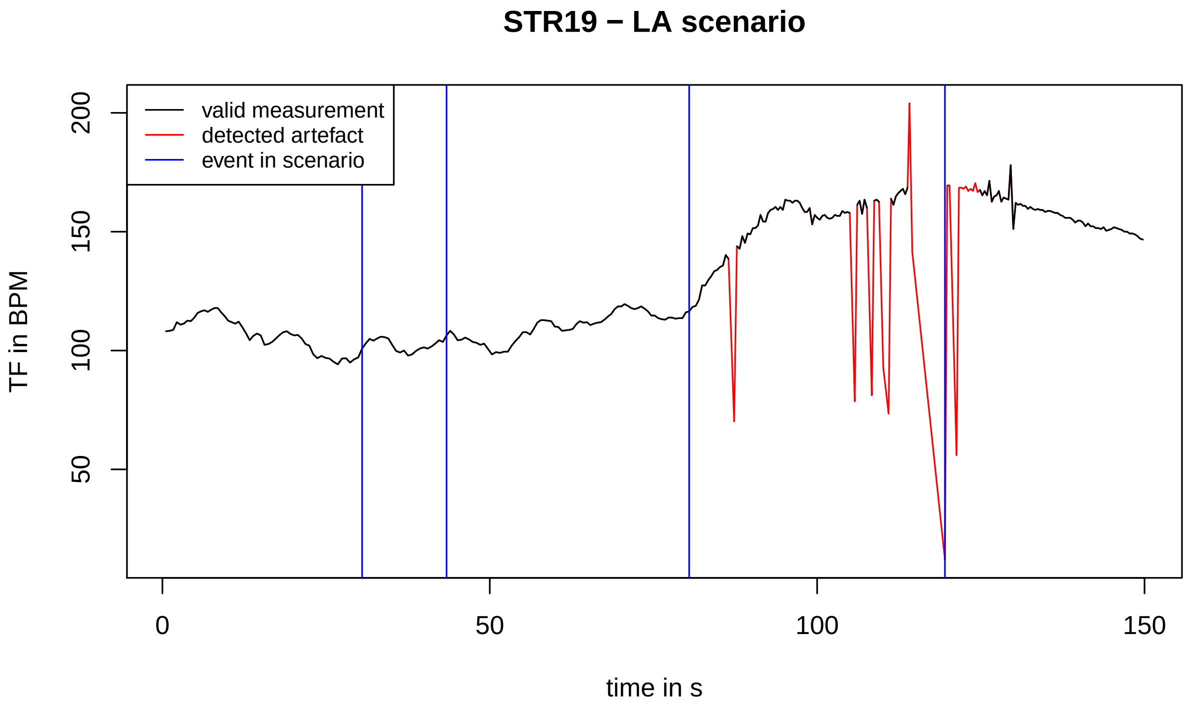
STR19 − LA scenario
valid measurement
detected artefact
event in scenario
TF in BPM
50
100
150
200
0
50
100
150
time in s

#### STR21 − LA scenario

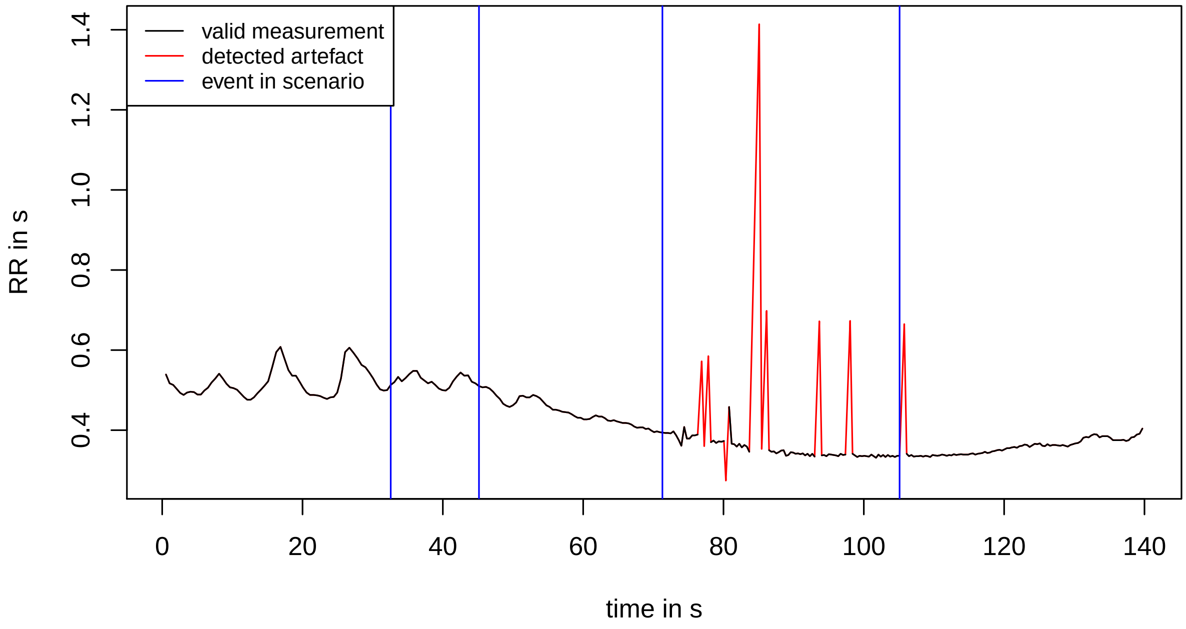


#### STR21 − LA scenario

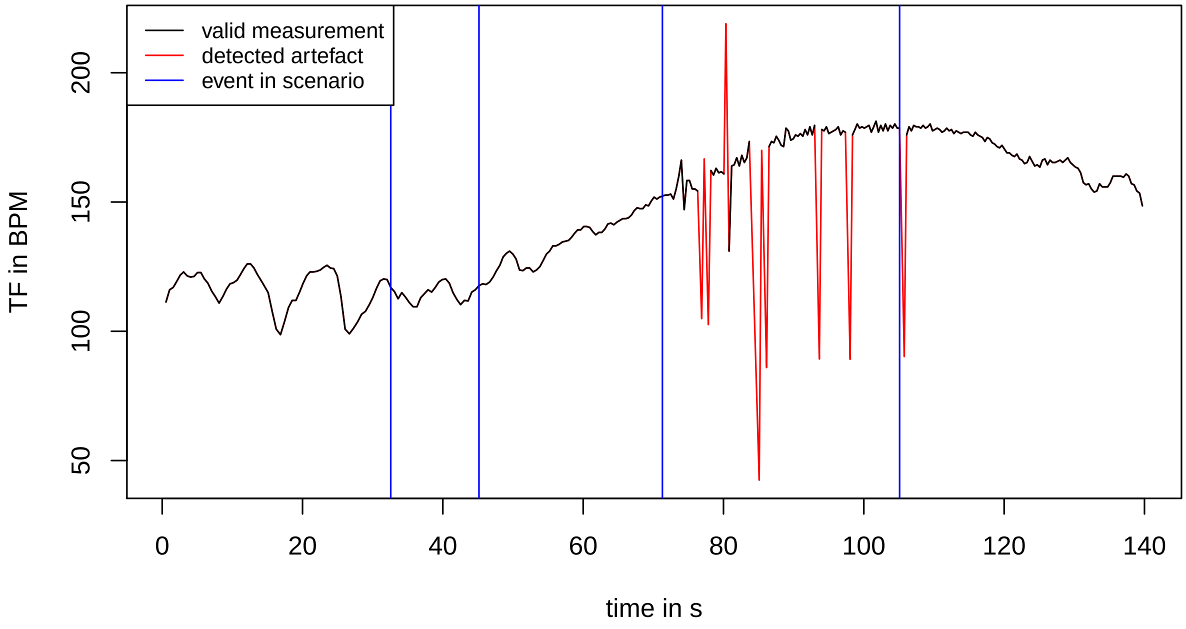

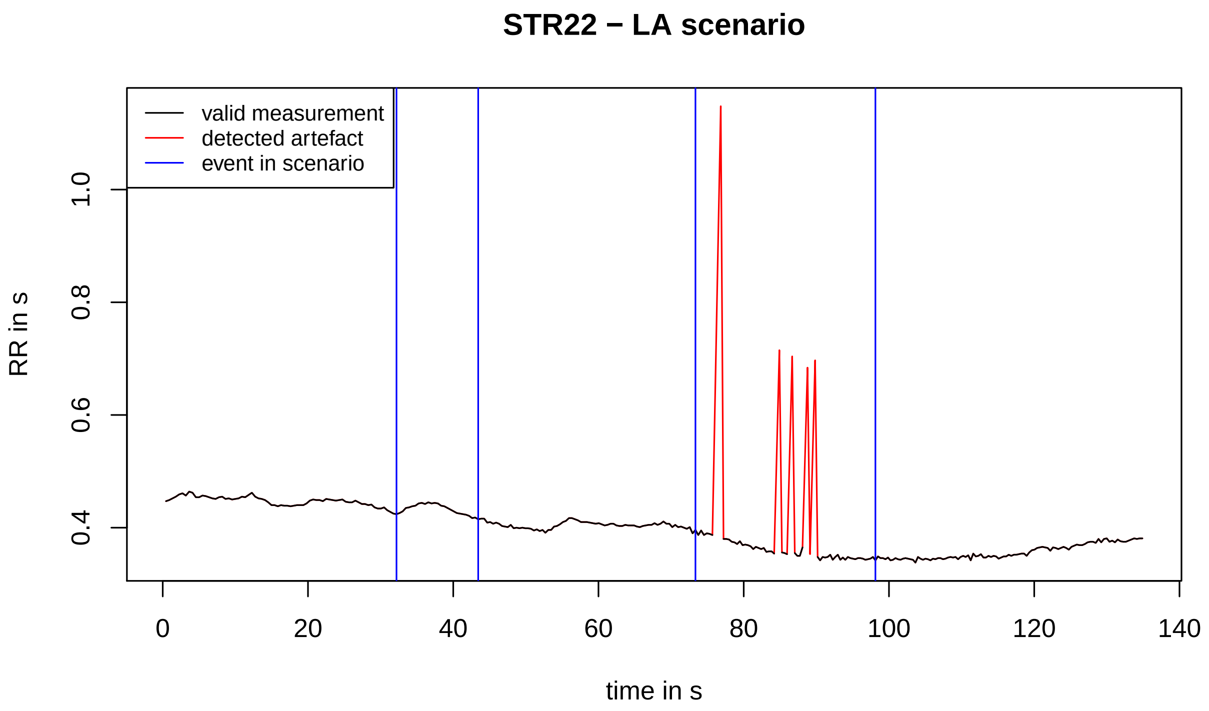
STR22 − LA scenario
valid measurement
detected artefact
event in scenario
RR in s
0.4
0.6
0.8
1.0
0
20
40
60
80
100
120
140
time in s

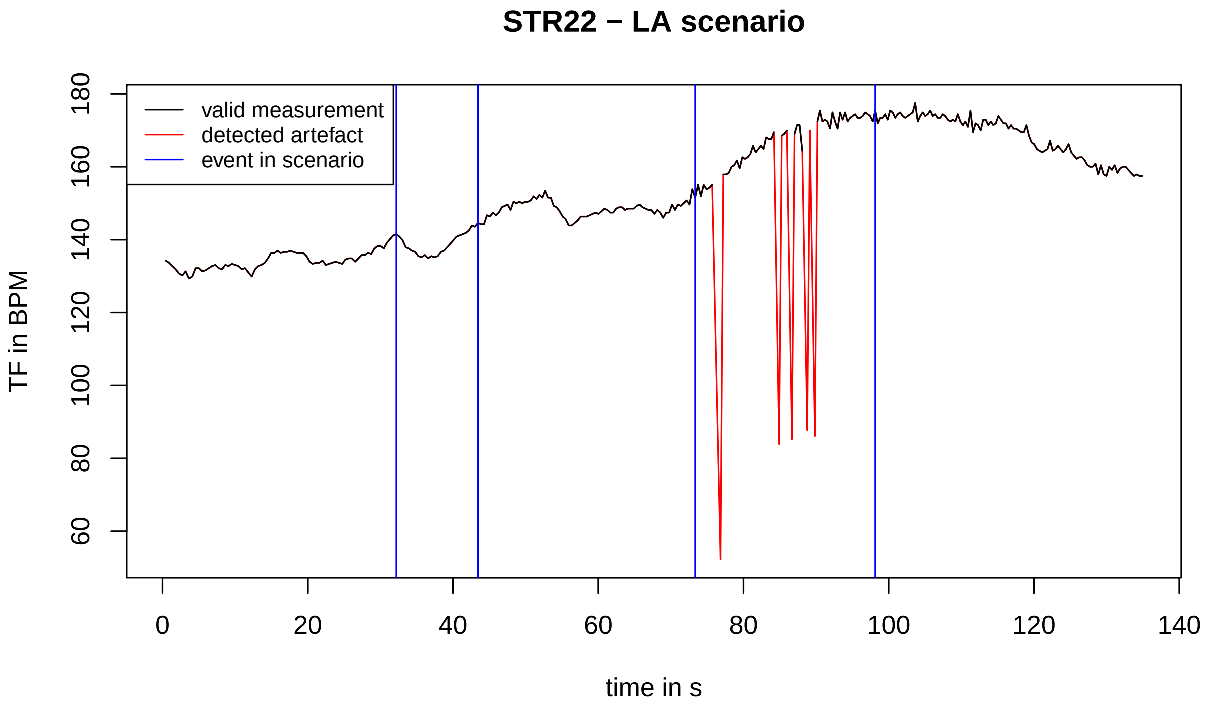
STR22 − LA scenario
valid measurement
detected artefact
event in scenario
TF in BPM
60
80
100
120
140
160
180
0
20
40
60
80
100
120
140
time in s

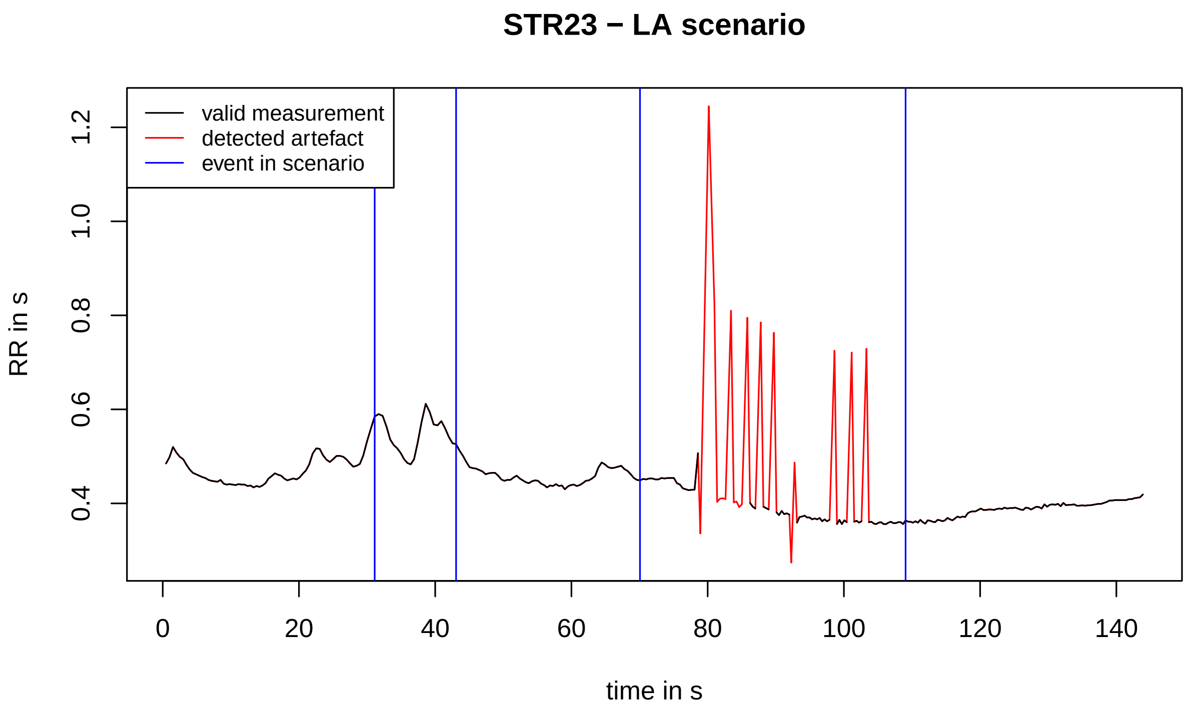
STR23 − LA scenario
valid measurement
detected artefact
event in scenario
RR in s
1.2
1.0
0.8
0.6
0.4
0
20
40
60
80
100
120
140
time in s

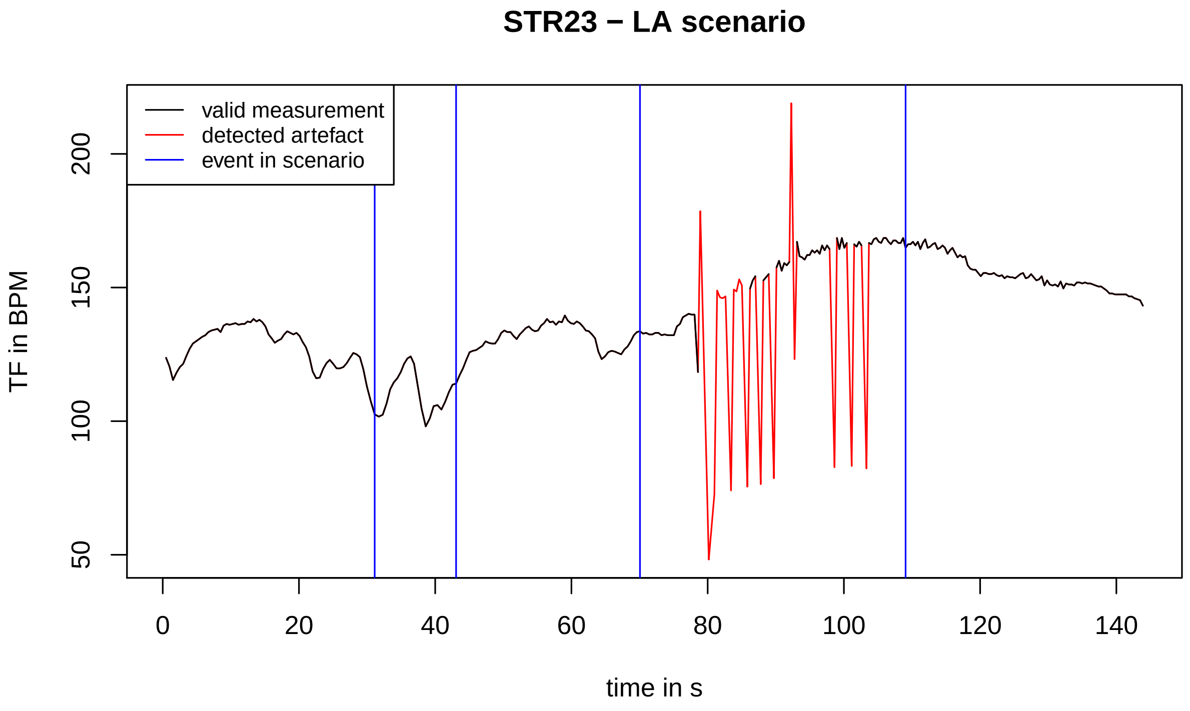
STR23 − LA scenario
valid measurement
detected artefact
event in scenario
TF in BPM
200
150
100
50
0
20
40
60
80
100
120
140
time in s

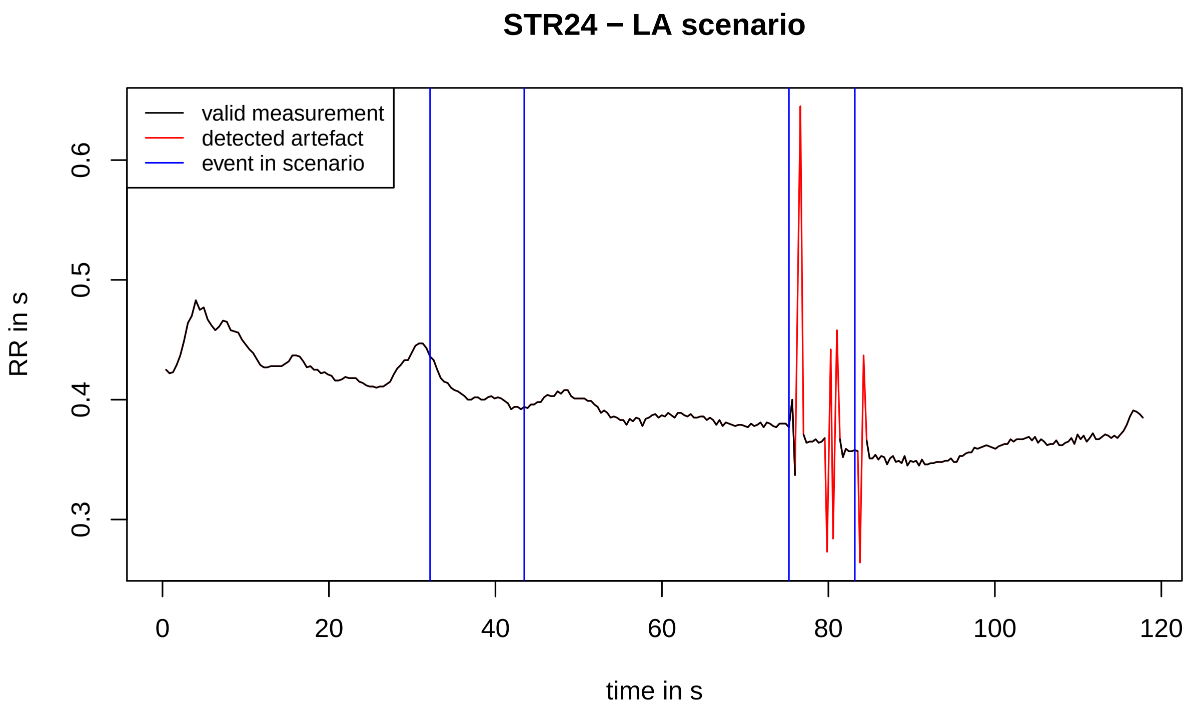
STR24 − LA scenario
valid measurement
detected artefact
event in scenario
RR in s
0.3
0.4
0.5
0.6
0
20
40
60
80
100
120
time in s

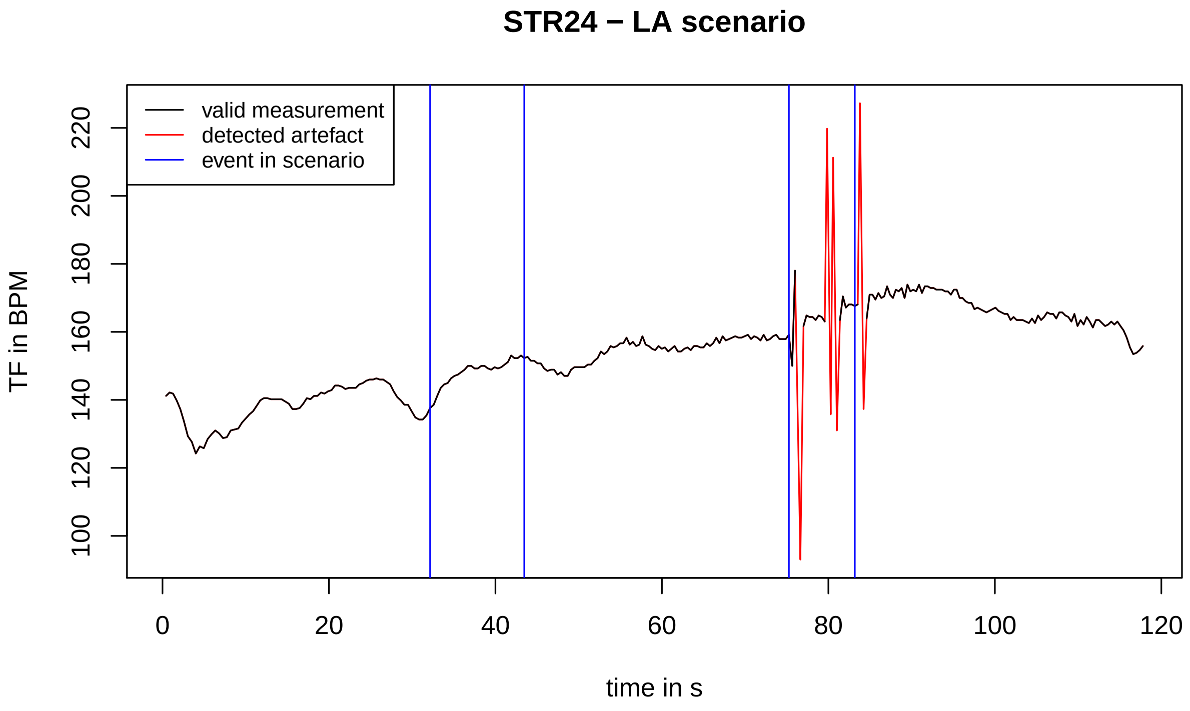
STR24 − LA scenario
valid measurement
detected artefact
event in scenario
TF in BPM
100
120
140
160
180
200
220
0
20
40
60
80
100
120
time in s

#### STR25 − LA scenario

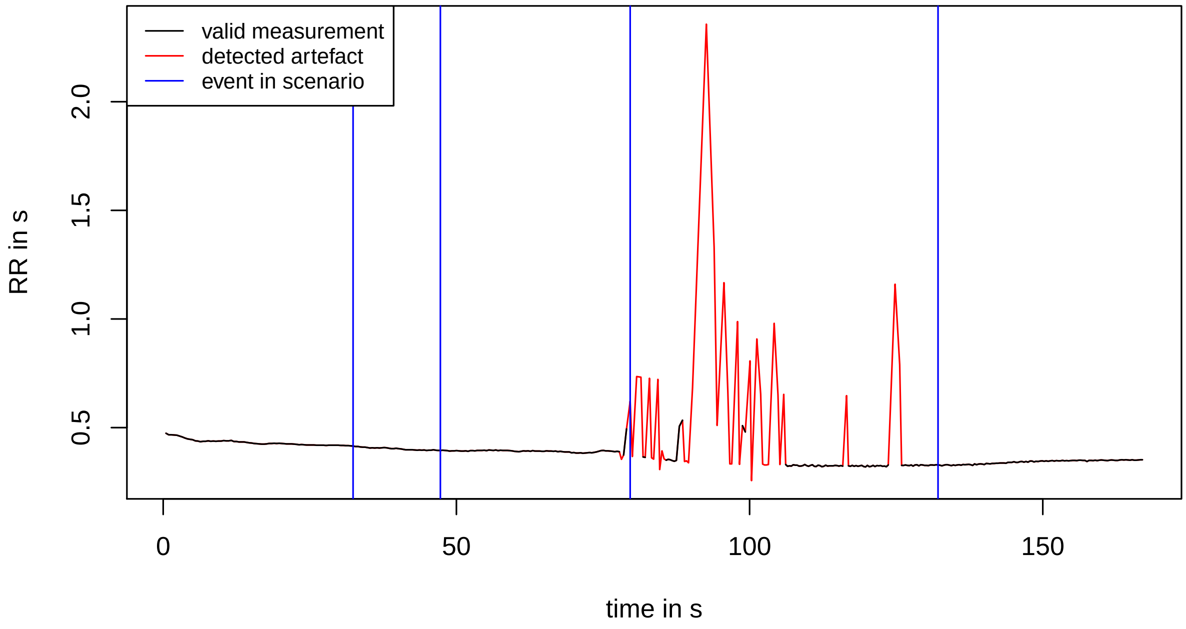


#### STR25 − LA scenario

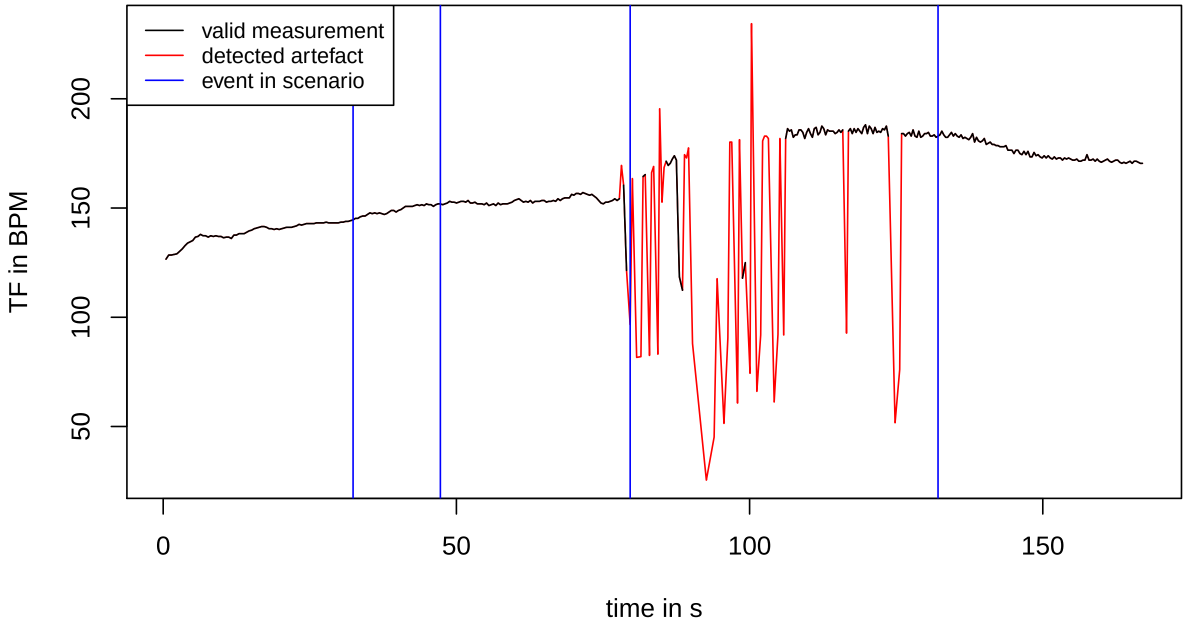

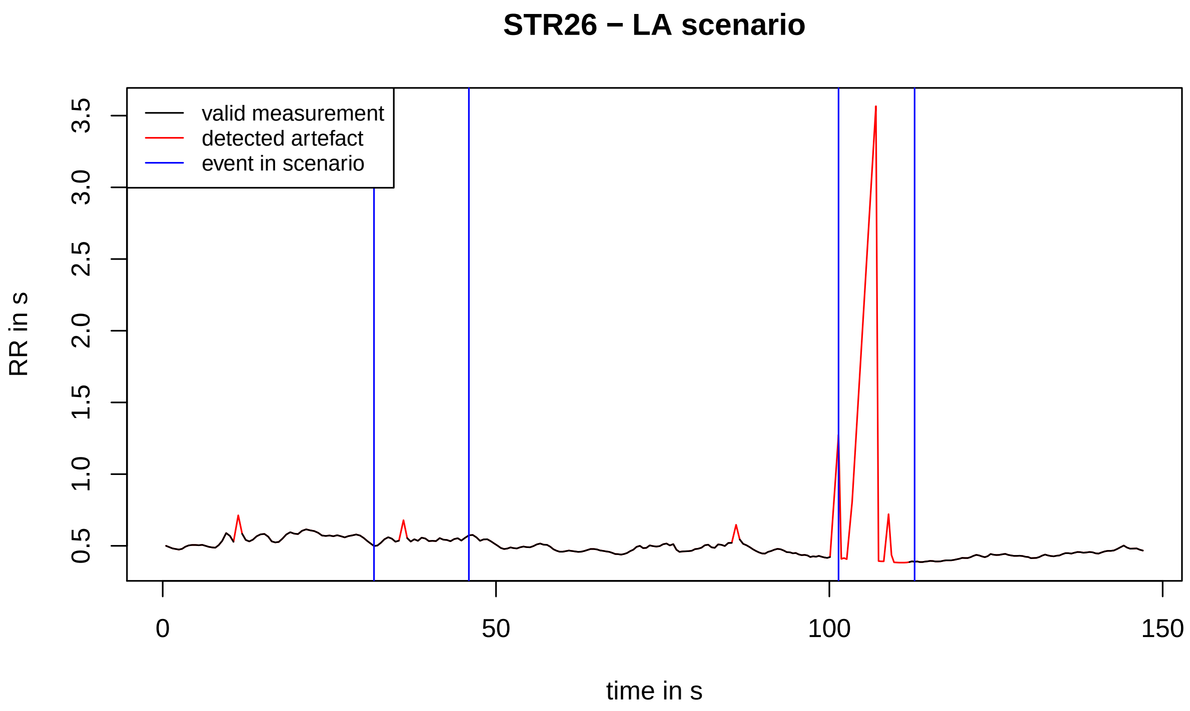
STR26 − LA scenario
valid measurement
detected artefact
event in scenario
RR in s
0.5
1.0
1.5
2.0
2.5
3.0
3.5
0
50
100
150
time in s

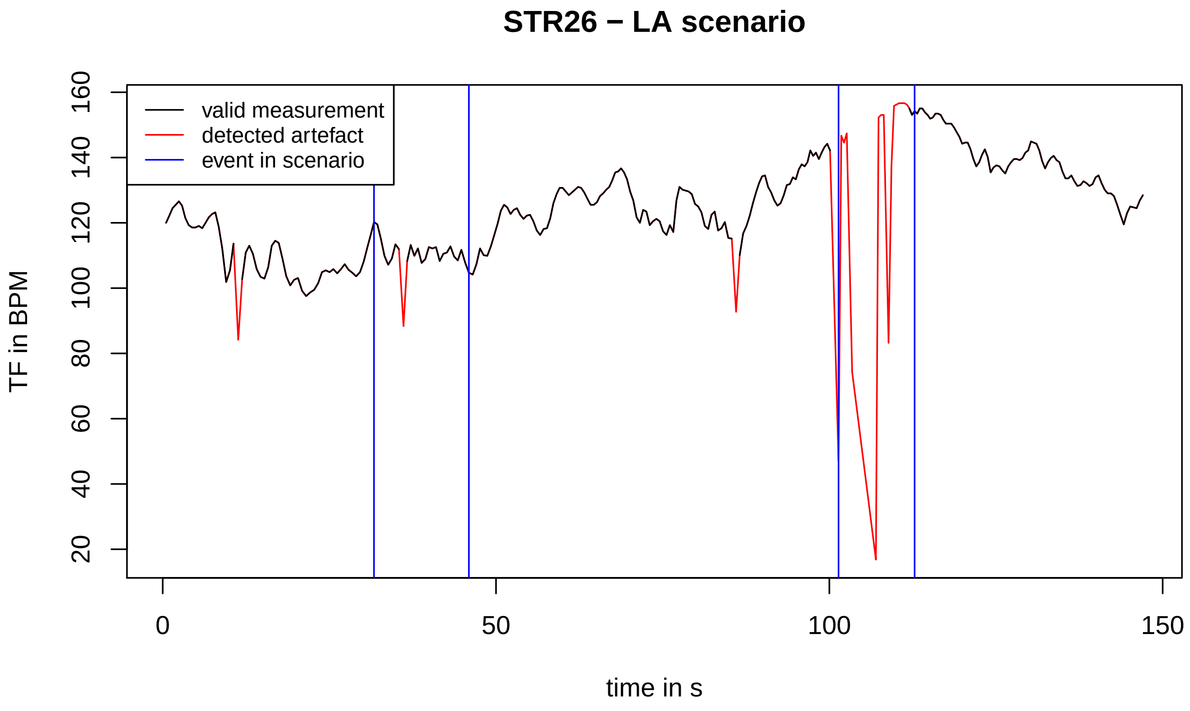
STR26 − LA scenario
valid measurement
detected artefact
event in scenario
TF in BPM
20
40
60
80
100
120
140
160
0
50
100
150
time in s

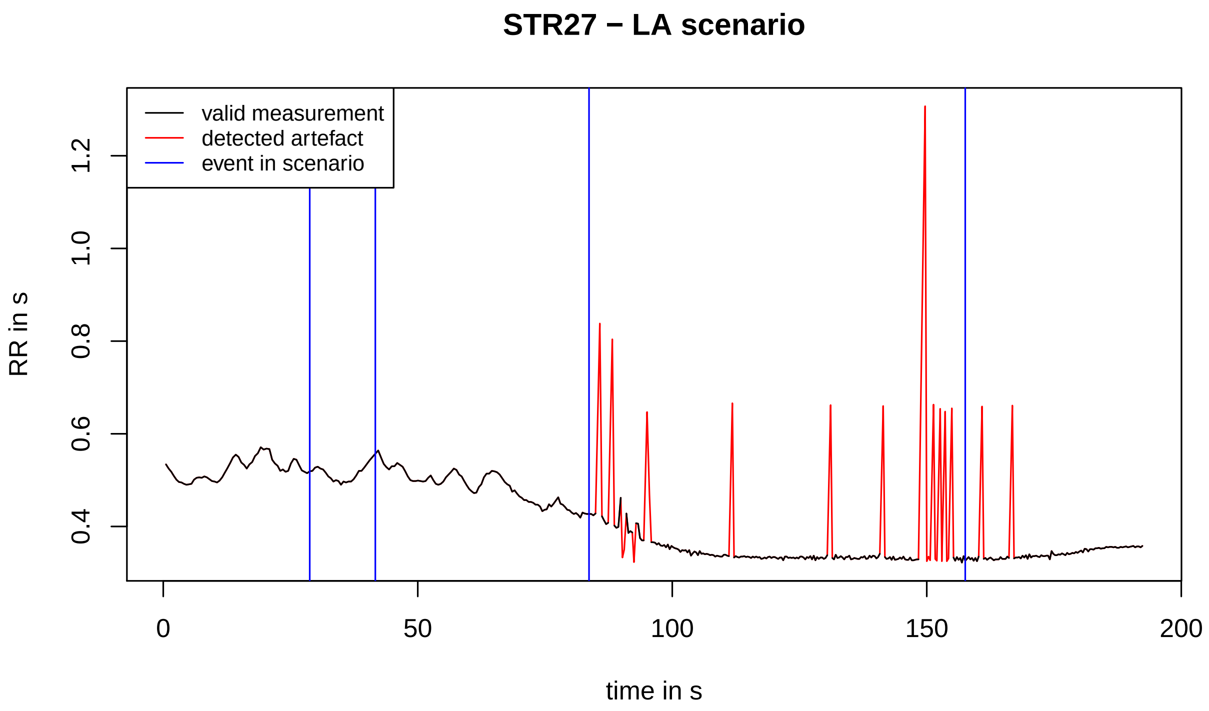
STR27 − LA scenario
valid measurement
detected artefact
event in scenario
RR in s
0.4
0.6
0.8
1.0
1.2
0
50
100
150
200
time in s

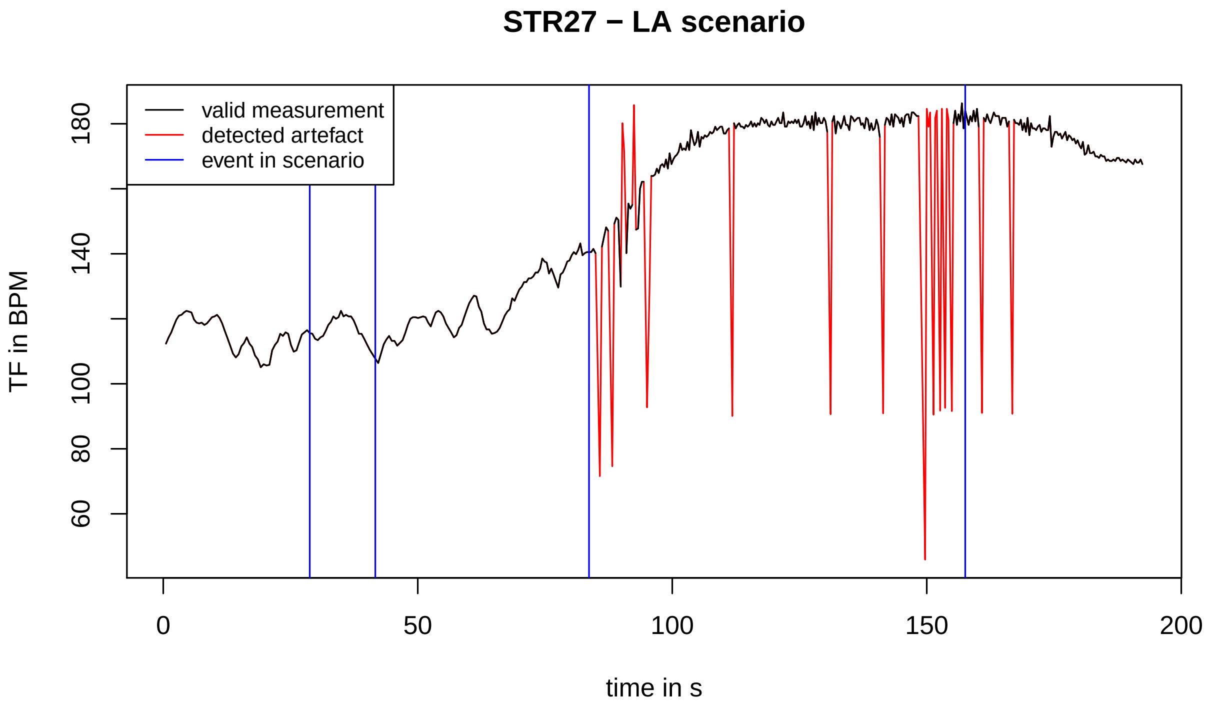
STR27 − LA scenario
valid measurement
detected artefact
event in scenario
TF in BPM
60
80
100
140
180
0
50
100
150
200
time in s

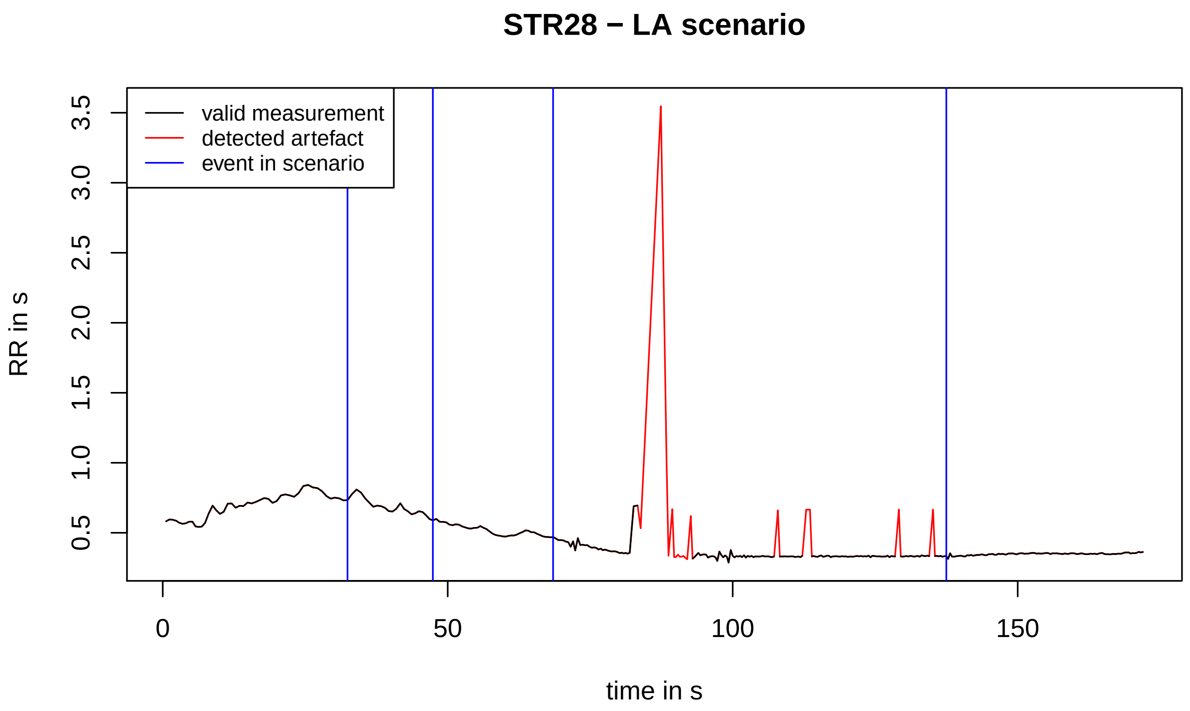
STR28 − LA scenario
valid measurement
detected artefact
event in scenario
RR in s
0.5
1.0
1.5
2.0
2.5
3.0
3.5
0
50
100
150
time in s

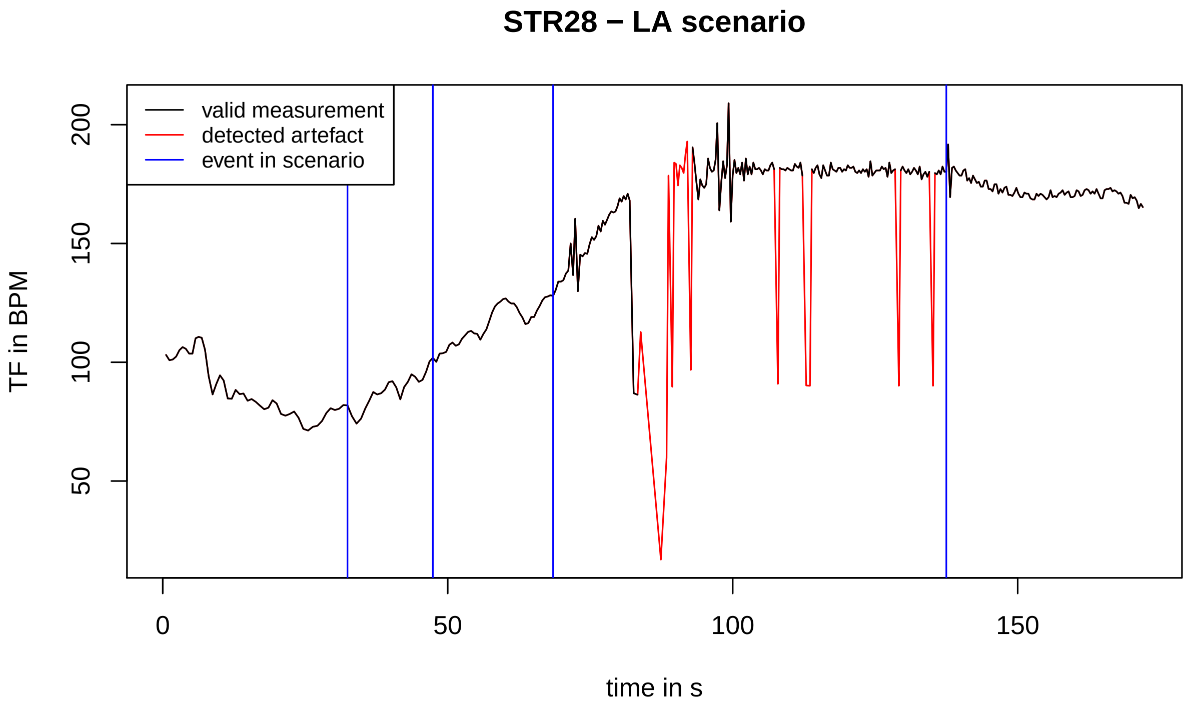
STR28 − LA scenario
valid measurement
detected artefact
event in scenario
TF in BPM
50
100
150
200
0
50
100
150
time in s

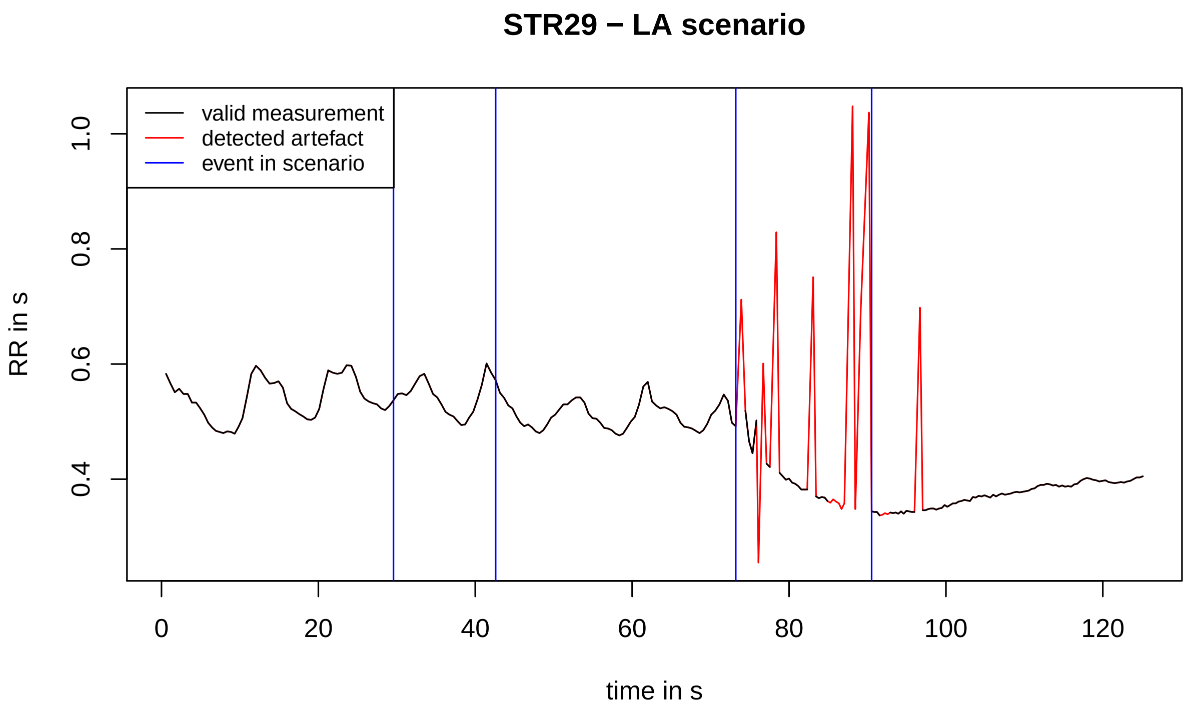
STR29 − LA scenario
valid measurement
detected artefact
event in scenario
RR in s
0.4
0.6
0.8
1.0
0
20
40
60
80
100
120
time in s

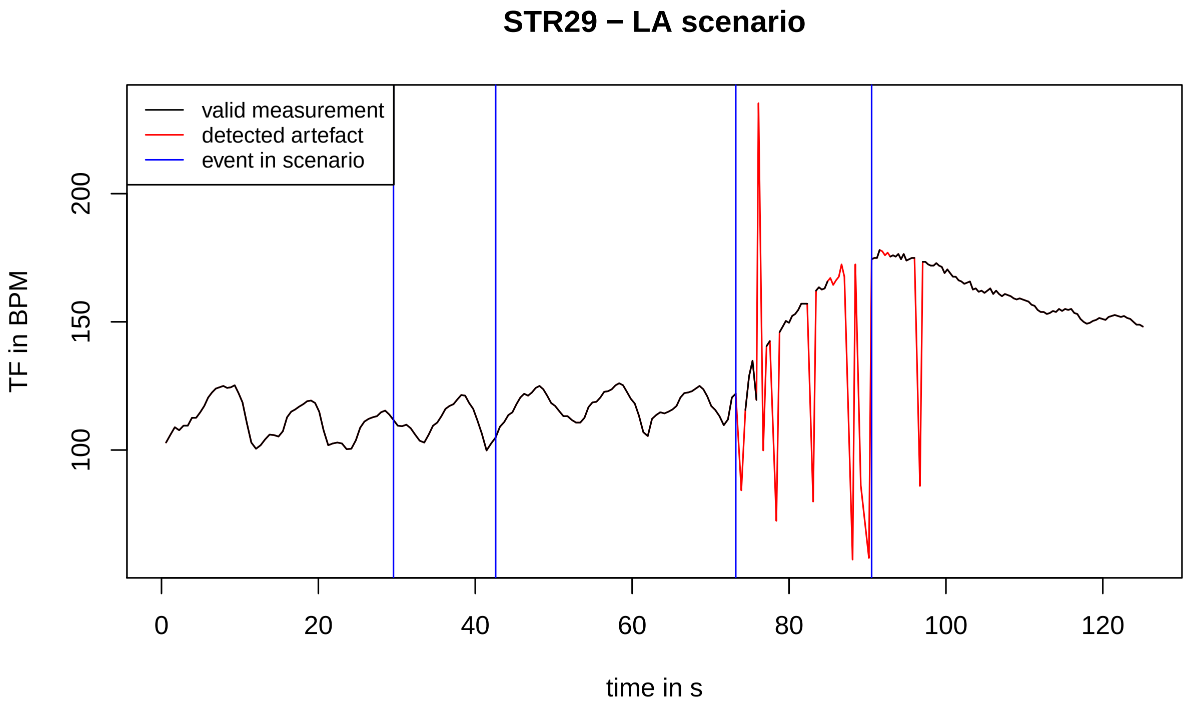
STR29 − LA scenario
valid measurement
detected artefact
event in scenario
TF in BPM
100
150
200
0
20
40
60
80
100
120
time in s

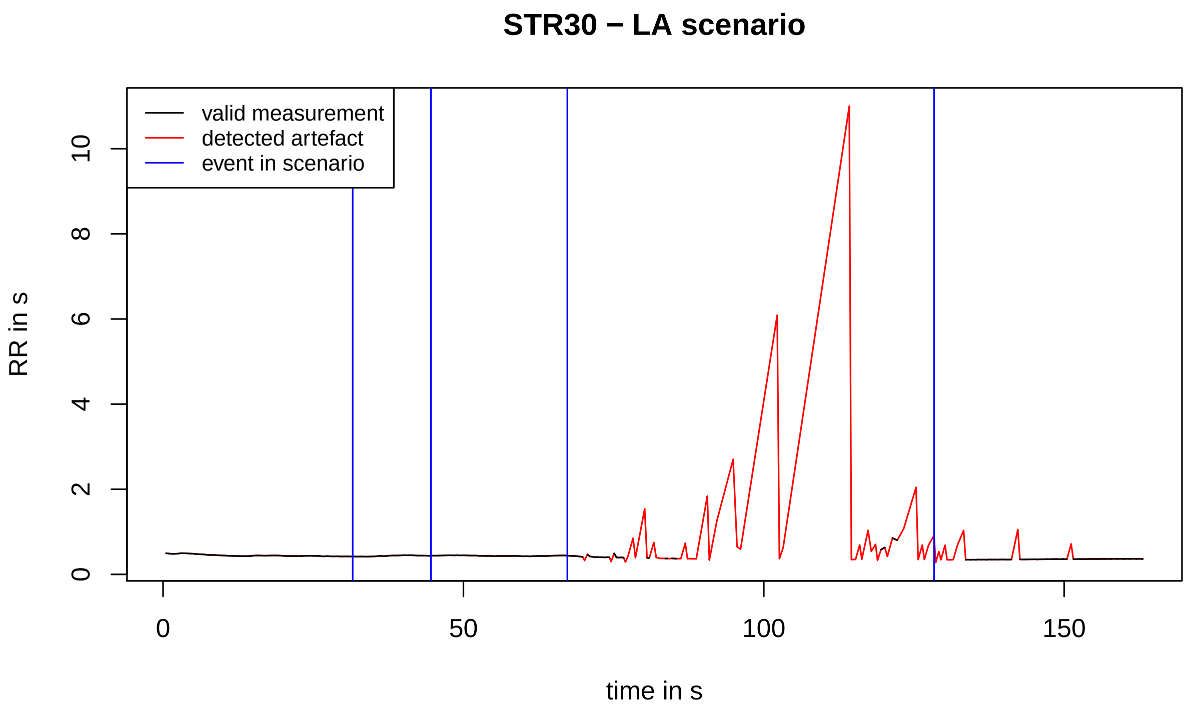
STR30 − LA scenario
valid measurement
detected artefact
event in scenario
RR in s
0
2
4
6
8
10
0
50
100
150
time in s

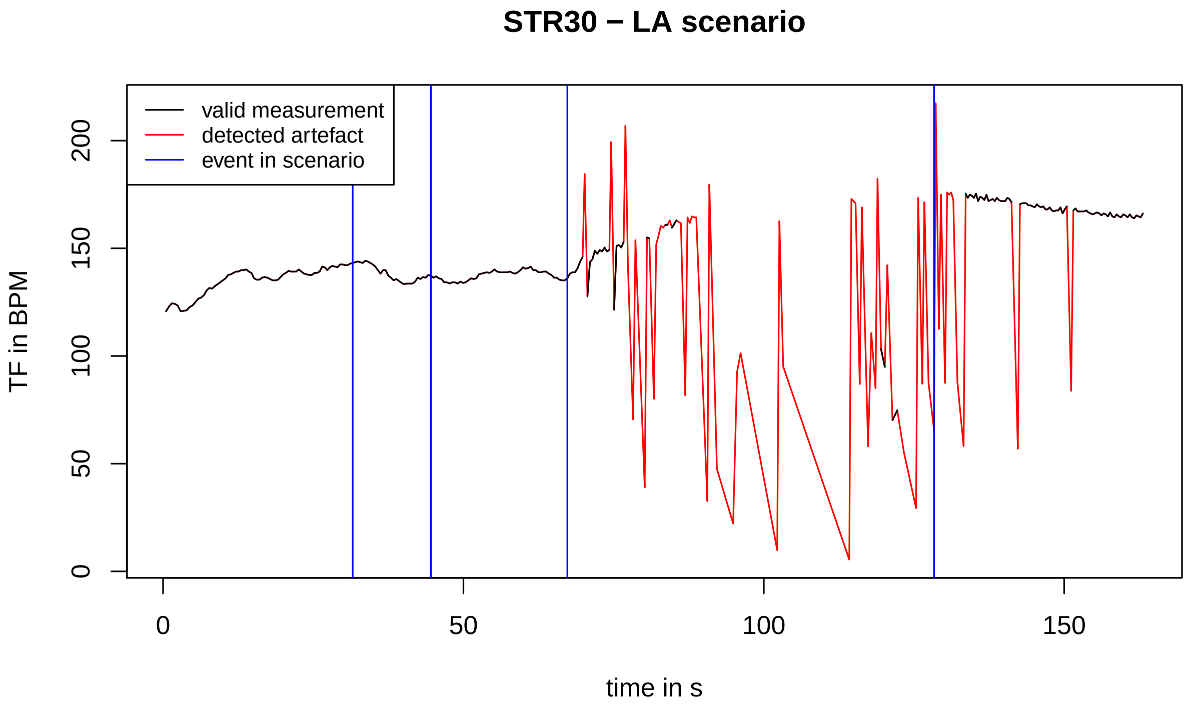
STR30 − LA scenario
valid measurement
detected artefact
event in scenario
TF in BPM
0
50
100
150
200
0
50
100
150
time in s

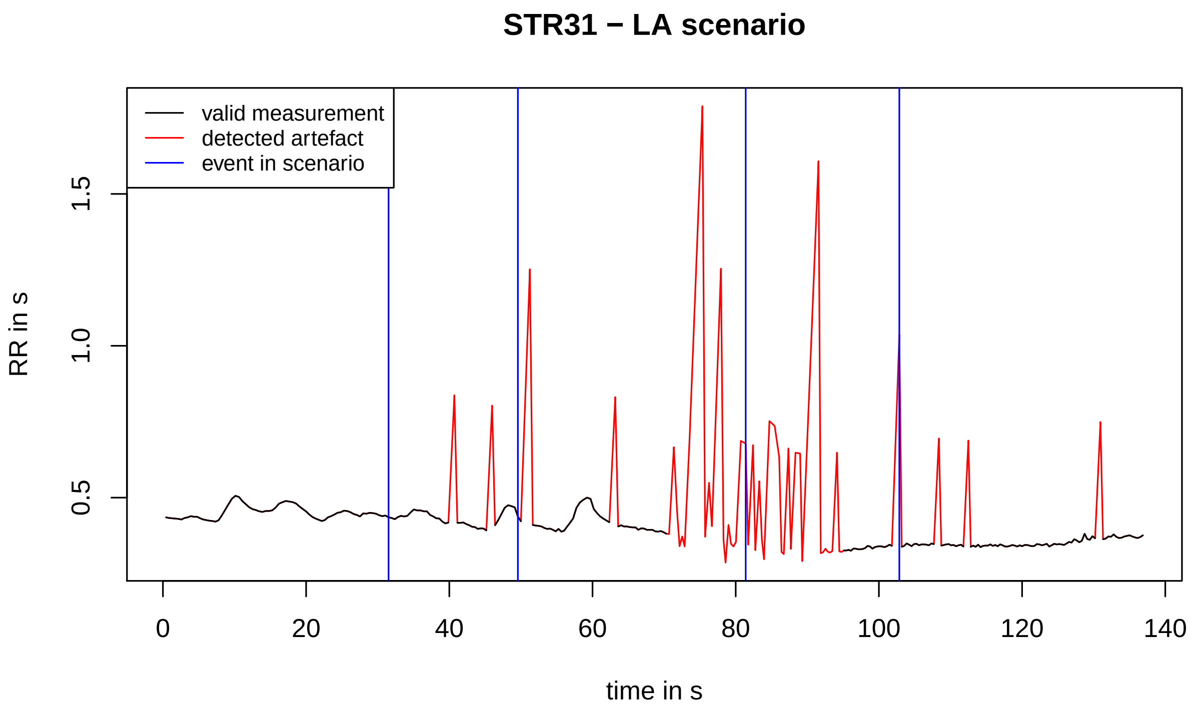
STR31 − LA scenario
valid measurement
detected artefact
event in scenario
RR in s
0.5
1.0
1.5
0
20
40
60
80
100
120
140
time in s

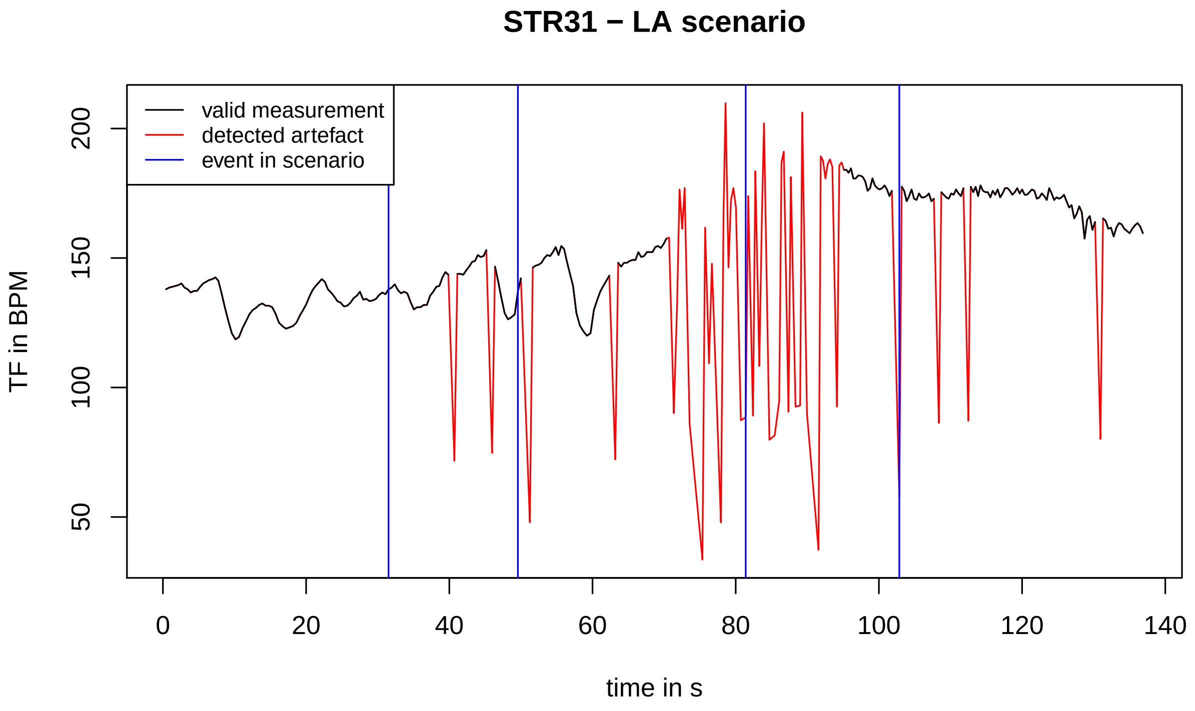
STR31 − LA scenario
valid measurement
detected artefact
event in scenario
TF in BPM
50
100
150
200
0
20
40
60
80
100
120
140
time in s

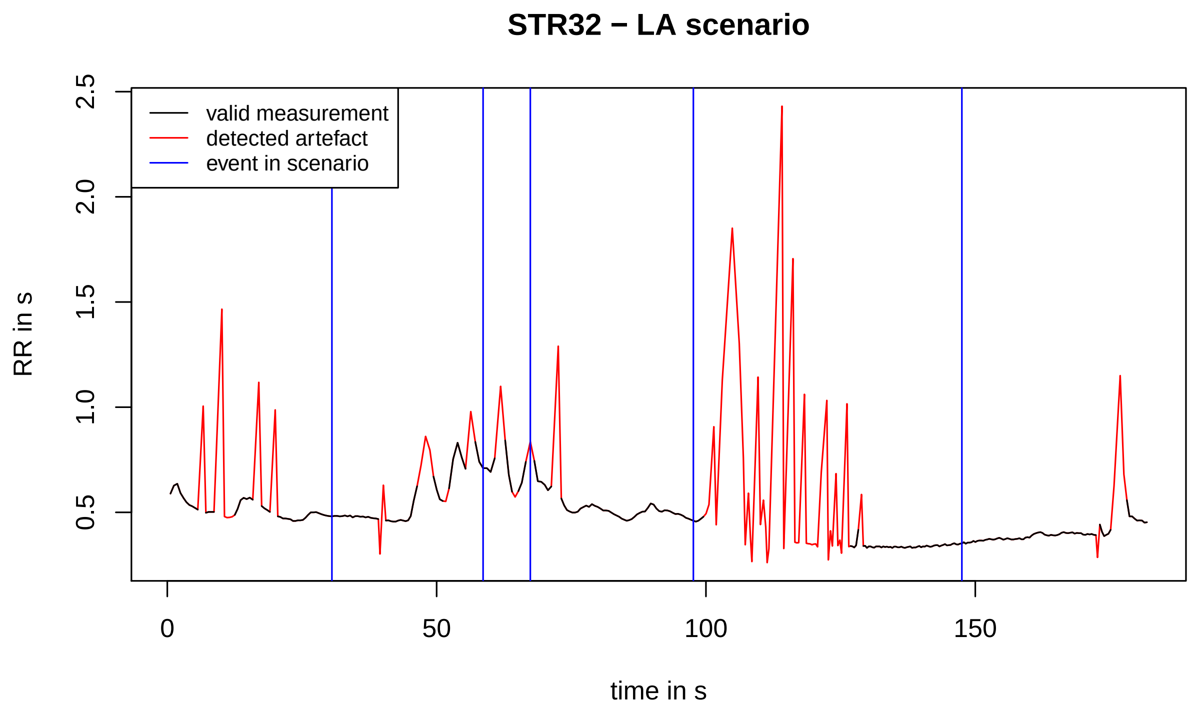
STR32 − LA scenario
valid measurement
detected artefact
event in scenario
RR in s
0.5
1.0
1.5
2.0
2.5
0
50
100
150
time in s

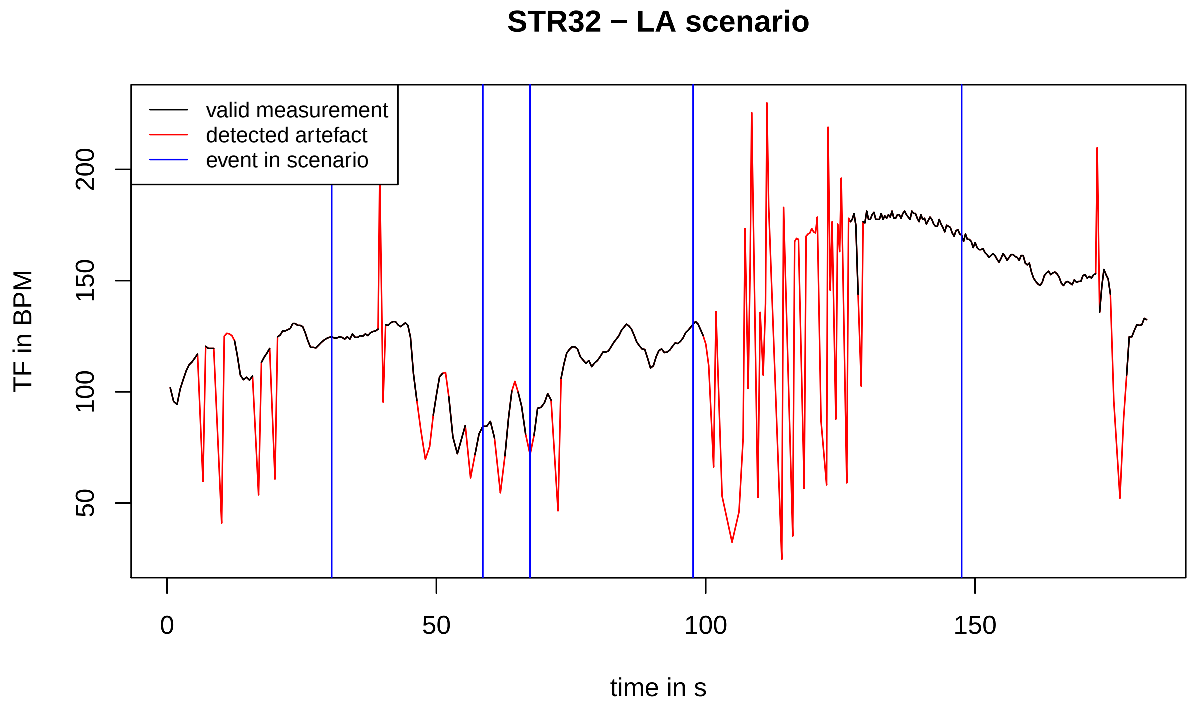
STR32 − LA scenario
valid measurement
detected artefact
event in scenario
TF in BPM
50
100
150
200
0
50
100
150
time in s

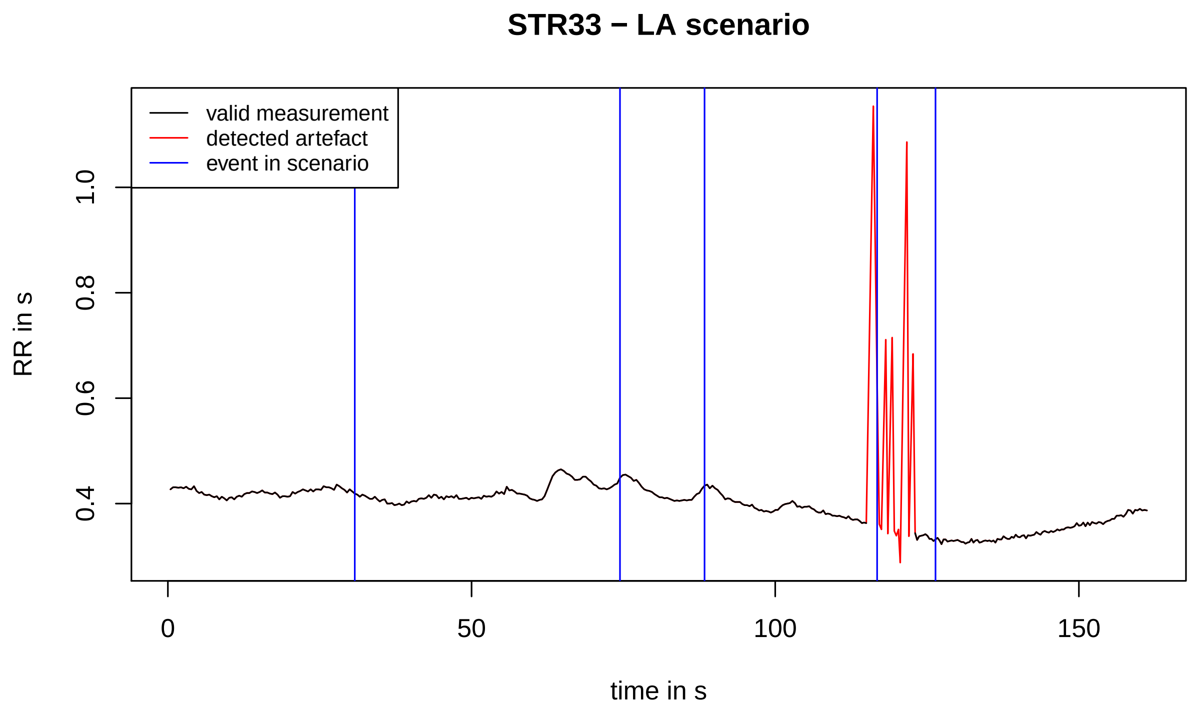
STR33 − LA scenario
valid measurement
detected artefact
event in scenario
RR in s
0.4
0.6
0.8
1.0
0
50
100
150
time in s

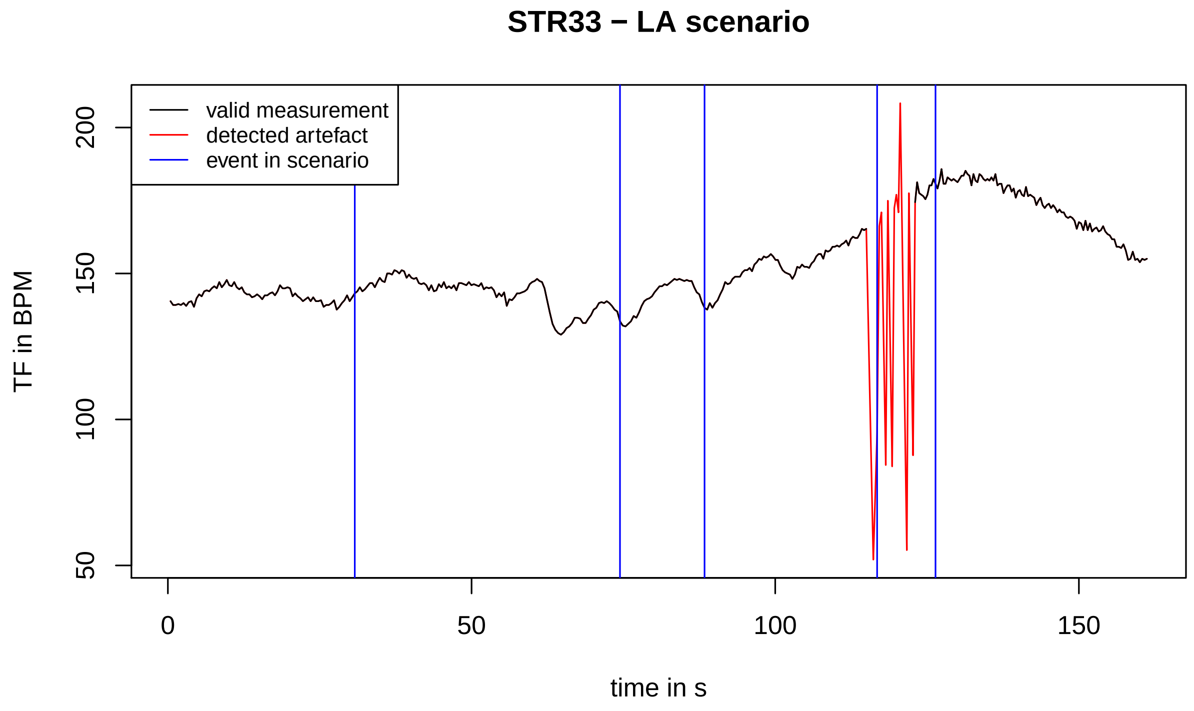
STR33 − LA scenario
valid measurement
detected artefact
event in scenario
TF in BPM
50
100
150
200
0
50
100
150
time in s

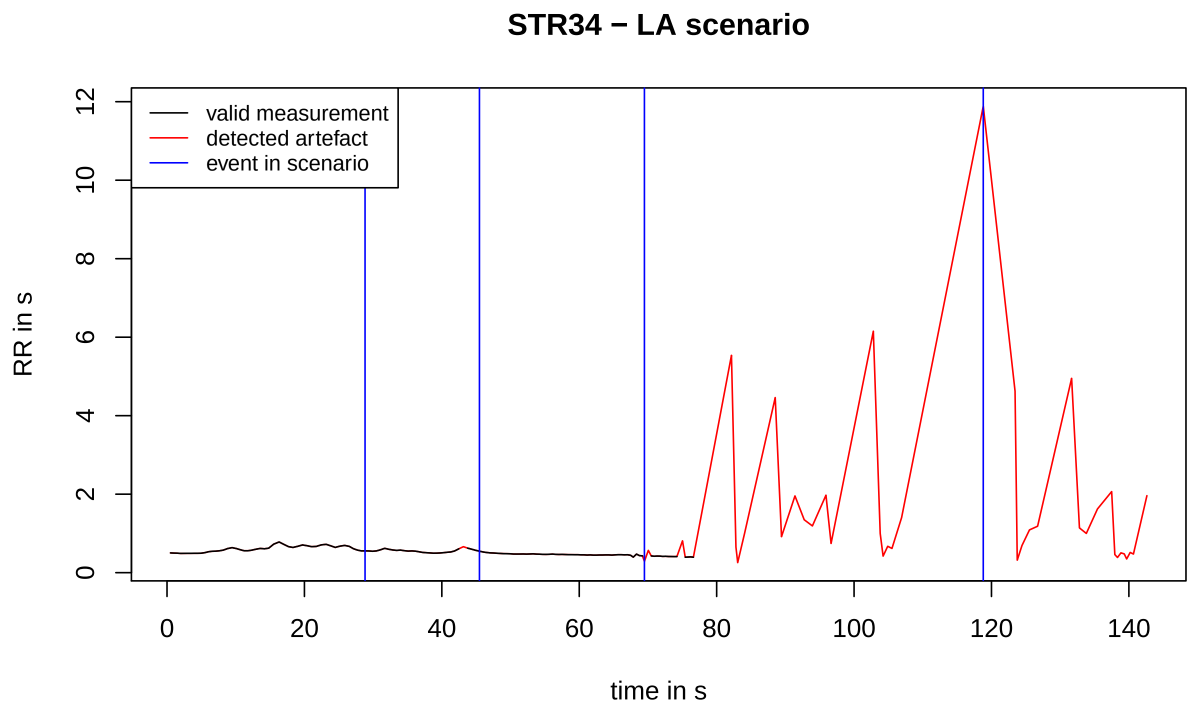
STR34 − LA scenario
valid measurement
detected artefact
event in scenario
RR in s
0
2
4
6
8
10
12
0
20
40
60
80
100
120
140
time in s

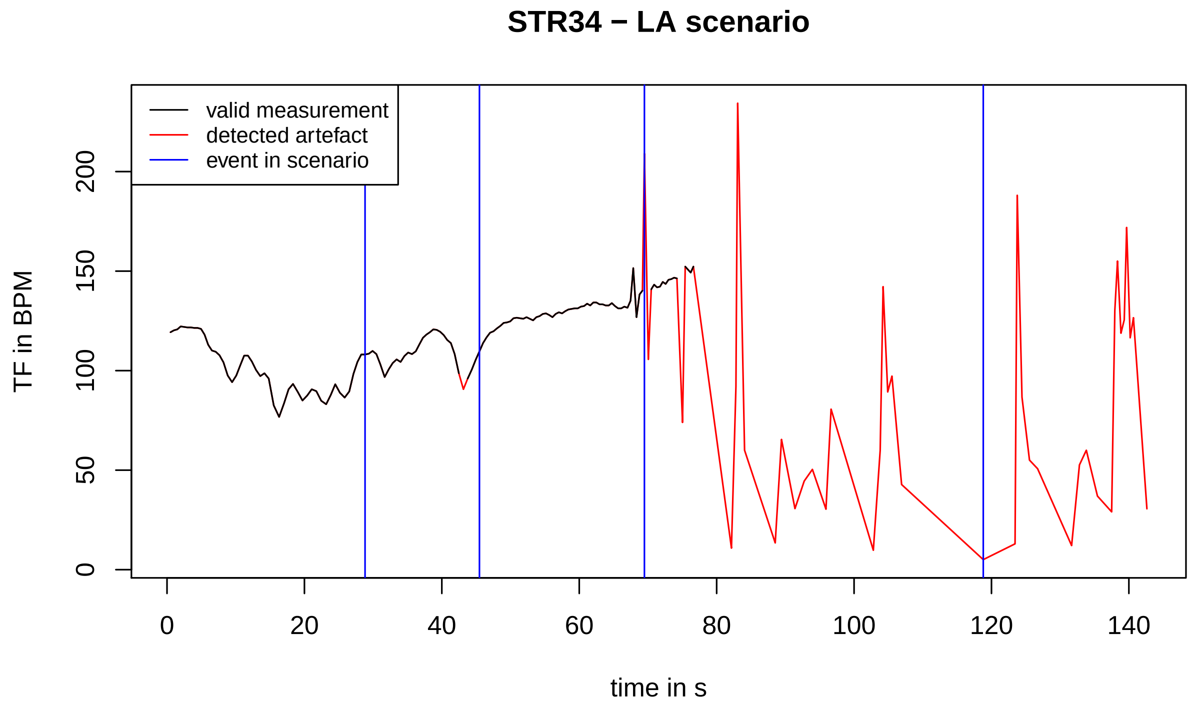
STR34 − LA scenario
valid measurement
detected artefact
event in scenario
TF in BPM
0
50
100
150
200
0
20
40
60
80
100
120
140
time in s

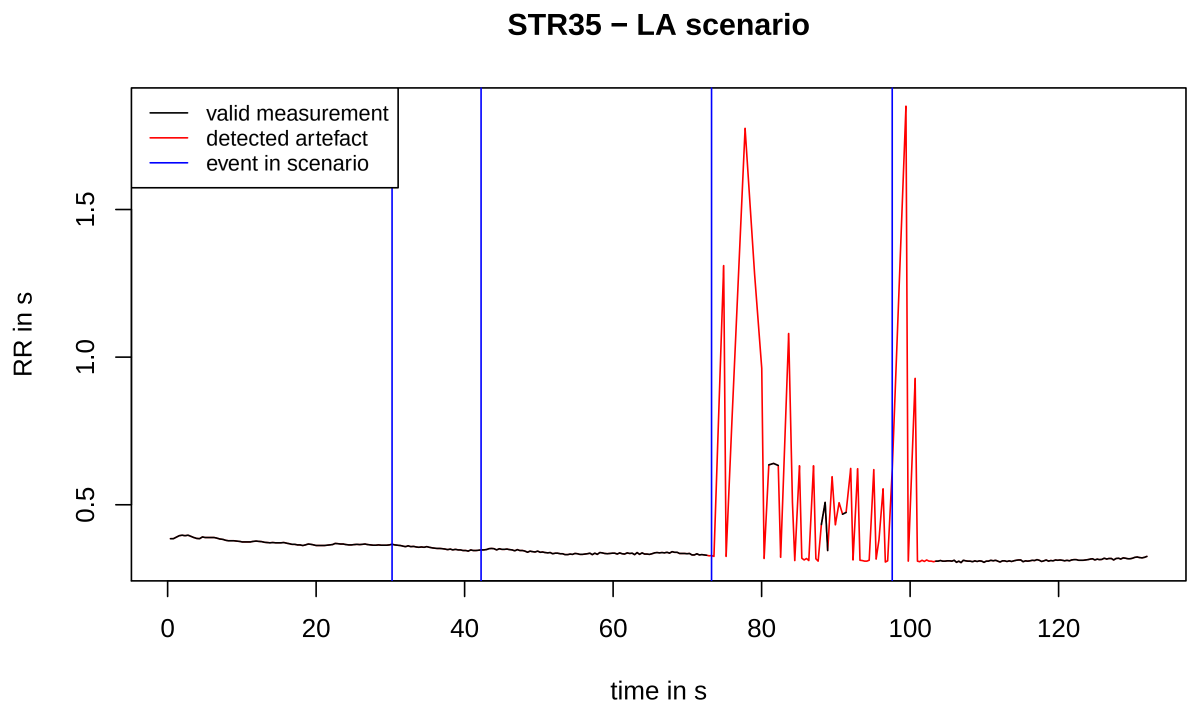
STR35 − LA scenario
valid measurement
detected artefact
event in scenario
RR in s
0.5
1.0
1.5
0
20
40
60
80
100
120
time in s

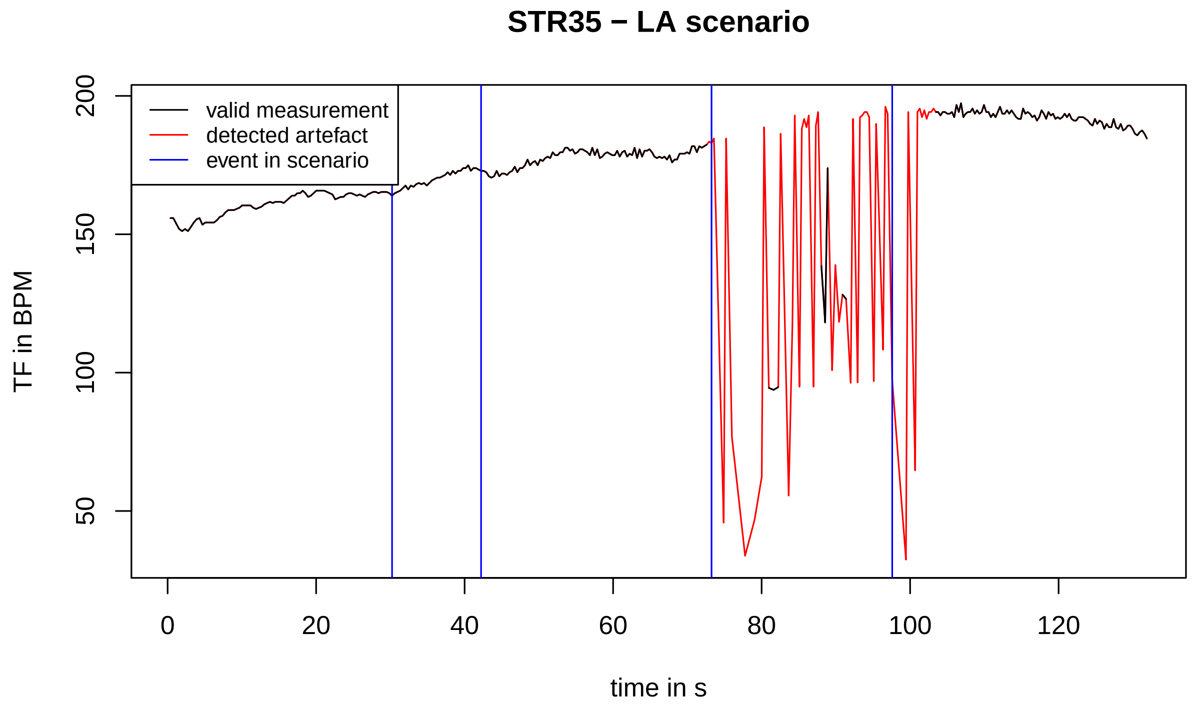
STR35 − LA scenario
valid measurement
detected artefact
event in scenario
TF in BPM
50
100
150
200
0
20
40
60
80
100
120
time in s

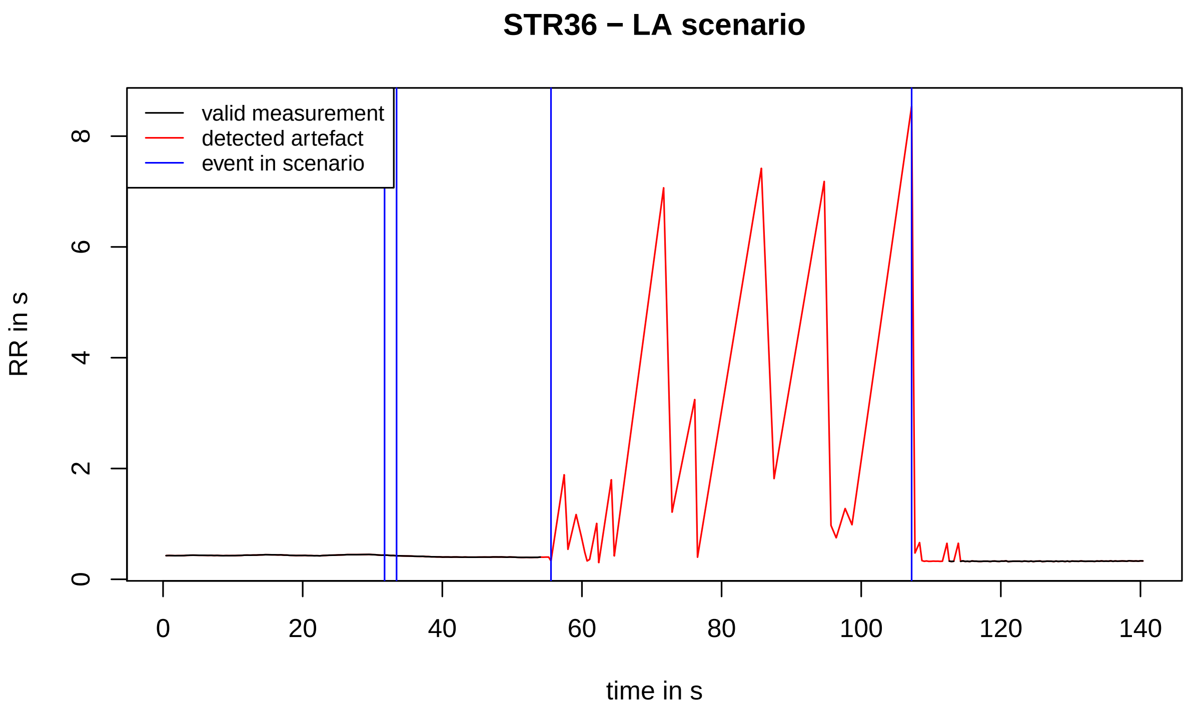
STR36 − LA scenario
valid measurement
detected artefact
event in scenario
RR in s
0
2
4
6
8
0
20
40
60
80
100
120
140
time in s

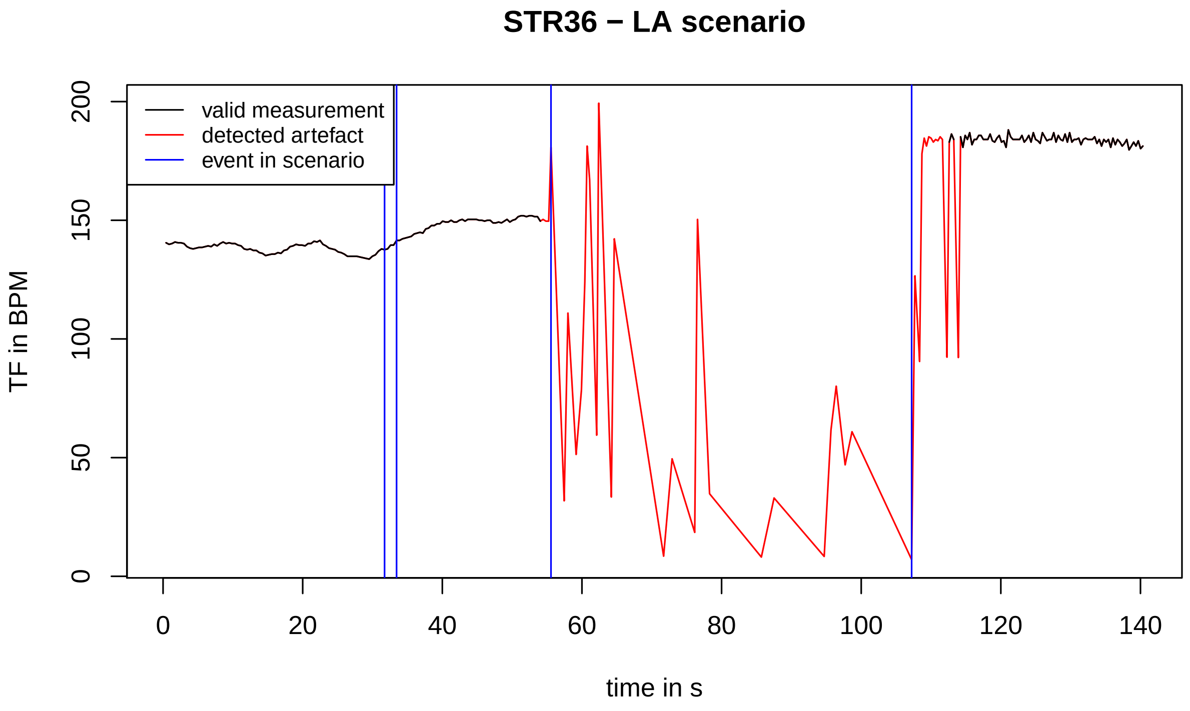
STR36 − LA scenario
valid measurement
detected artefact
event in scenario
TF in BPM
0
50
100
150
200
0
20
40
60
80
100
120
140
time in s

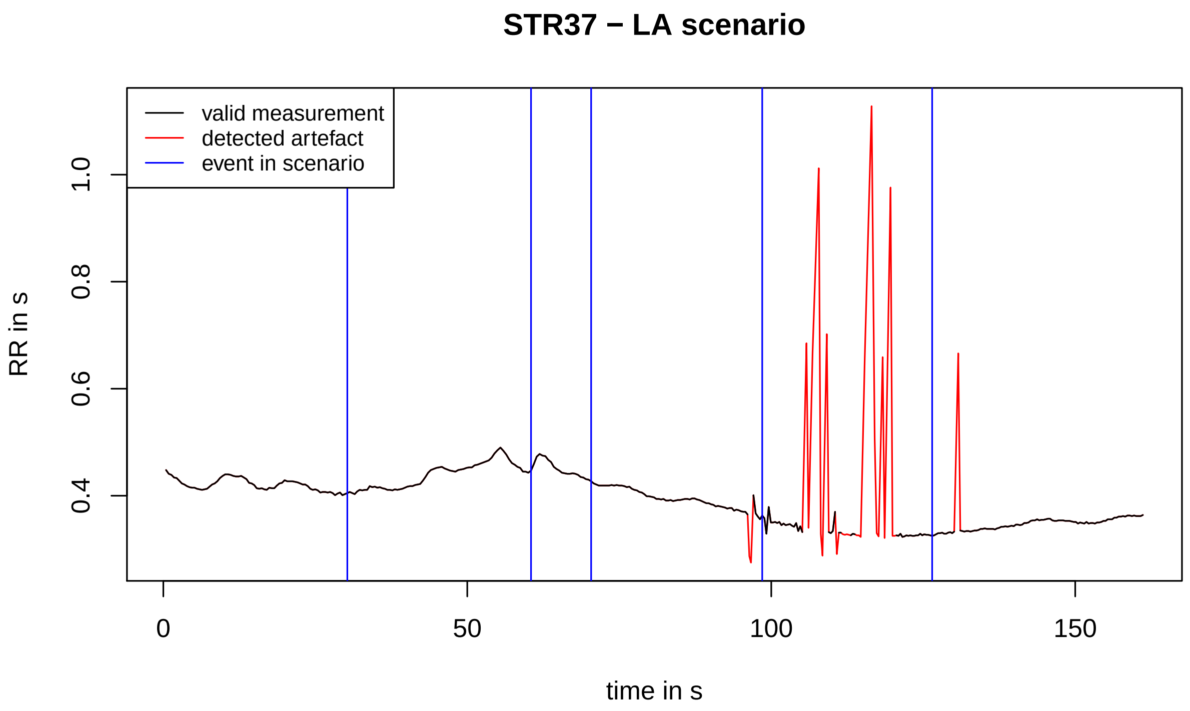
STR37 − LA scenario
valid measurement
detected artefact
event in scenario
RR in s
1.0
0.8
0.6
0.4
0
50
100
150
time in s

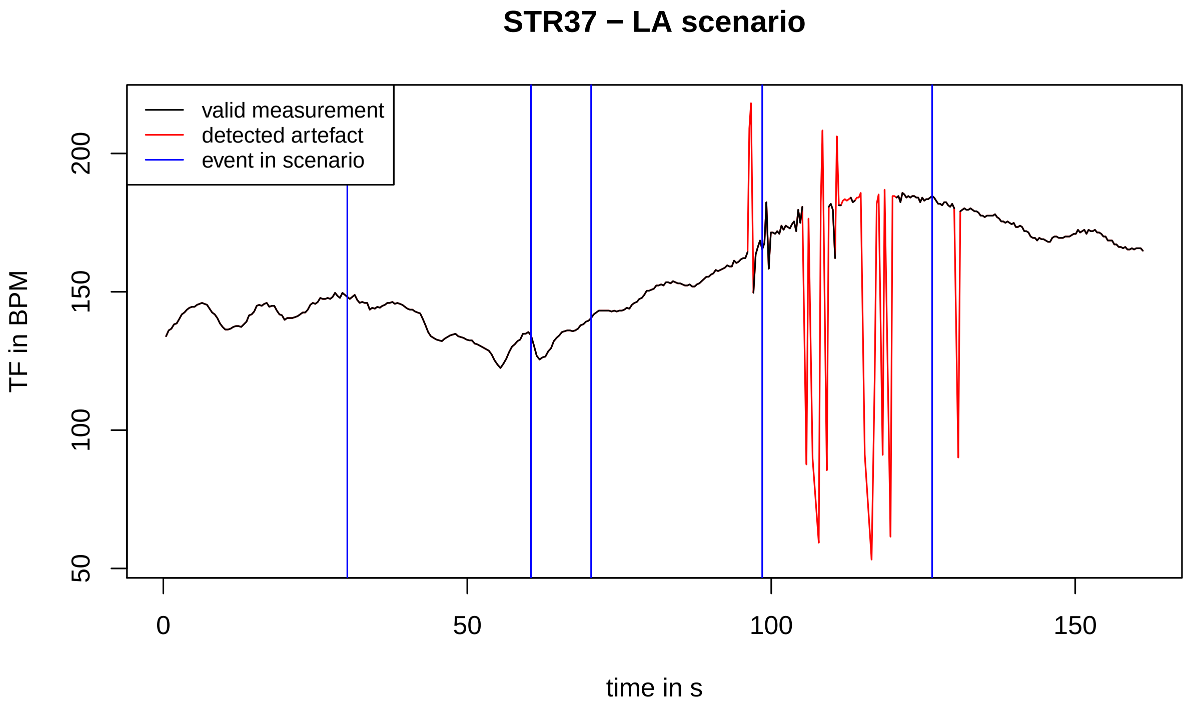
STR37 − LA scenario
valid measurement
detected artefact
event in scenario
TF in BPM
200
150
100
50
0
50
100
150
time in s

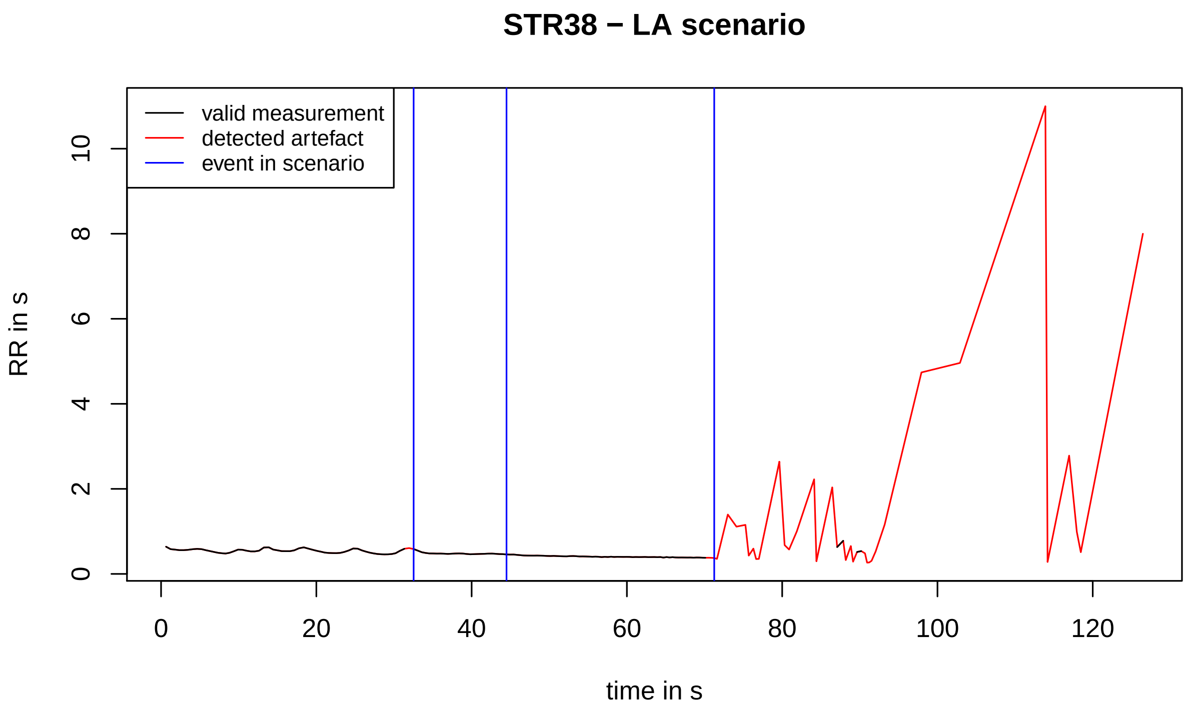
STR38 − LA scenario
valid measurement
detected artefact
event in scenario
RR in s
0
2
4
6
8
10
0
20
40
60
80
100
120
time in s

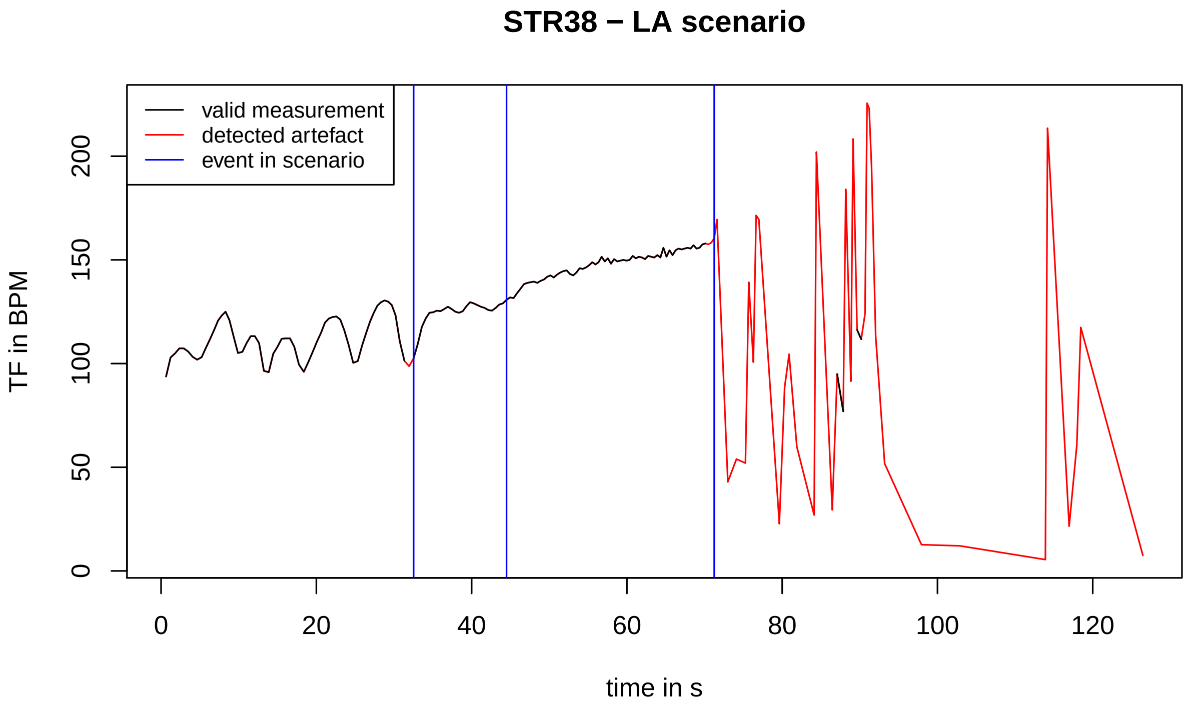
STR38 − LA scenario
valid measurement
detected artefact
event in scenario
TF in BPM
0
50
100
150
200
0
20
40
60
80
100
120
time in s

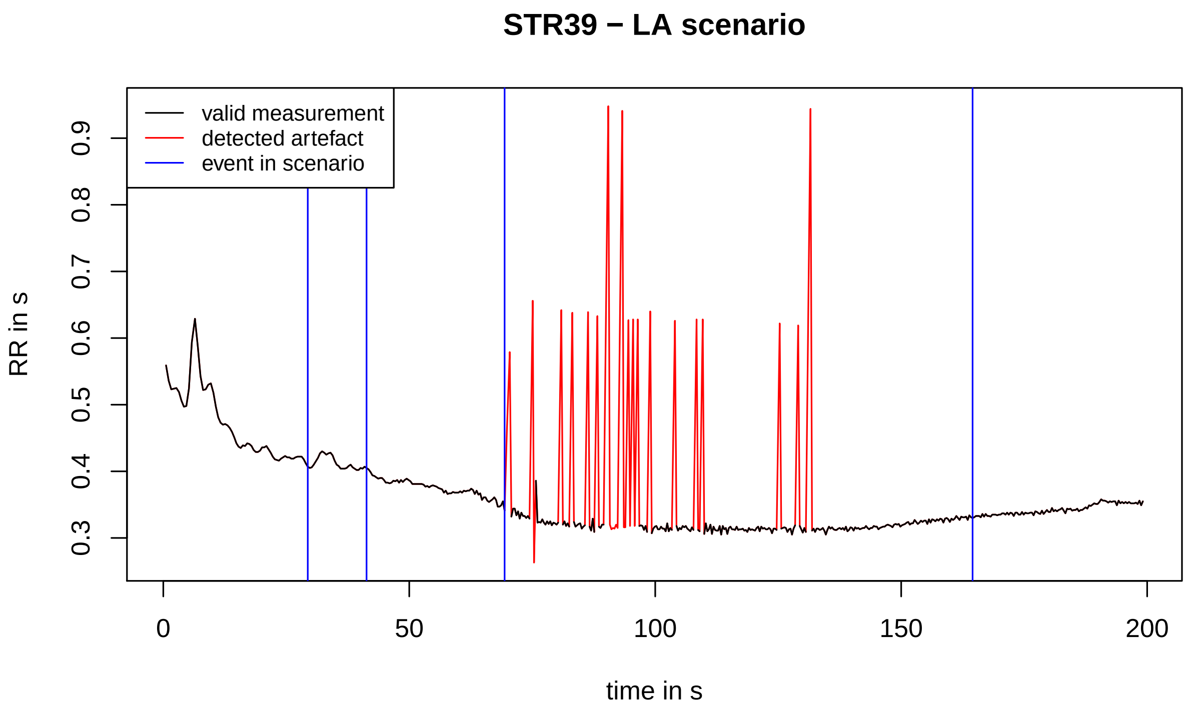
STR39 − LA scenario
valid measurement
detected artefact
event in scenario
RR in s
0.3
0.4
0.5
0.6
0.7
0.8
0.9
0
50
100
150
200
time in s

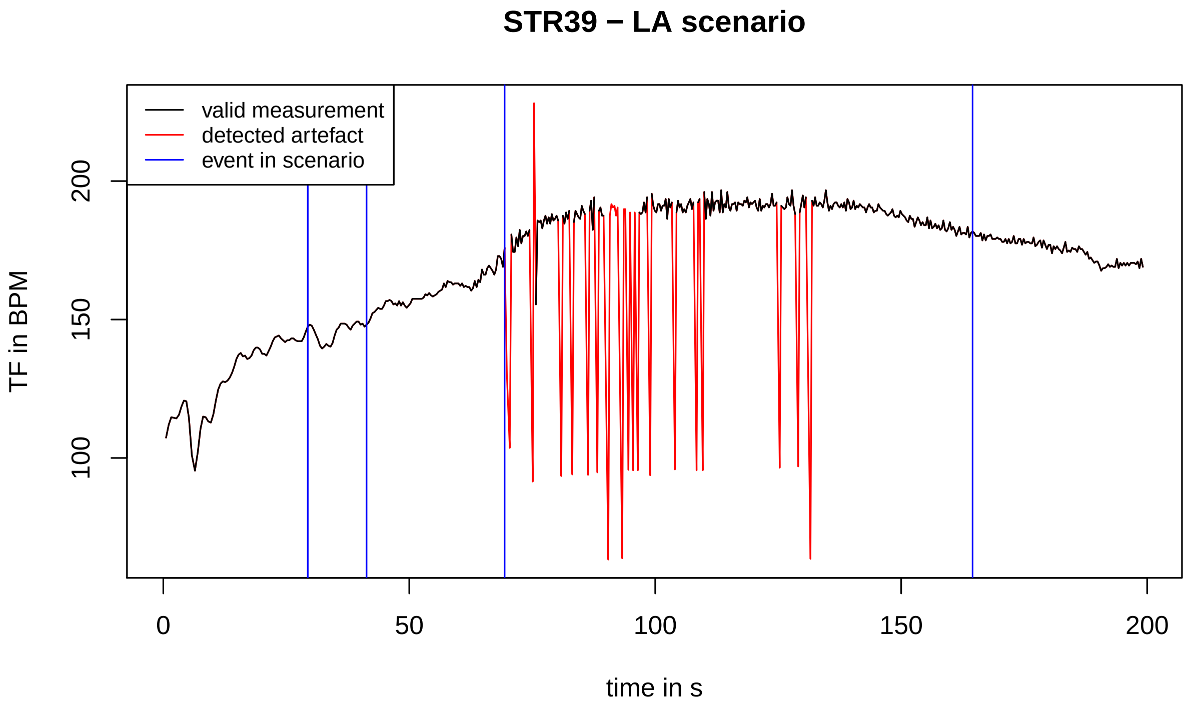
STR39 − LA scenario
valid measurement
detected artefact
event in scenario
TF in BPM
100
150
200
0
50
100
150
200
time in s

#### STR40 − LA scenario

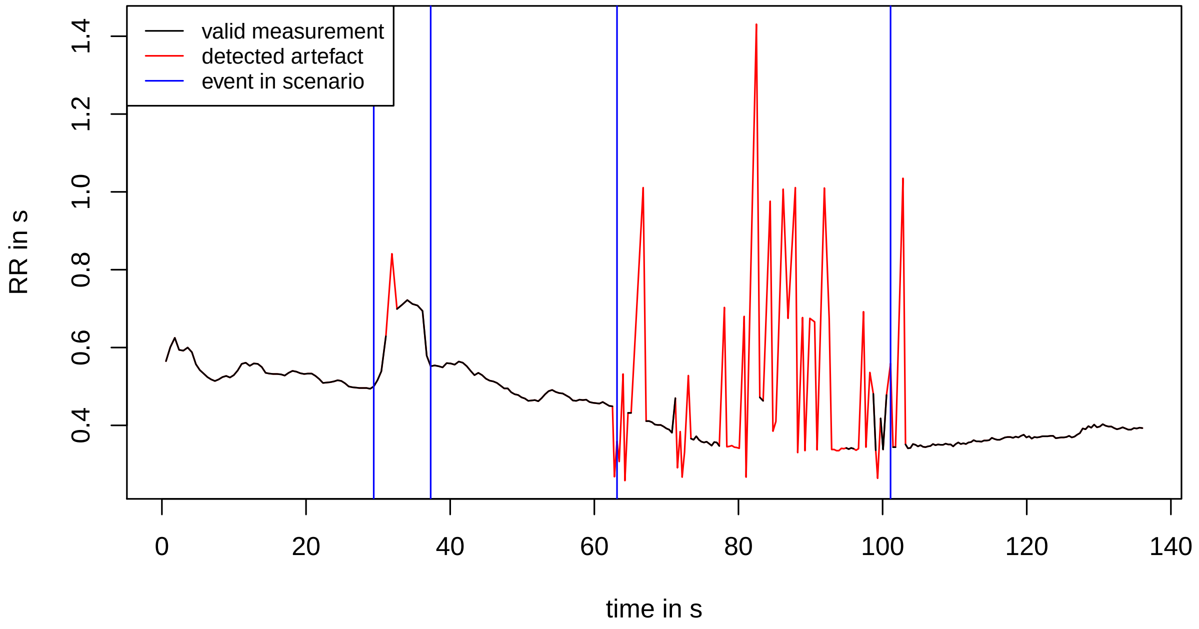


#### STR40 − LA scenario

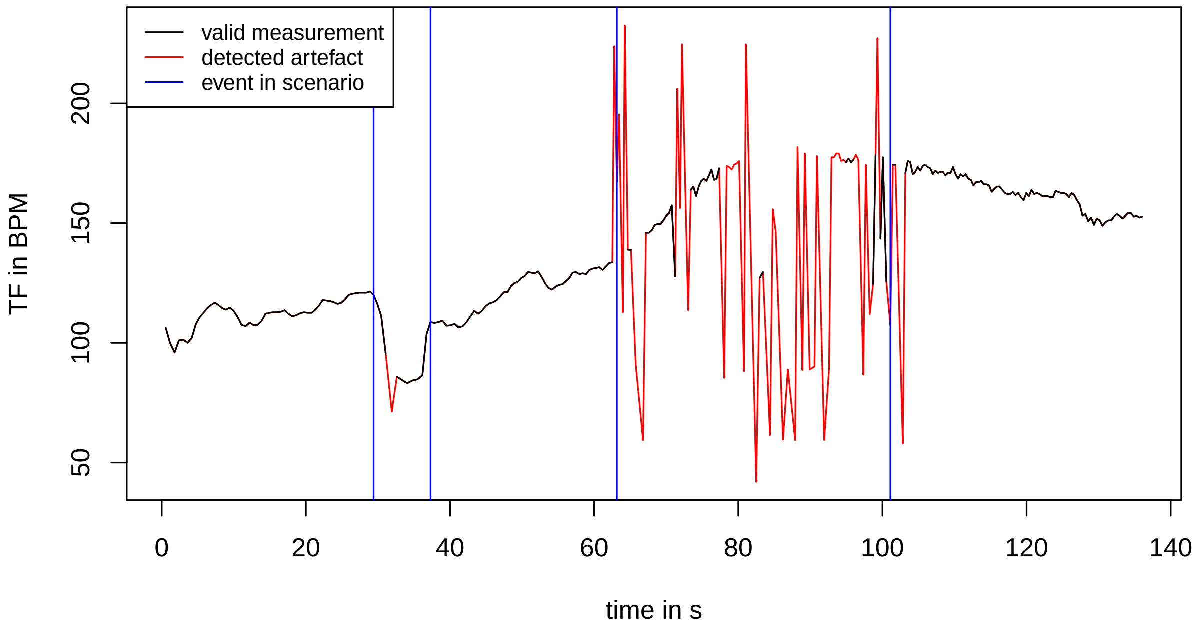

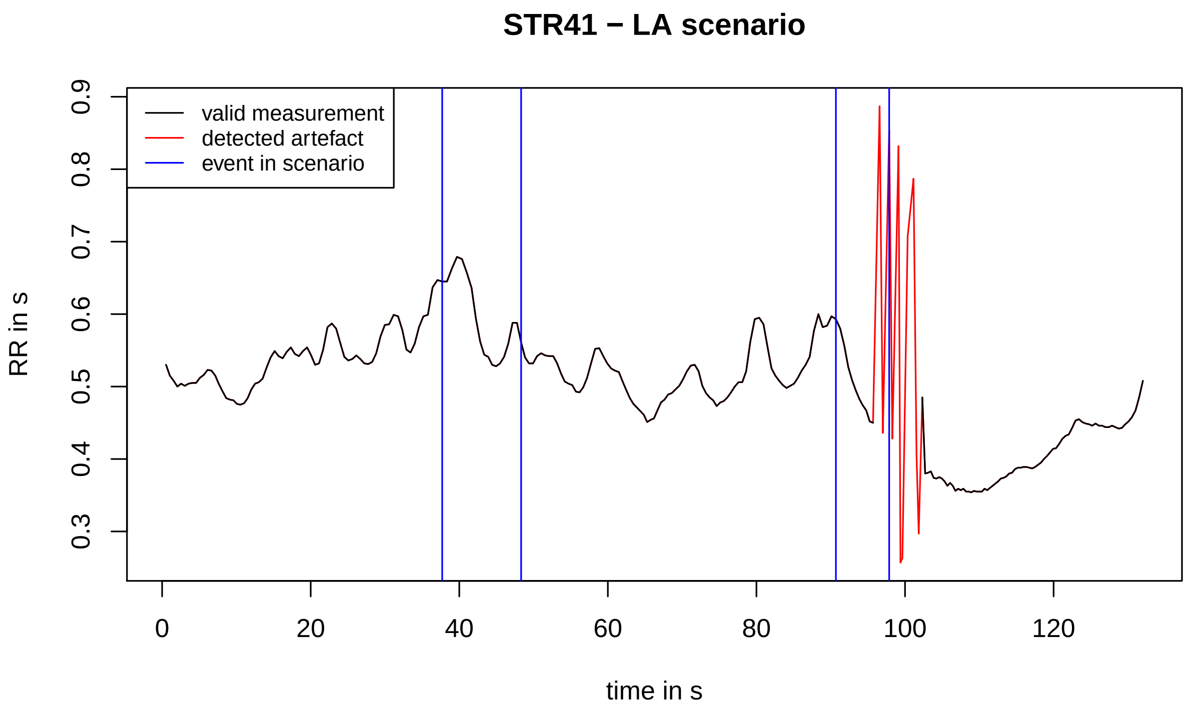
STR41 − LA scenario
RR in s
time in s
valid measurement
detected artefact
event in scenario
0.3
0.4
0.5
0.6
0.7
0.8
0.9
0
20
40
60
80
100
120

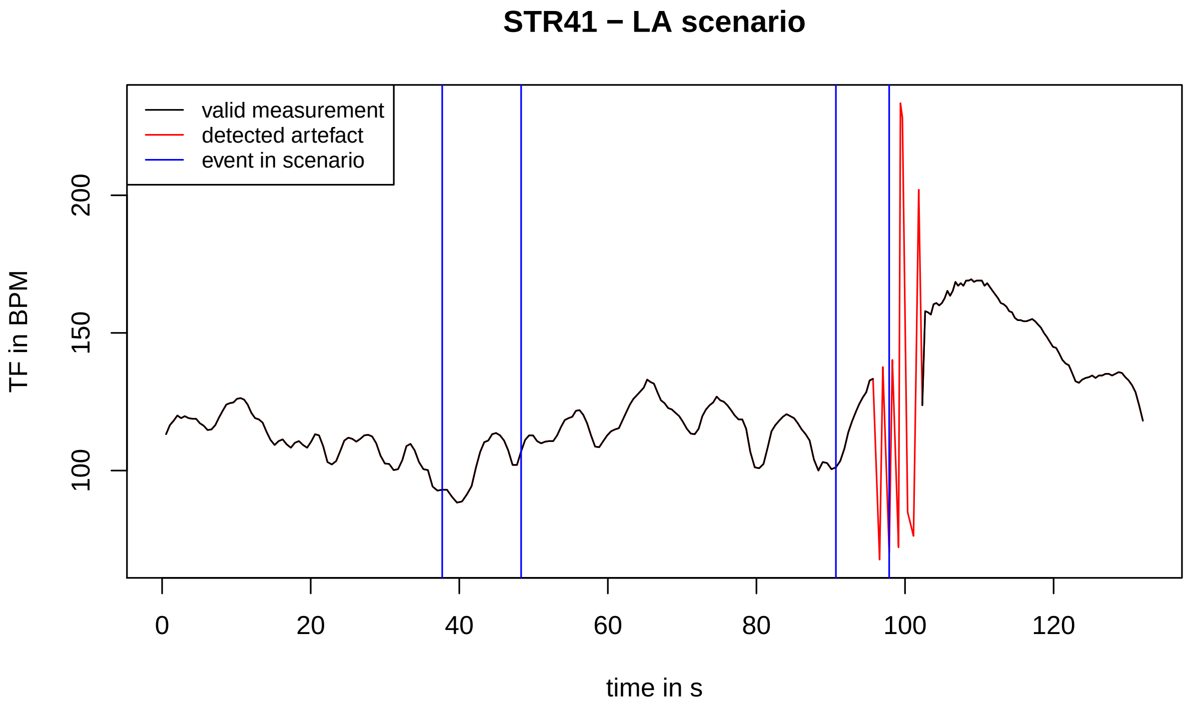
STR41 − LA scenario
TF in BPM
time in s
valid measurement
detected artefact
event in scenario
100
150
200
0
20
40
60
80
100
120

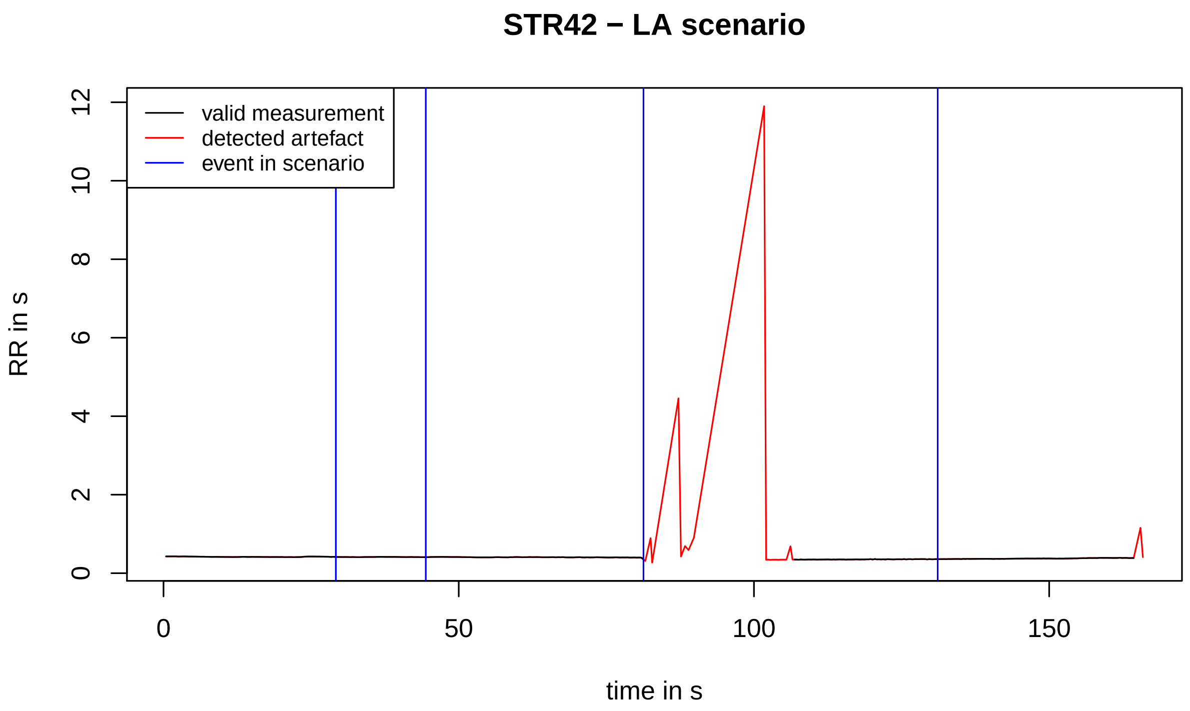
STR42 − LA scenario
valid measurement
detected artefact
event in scenario
RR in s
0
2
4
6
8
10
12
0
50
100
150
time in s

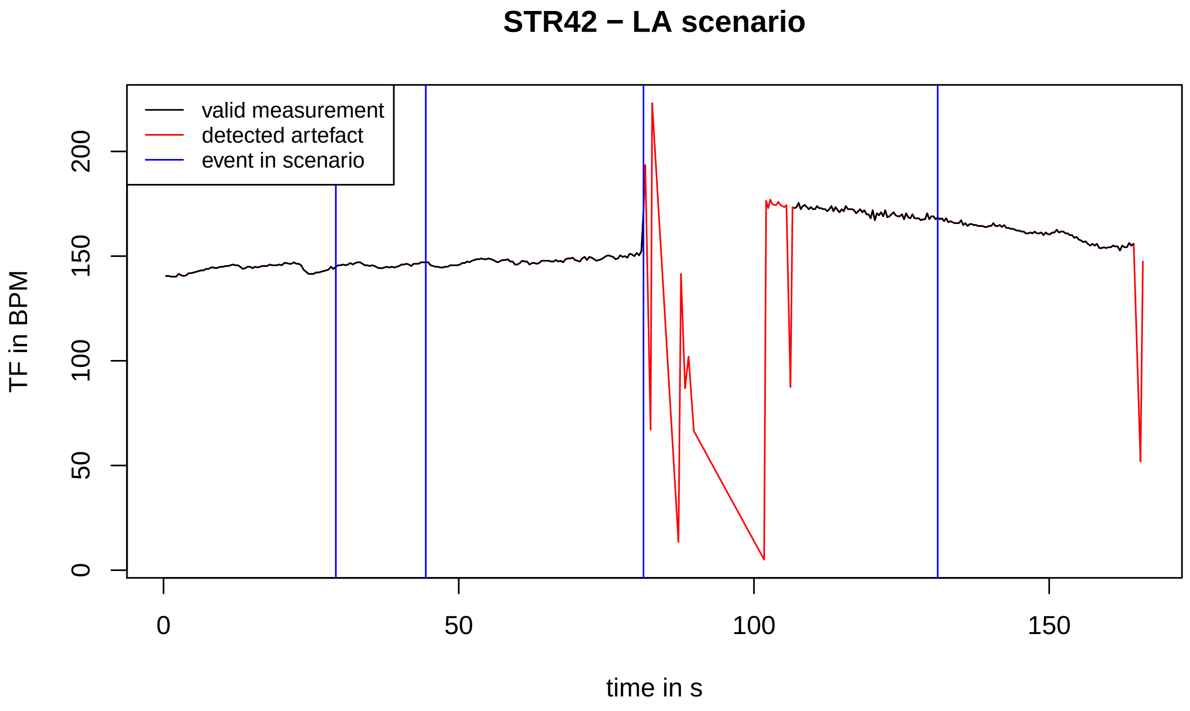
STR42 − LA scenario
valid measurement
detected artefact
event in scenario
TF in BPM
0
50
100
150
200
0
50
100
150
time in s

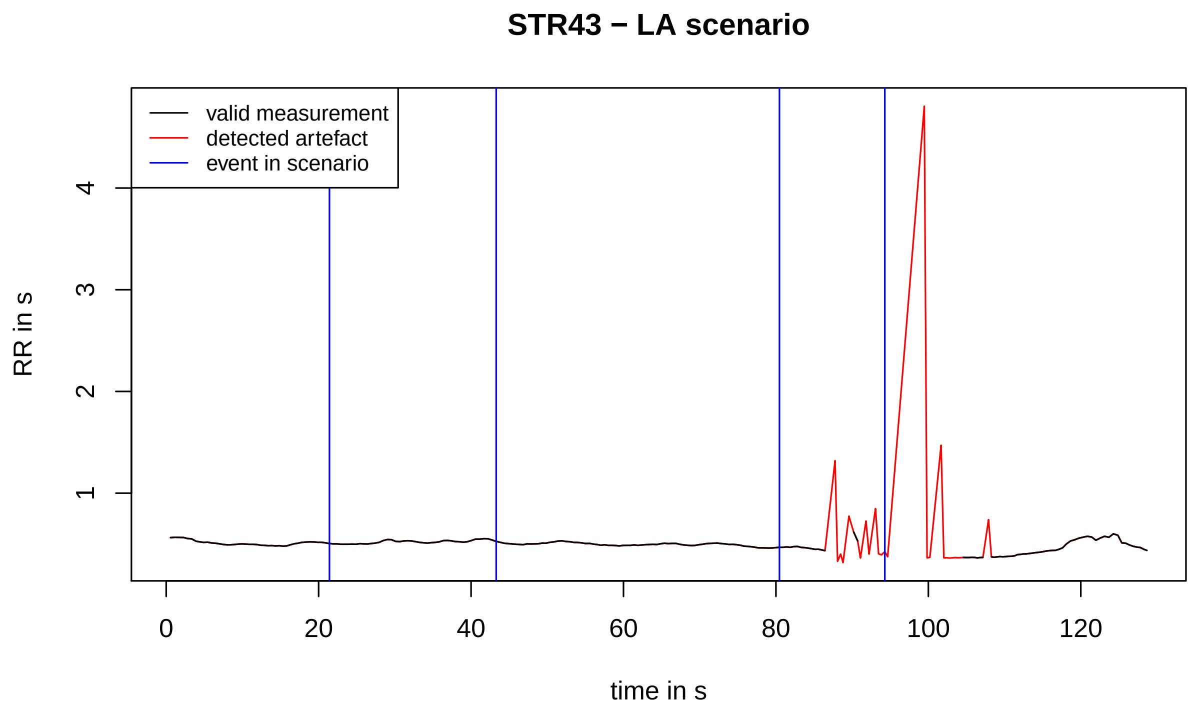
STR43 − LA scenario
valid measurement
detected artefact
event in scenario
RR in s
time in s

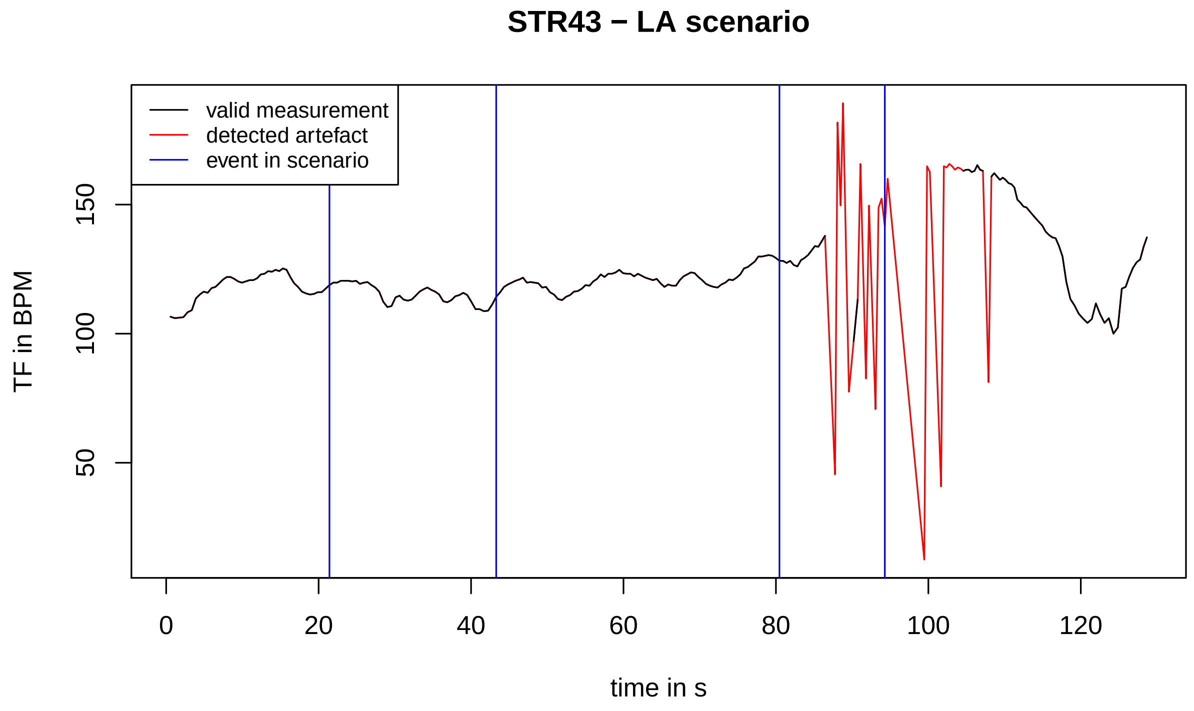
STR43 − LA scenario
valid measurement
detected artefact
event in scenario
TF in BPM
time in s

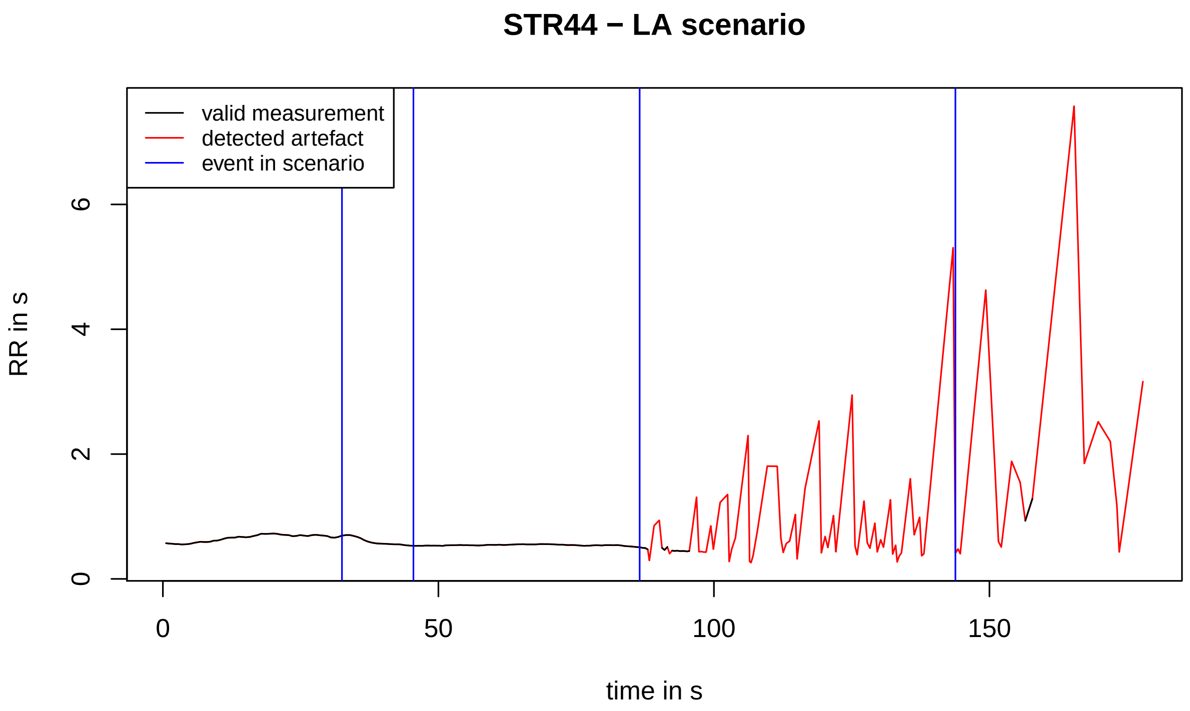

STR44 − LA scenario
valid measurement
detected artefact
event in scenario
RR in s
0
2
4
6
0
50
100
150
time in s


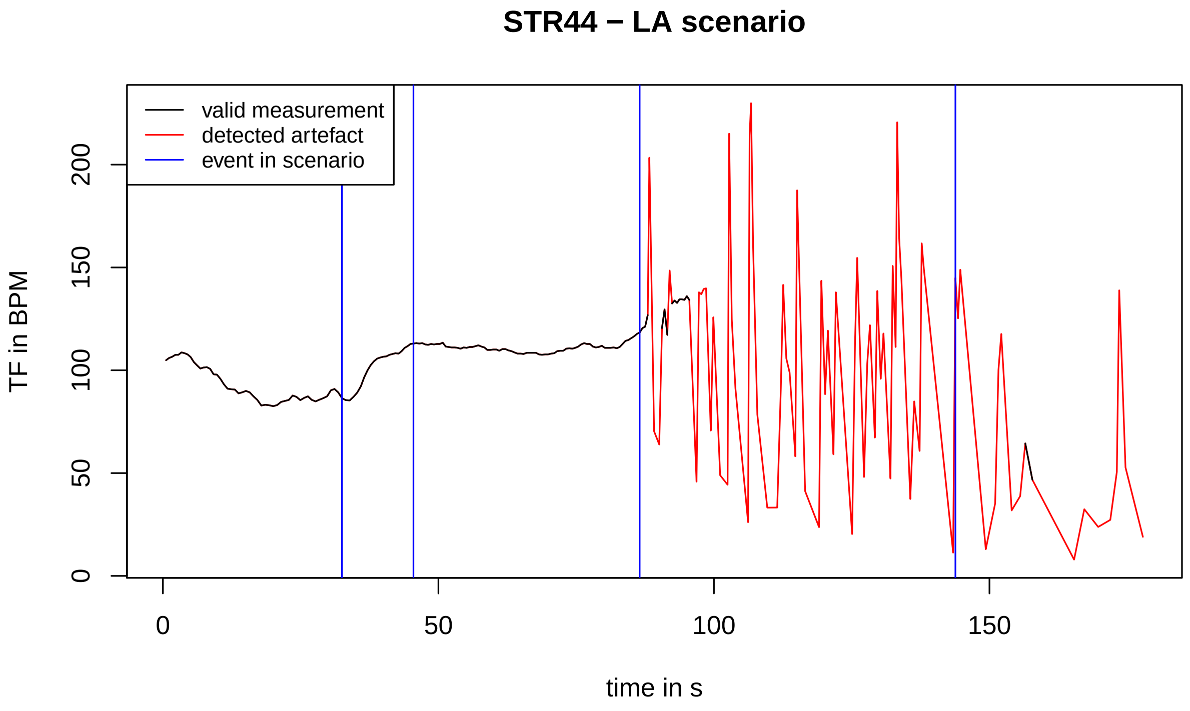

STR44 − LA scenario
valid measurement
detected artefact
event in scenario
TF in BPM
0
50
100
150
200
0
50
100
150
time in s

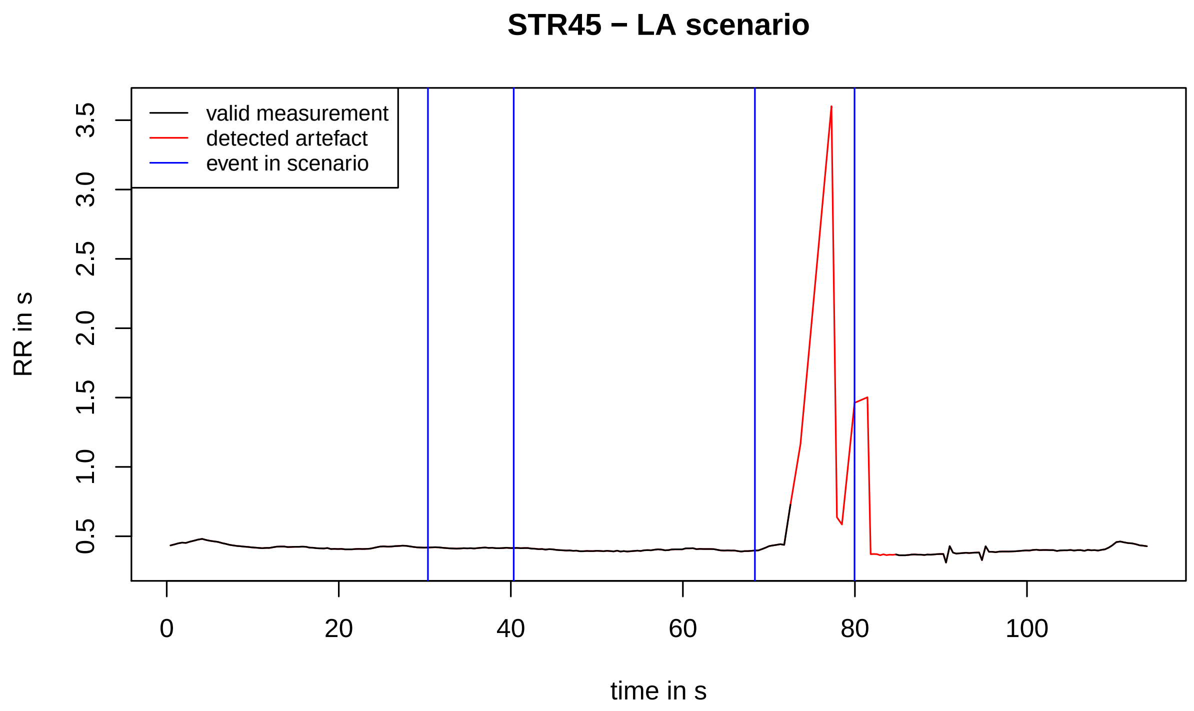
STR45 − LA scenario
valid measurement
detected artefact
event in scenario
RR in s
0.5
1.0
1.5
2.0
2.5
3.0
3.5
0
20
40
60
80
100
time in s

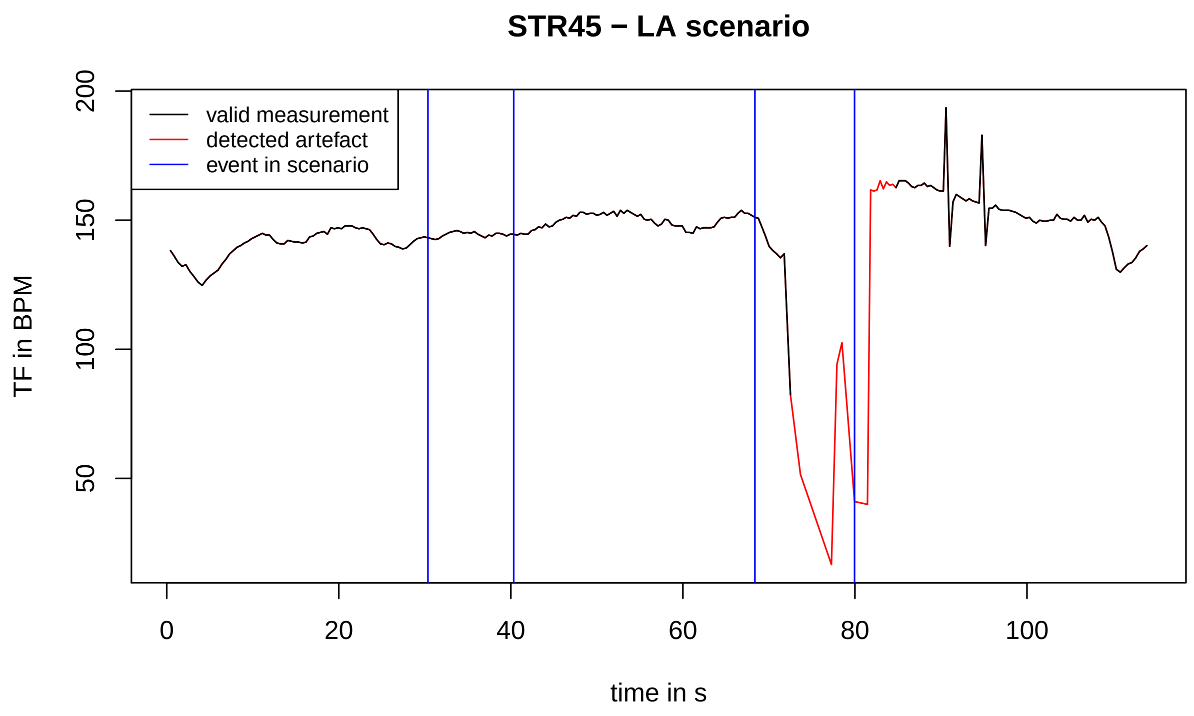
STR45 − LA scenario
valid measurement
detected artefact
event in scenario
TF in BPM
50
100
150
200
0
20
40
60
80
100
time in s

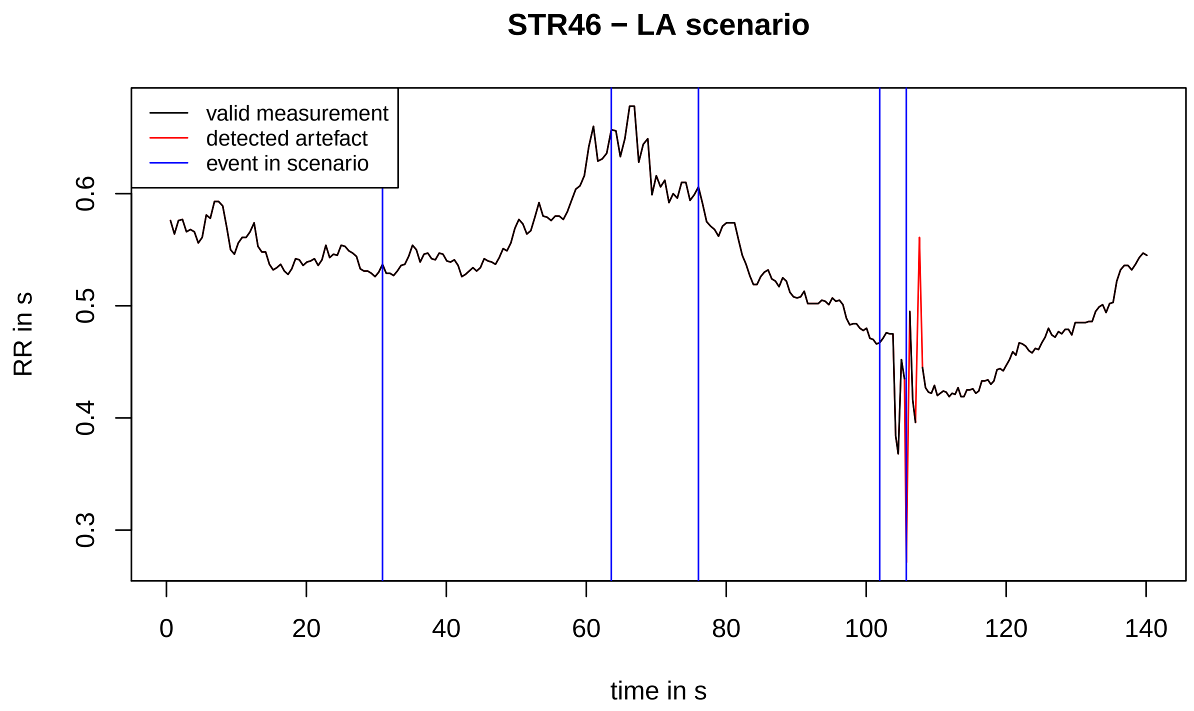
STR46 − LA scenario
valid measurement
detected artefact
event in scenario
RR in s
0.3
0.4
0.5
0.6
0
20
40
60
80
100
120
140
time in s

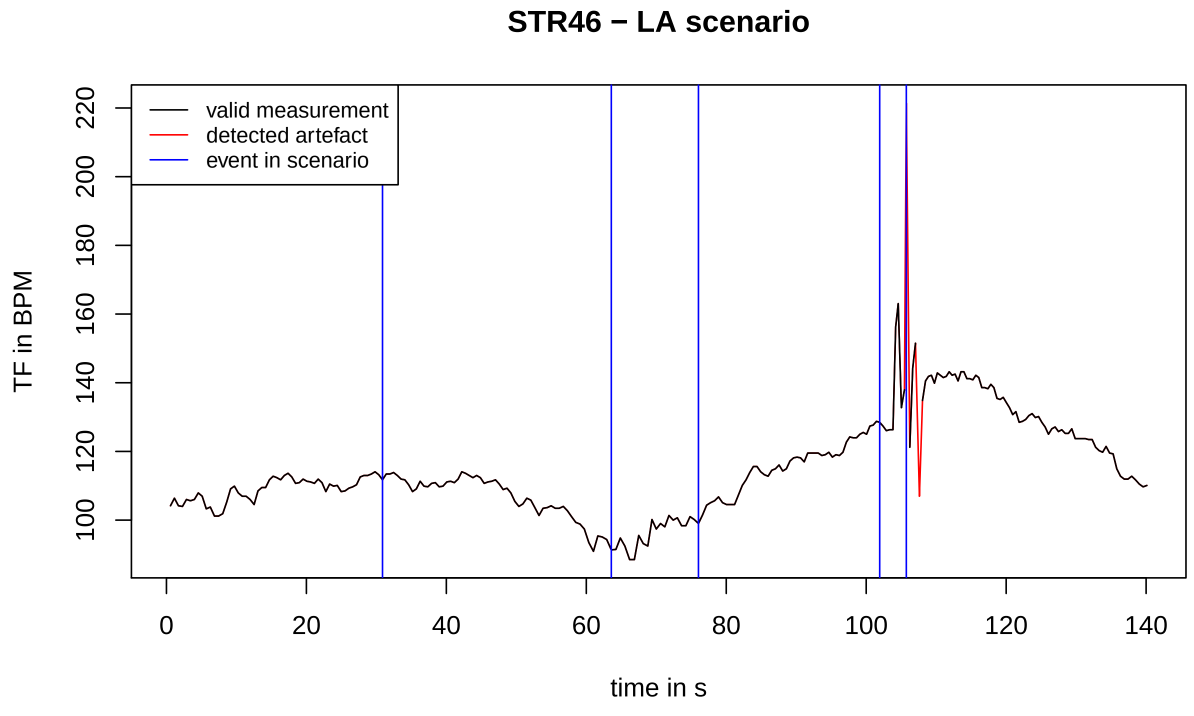
STR46 − LA scenario
valid measurement
detected artefact
event in scenario
TF in BPM
100
120
140
160
180
200
220
0
20
40
60
80
100
120
140
time in s

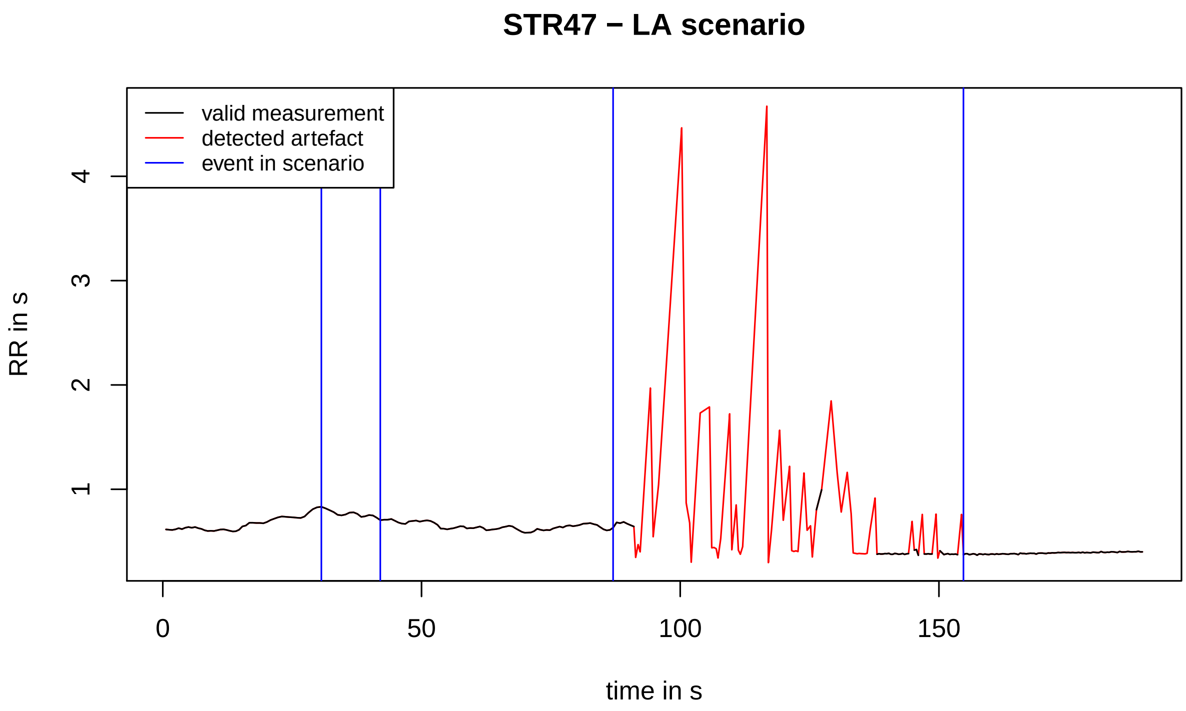
STR47 − LA scenario
valid measurement
detected artefact
event in scenario
RR in s
1
2
3
4
0
50
100
150
time in s

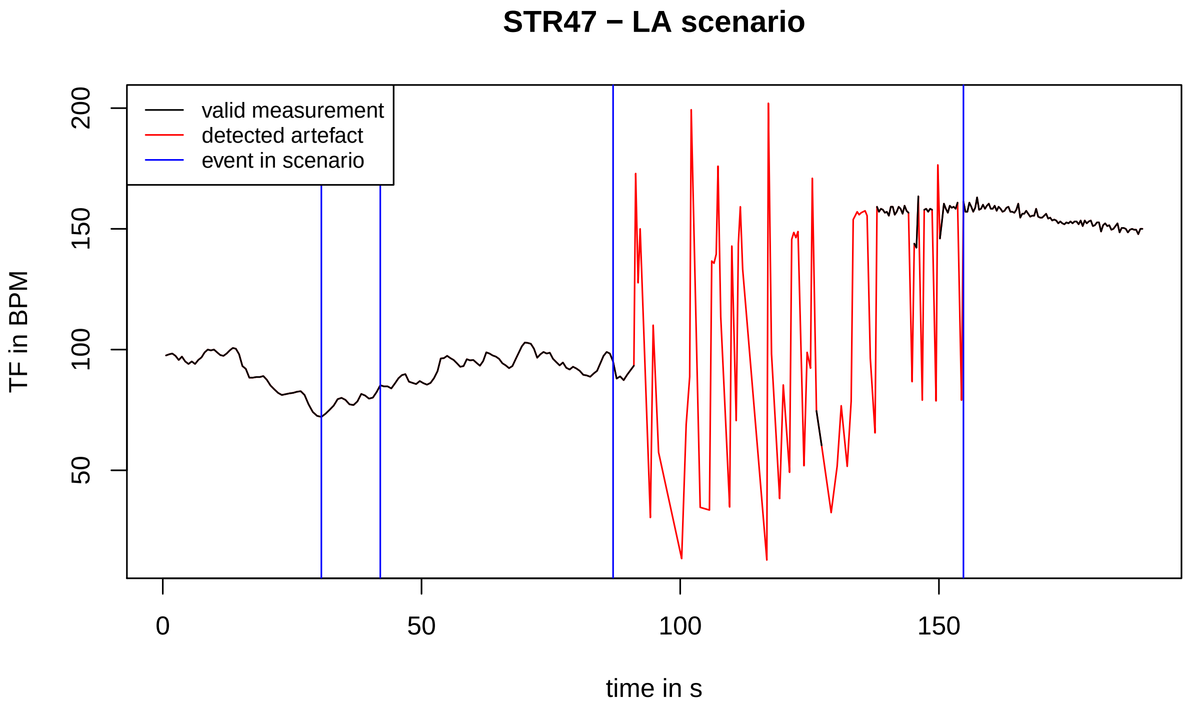
STR47 − LA scenario
valid measurement
detected artefact
event in scenario
TF in BPM
50
100
150
200
0
50
100
150
time in s

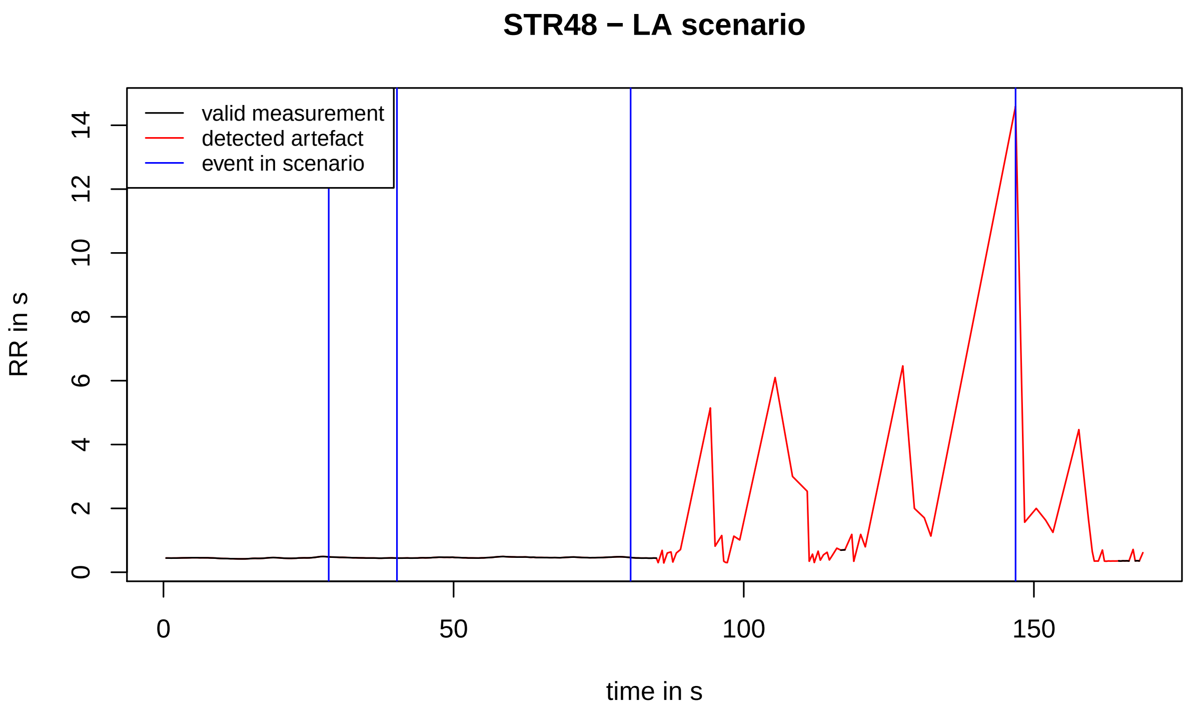
STR48 − LA scenario
valid measurement
detected artefact
event in scenario
RR in s
0
2
4
6
8
10
12
14
0
50
100
150
time in s

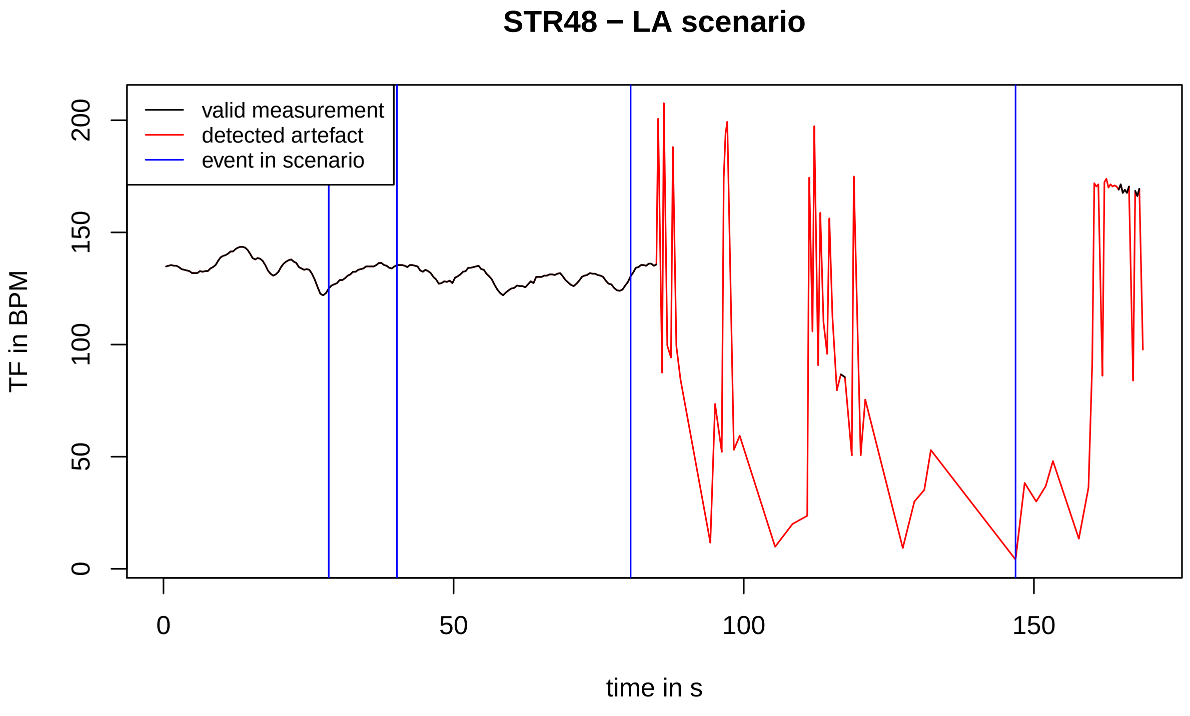
STR48 − LA scenario
valid measurement
detected artefact
event in scenario
TF in BPM
0
50
100
150
200
0
50
100
150
time in s

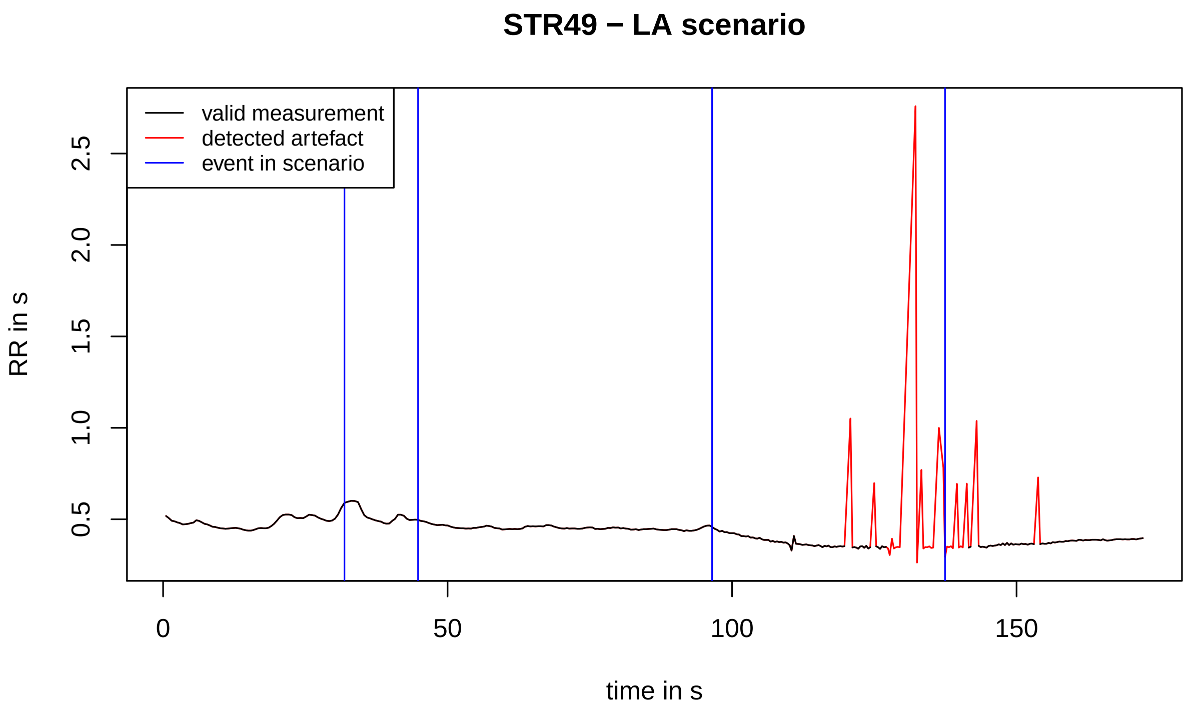
STR49 − LA scenario
valid measurement
detected artefact
event in scenario
RR in s
0.5
1.0
1.5
2.0
2.5
0
50
100
150
time in s

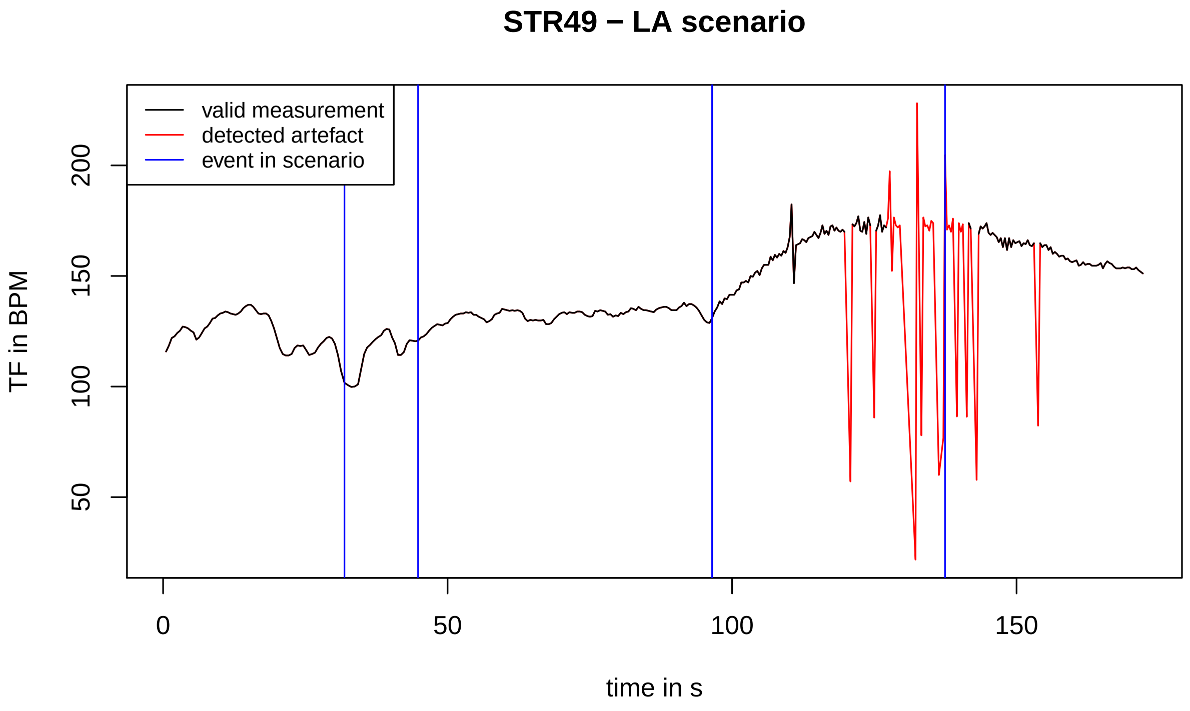
STR49 − LA scenario
valid measurement
detected artefact
event in scenario
TF in BPM
50
100
150
200
0
50
100
150
time in s

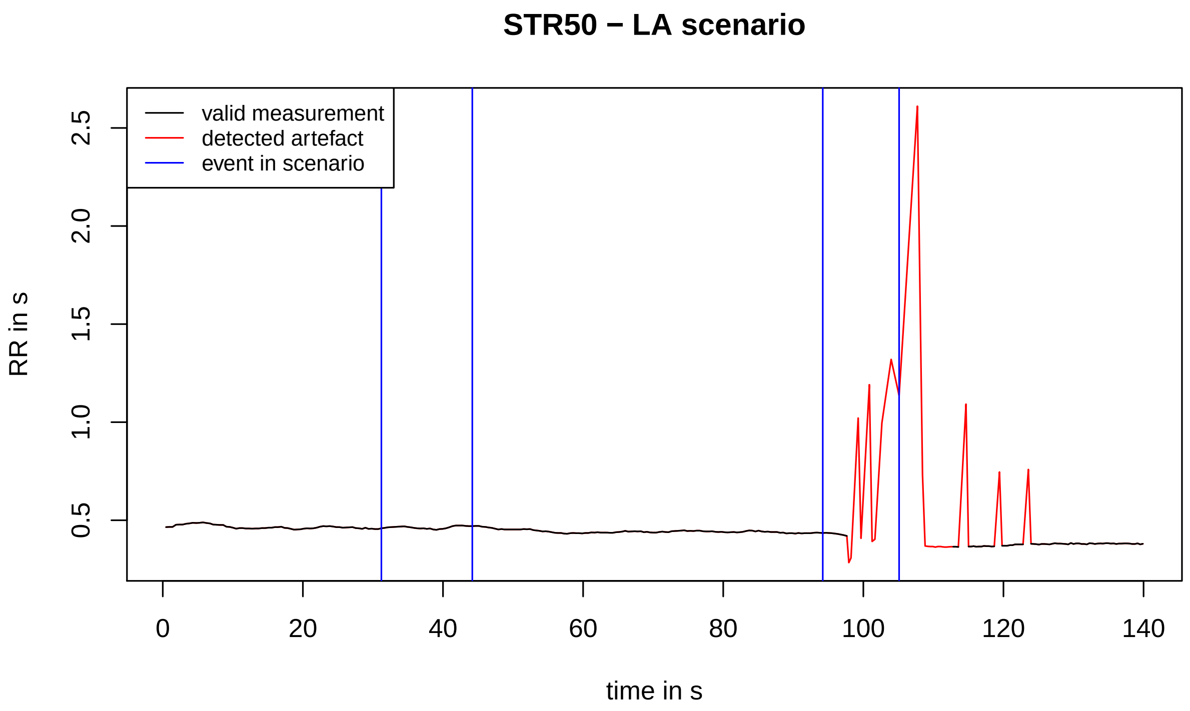
STR50 − LA scenario
valid measurement
detected artefact
event in scenario
RR in s
0.5
1.0
1.5
2.0
2.5
0
20
40
60
80
100
120
140
time in s

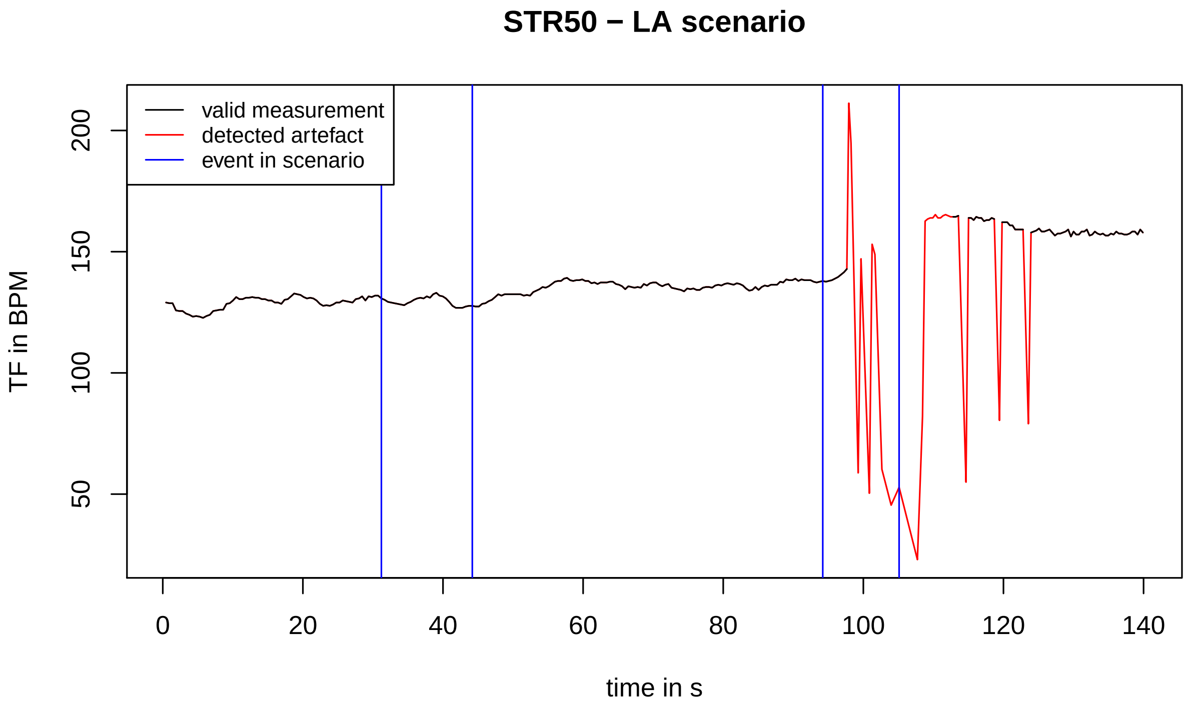
STR50 − LA scenario
valid measurement
detected artefact
event in scenario
TF in BPM
50
100
150
200
0
20
40
60
80
100
120
140
time in s

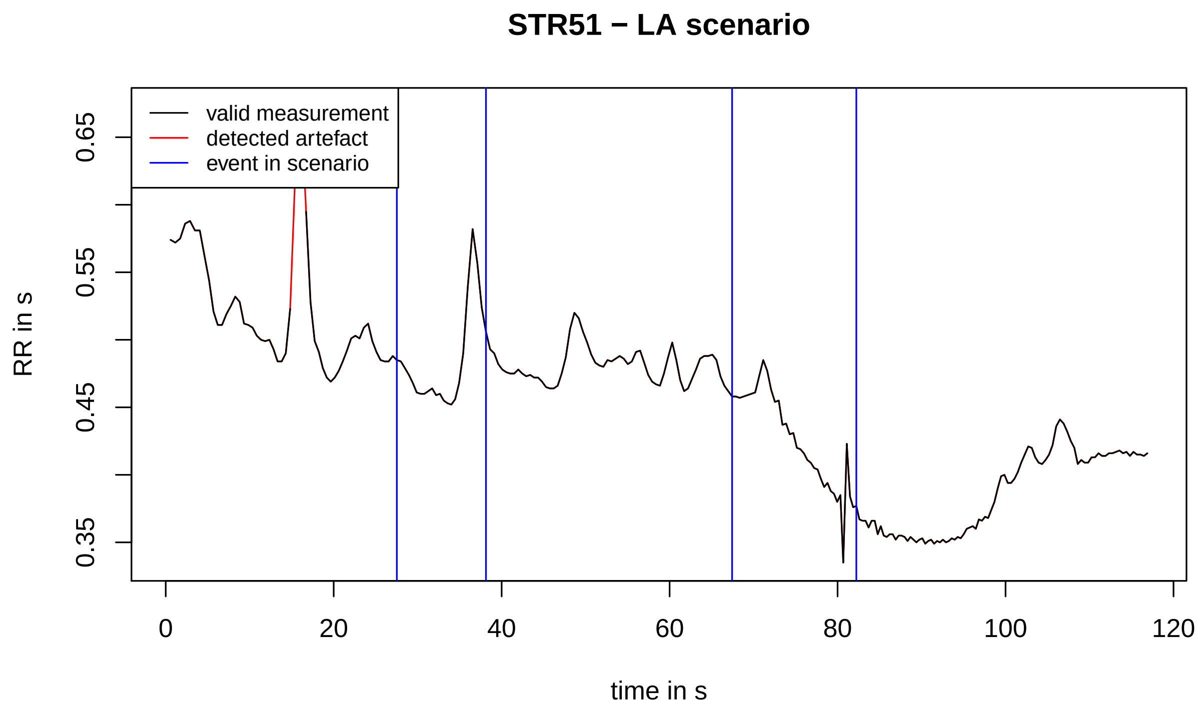
STR51 − LA scenario
valid measurement
detected artefact
event in scenario
RR in s
0.35
0.45
0.55
0.65
0
20
40
60
80
100
120
time in s

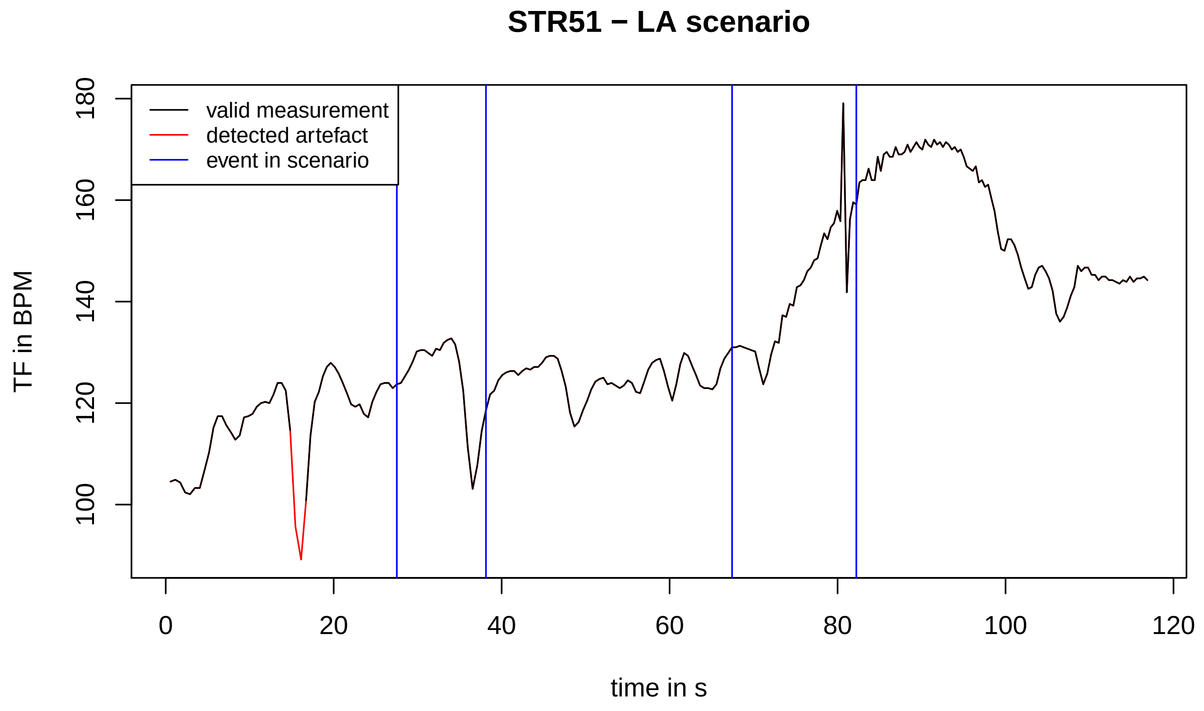
STR51 − LA scenario
valid measurement
detected artefact
event in scenario
TF in BPM
100
120
140
160
180
0
20
40
60
80
100
120
time in s

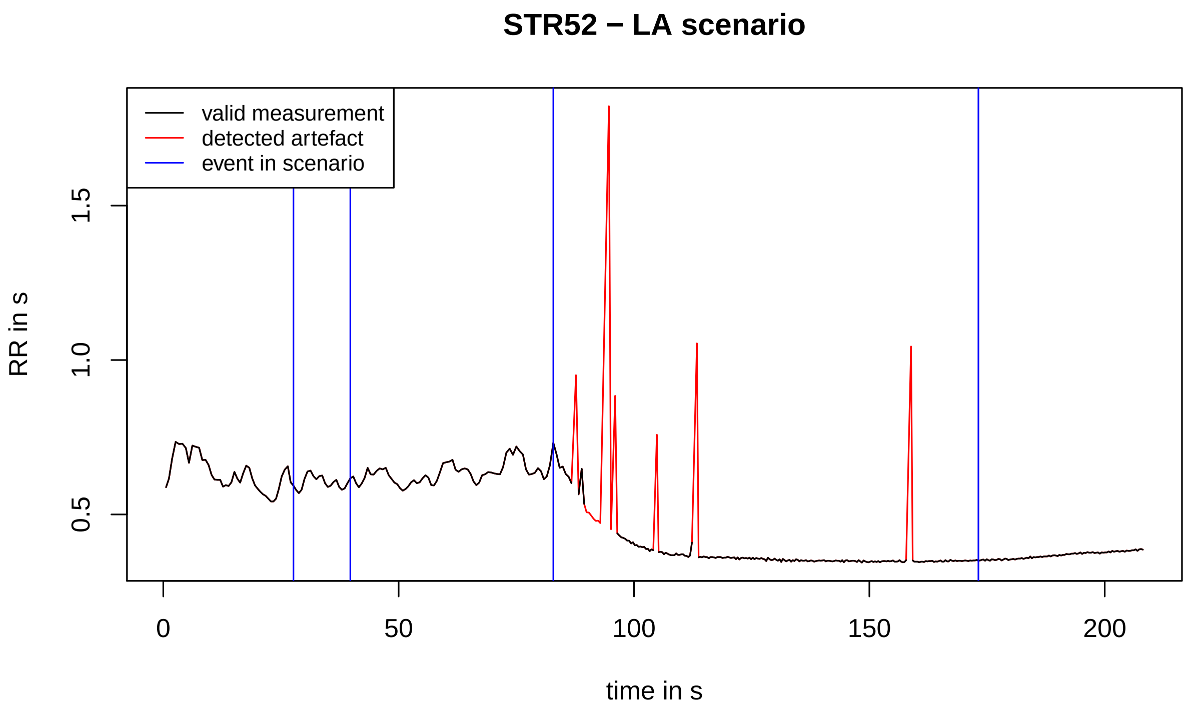
STR52 − LA scenario
valid measurement
detected artefact
event in scenario
RR in s
0.5
1.0
1.5
0
50
100
150
200
time in s

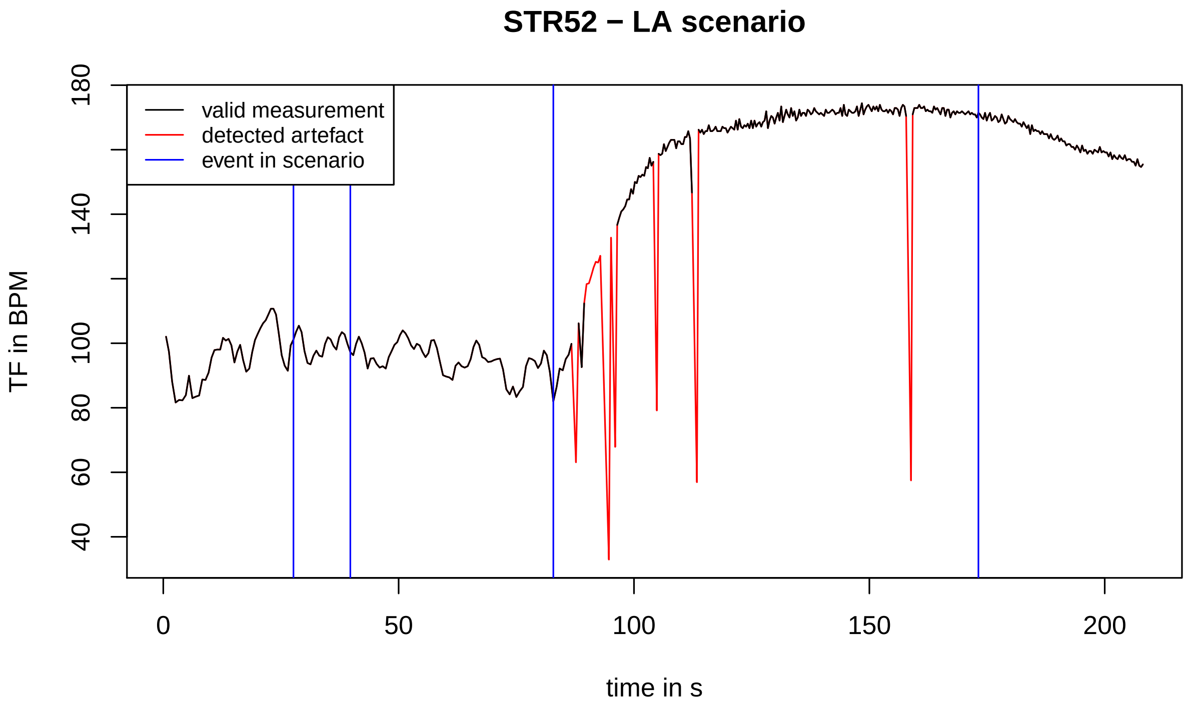
STR52 − LA scenario
valid measurement
detected artefact
event in scenario
TF in BPM
40
60
80
100
140
180
0
50
100
150
200
time in s

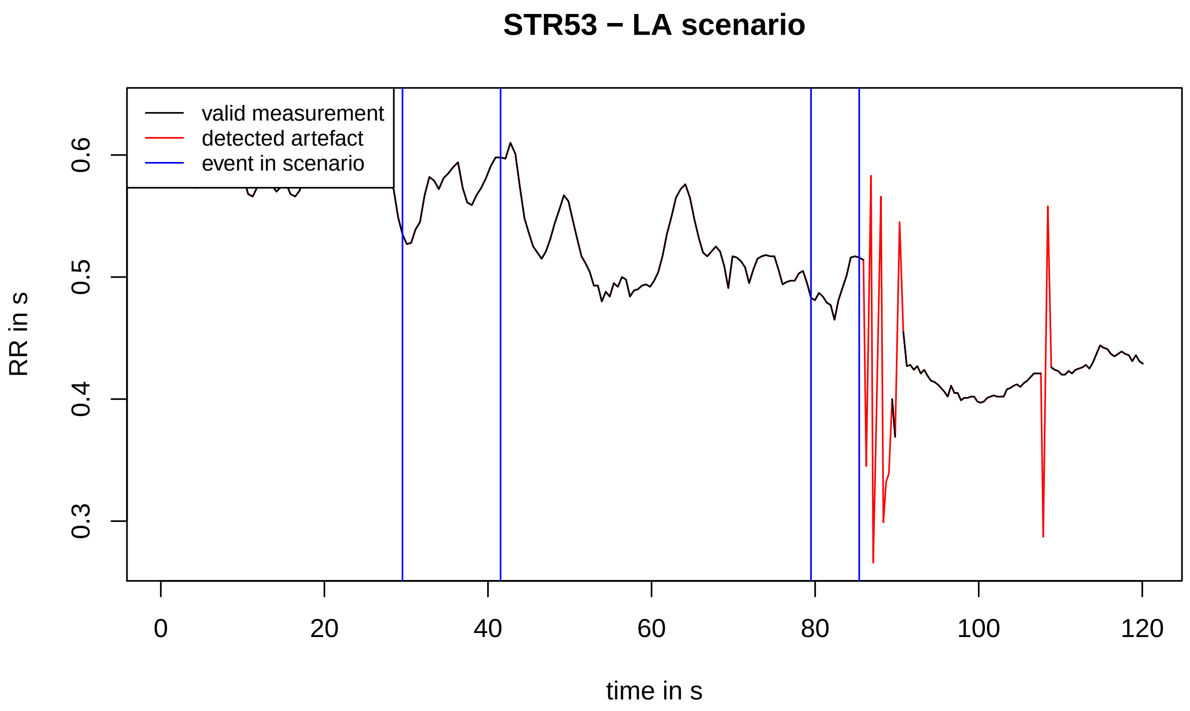
STR53 − LA scenario
valid measurement
detected artefact
event in scenario
RR in s
0.3
0.4
0.5
0.6
0
20
40
60
80
100
120
time in s

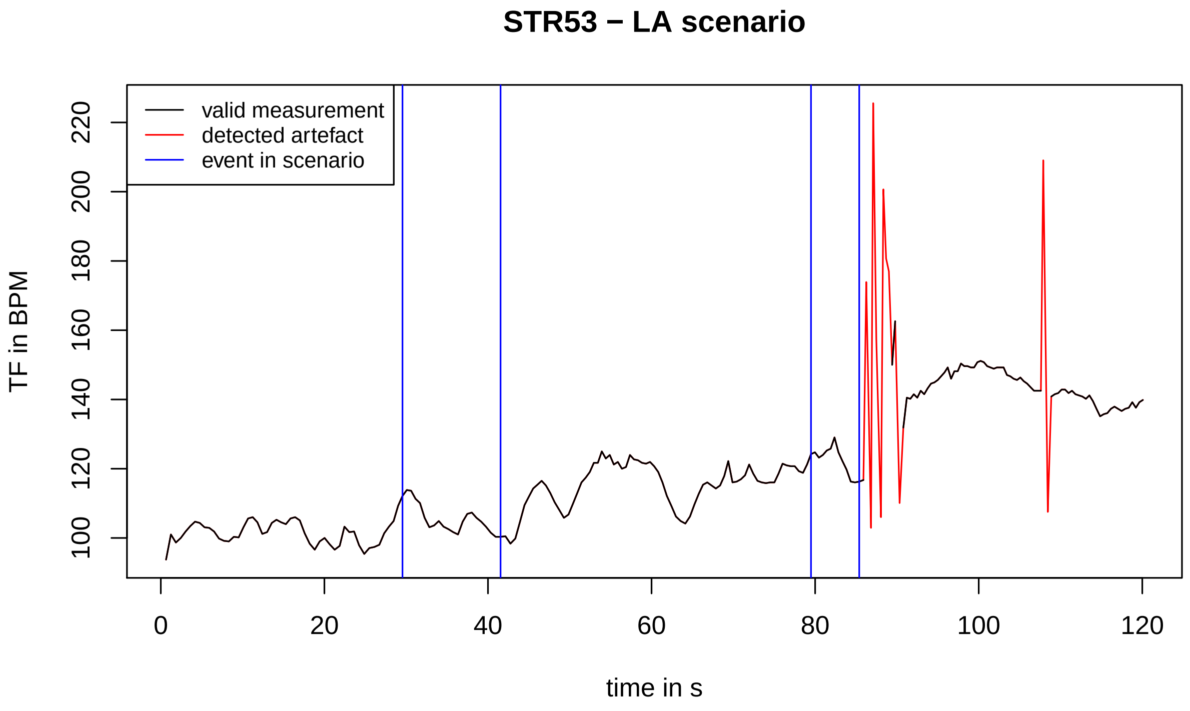
STR53 − LA scenario
valid measurement
detected artefact
event in scenario
TF in BPM
100
120
140
160
180
200
220
0
20
40
60
80
100
120
time in s

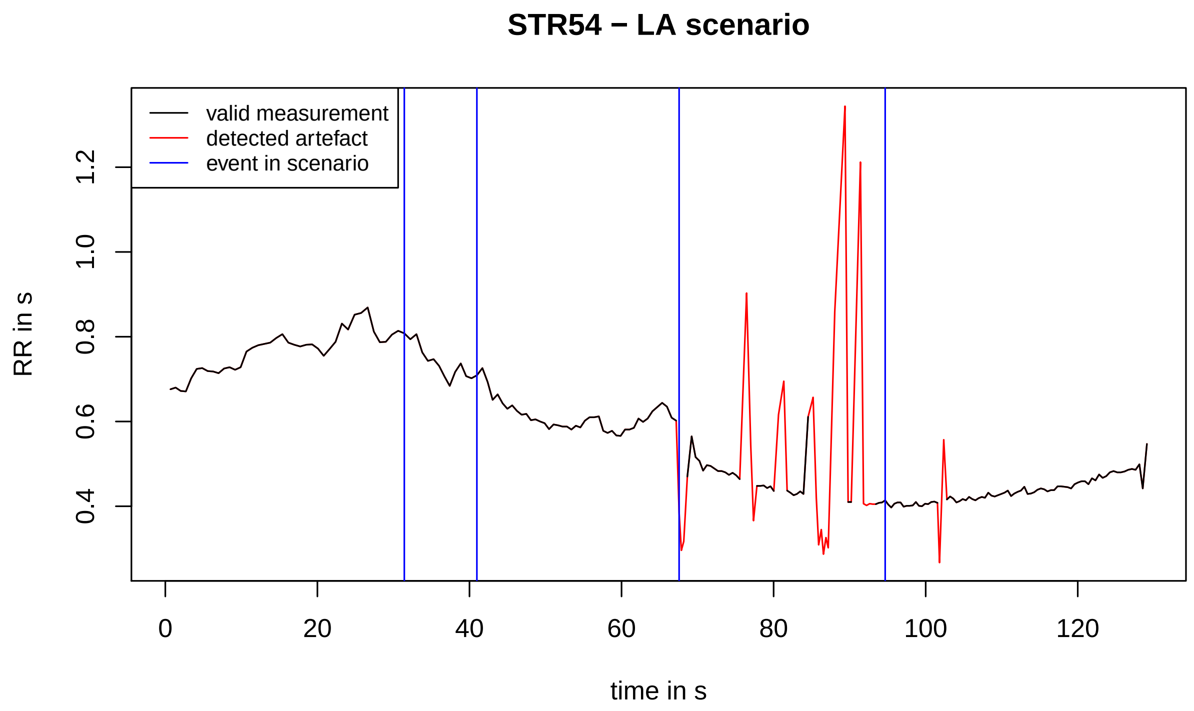
STR54 − LA scenario
valid measurement
detected artefact
event in scenario
RR in s
0.4
0.6
0.8
1.0
1.2
0
20
40
60
80
100
120
time in s

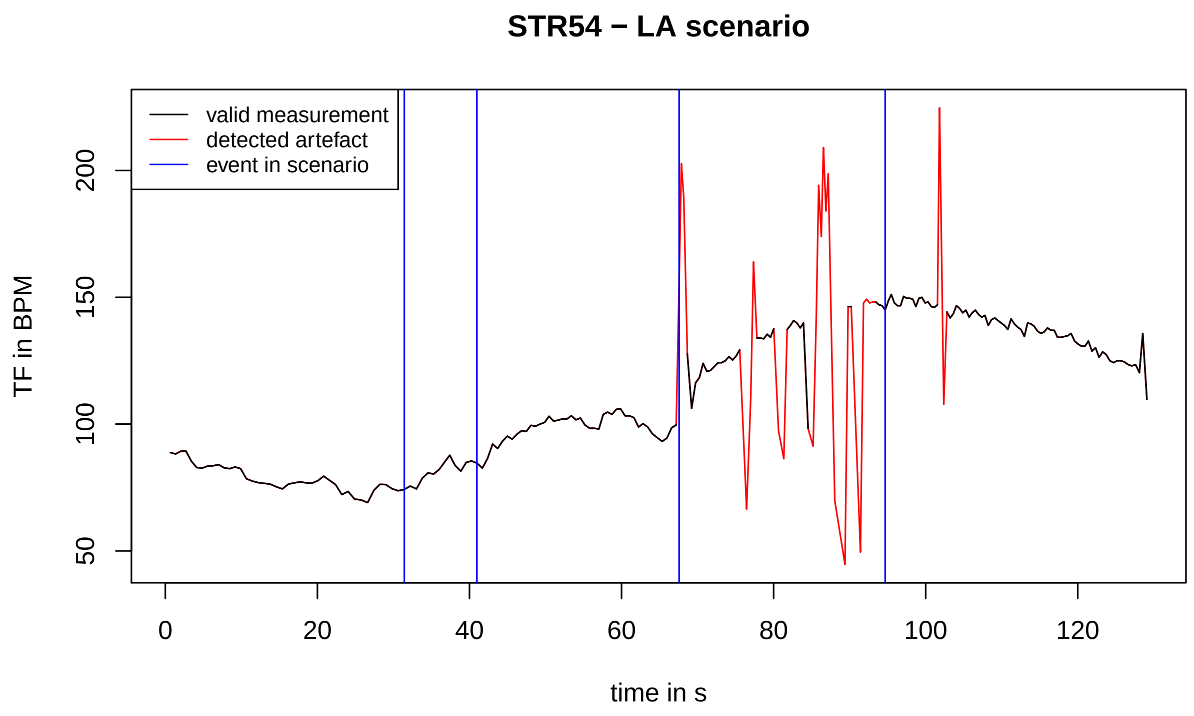
STR54 − LA scenario
valid measurement
detected artefact
event in scenario
TF in BPM
50
100
150
200
0
20
40
60
80
100
120
time in s

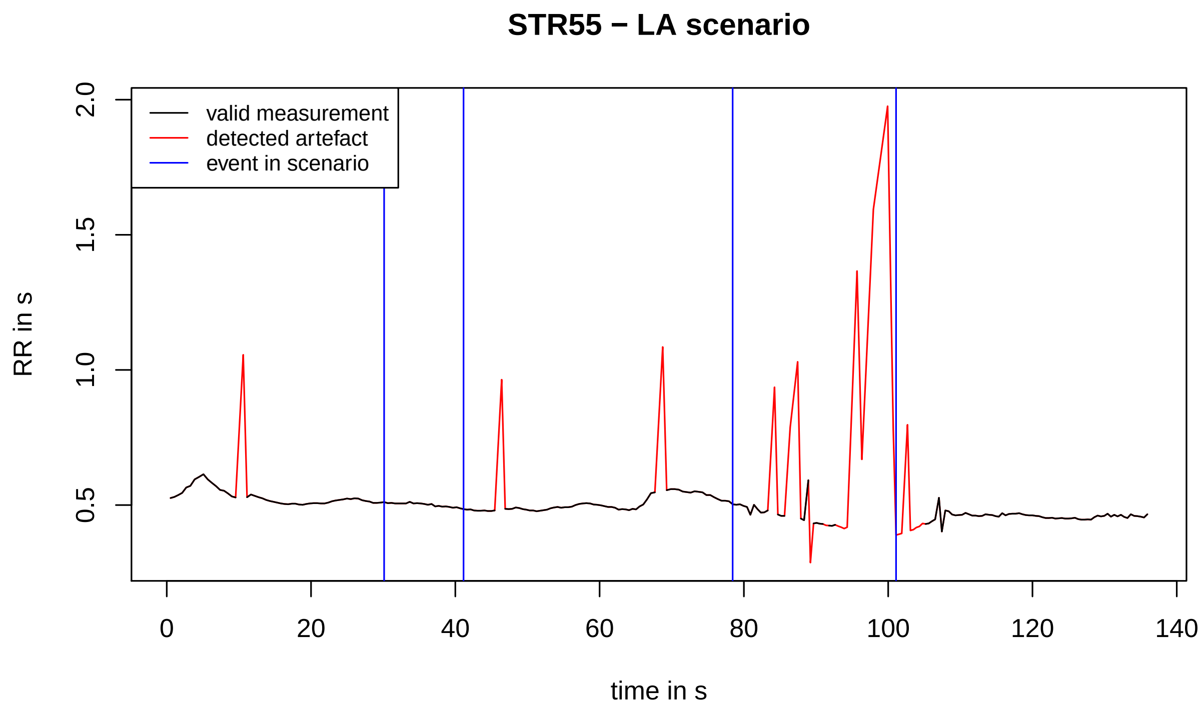
STR55 − LA scenario
valid measurement
detected artefact
event in scenario
RR in s
2.0
1.5
1.0
0.5
0
20
40
60
80
100
120
140
time in s

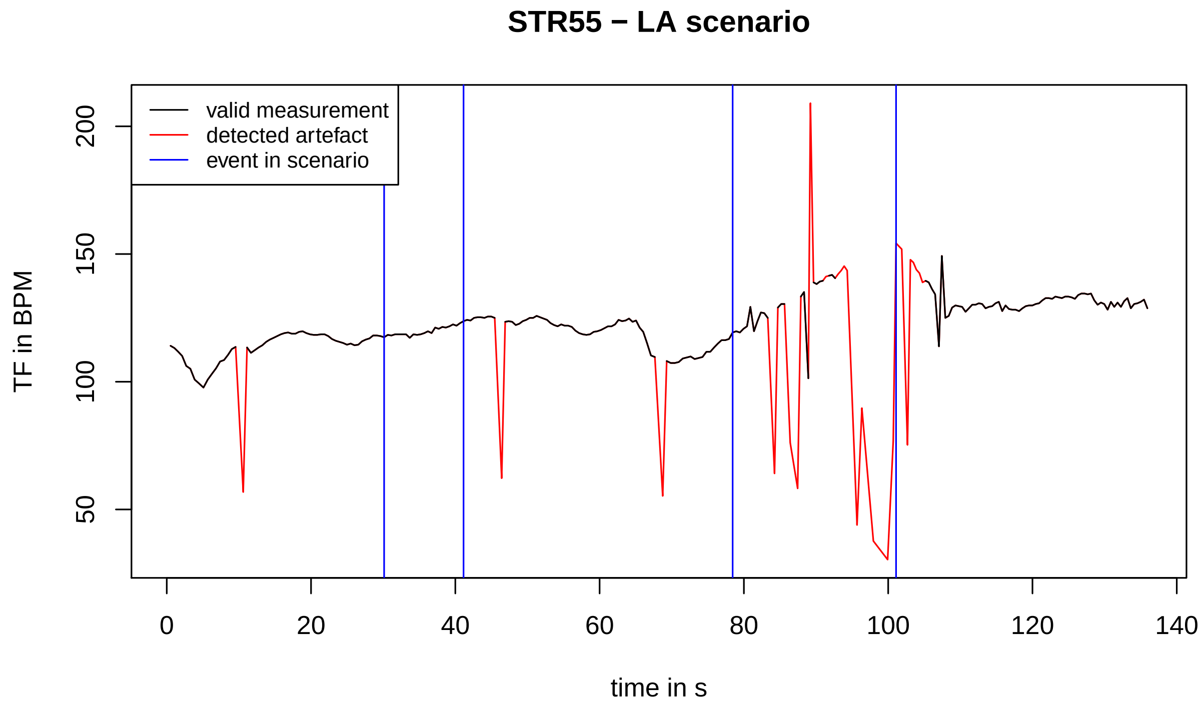
STR55 − LA scenario
valid measurement
detected artefact
event in scenario
TF in BPM
200
150
100
50
0
20
40
60
80
100
120
140
time in s

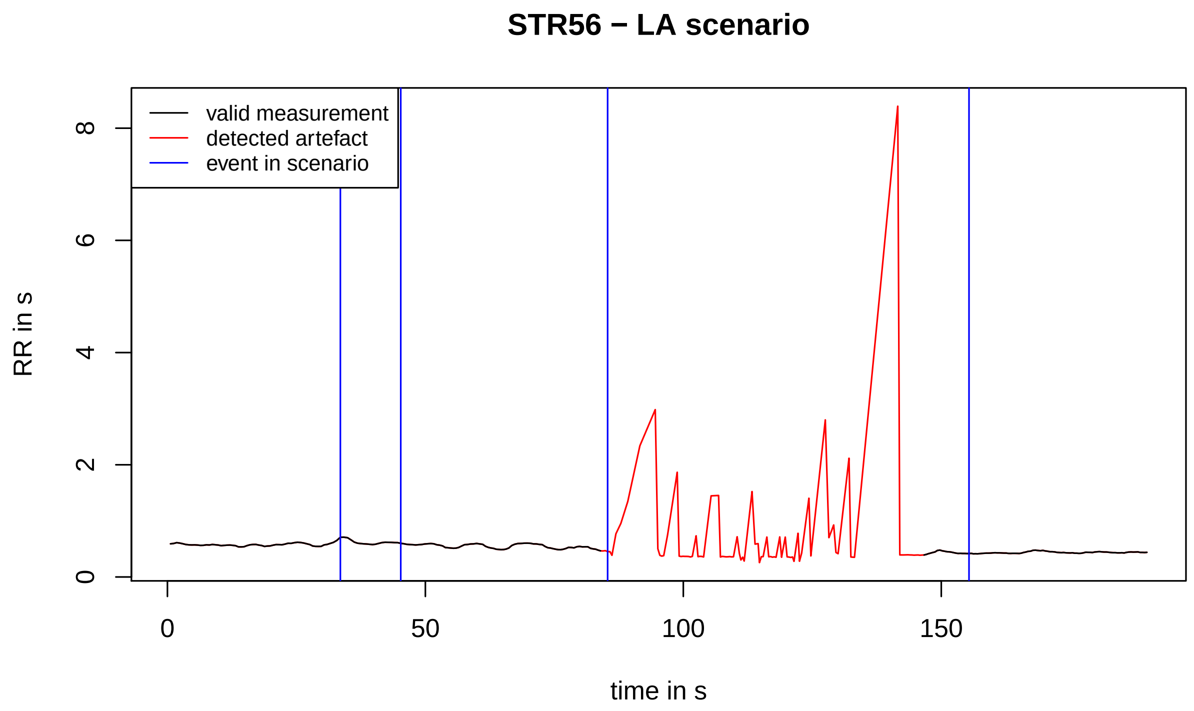
STR56 − LA scenario
valid measurement
detected artefact
event in scenario
RR in s
time in s
0
2
4
6
8
0
50
100
150

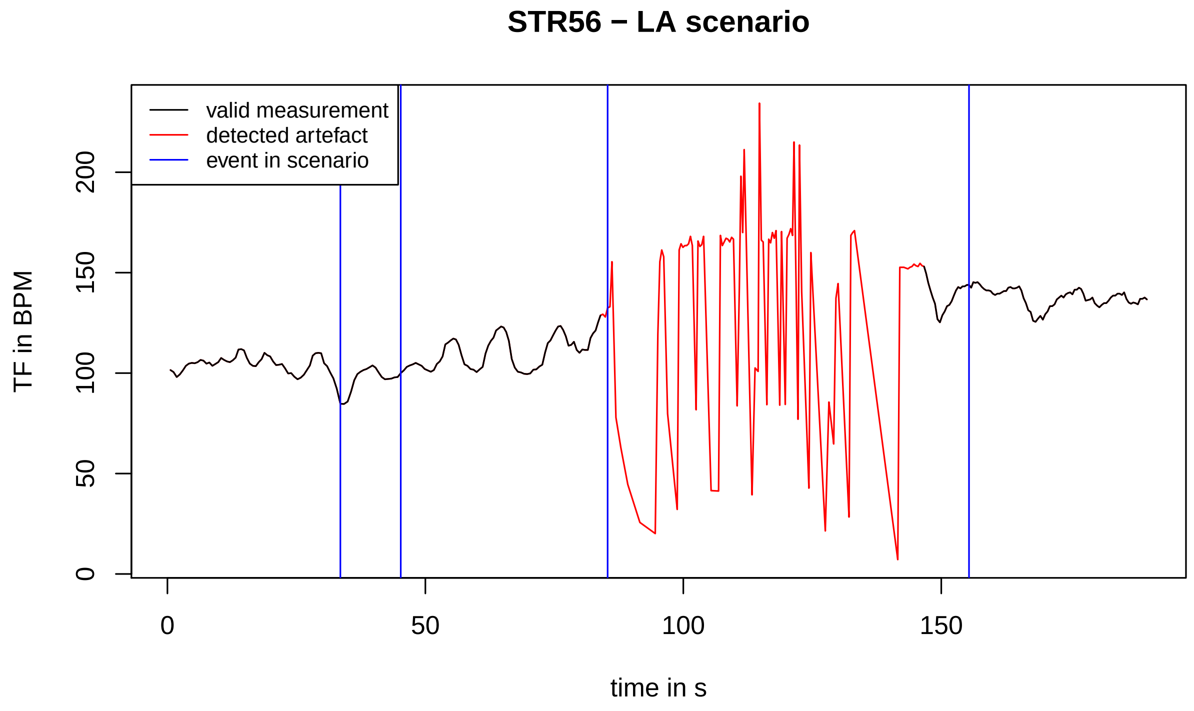
STR56 − LA scenario
valid measurement
detected artefact
event in scenario
TF in BPM
time in s
0
50
100
150
200
0
50
100
150

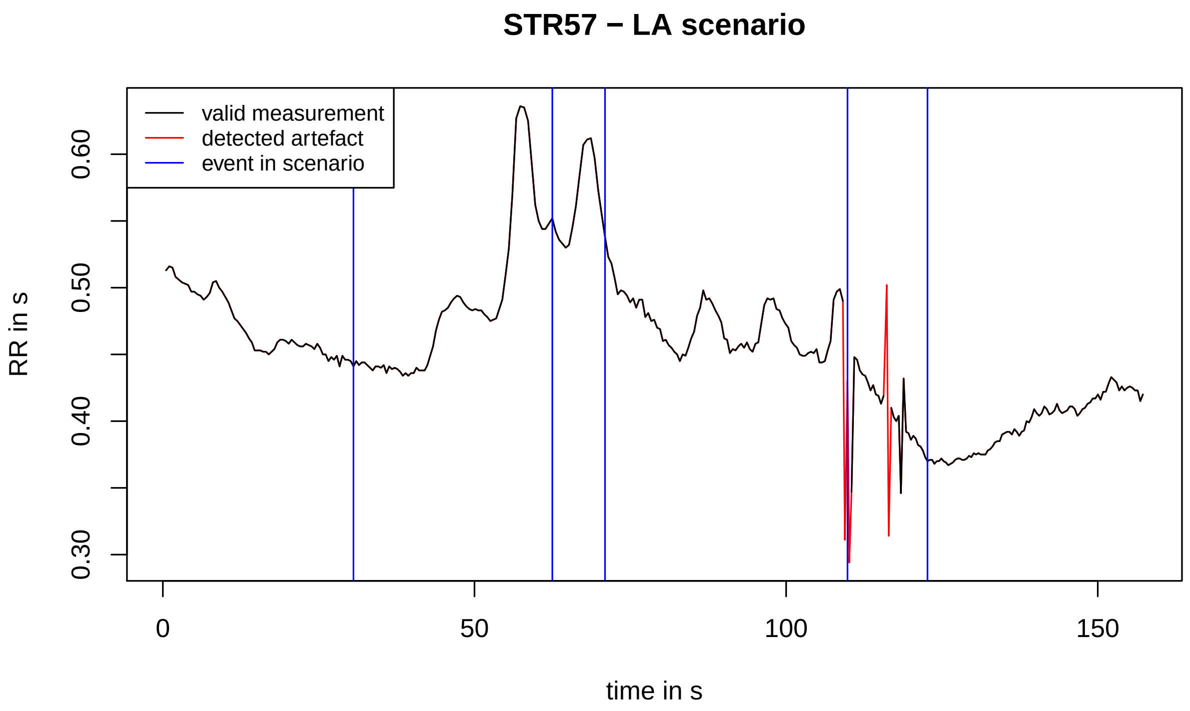
STR57 − LA scenario
valid measurement
detected artefact
event in scenario
RR in s
0.30
0.40
0.50
0.60
0
50
100
150
time in s

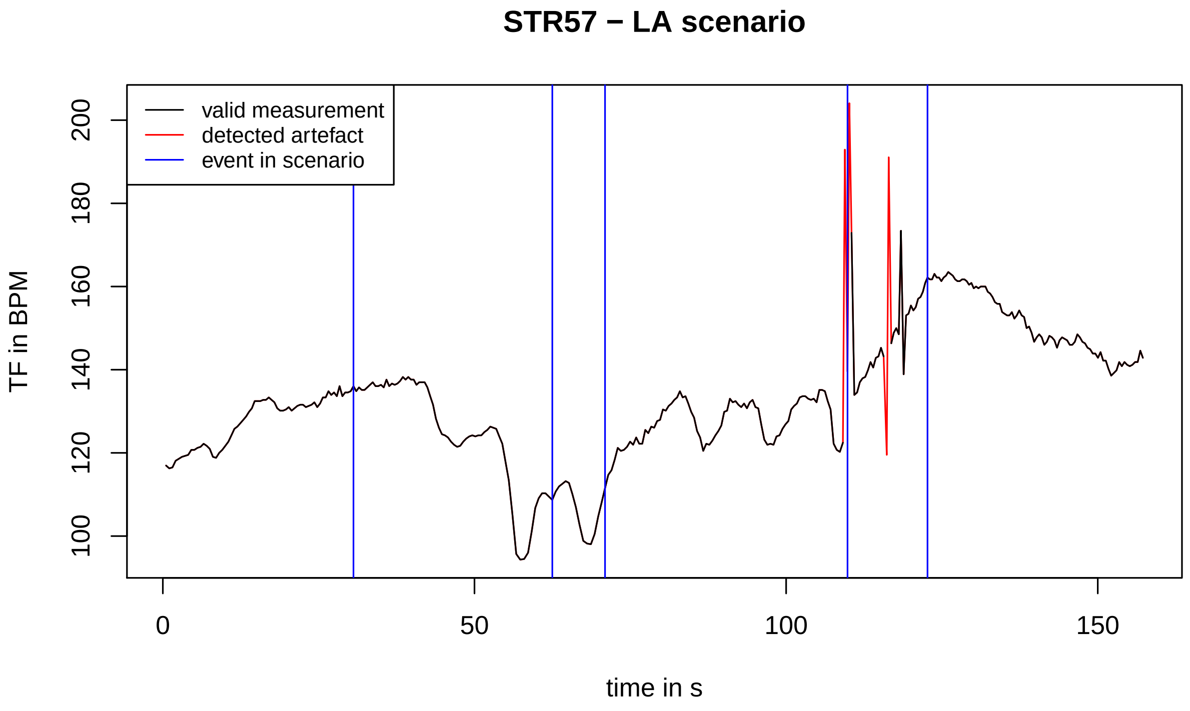
STR57 − LA scenario
valid measurement
detected artefact
event in scenario
TF in BPM
100
120
140
160
180
200
0
50
100
150
time in s

#### STR58 − LA scenario

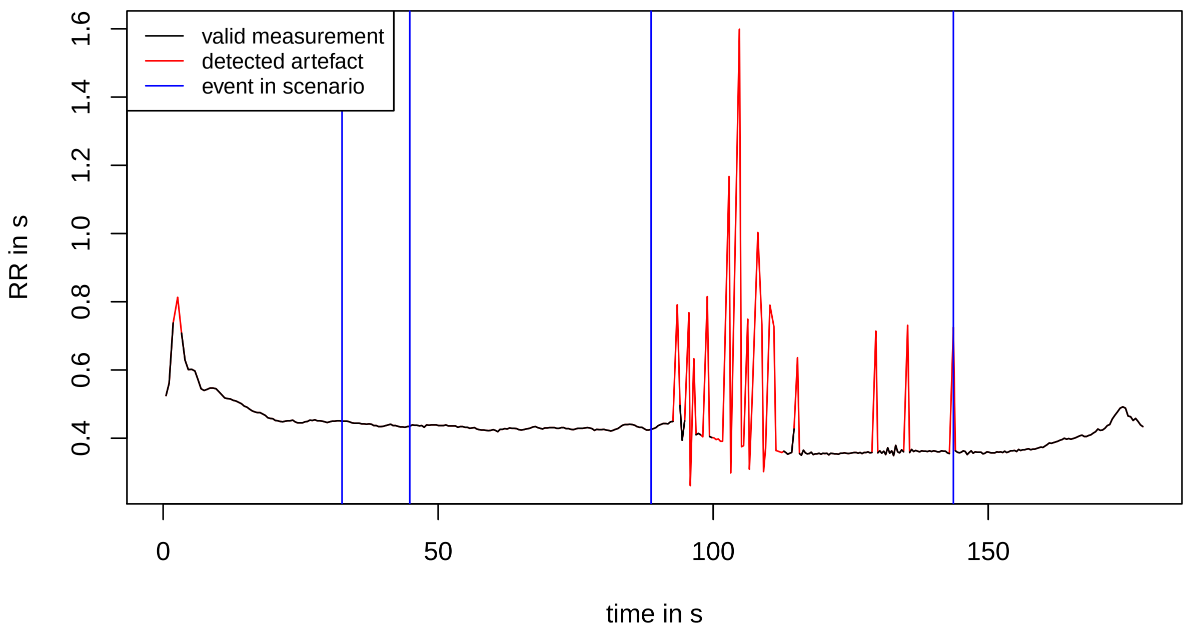


#### STR58 − LA scenario

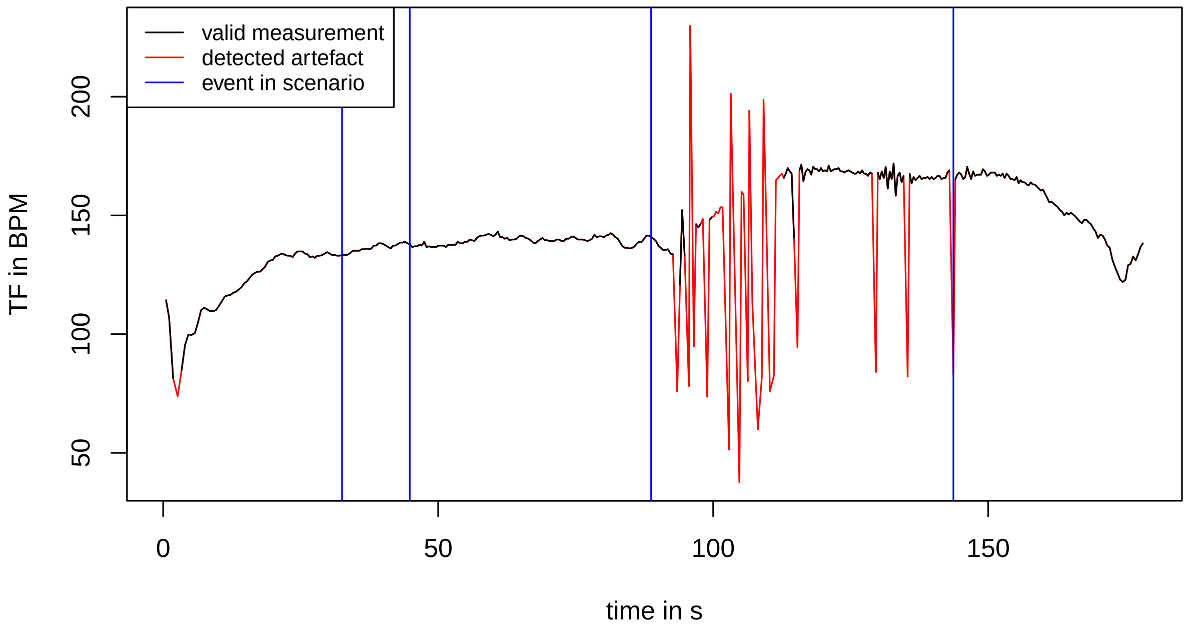

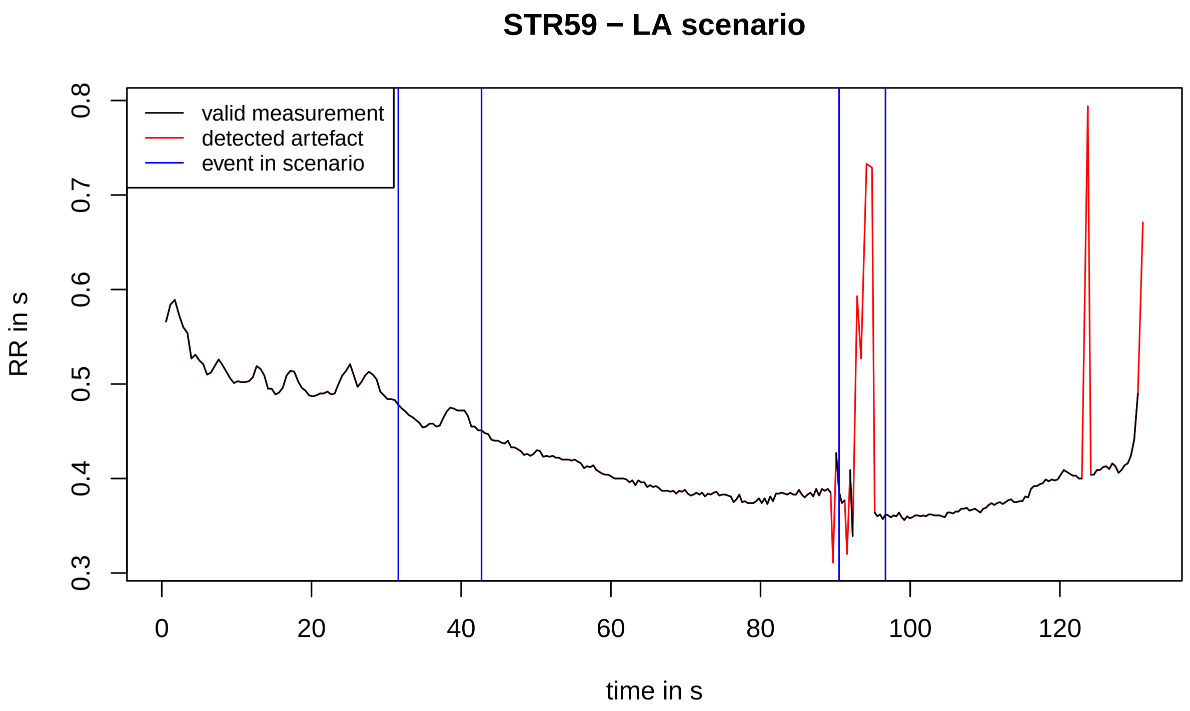
STR59 − LA scenario
valid measurement
detected artefact
event in scenario
RR in s
0.3
0.4
0.5
0.6
0.7
0.8
0
20
40
60
80
100
120
time in s

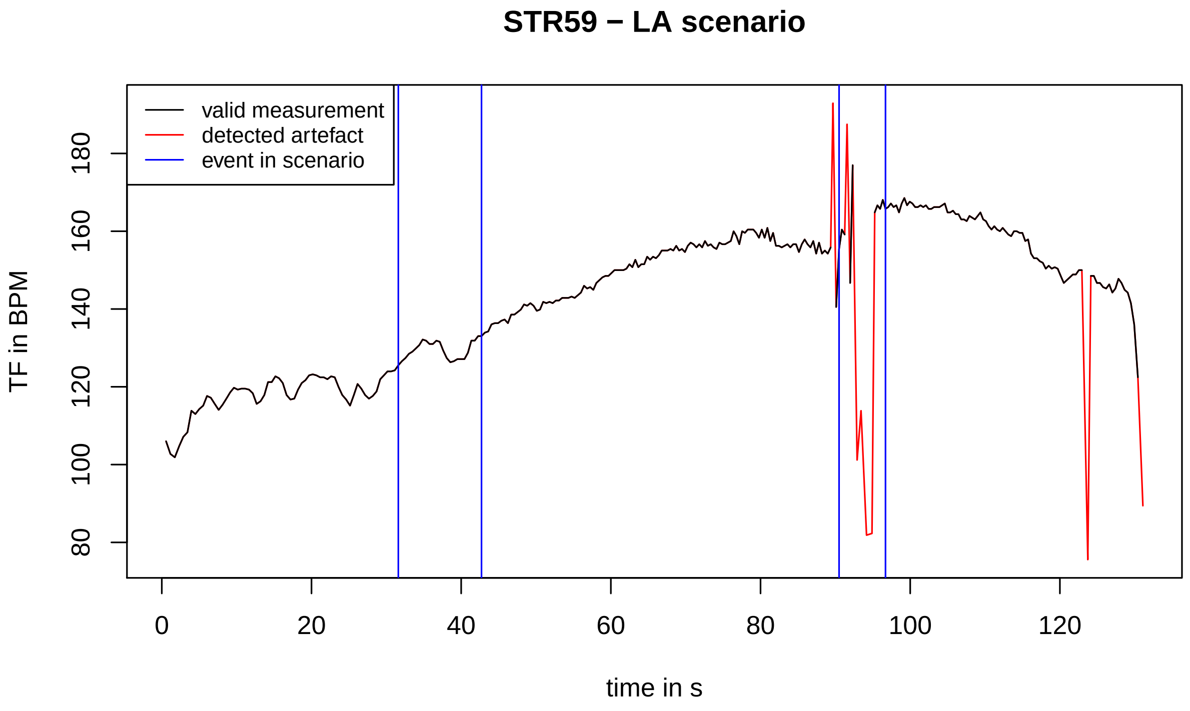
STR59 − LA scenario
valid measurement
detected artefact
event in scenario
TF in BPM
80
100
120
140
160
180
0
20
40
60
80
100
120
time in s

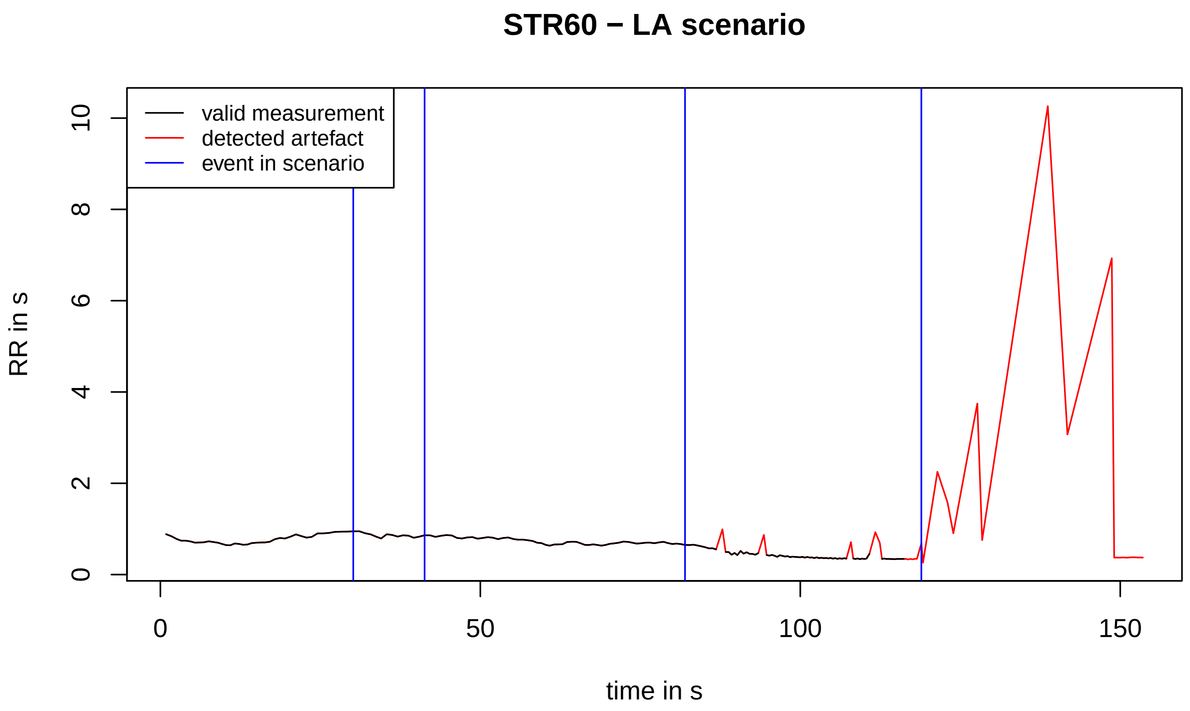
STR60 − LA scenario
valid measurement
detected artefact
event in scenario
RR in s
0
2
4
6
8
10
0
50
100
150
time in s

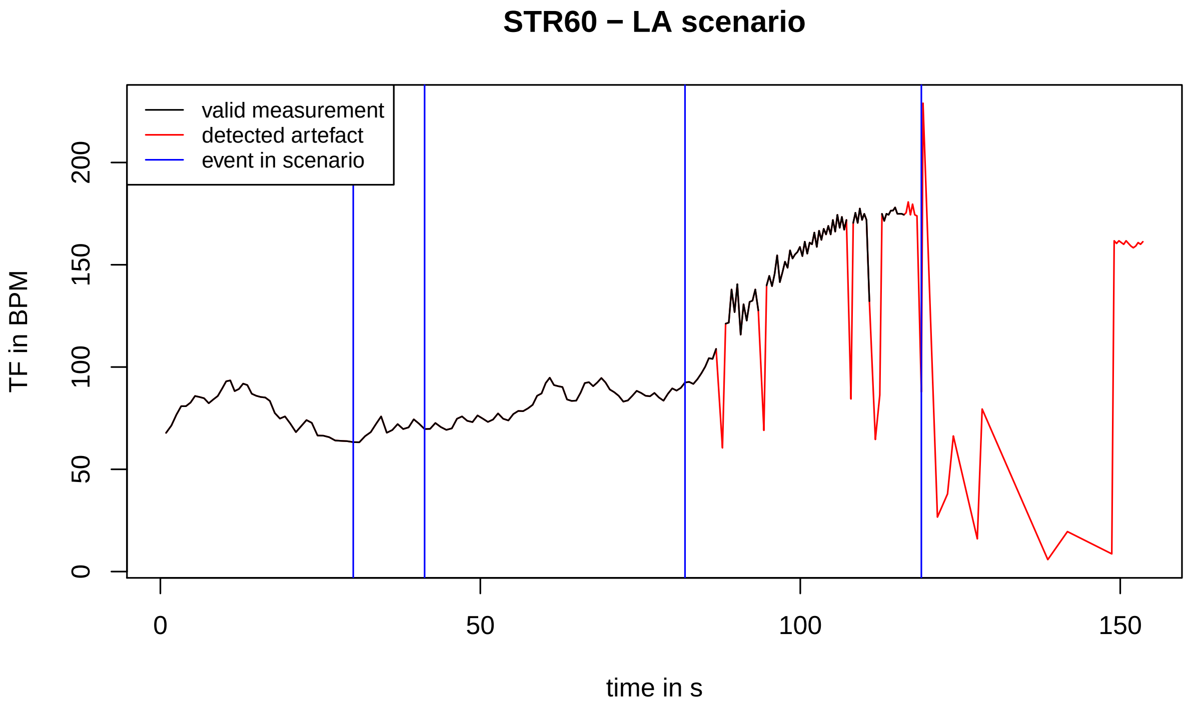
STR60 − LA scenario
valid measurement
detected artefact
event in scenario
TF in BPM
0
50
100
150
200
0
50
100
150
time in s

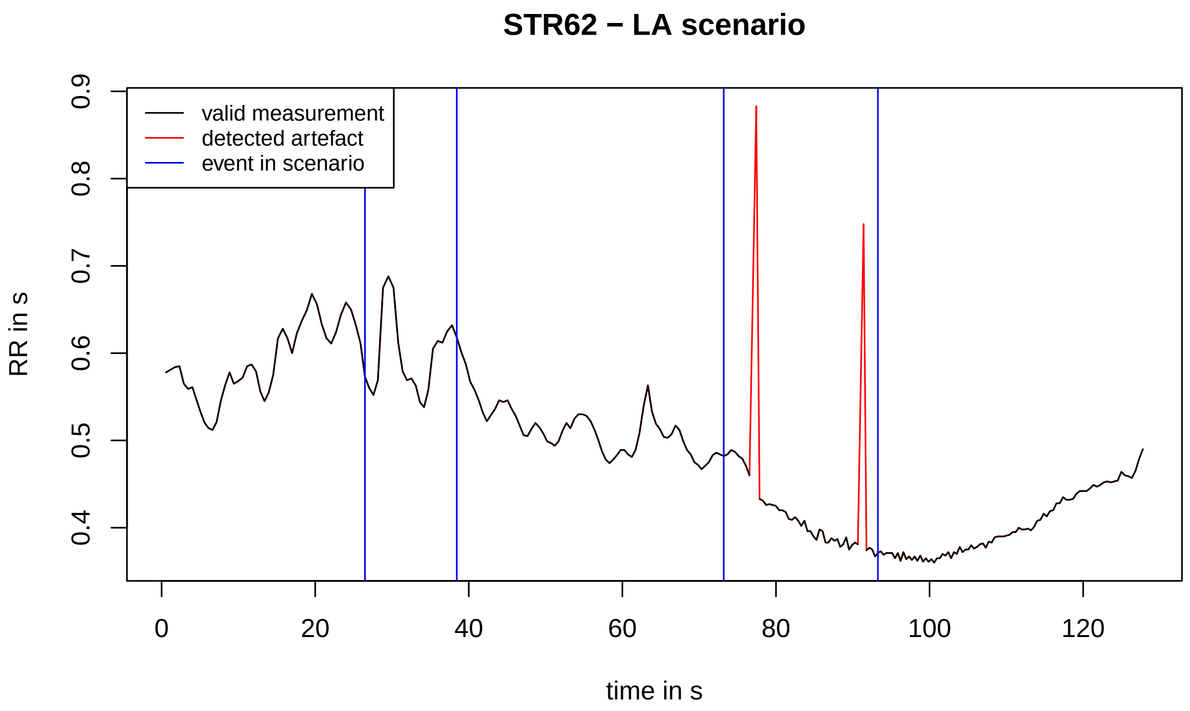
STR62 − LA scenario
valid measurement
detected artefact
event in scenario
RR in s
0.4
0.5
0.6
0.7
0.8
0.9
0
20
40
60
80
100
120
time in s

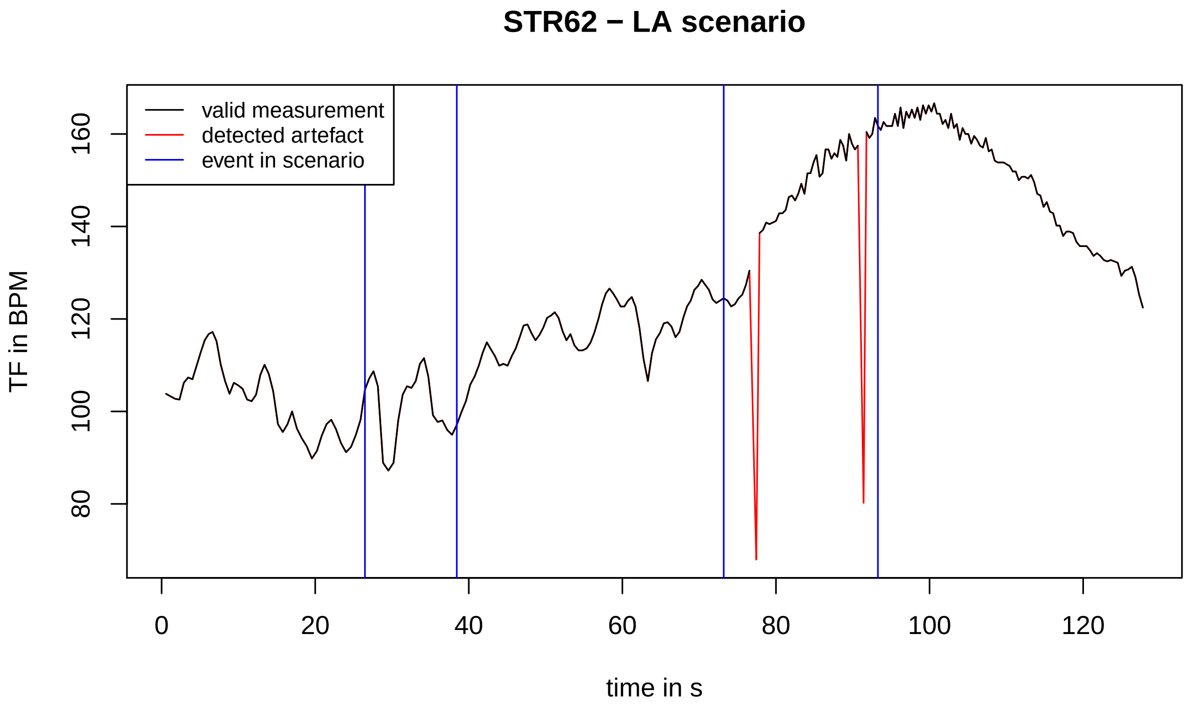
STR62 − LA scenario
valid measurement
detected artefact
event in scenario
TF in BPM
80
100
120
140
160
0
20
40
60
80
100
120
time in s

### Heart rate variability for high anxiety scenario

The file contains plots of time-series of heart rate and RR intervals in the high anxiety scenario. In each plot, the time series line consists of the black and red parts. The black parts denote valid data. The red parts denote automatically detected artefacts. To detect artifact, we used kernel smoothing with bandwidth 10 s and Gaussian kernel. In case the relative difference between the smooth line and raw data was larger than 20%, we called the raw data value an artifact. The vertical blue lines in each plot denotes events in the scenario.

| participant ID | identified problem with the recording |
|---|---|
| STR01 | - |
| STR02 | RR intervals recording failed, recording missing |
| STR03 | RR intervals recording failed after approximately 1.5 minutes |
| STR04 | - |
| STR05 | - |
| STR06 | participant ID was not assigned |
| STR07 | - |
| STR08 | RR intervals recording failed after approximately 2 minutes |
| STR09 | - |
| STR10 | - |
| STR11 | - |
| STR12 | - |
| STR13 | - |
| STR14 | participant ID was not assigned |
| STR15 | - |
| STR16 | - |
| STR17 | - |
| STR18 | - |
| STR19 | - |
| STR20 | - |
| STR21 | failed to assign the second event in scenario |
| STR22 | failed to assign events in scenario (except the last one) |
| STR23 | failed to assign events in scenario |
| STR24 | - |
| STR25 | - |
| STR26 | RR intervals recording failed after approximately 2 minutes |
| STR27 | failed to assign last two events in the scenario |
| STR28 | - |
| STR29 | - |
| STR30 | - |
| STR31 | - |
| STR32 | - |
| STR33 | - |
| STR34 | - |
| STR35 | failed to assign last two events in scenario |
| STR36 | - |
| STR37 | RR intervals recording failed after approximately 1.5 minutes |

| | |
|---|---|
| STR38 | - |
| STR39 | - |
| STR40 | - |
| STR41 | - |
| STR42 | - |
| STR43 | failed to assign the second event in scenario |
| STR44 | - |
| STR45 | - |
| STR46 | - |
| STR47 | - |
| STR48 | - |
| STR49 | - |
| STR50 | - |
| STR51 | - |
| STR52 | - |
| STR53 | - |
| STR54 | - |
| STR55 | - |
| STR56 | - |
| STR57 | - |
| STR58 | - |
| STR59 | - |
| STR60 | - |
| STR61 | failed to assign last two events in the scenario |
| STR62 | - |

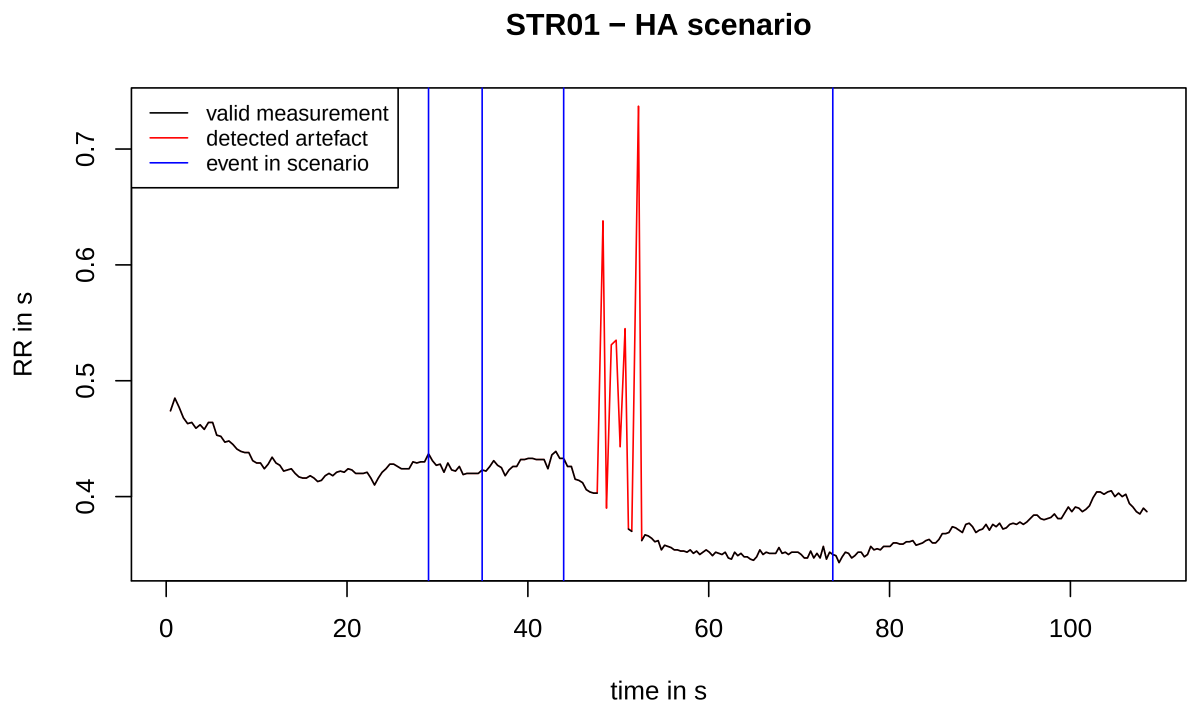
STR01 − HA scenario
valid measurement
detected artefact
event in scenario
RR in s
0.4
0.5
0.6
0.7
0
20
40
60
80
100
time in s

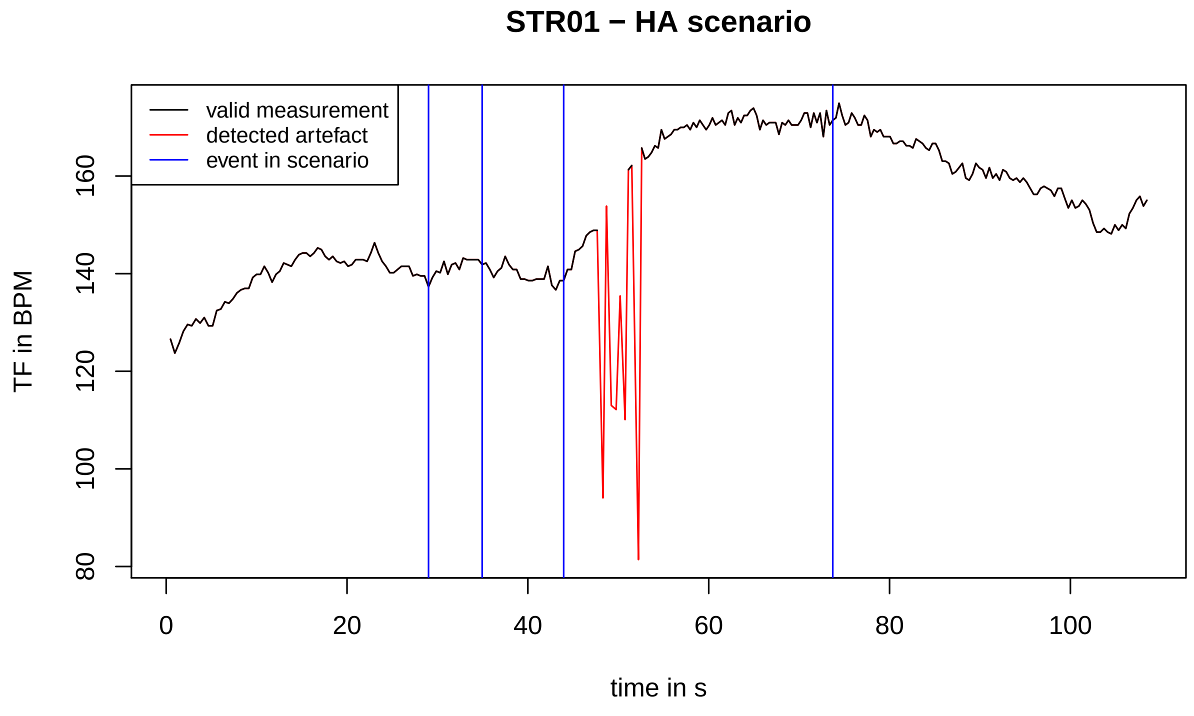
STR01 − HA scenario
valid measurement
detected artefact
event in scenario
TF in BPM
80
100
120
140
160
0
20
40
60
80
100
time in s

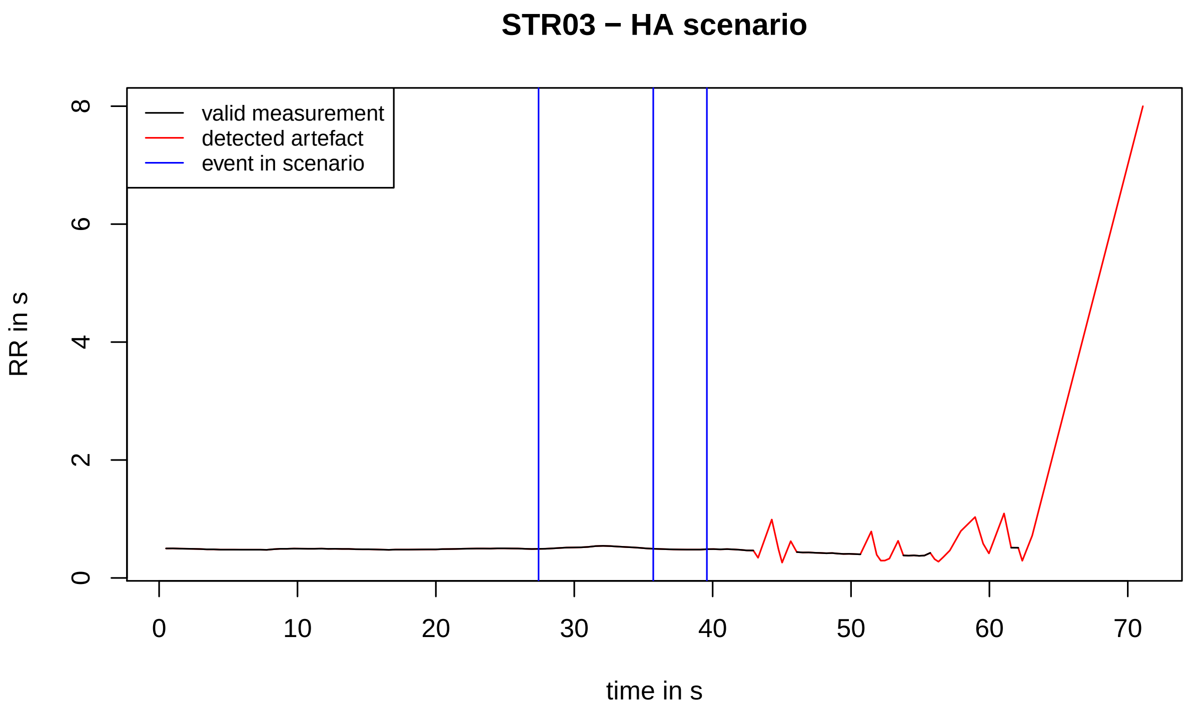
STR03 − HA scenario
valid measurement
detected artefact
event in scenario
RR in s
0
2
4
6
8
0
10
20
30
40
50
60
70
time in s

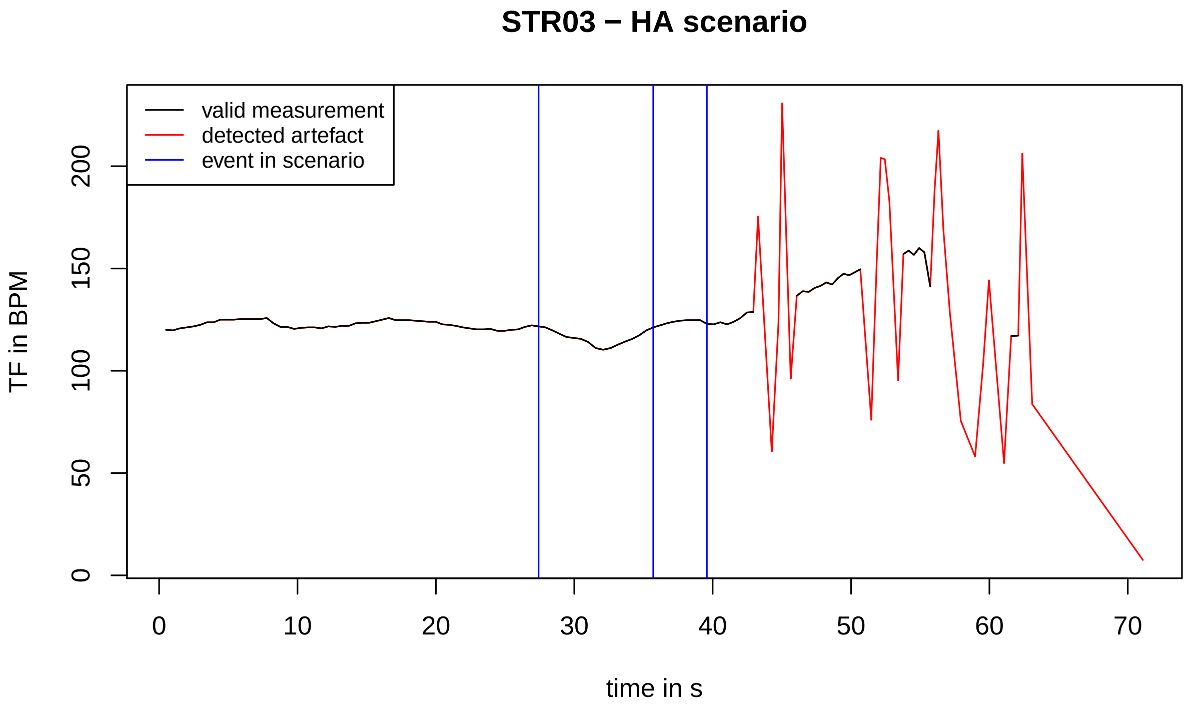
STR03 − HA scenario
valid measurement
detected artefact
event in scenario
TF in BPM
0
50
100
150
200
0
10
20
30
40
50
60
70
time in s

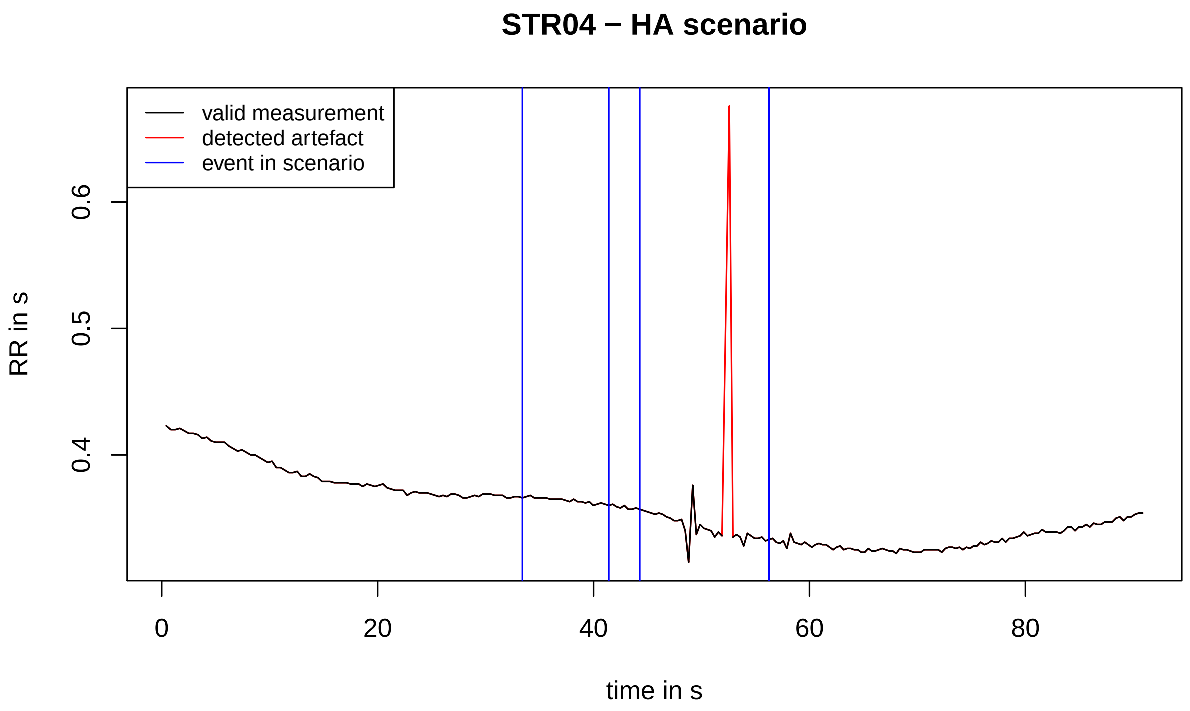
STR04 − HA scenario
valid measurement
detected artefact
event in scenario
RR in s
0.4
0.5
0.6
0
20
40
60
80
time in s

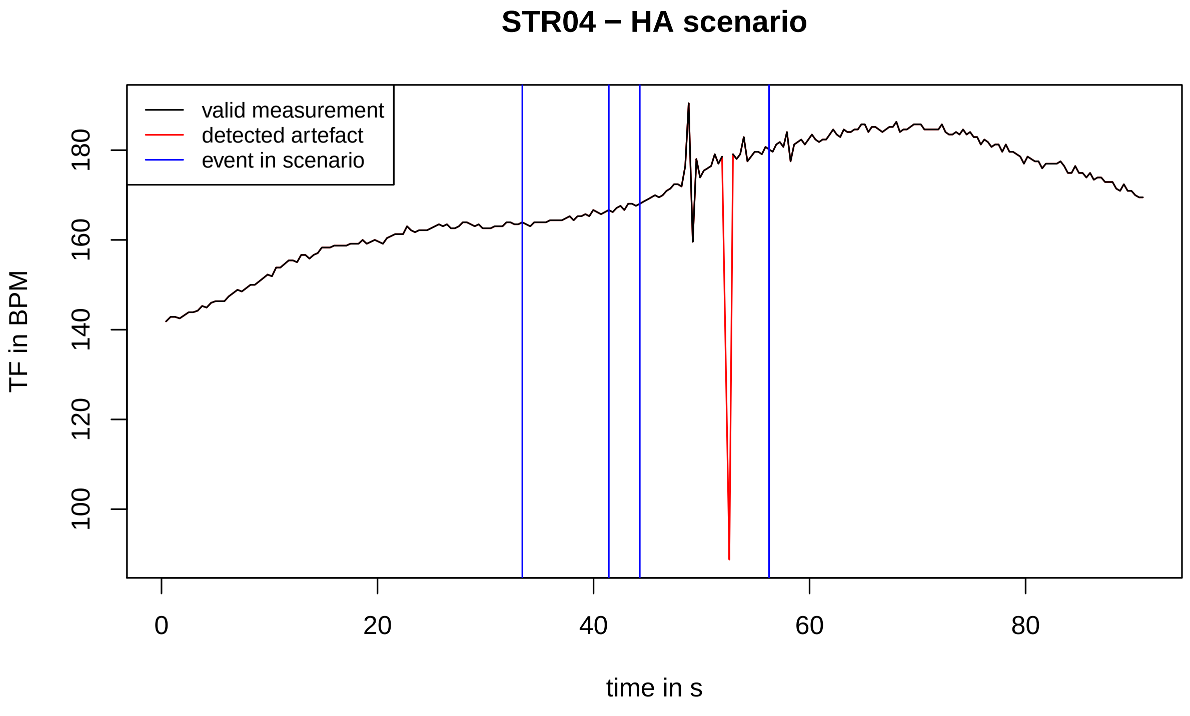
STR04 − HA scenario
valid measurement
detected artefact
event in scenario
TF in BPM
100
120
140
160
180
0
20
40
60
80
time in s

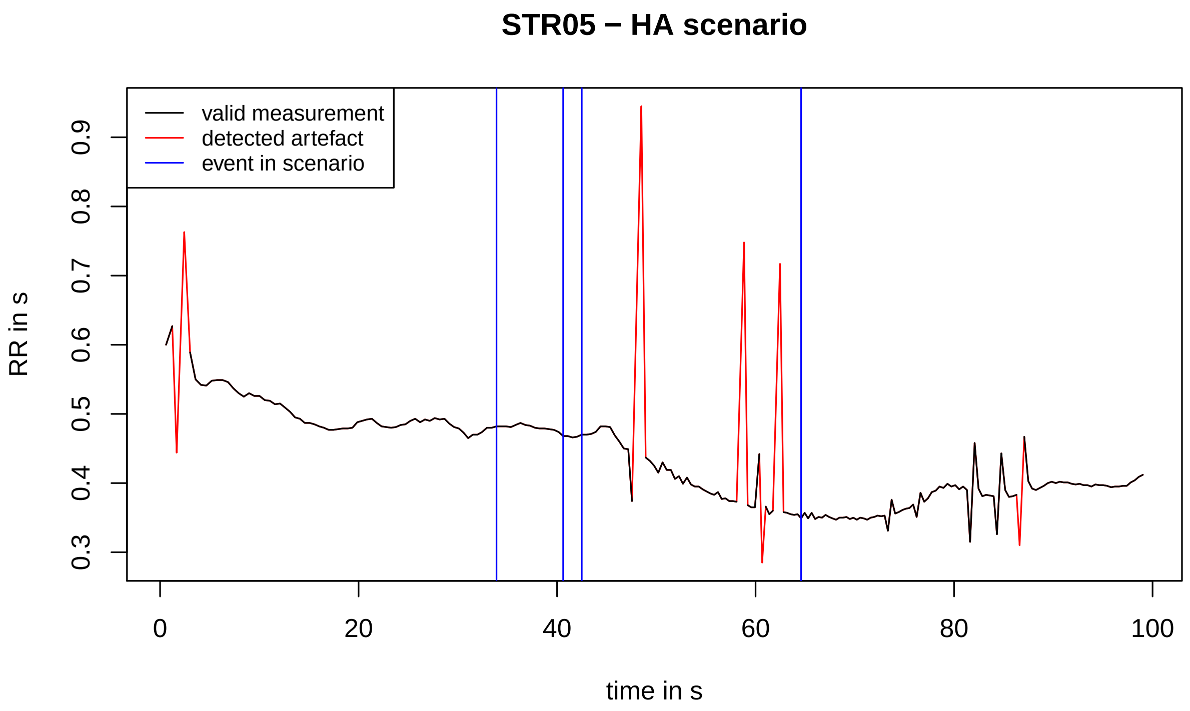
STR05 − HA scenario
valid measurement
detected artefact
event in scenario
RR in s
0.3
0.4
0.5
0.6
0.7
0.8
0.9
time in s
0
20
40
60
80
100

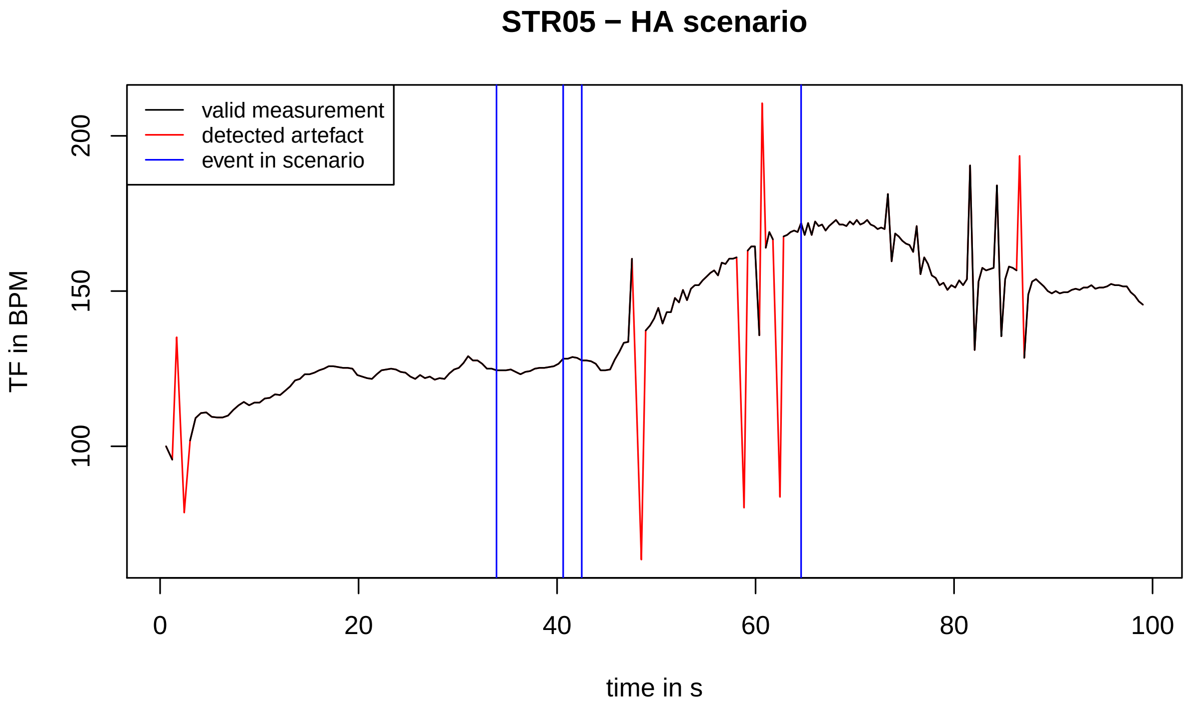
STR05 − HA scenario
valid measurement
detected artefact
event in scenario
TF in BPM
100
150
200
time in s
0
20
40
60
80
100

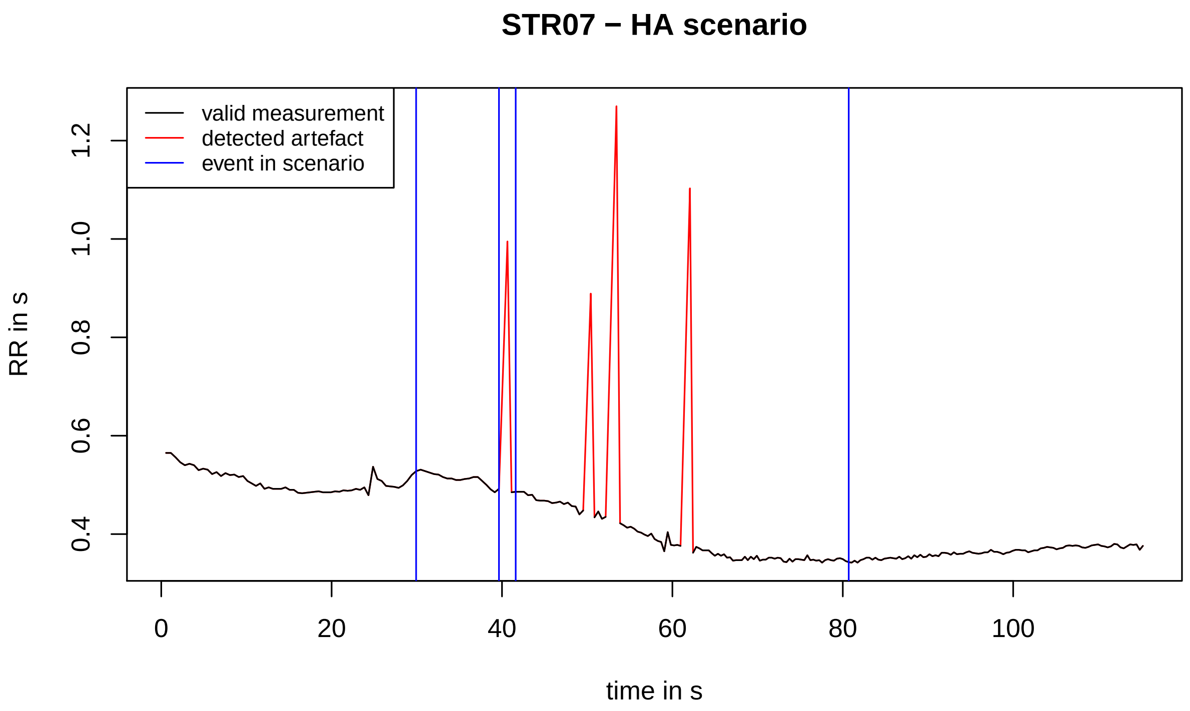
STR07 − HA scenario
valid measurement
detected artefact
event in scenario
RR in s
0.4
0.6
0.8
1.0
1.2
0
20
40
60
80
100
time in s

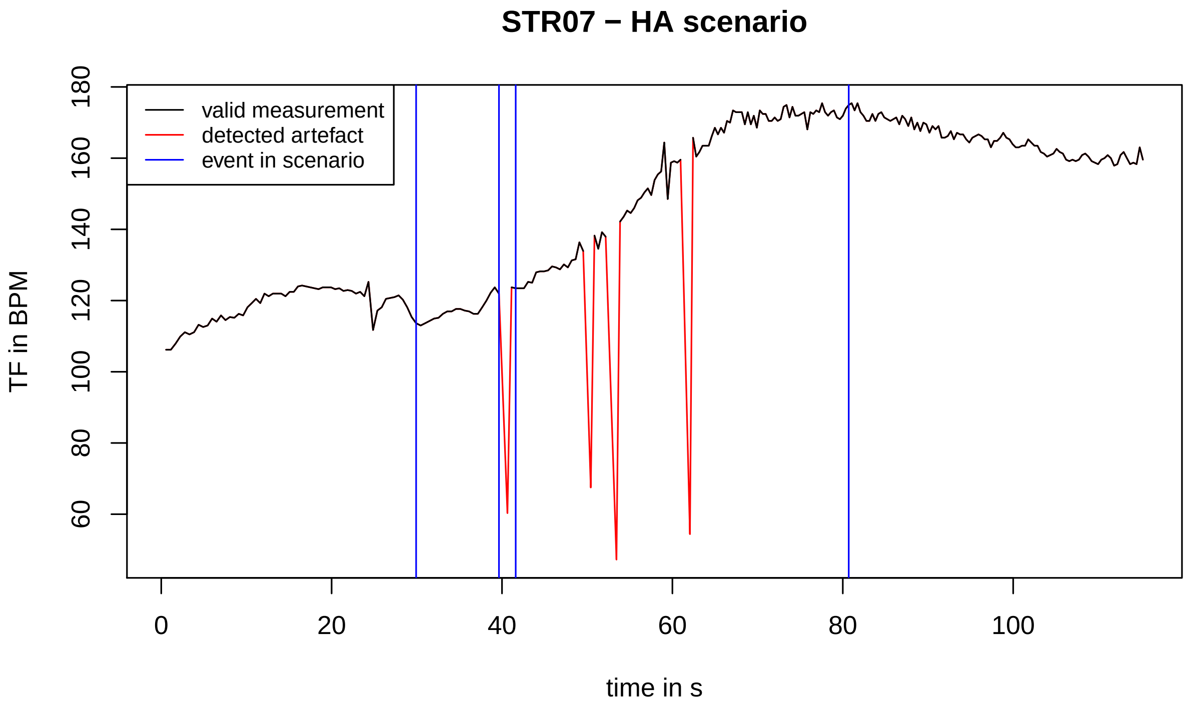
STR07 − HA scenario
valid measurement
detected artefact
event in scenario
TF in BPM
60
80
100
120
140
160
180
0
20
40
60
80
100
time in s

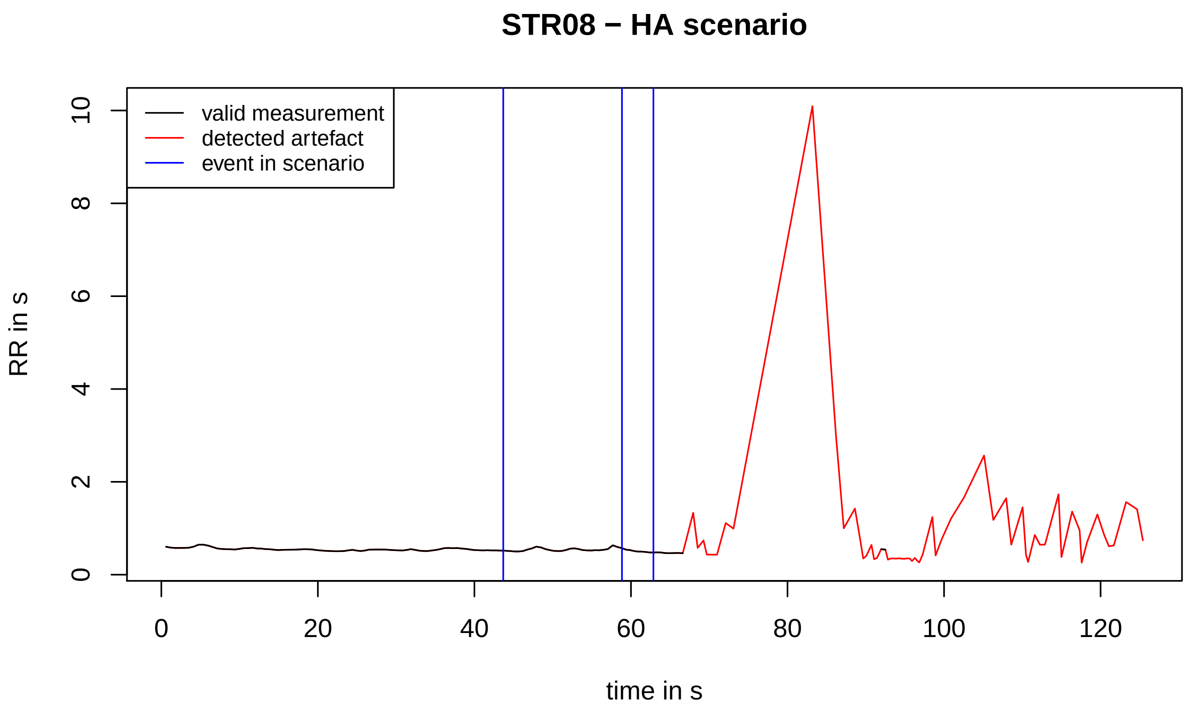
STR08 − HA scenario
valid measurement
detected artefact
event in scenario
RR in s
0
2
4
6
8
10
0
20
40
60
80
100
120
time in s

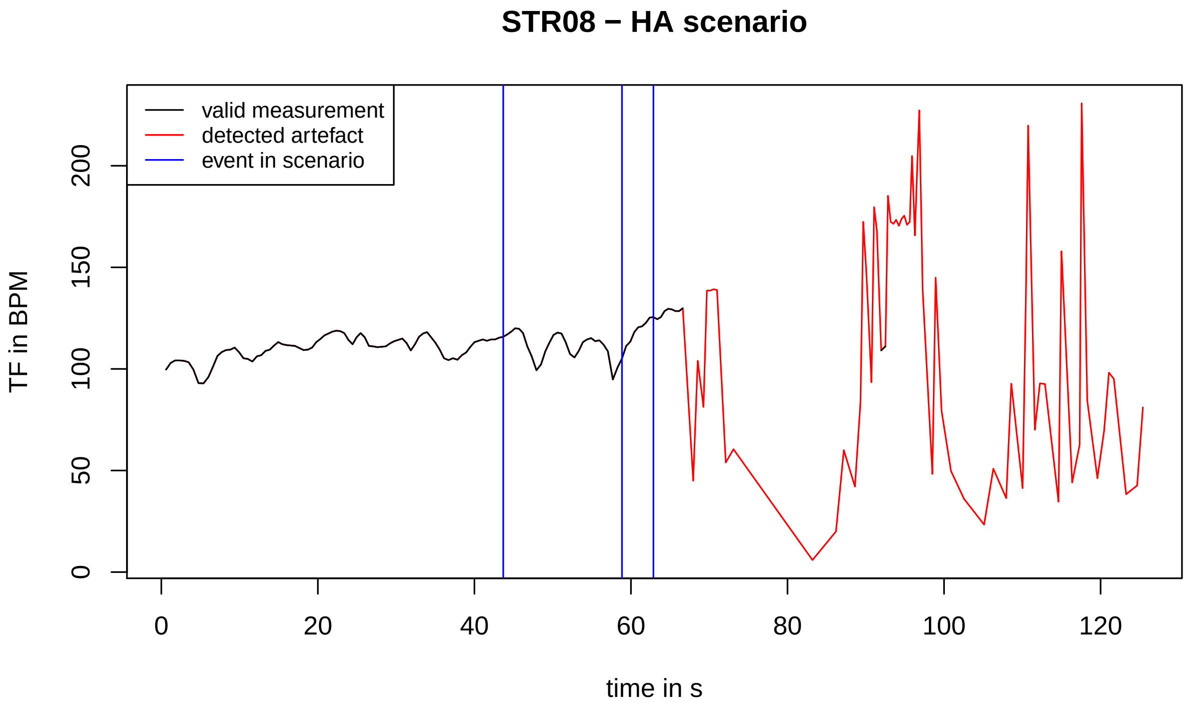
STR08 − HA scenario
valid measurement
detected artefact
event in scenario
TF in BPM
0
50
100
150
200
0
20
40
60
80
100
120
time in s

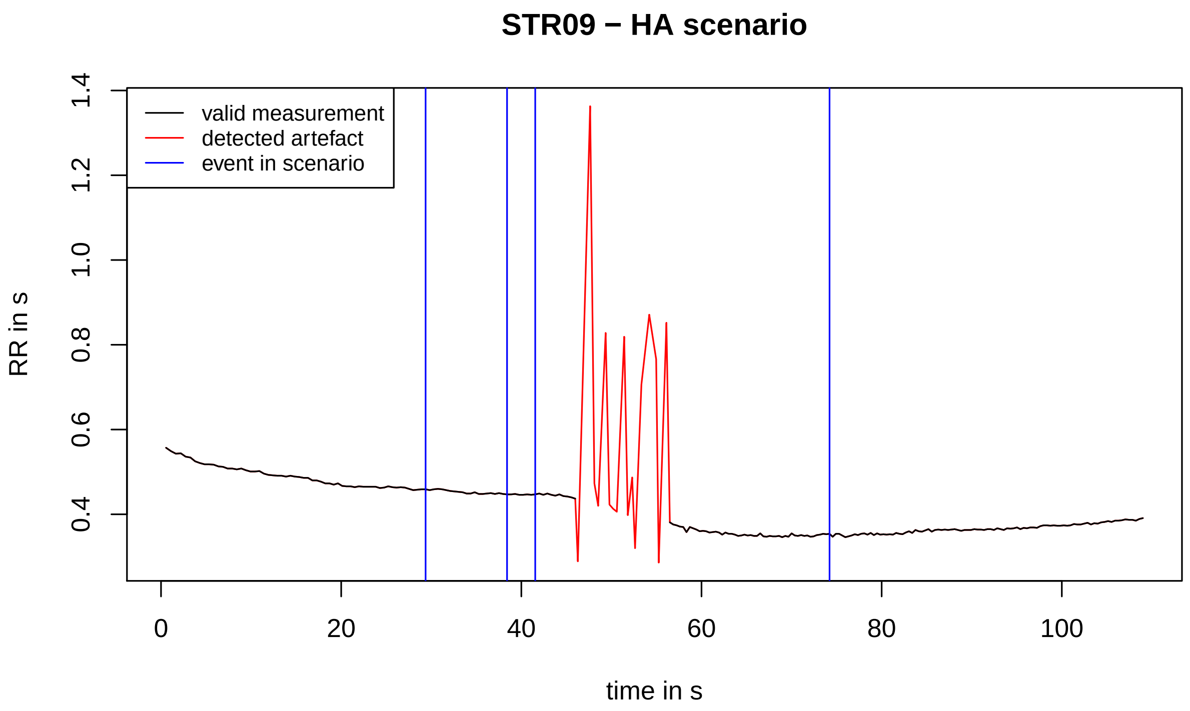
STR09 − HA scenario
valid measurement
detected artefact
event in scenario
RR in s
time in s
0.4
0.6
0.8
1.0
1.2
1.4
0
20
40
60
80
100

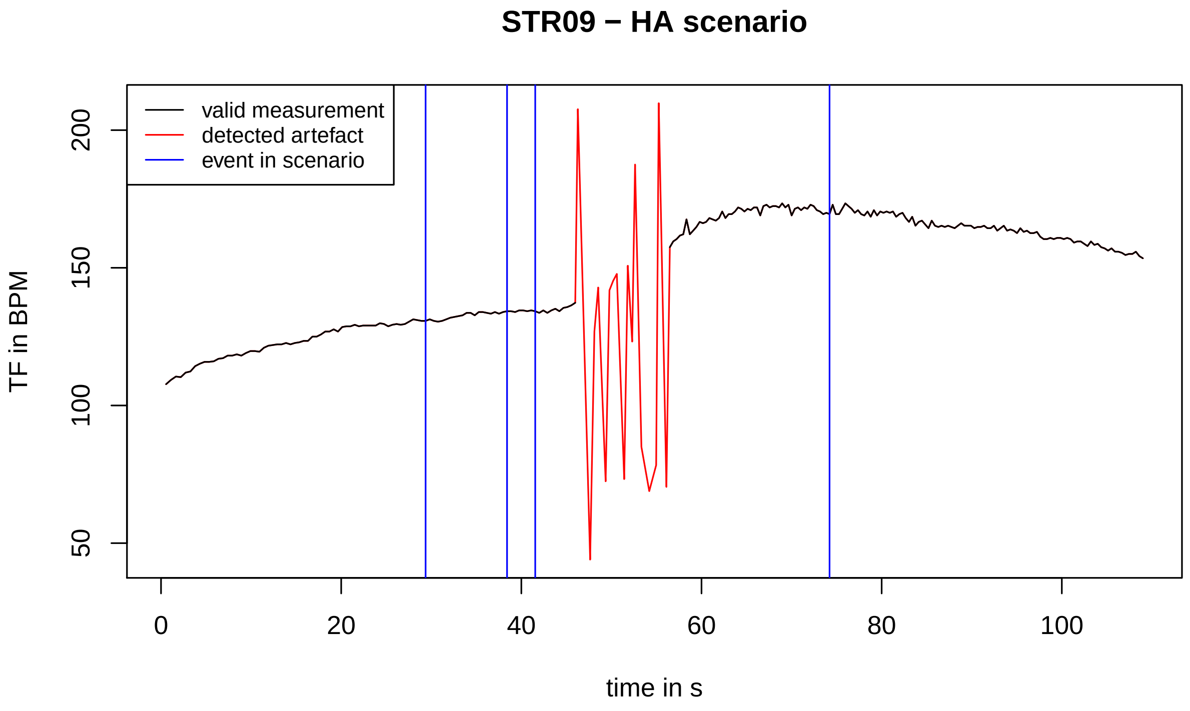
STR09 − HA scenario
valid measurement
detected artefact
event in scenario
TF in BPM
time in s
50
100
150
200
0
20
40
60
80
100

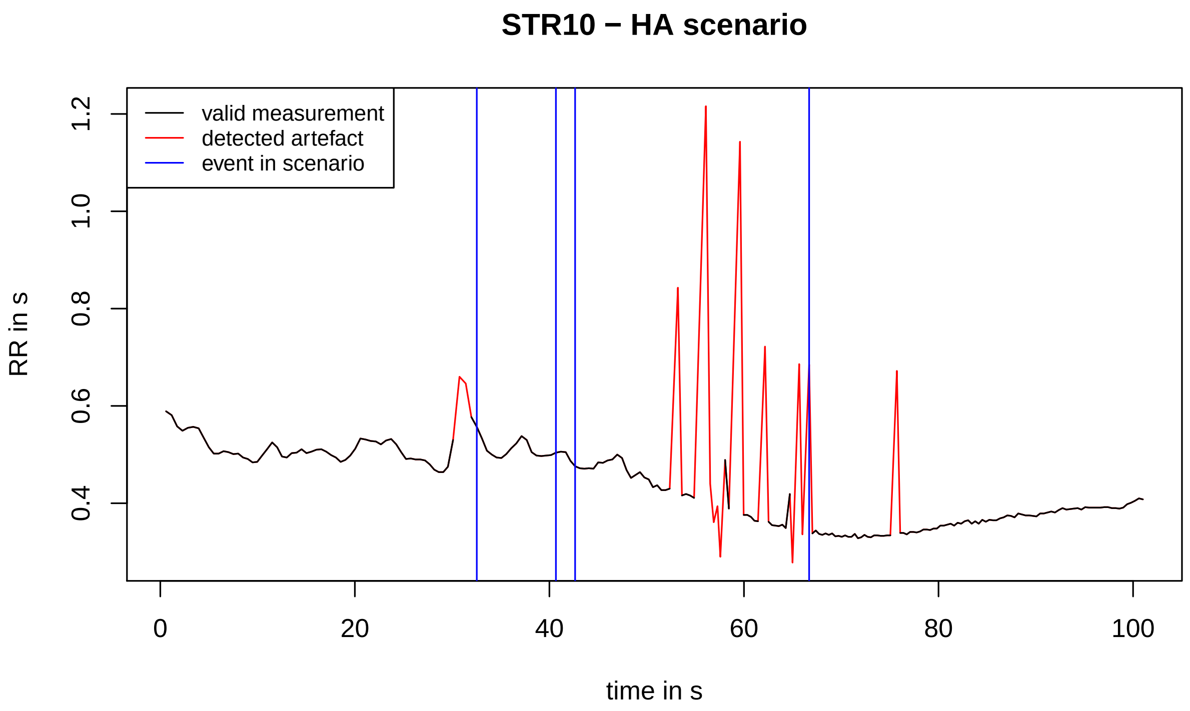
STR10 − HA scenario
valid measurement
detected artefact
event in scenario
RR in s
0.4
0.6
0.8
1.0
1.2
0
20
40
60
80
100
time in s

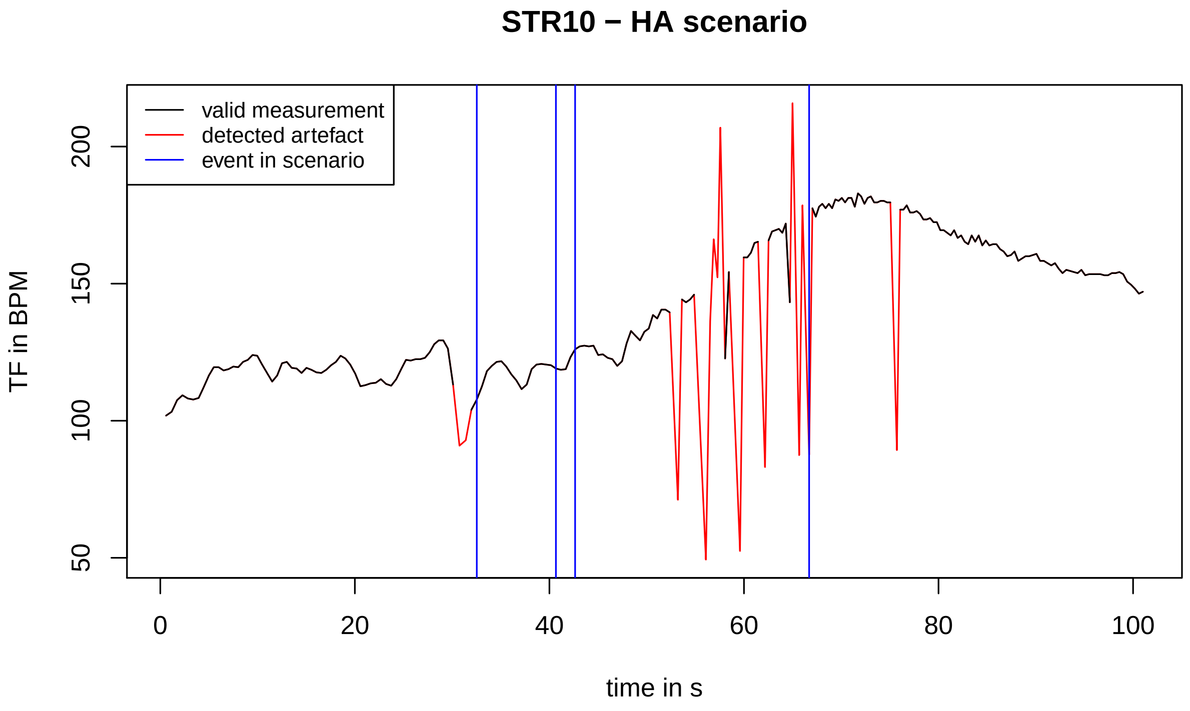
STR10 − HA scenario
valid measurement
detected artefact
event in scenario
TF in BPM
50
100
150
200
0
20
40
60
80
100
time in s

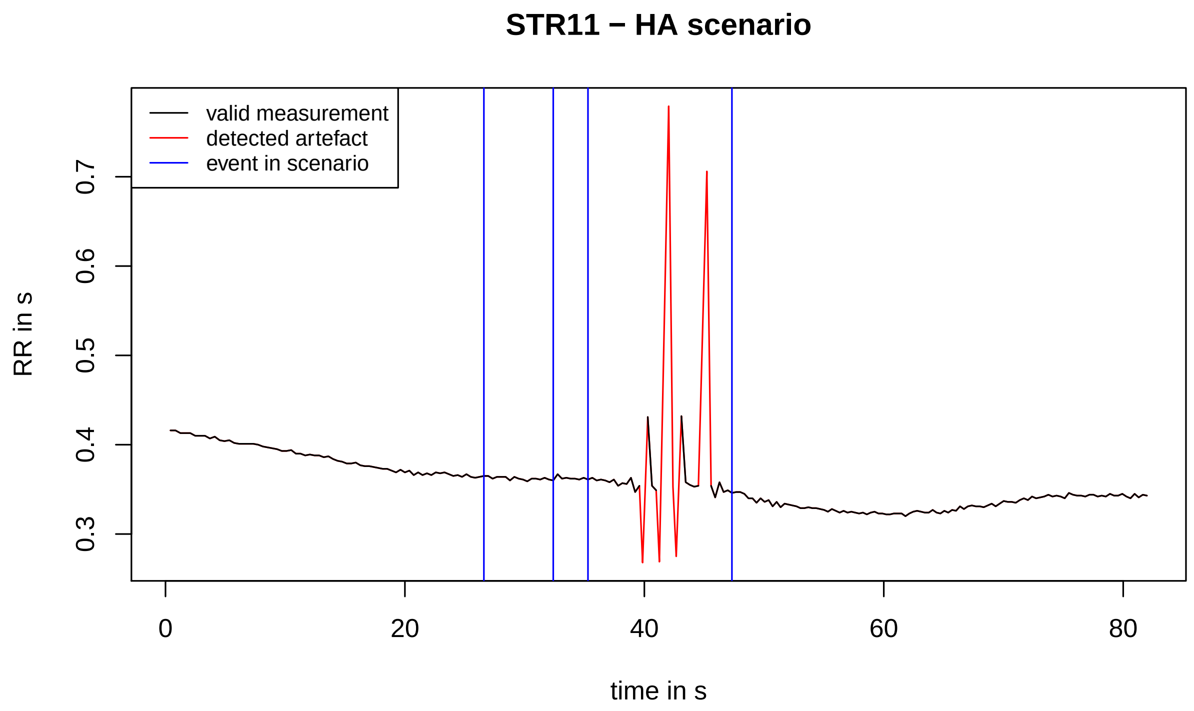
STR11 − HA scenario
valid measurement
detected artefact
event in scenario
RR in s
0.3
0.4
0.5
0.6
0.7
0
20
40
60
80
time in s

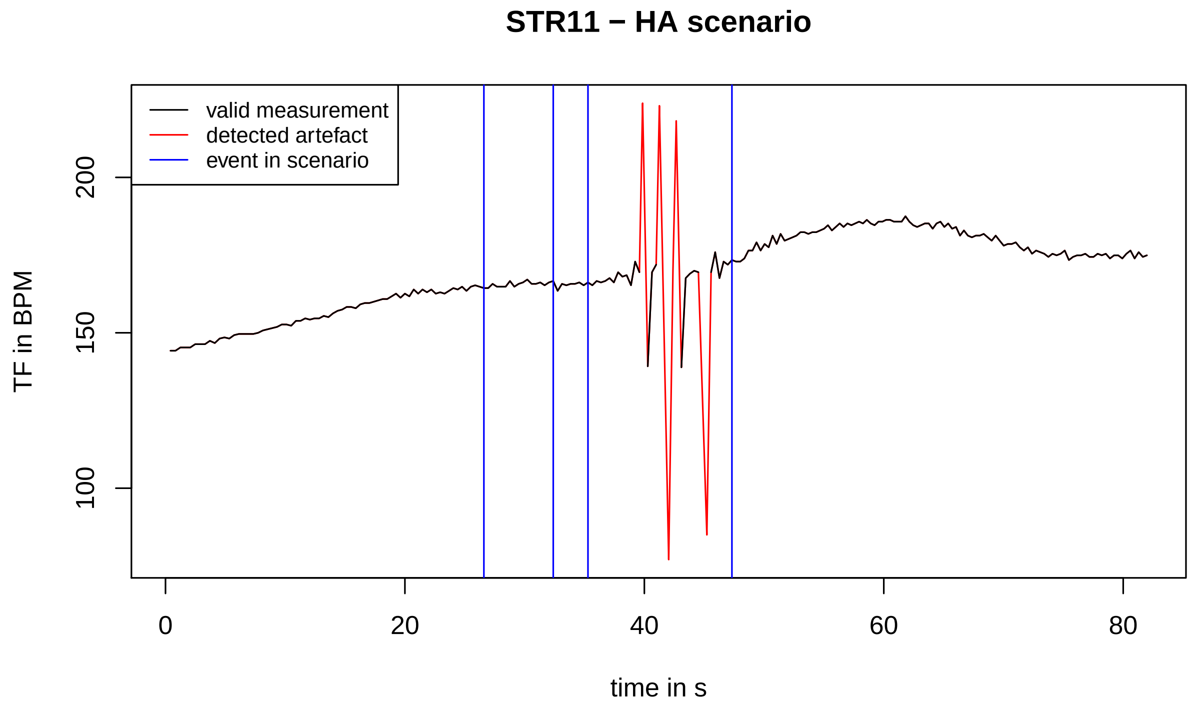
STR11 − HA scenario
valid measurement
detected artefact
event in scenario
TF in BPM
100
150
200
0
20
40
60
80
time in s

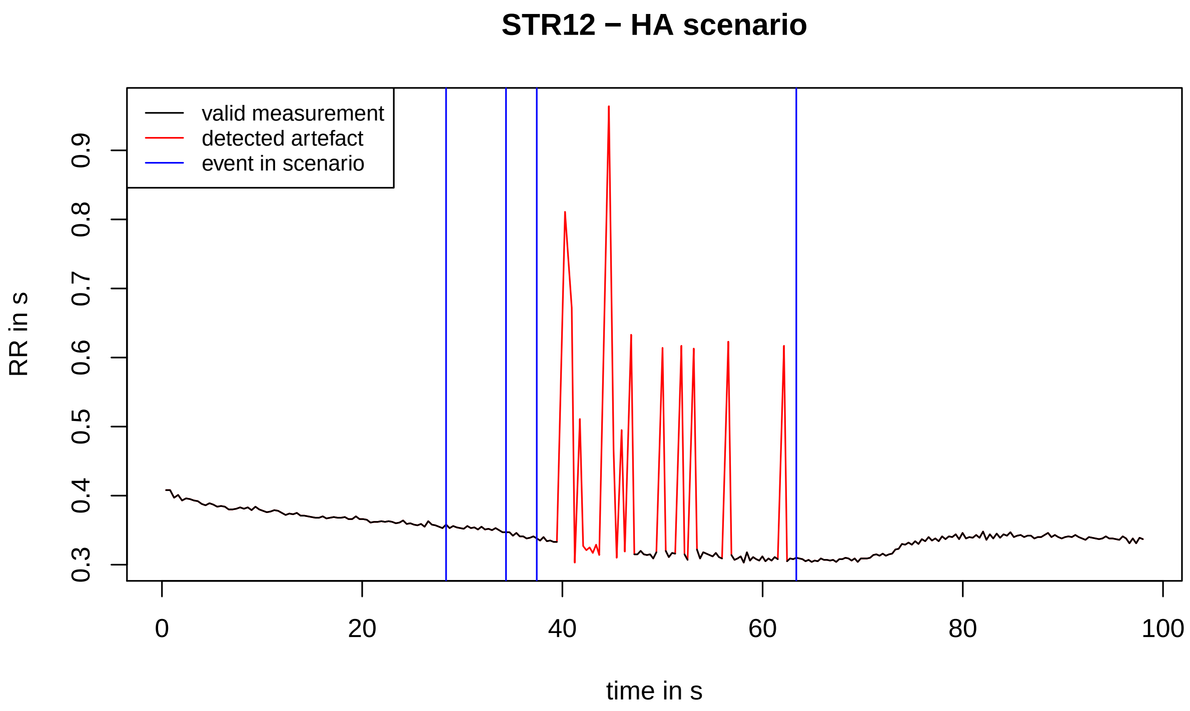
STR12 − HA scenario
valid measurement
detected artefact
event in scenario
RR in s
0.3
0.4
0.5
0.6
0.7
0.8
0.9
0
20
40
60
80
100
time in s

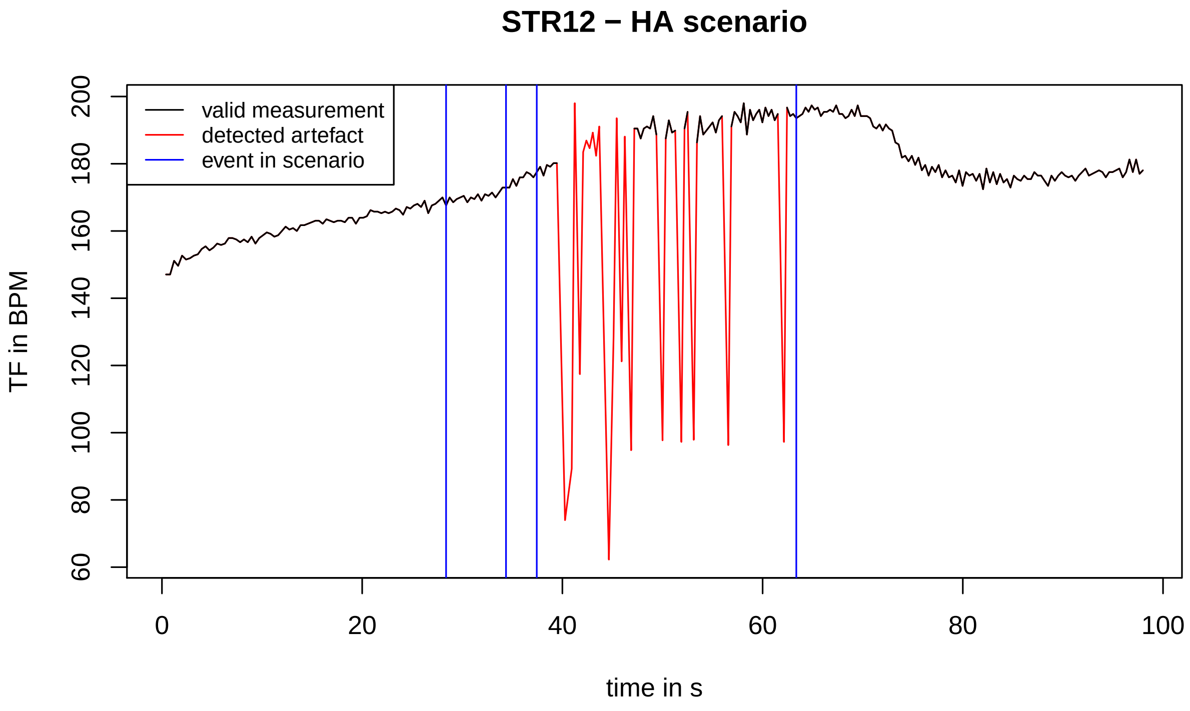
STR12 − HA scenario
valid measurement
detected artefact
event in scenario
TF in BPM
60
80
100
120
140
160
180
200
0
20
40
60
80
100
time in s

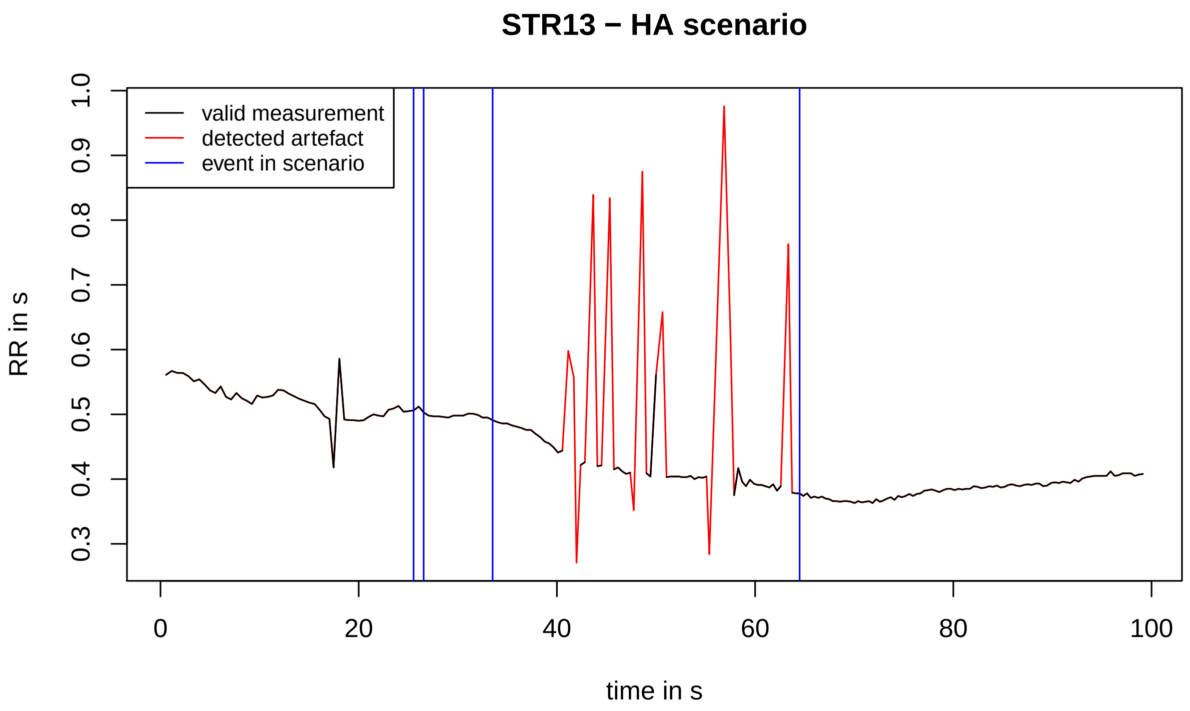
STR13 − HA scenario
valid measurement
detected artefact
event in scenario
RR in s
0.3 0.4 0.5 0.6 0.7 0.8 0.9 1.0
0 20 40 60 80 100
time in s

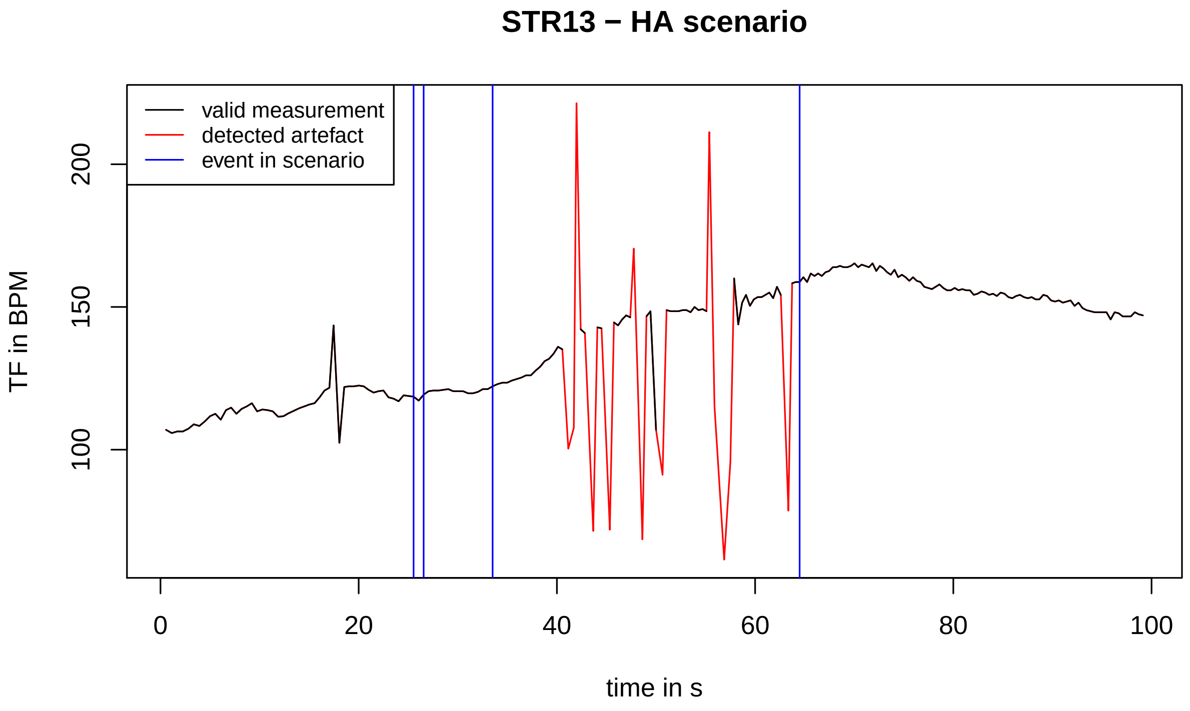
STR13 − HA scenario
valid measurement
detected artefact
event in scenario
TF in BPM
100 150 200
0 20 40 60 80 100
time in s

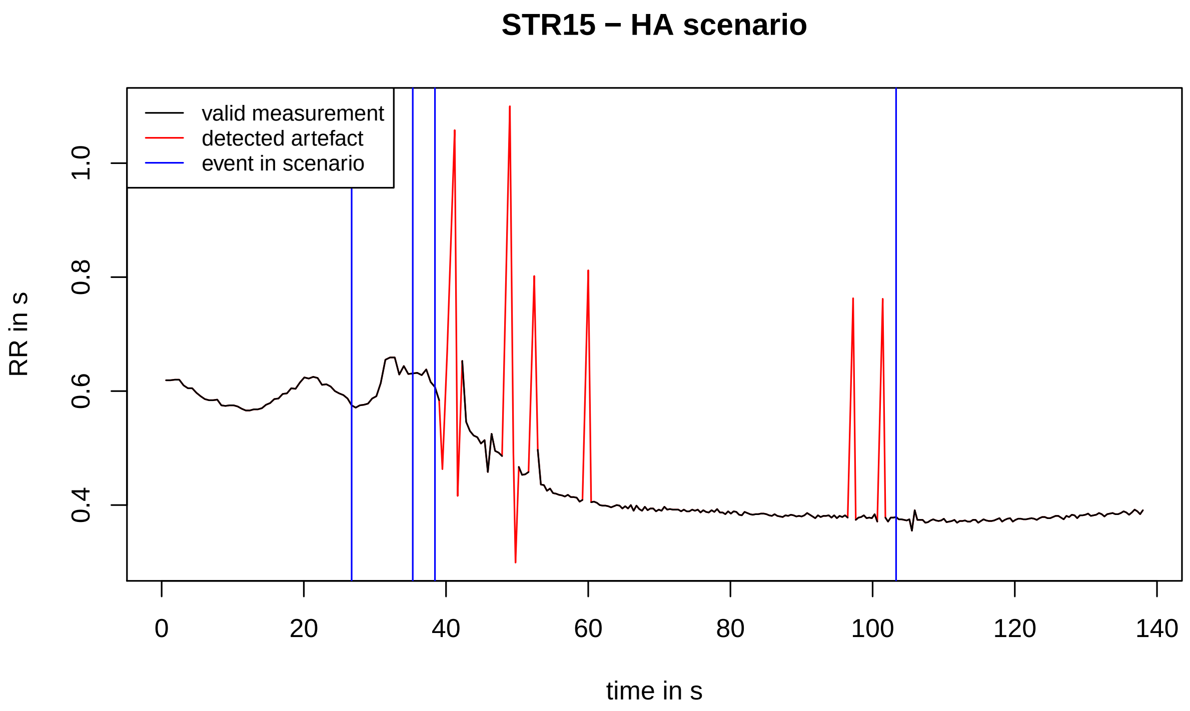
STR15 − HA scenario
valid measurement
detected artefact
event in scenario
RR in s
time in s

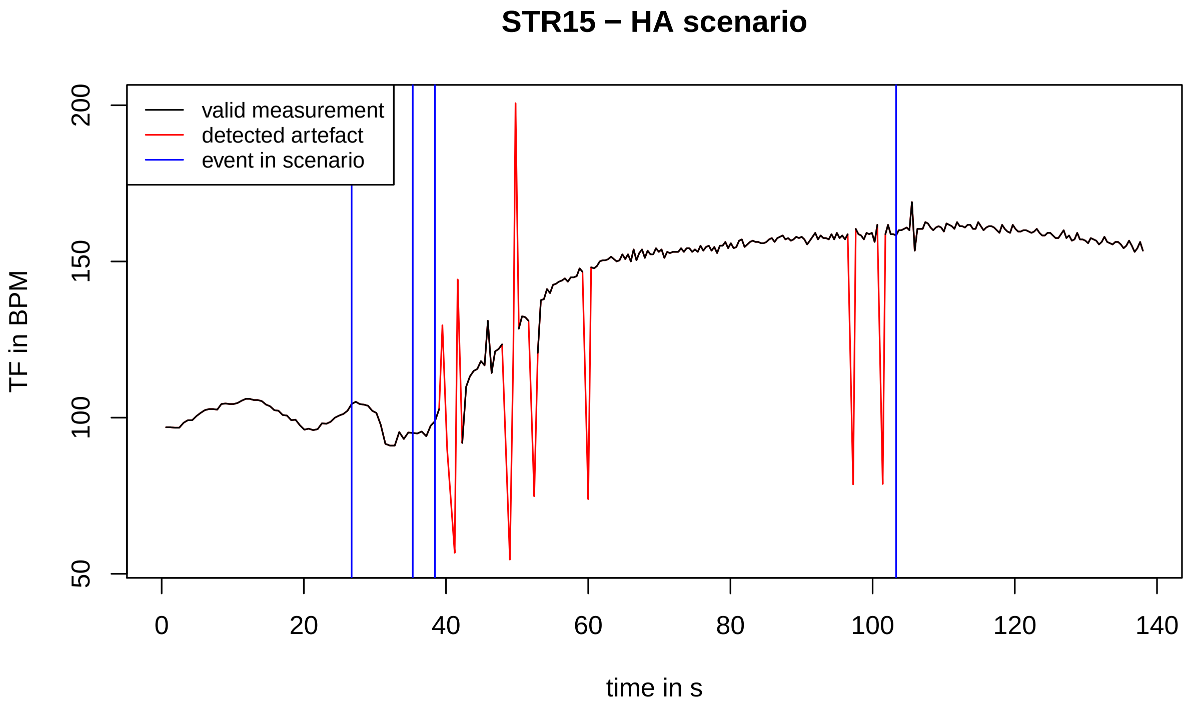
STR15 − HA scenario
valid measurement
detected artefact
event in scenario
TF in BPM
time in s

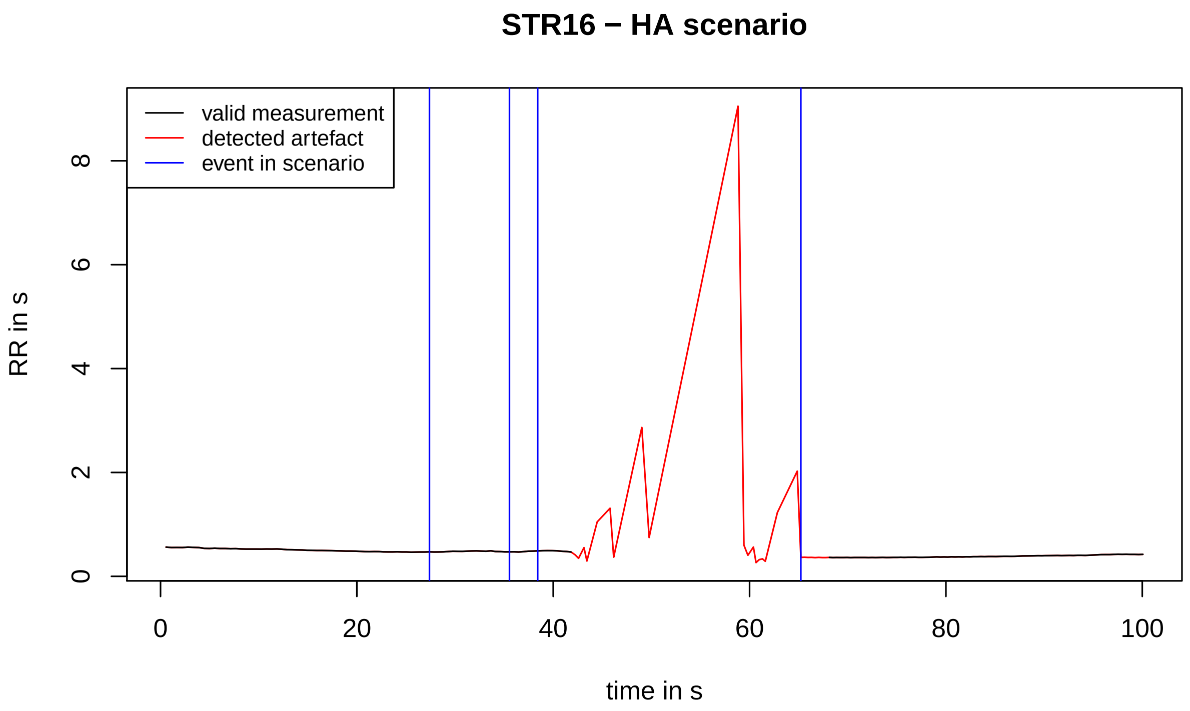
STR16 − HA scenario
valid measurement
detected artefact
event in scenario
RR in s
0
2
4
6
8
0
20
40
60
80
100
time in s

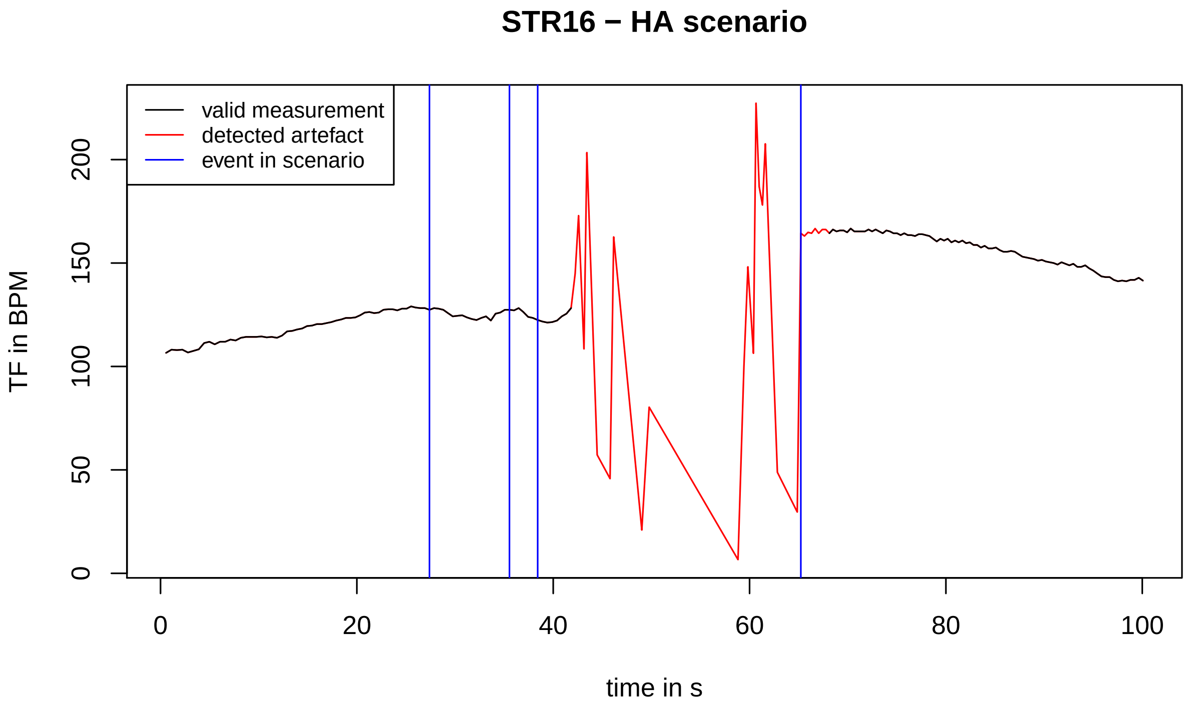
STR16 − HA scenario
valid measurement
detected artefact
event in scenario
TF in BPM
0
50
100
150
200
0
20
40
60
80
100
time in s

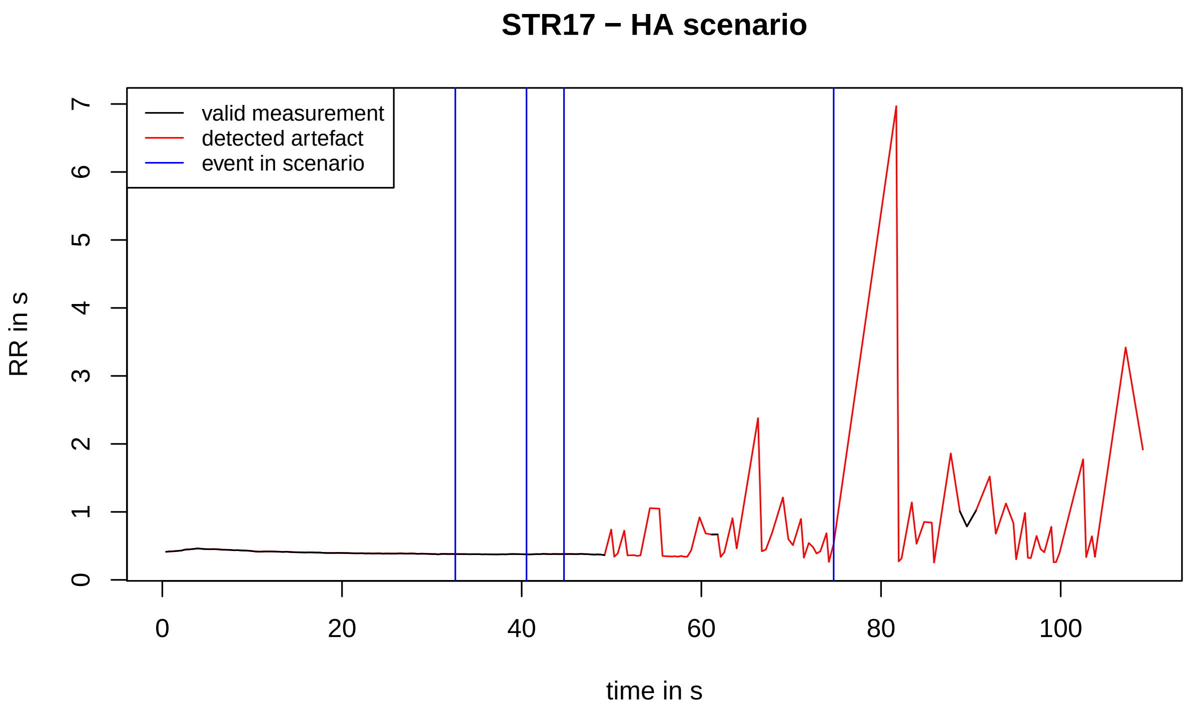
STR17 − HA scenario
valid measurement
detected artefact
event in scenario
RR in s
time in s
0
1
2
3
4
5
6
7
0
20
40
60
80
100

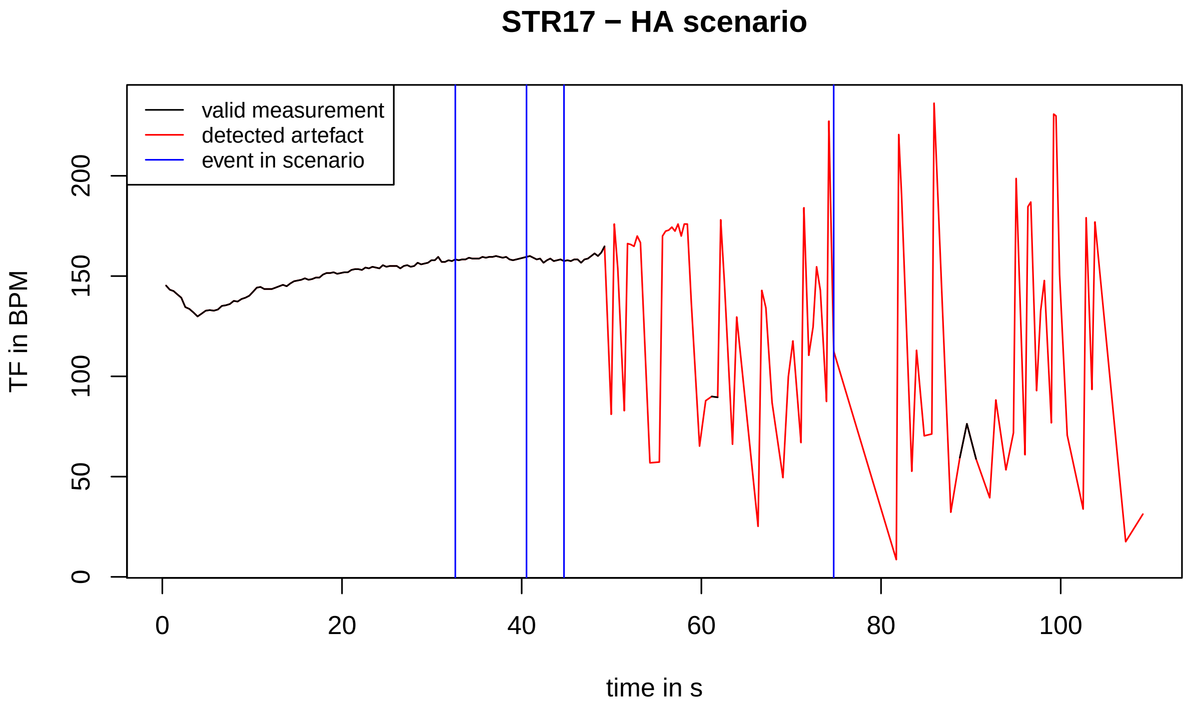
STR17 − HA scenario
valid measurement
detected artefact
event in scenario
TF in BPM
time in s
0
50
100
150
200
0
20
40
60
80
100

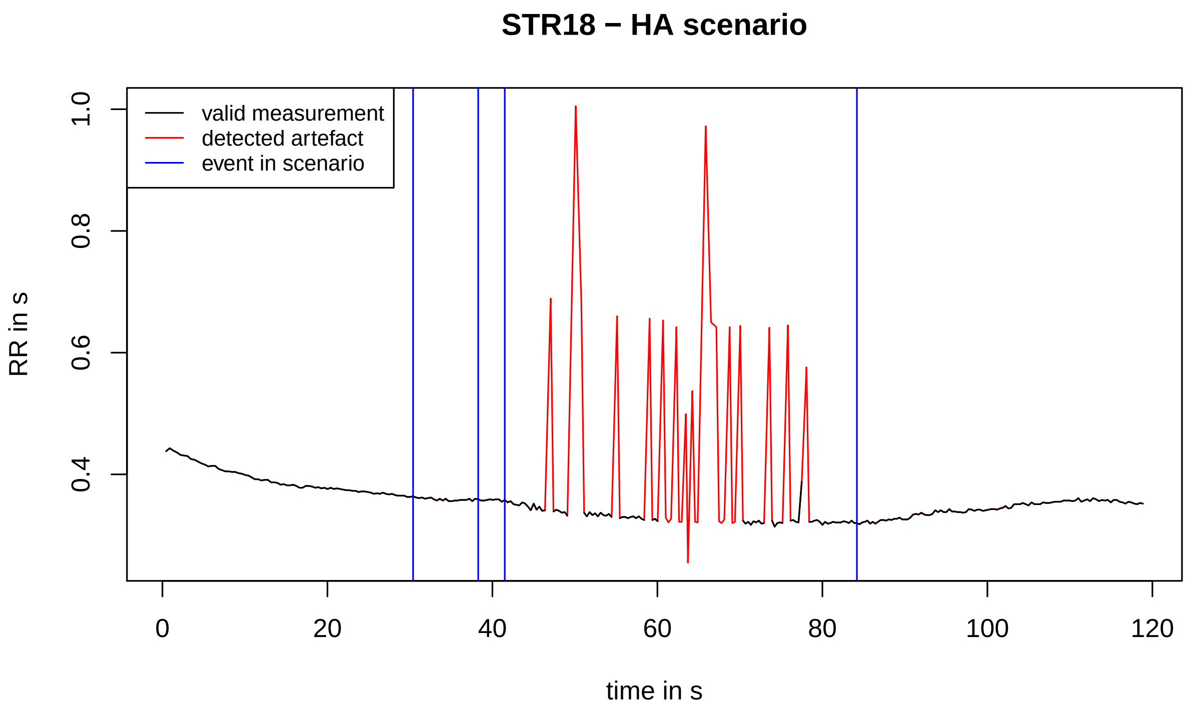
STR18 − HA scenario
valid measurement
detected artefact
event in scenario
RR in s
1.0
0.8
0.6
0.4
0
20
40
60
80
100
120
time in s

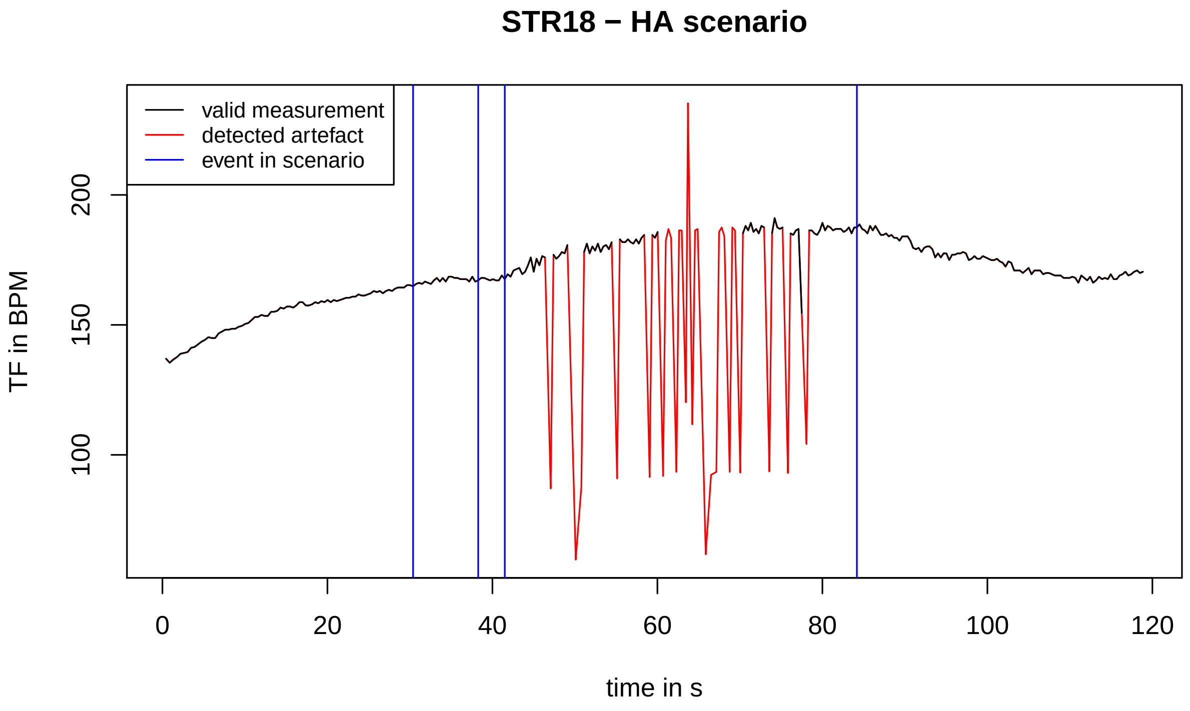
STR18 − HA scenario
valid measurement
detected artefact
event in scenario
TF in BPM
200
150
100
0
20
40
60
80
100
120
time in s

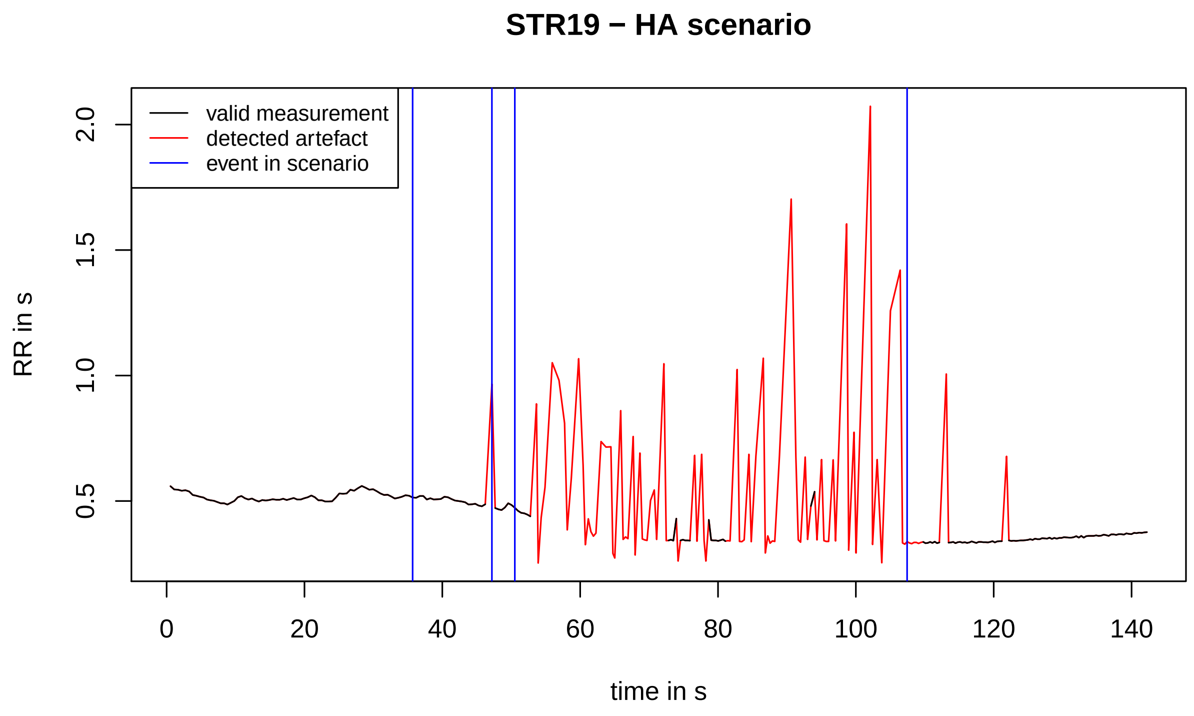

STR19 − HA scenario
valid measurement
detected artefact
event in scenario
RR in s
time in s
0.5
1.0
1.5
2.0
0
20
40
60
80
100
120
140


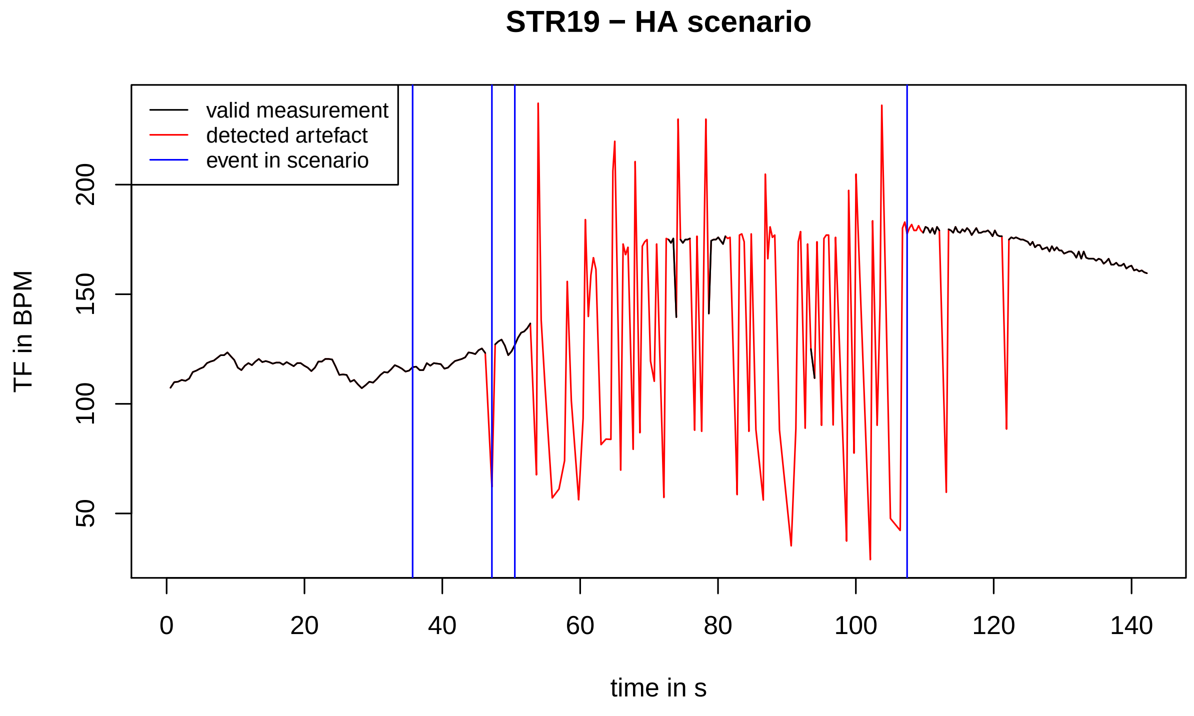

STR19 − HA scenario
valid measurement
detected artefact
event in scenario
TF in BPM
time in s
50
100
150
200
0
20
40
60
80
100
120
140

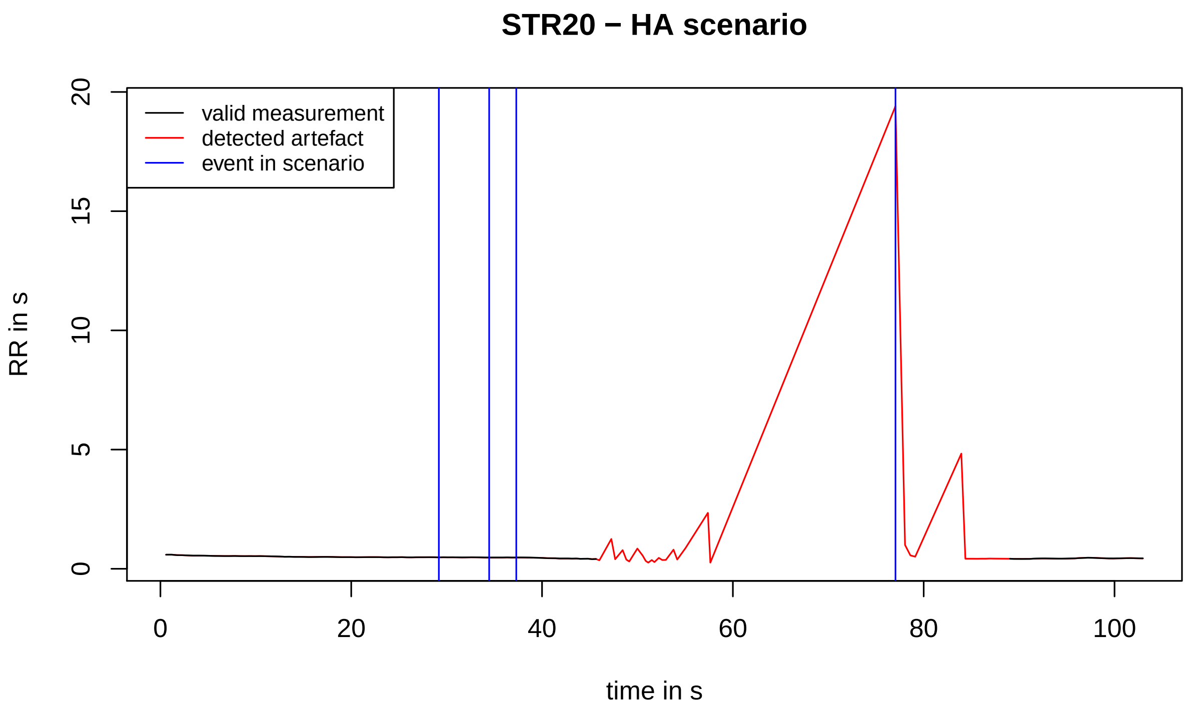
STR20 − HA scenario
valid measurement
detected artefact
event in scenario
RR in s
time in s
0
5
10
15
20
0
20
40
60
80
100

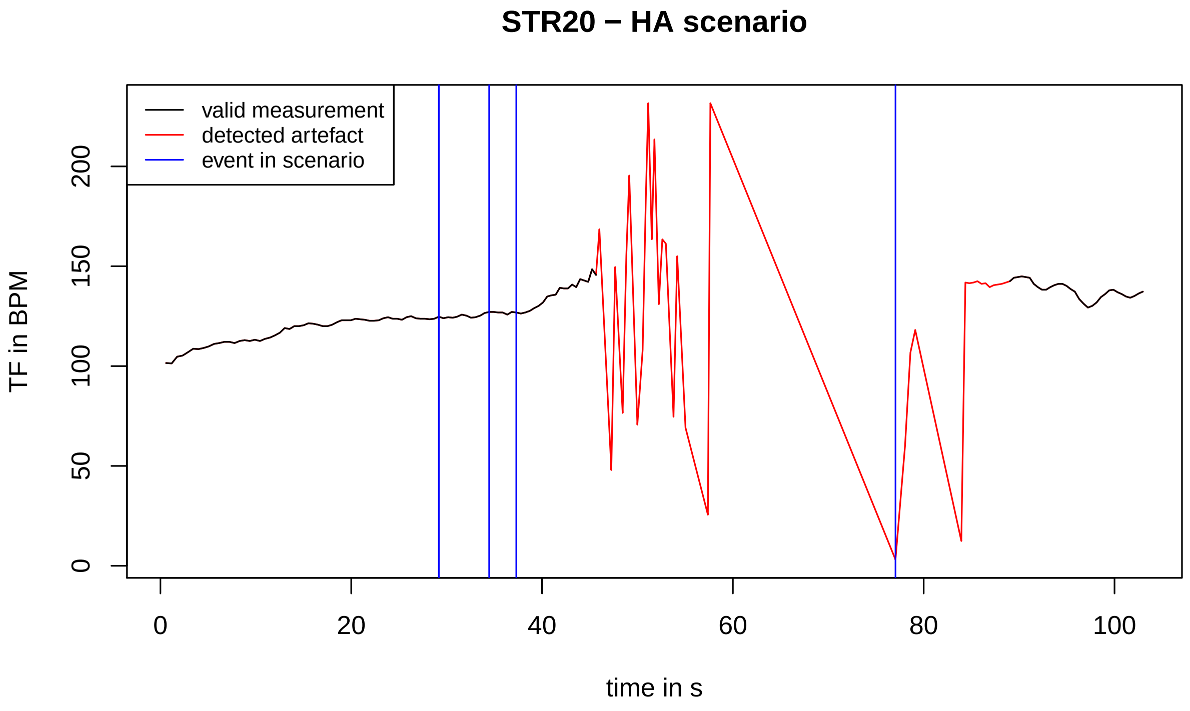
STR20 − HA scenario
valid measurement
detected artefact
event in scenario
TF in BPM
time in s
0
50
100
150
200
0
20
40
60
80
100

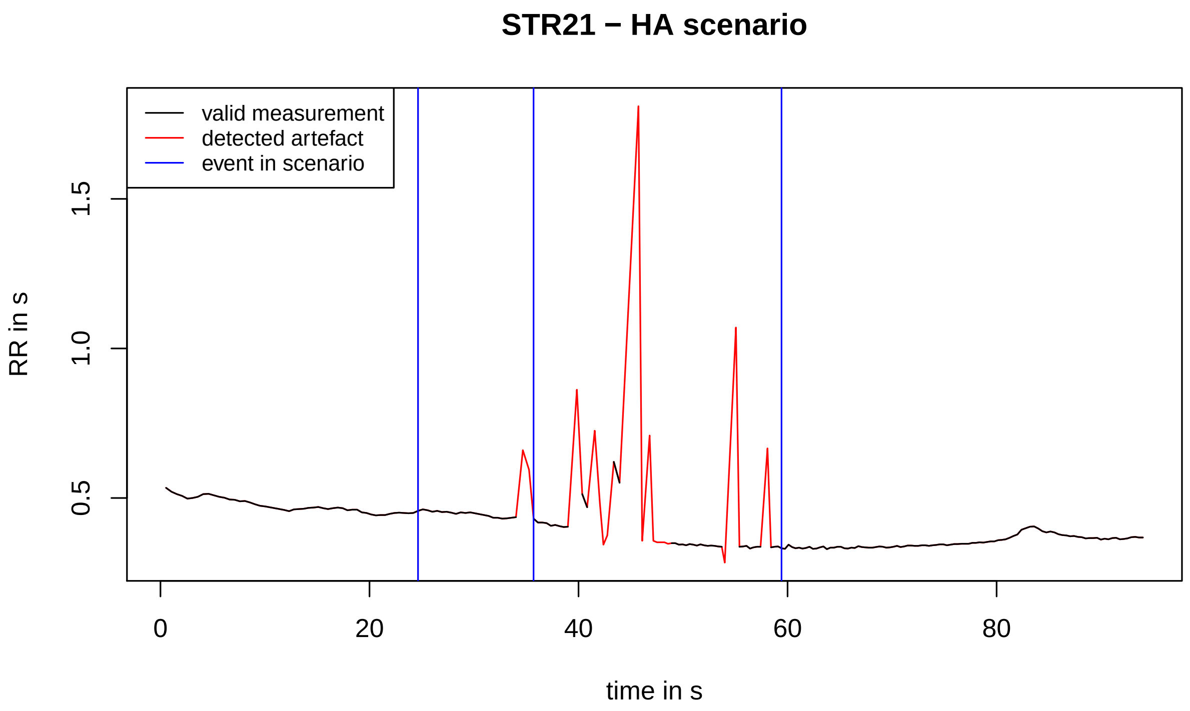
STR21 − HA scenario
valid measurement
detected artefact
event in scenario
RR in s
0.5
1.0
1.5
0
20
40
60
80
time in s

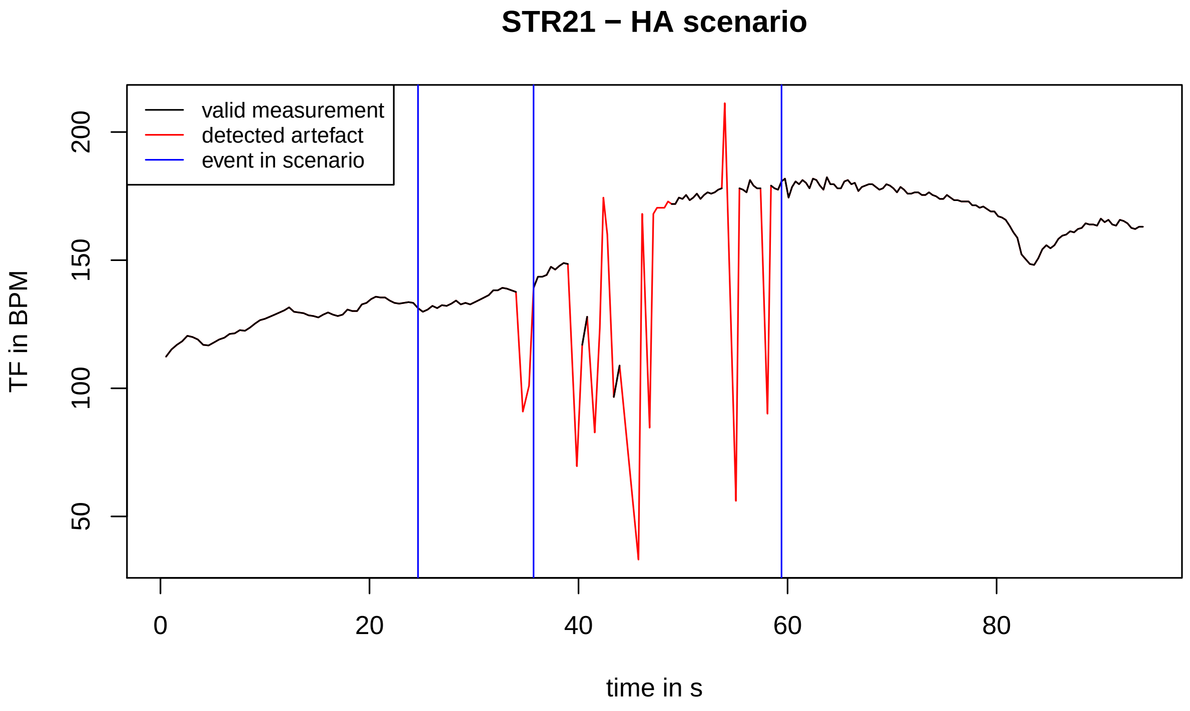
STR21 − HA scenario
valid measurement
detected artefact
event in scenario
TF in BPM
50
100
150
200
0
20
40
60
80
time in s

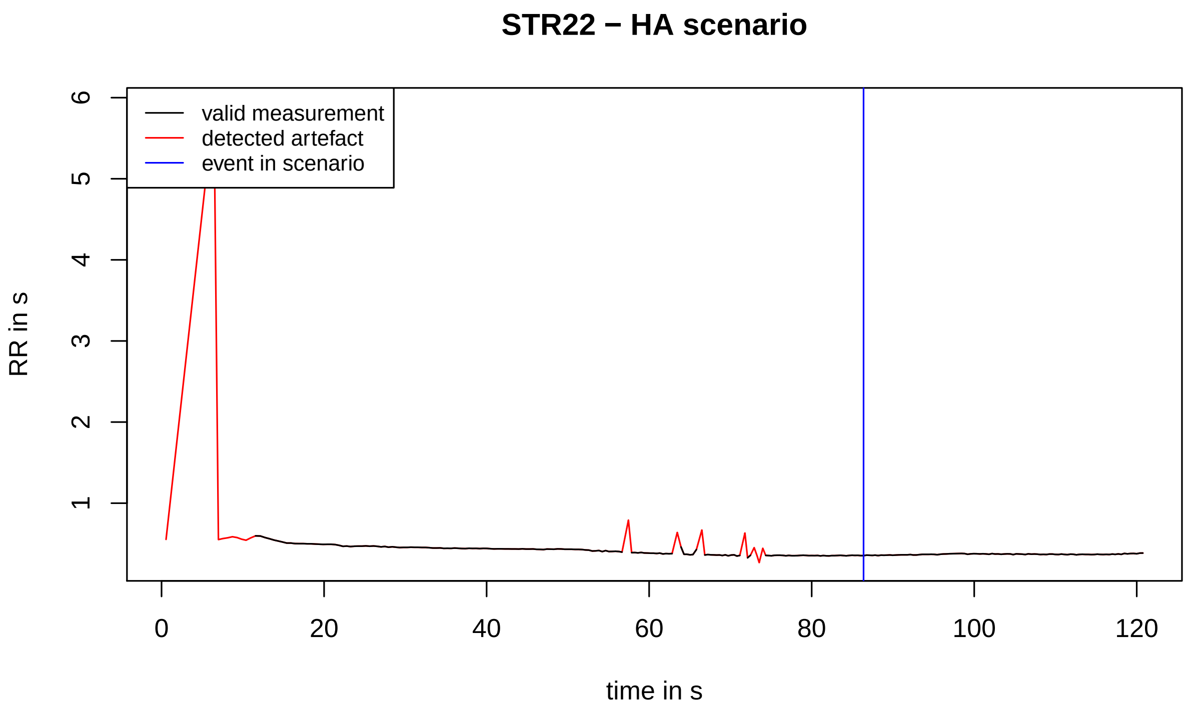
STR22 − HA scenario
valid measurement
detected artefact
event in scenario
RR in s
0
1
2
3
4
5
6
0
20
40
60
80
100
120
time in s

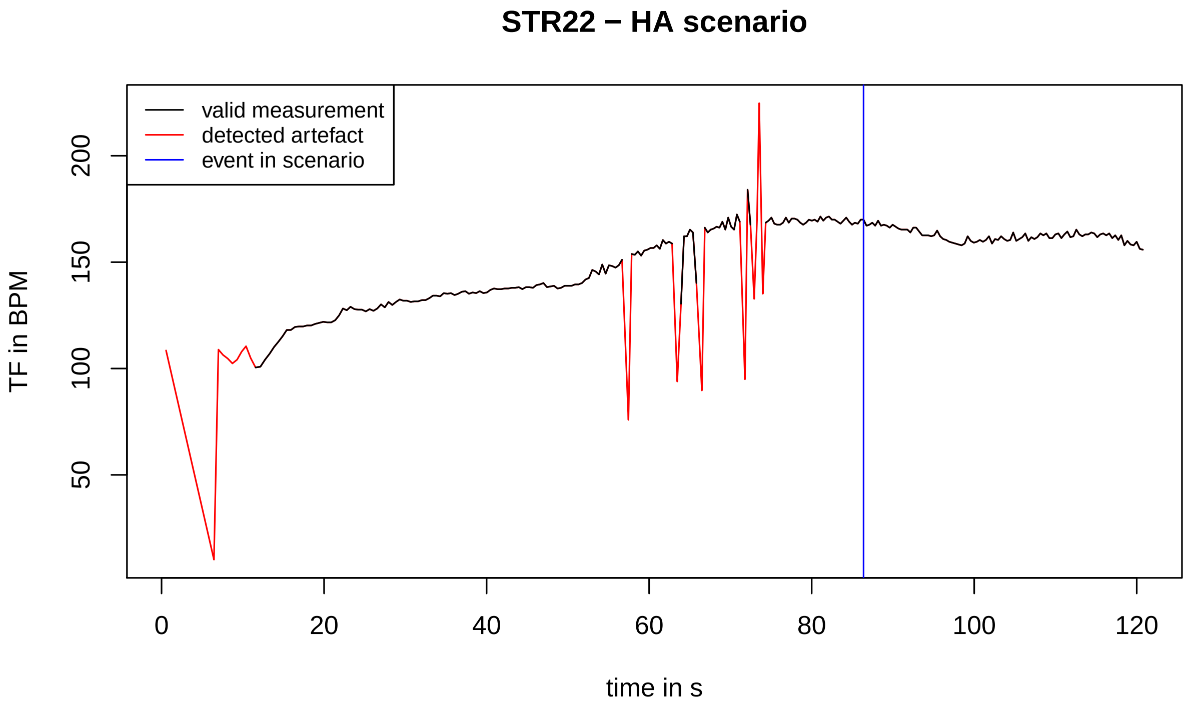
STR22 − HA scenario
valid measurement
detected artefact
event in scenario
TF in BPM
50
100
150
200
0
20
40
60
80
100
120
time in s

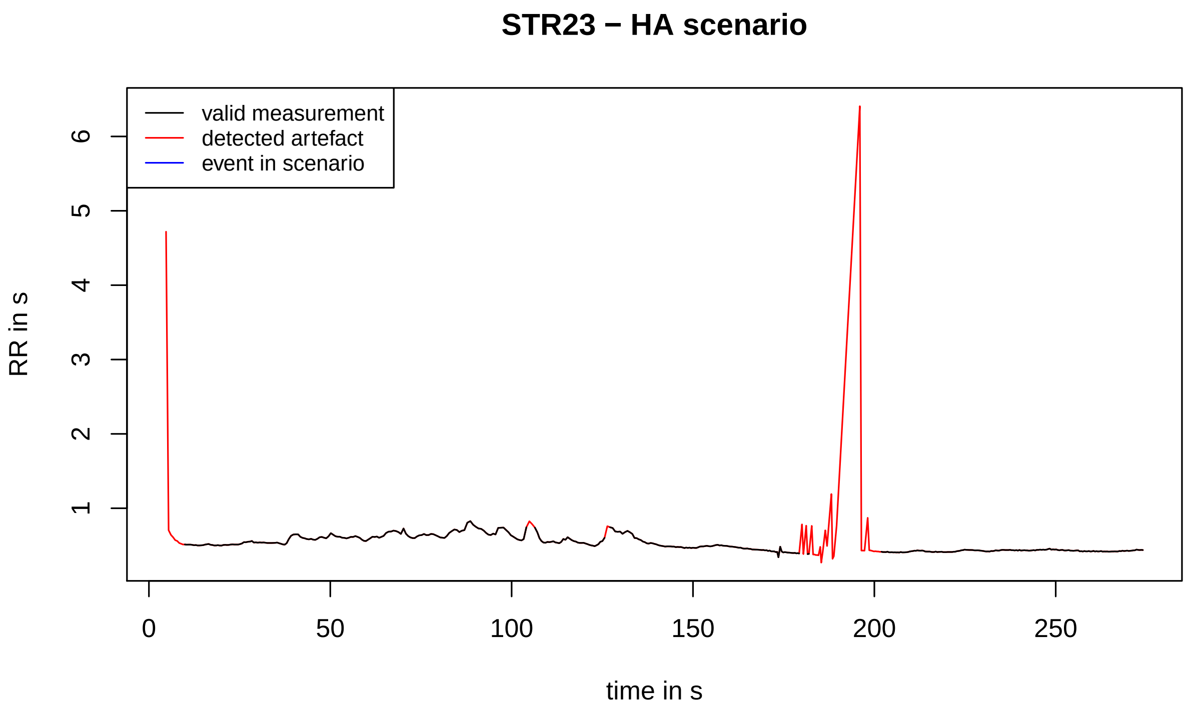
STR23 − HA scenario
valid measurement
detected artefact
event in scenario
RR in s
time in s
0
50
100
150
200
250
1
2
3
4
5
6

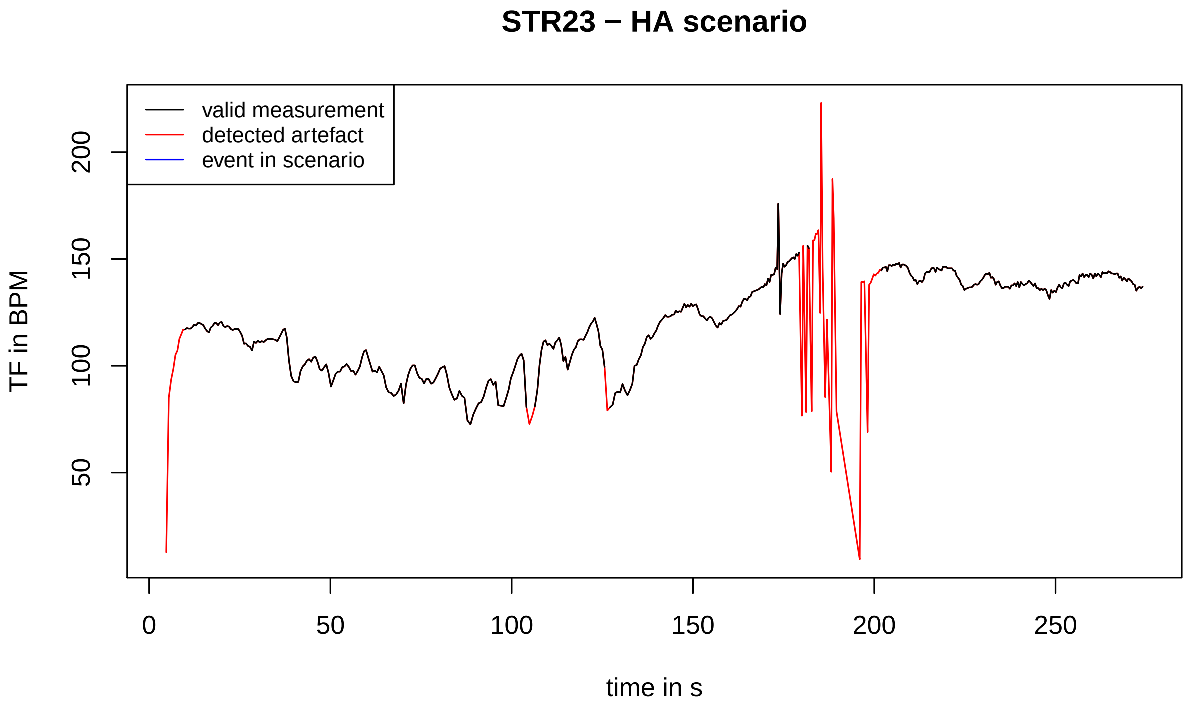
STR23 − HA scenario
valid measurement
detected artefact
event in scenario
TF in BPM
time in s
0
50
100
150
200
250
50
100
150
200

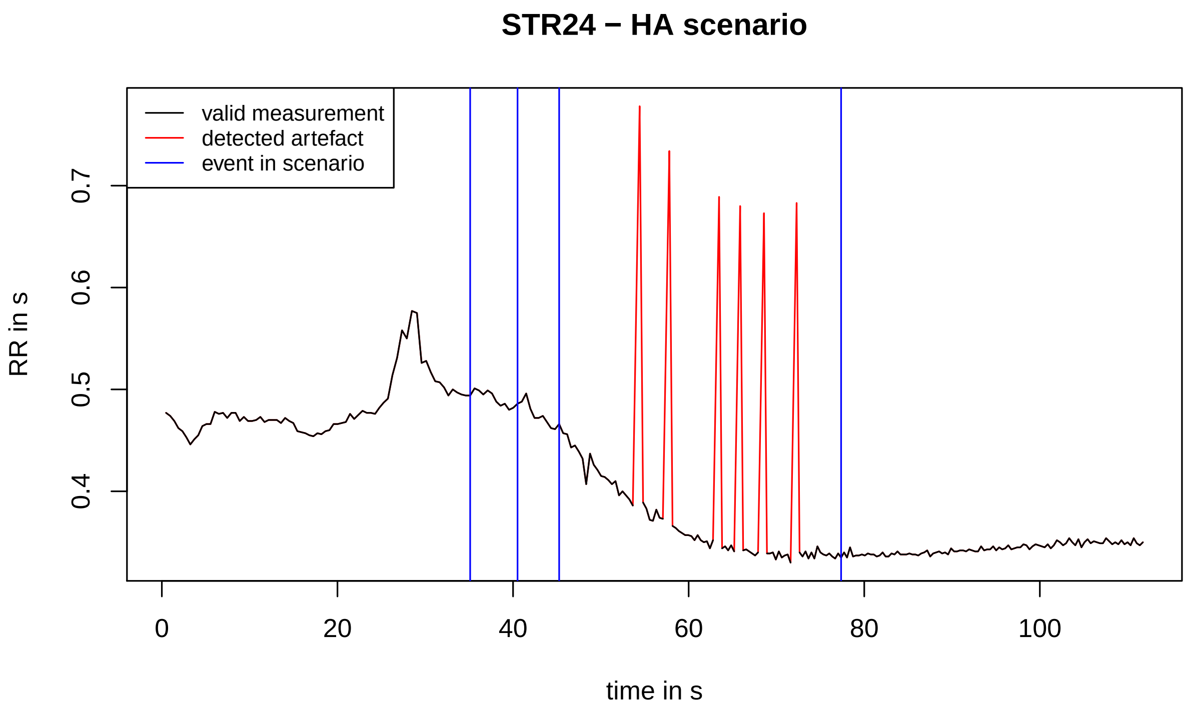
STR24 − HA scenario
valid measurement
detected artefact
event in scenario
RR in s
0.4
0.5
0.6
0.7
0
20
40
60
80
100
time in s

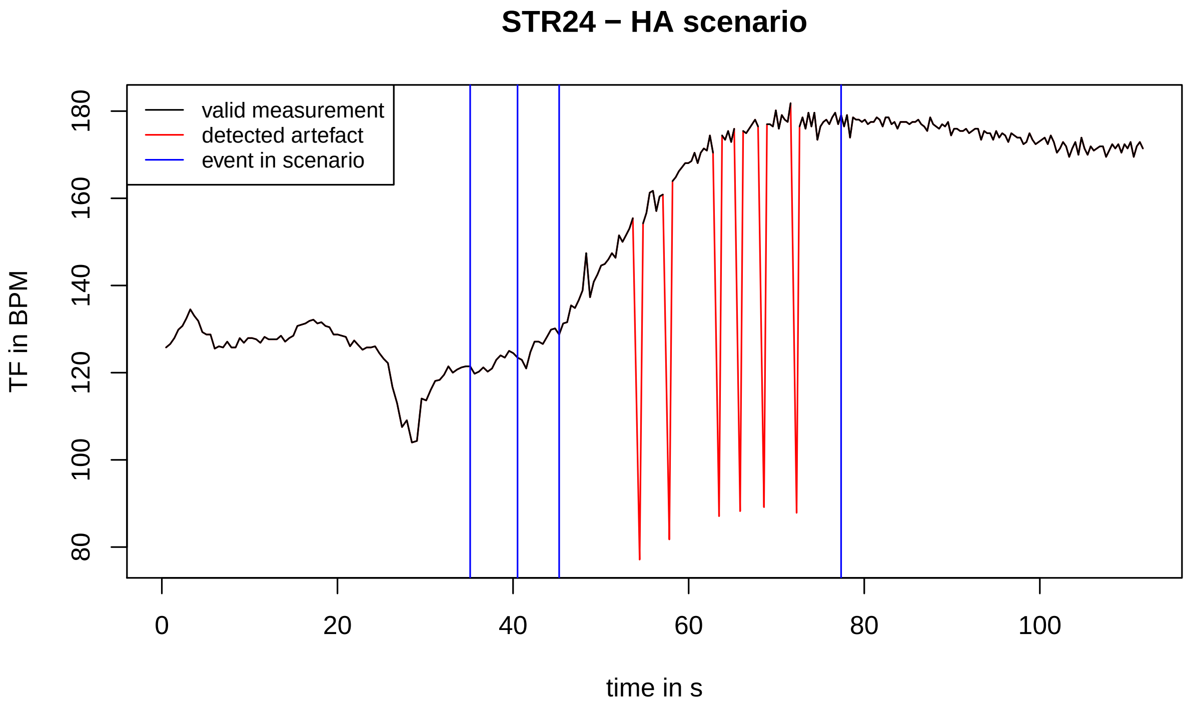
STR24 − HA scenario
valid measurement
detected artefact
event in scenario
TF in BPM
80
100
120
140
160
180
0
20
40
60
80
100
time in s

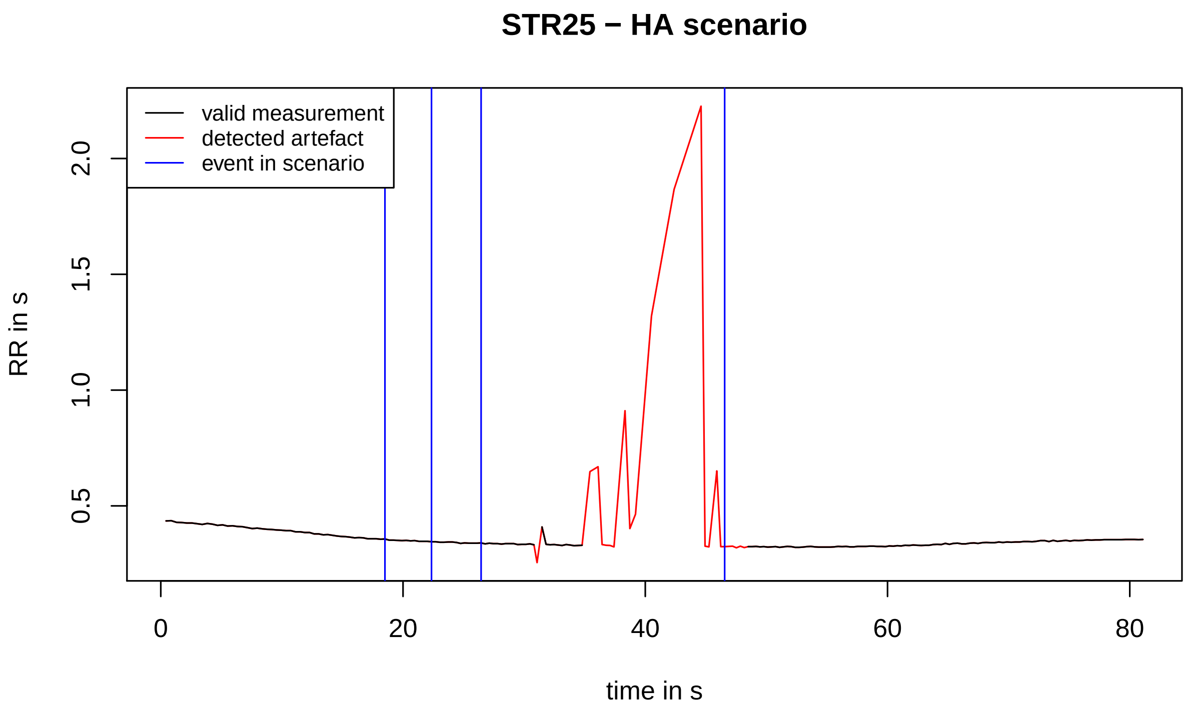
STR25 − HA scenario
valid measurement
detected artefact
event in scenario
RR in s
0.5
1.0
1.5
2.0
0
20
40
60
80
time in s

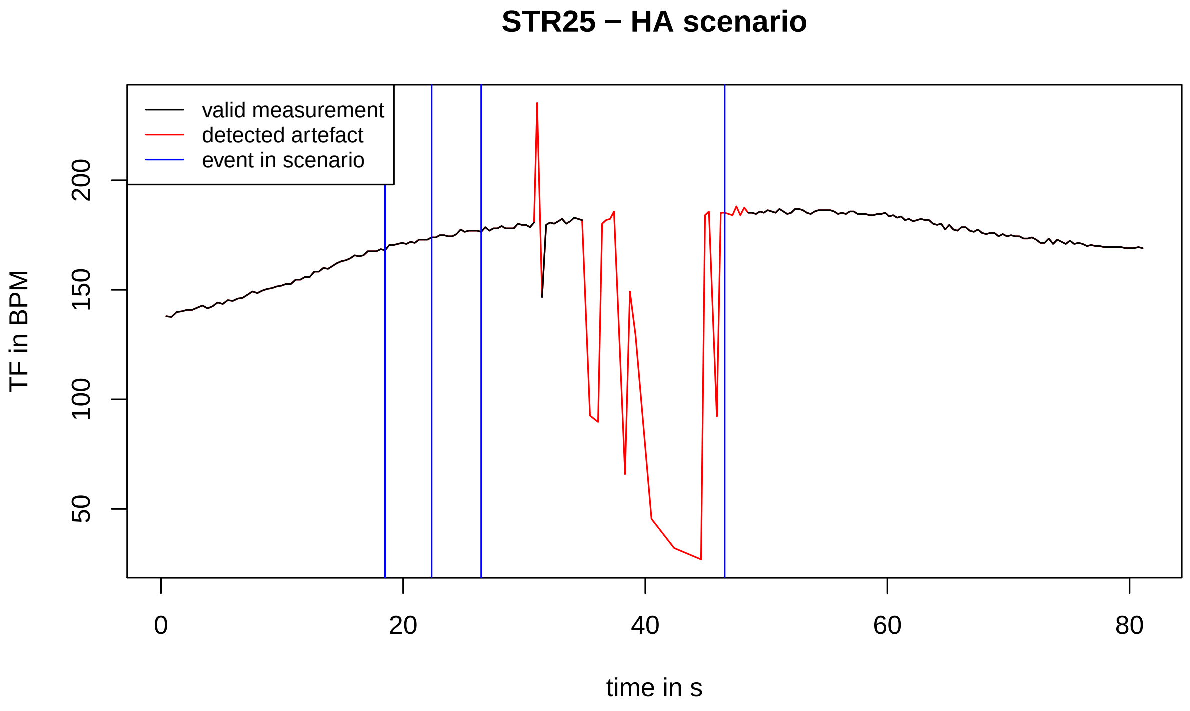
STR25 − HA scenario
valid measurement
detected artefact
event in scenario
TF in BPM
50
100
150
200
0
20
40
60
80
time in s

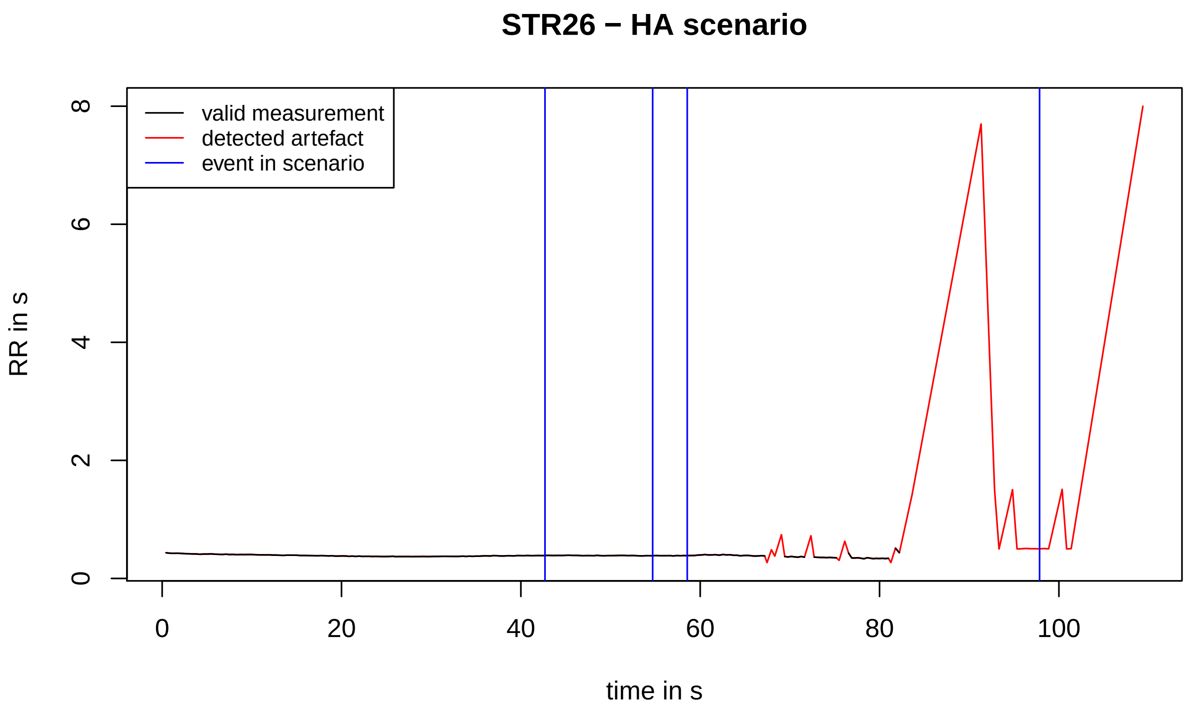
STR26 − HA scenario
valid measurement
detected artefact
event in scenario
RR in s
0
2
4
6
8
0
20
40
60
80
100
time in s

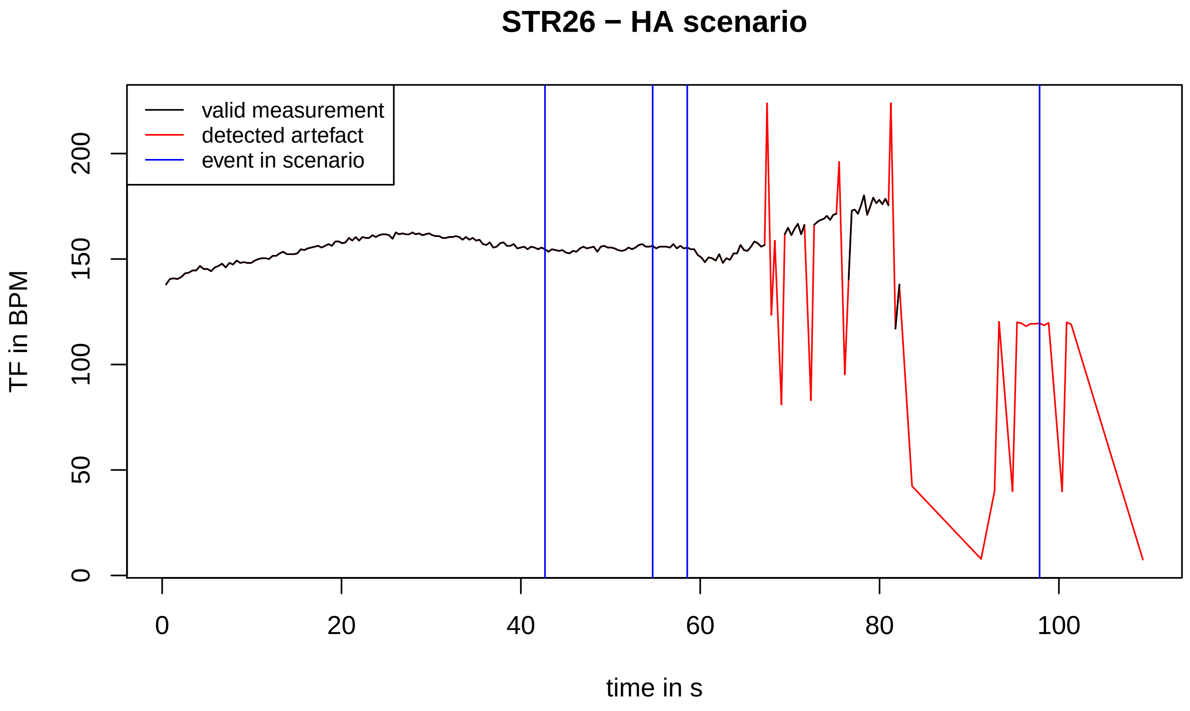
STR26 − HA scenario
valid measurement
detected artefact
event in scenario
TF in BPM
0
50
100
150
200
0
20
40
60
80
100
time in s

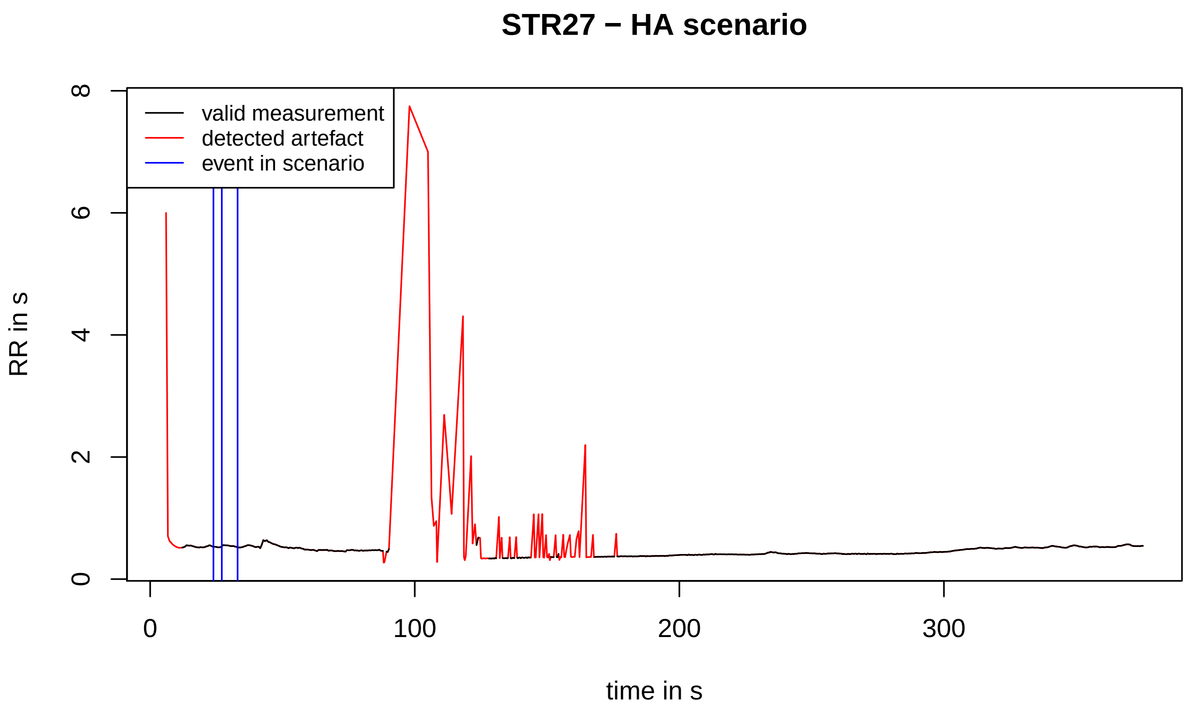
STR27 − HA scenario
valid measurement
detected artefact
event in scenario
RR in s
time in s
0
2
4
6
8
0
100
200
300

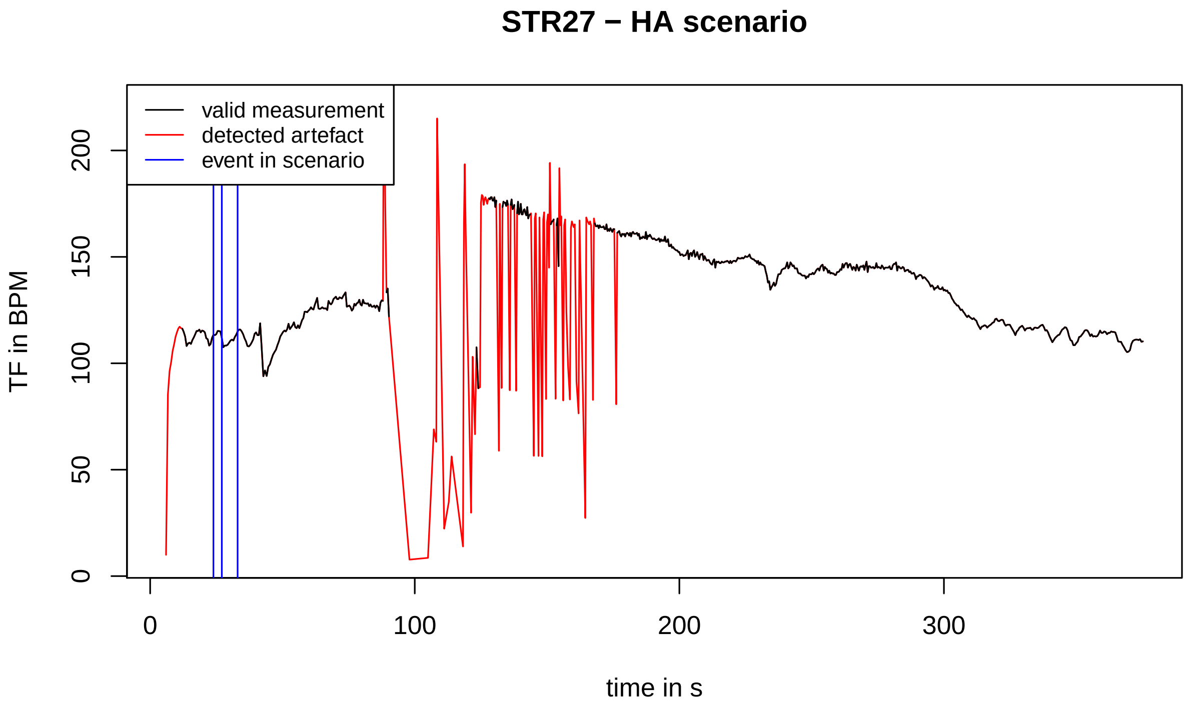
STR27 − HA scenario
valid measurement
detected artefact
event in scenario
TF in BPM
time in s
0
50
100
150
200
0
100
200
300

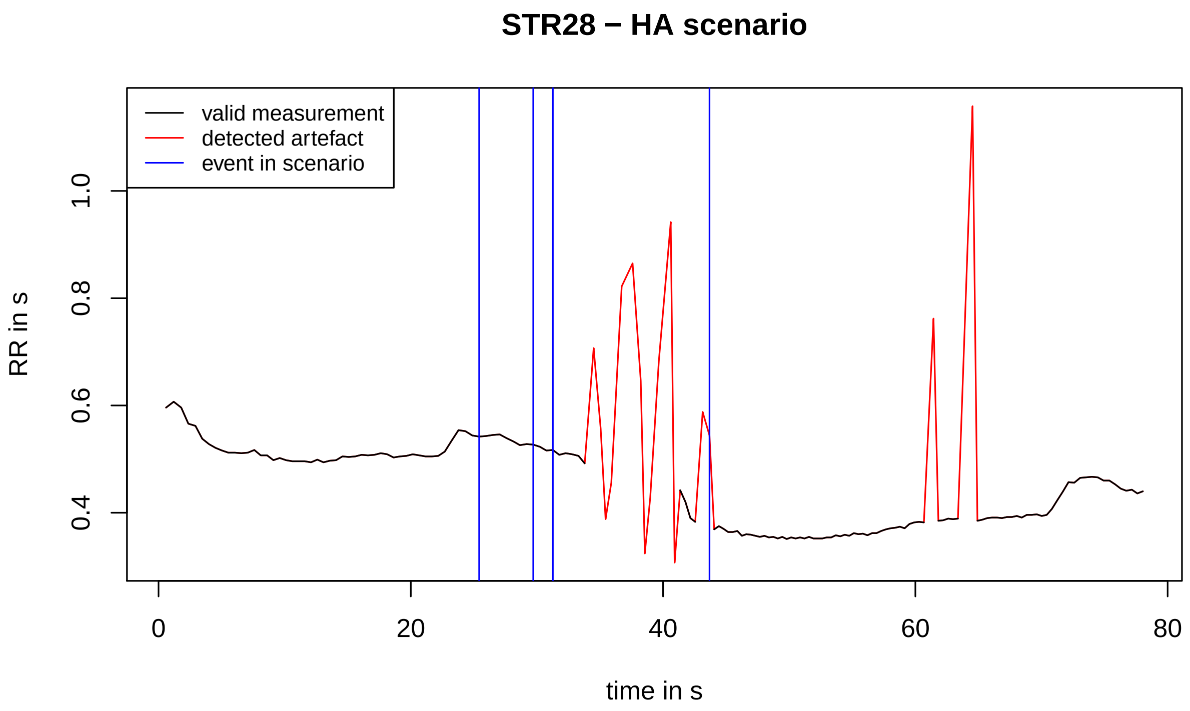
STR28 − HA scenario
valid measurement
detected artefact
event in scenario
RR in s
0.4
0.6
0.8
1.0
0
20
40
60
80
time in s

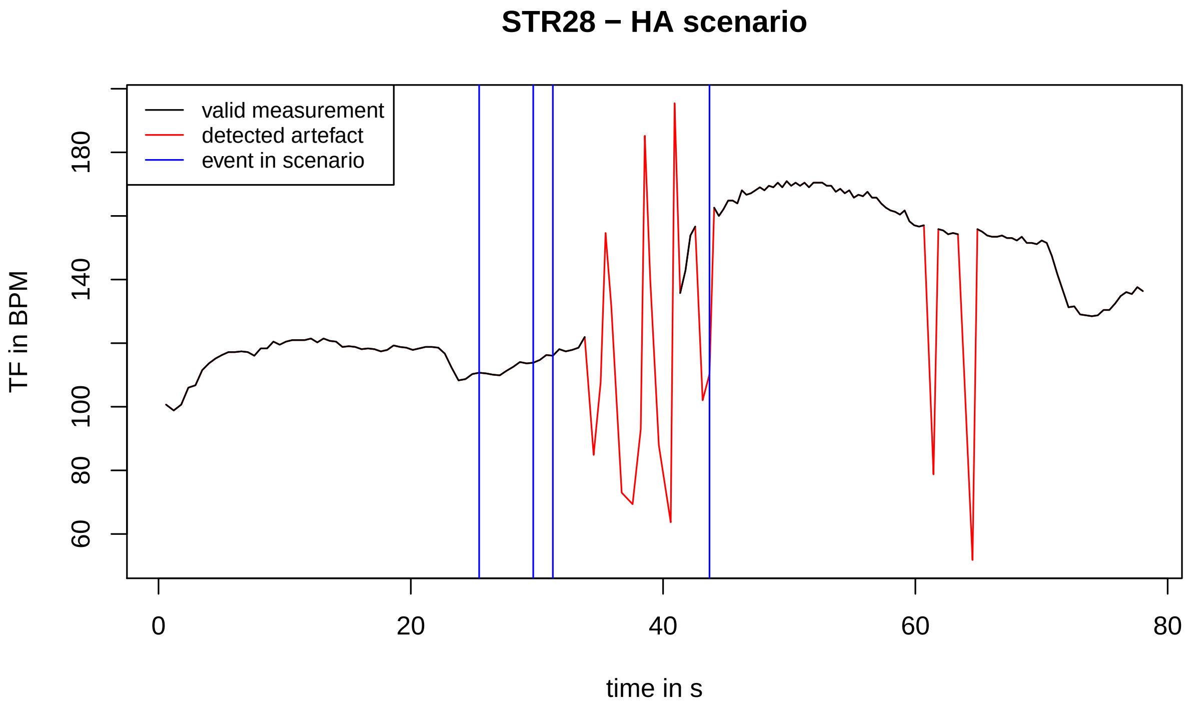
STR28 − HA scenario
valid measurement
detected artefact
event in scenario
TF in BPM
60
80
100
140
180
0
20
40
60
80
time in s

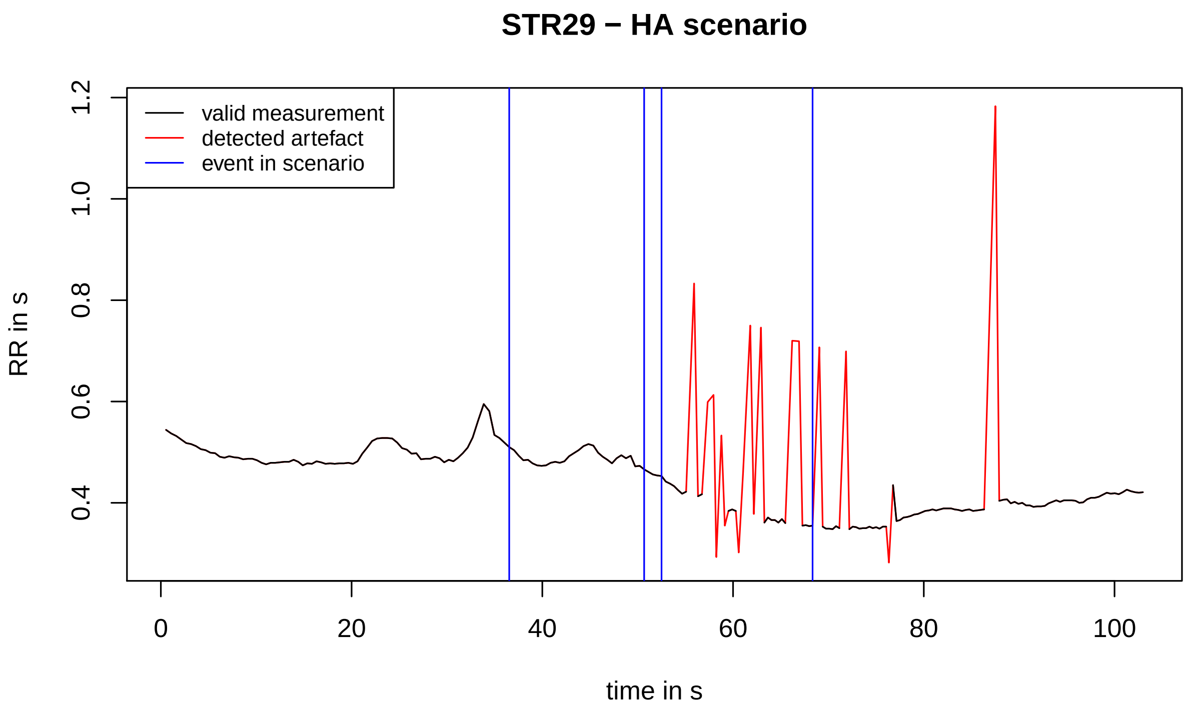

STR29 − HA scenario
valid measurement
detected artefact
event in scenario
RR in s
0.4
0.6
0.8
1.0
1.2
0
20
40
60
80
100
time in s


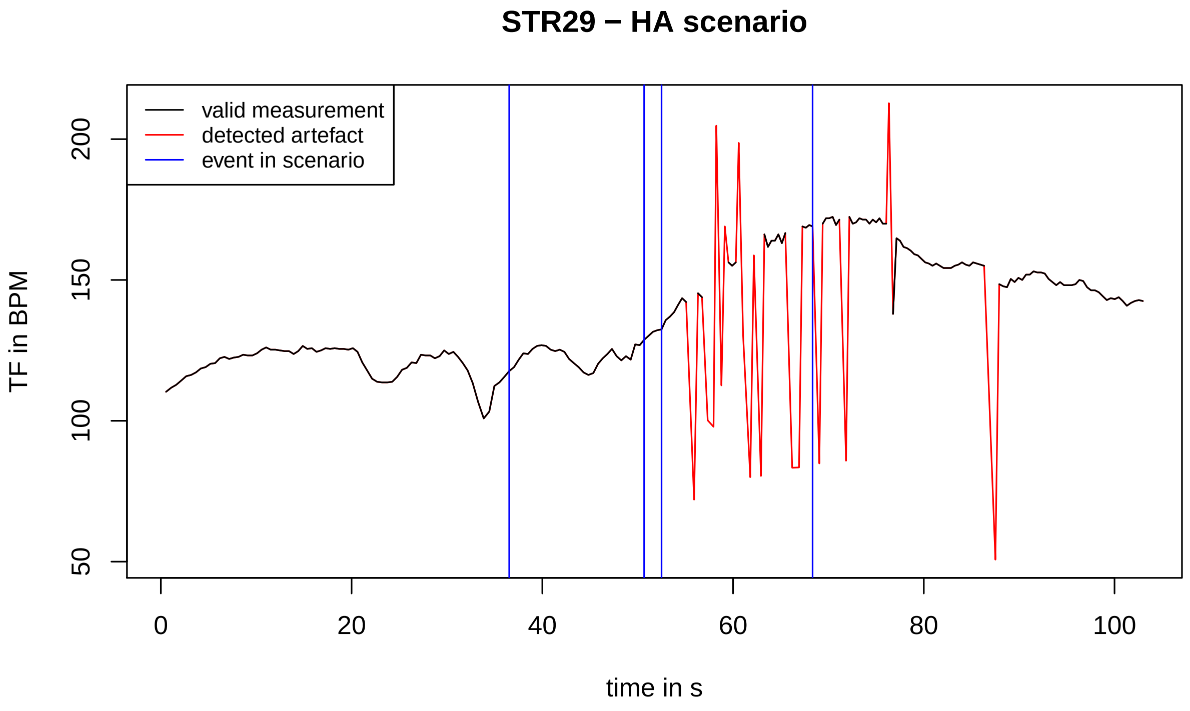

STR29 − HA scenario
valid measurement
detected artefact
event in scenario
TF in BPM
50
100
150
200
0
20
40
60
80
100
time in s

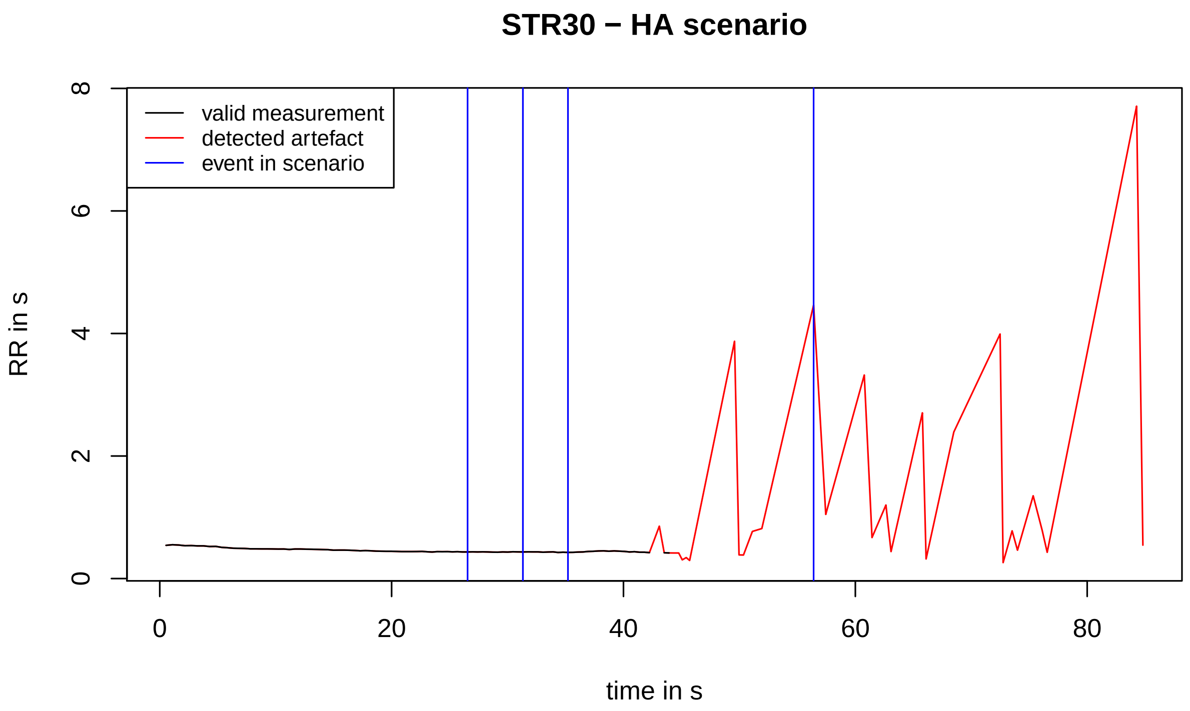
STR30 − HA scenario
valid measurement
detected artefact
event in scenario
RR in s
time in s
0
2
4
6
8
0
20
40
60
80

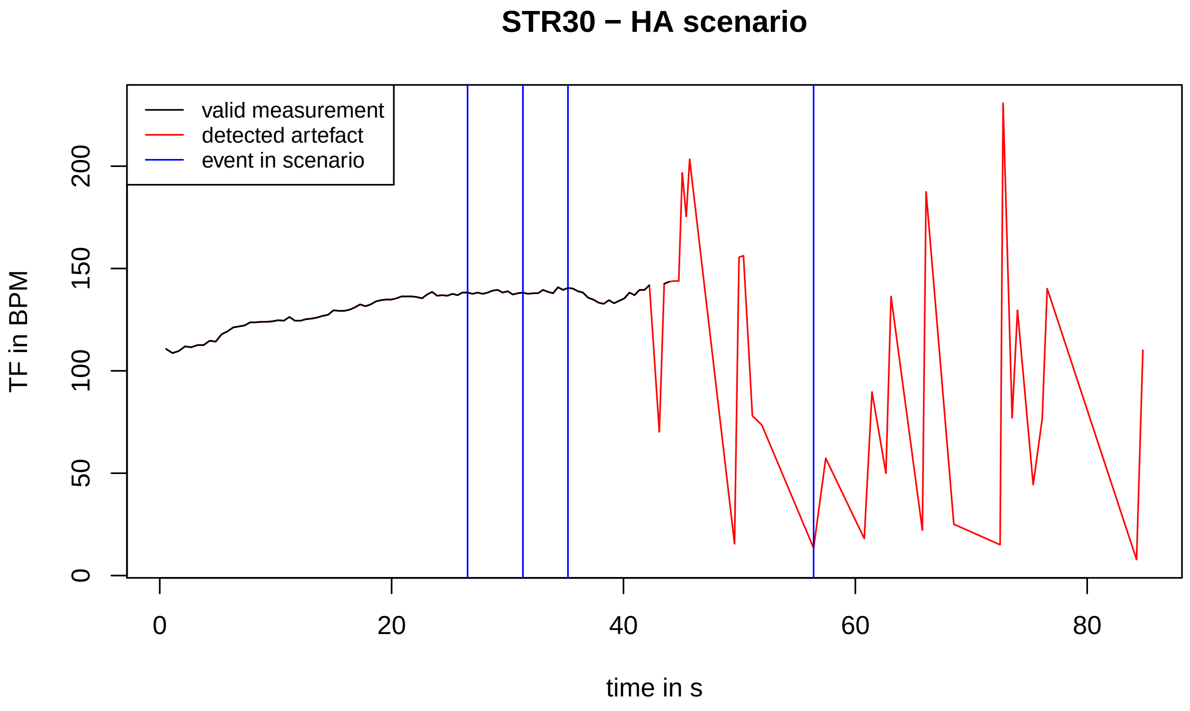
STR30 − HA scenario
valid measurement
detected artefact
event in scenario
TF in BPM
time in s
0
50
100
150
200
0
20
40
60
80

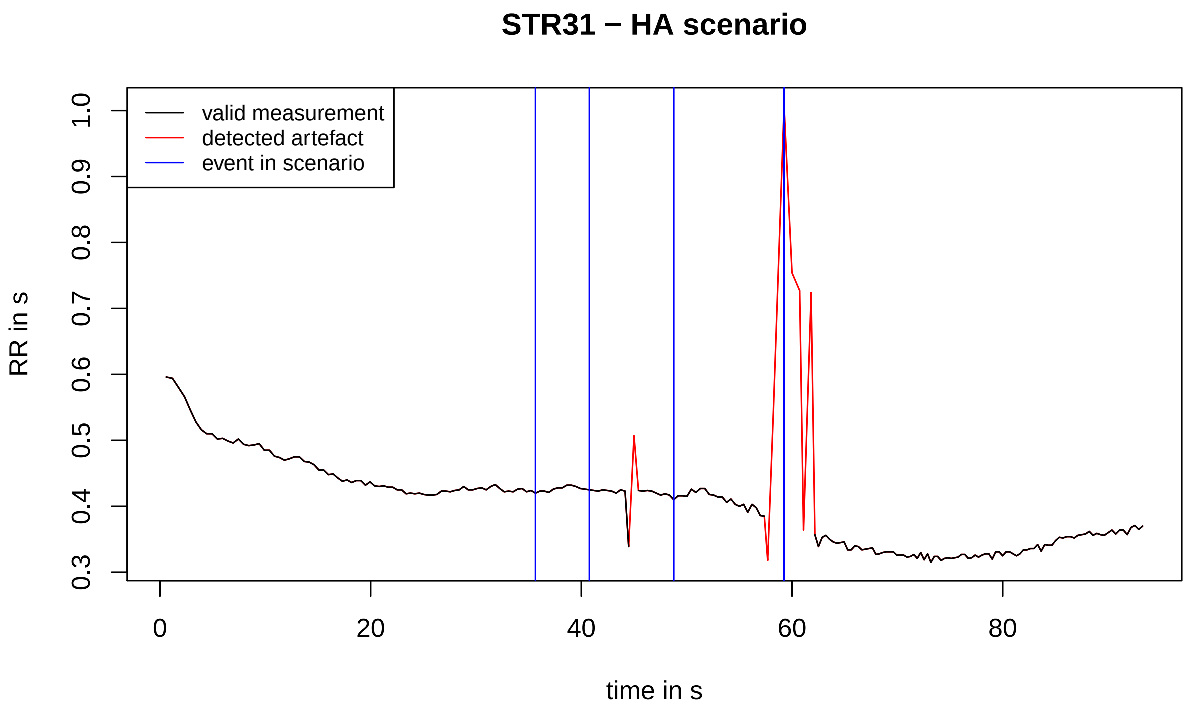
STR31 − HA scenario
valid measurement
detected artefact
event in scenario
RR in s
0.3
0.4
0.5
0.6
0.7
0.8
0.9
1.0
0
20
40
60
80
time in s

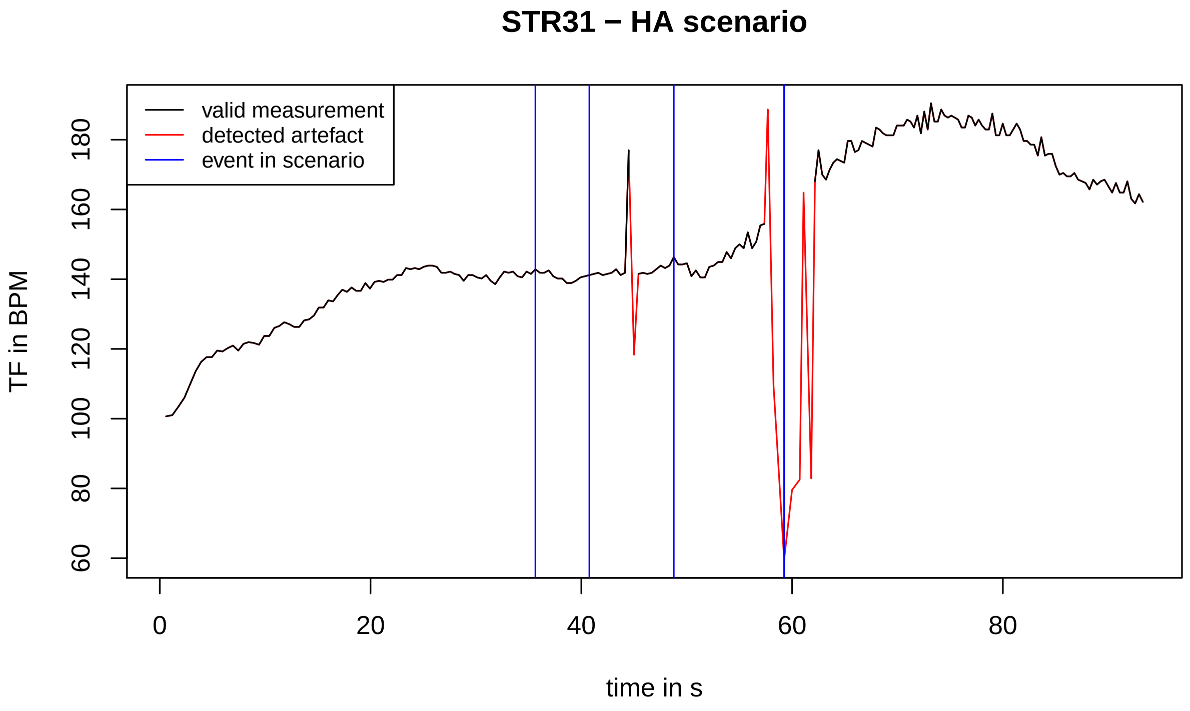
STR31 − HA scenario
valid measurement
detected artefact
event in scenario
TF in BPM
60
80
100
120
140
160
180
0
20
40
60
80
time in s

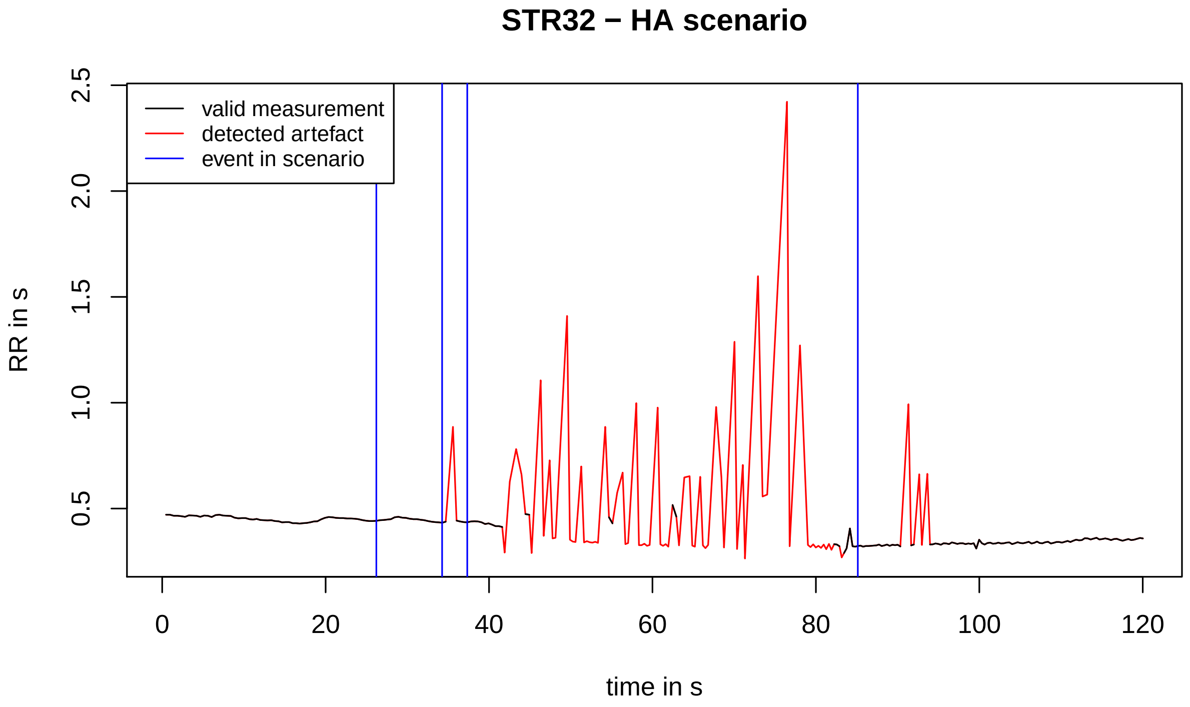
STR32 − HA scenario
RR in s
2.5
2.0
1.5
1.0
0.5
0
20
40
60
80
100
120
time in s
valid measurement
detected artefact
event in scenario

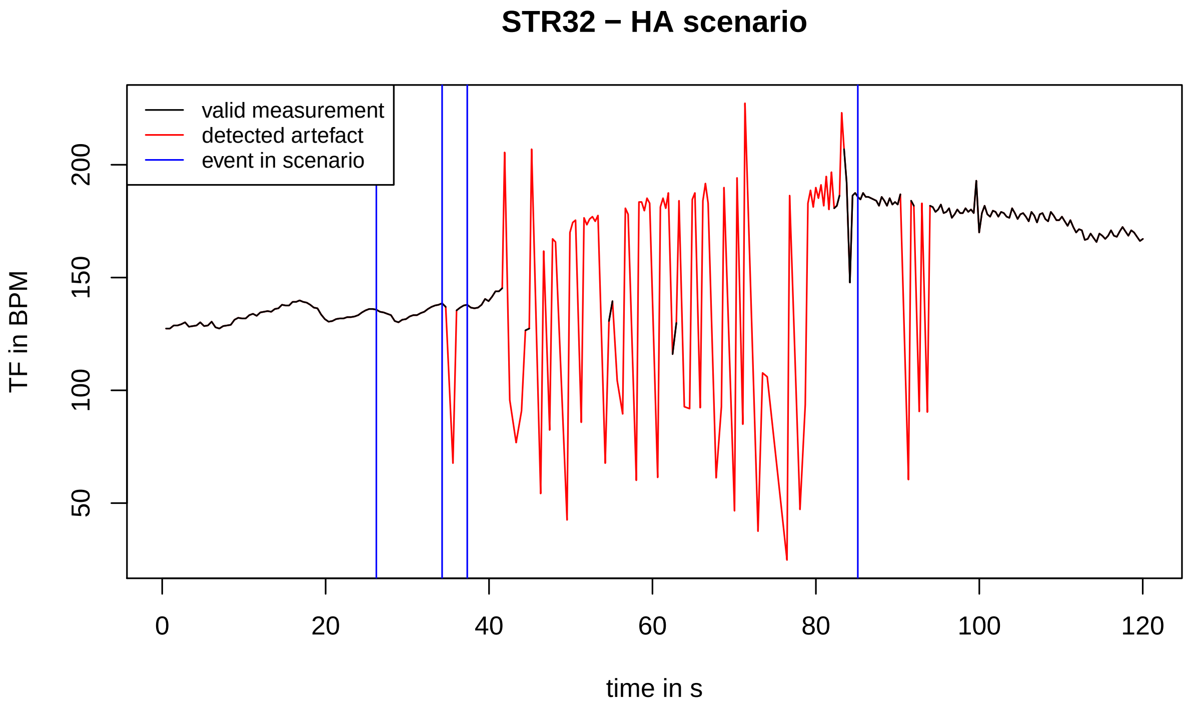
STR32 − HA scenario
TF in BPM
200
150
100
50
0
20
40
60
80
100
120
time in s
valid measurement
detected artefact
event in scenario

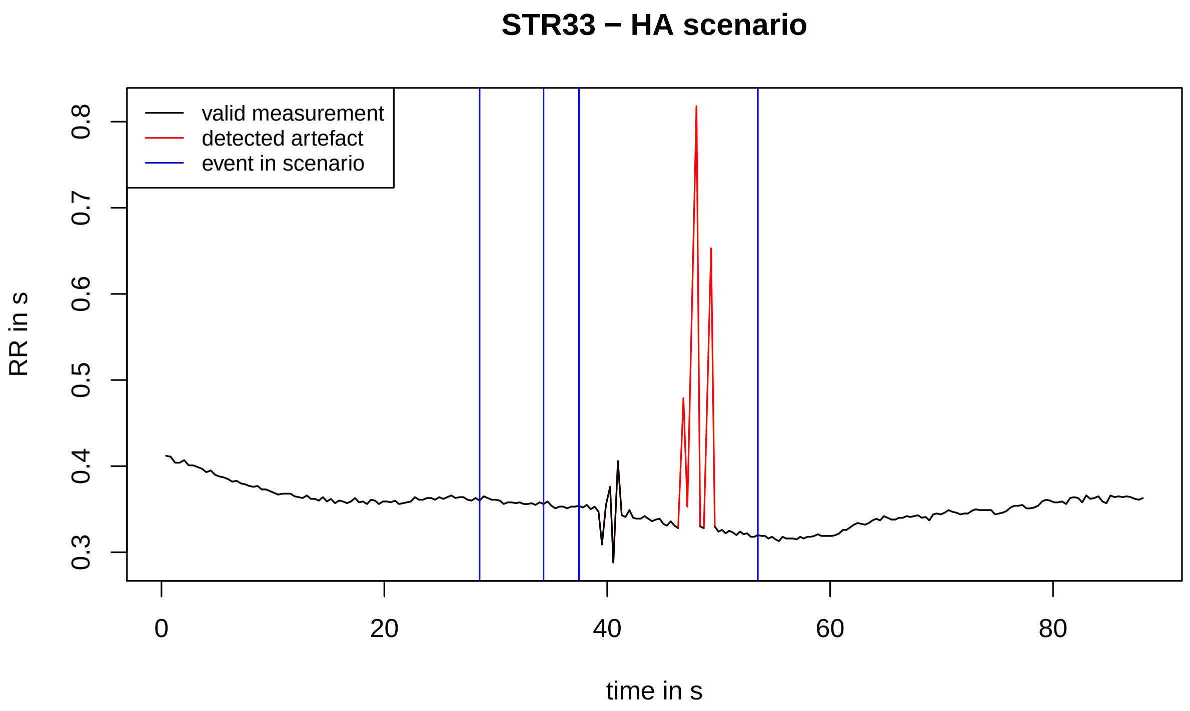
STR33 − HA scenario
valid measurement
detected artefact
event in scenario
RR in s
0.3
0.4
0.5
0.6
0.7
0.8
0
20
40
60
80
time in s

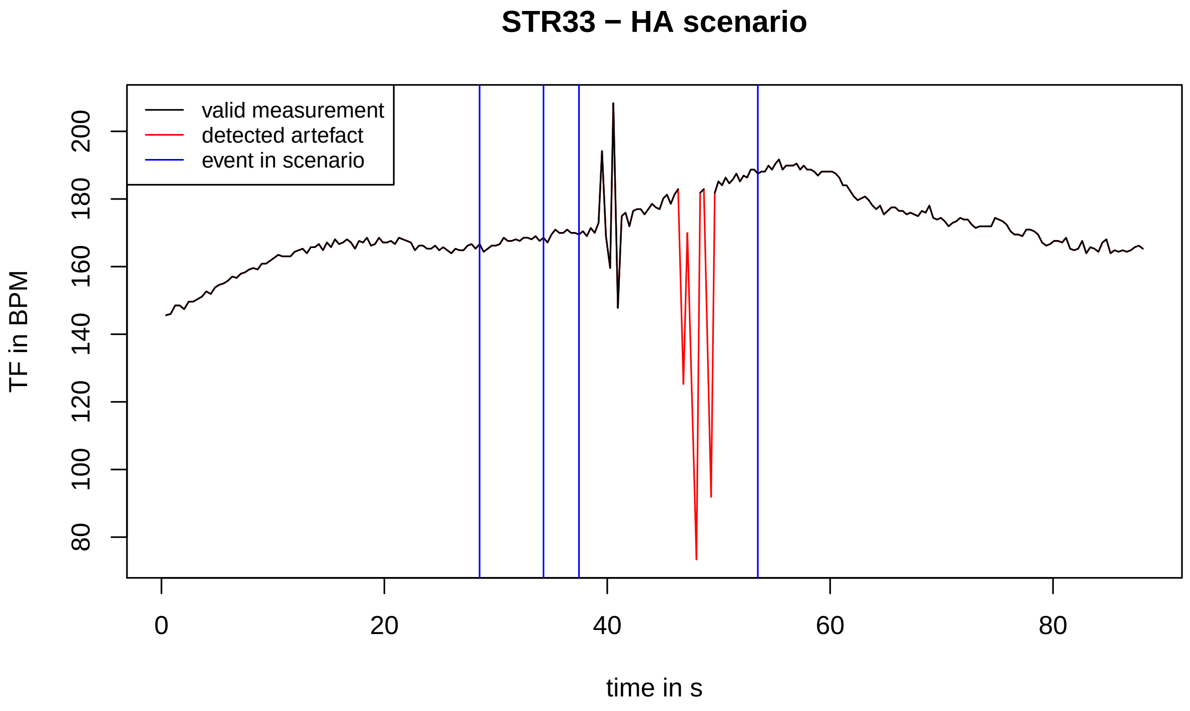
STR33 − HA scenario
valid measurement
detected artefact
event in scenario
TF in BPM
80
100
120
140
160
180
200
0
20
40
60
80
time in s

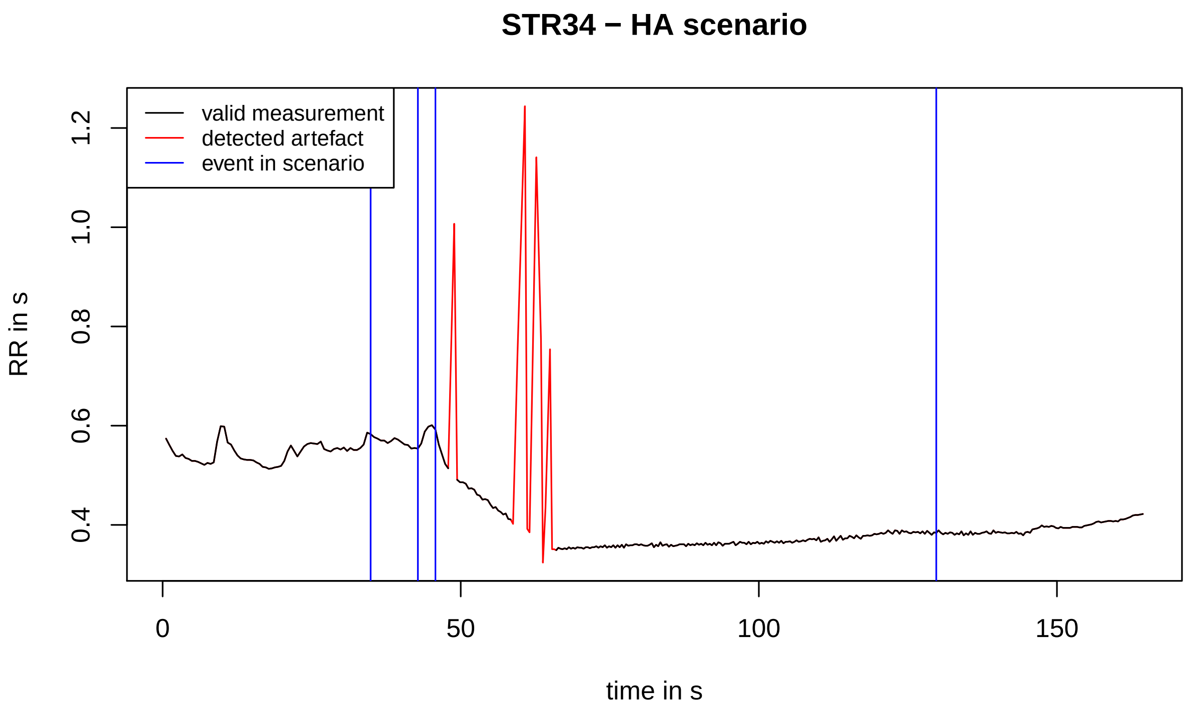
STR34 − HA scenario
valid measurement
detected artefact
event in scenario
RR in s
0.4
0.6
0.8
1.0
1.2
0
50
100
150
time in s

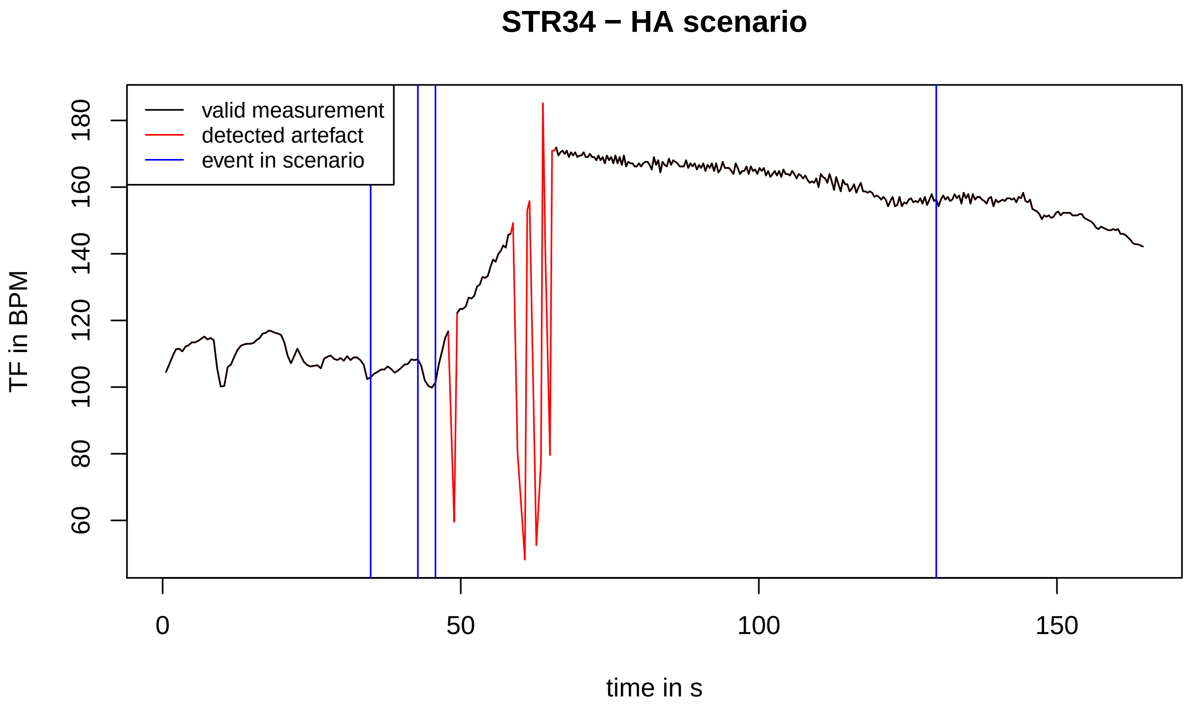
STR34 − HA scenario
valid measurement
detected artefact
event in scenario
TF in BPM
60
80
100
120
140
160
180
0
50
100
150
time in s

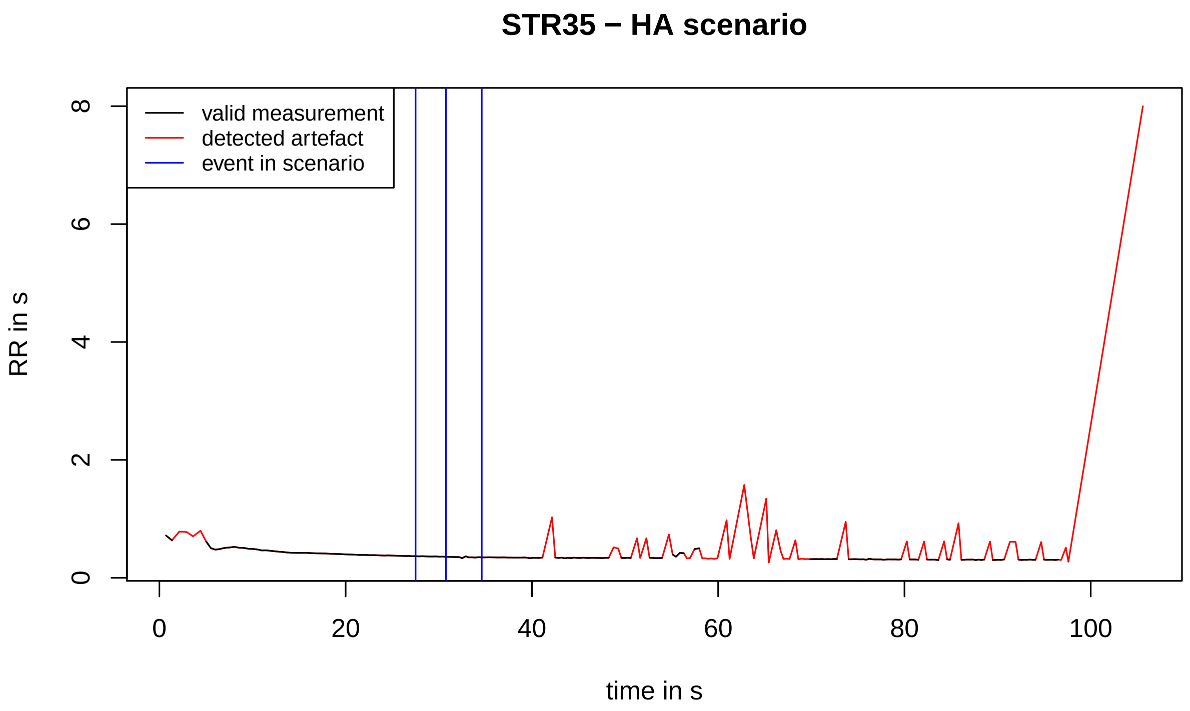
STR35 − HA scenario
valid measurement
detected artefact
event in scenario
RR in s
0
2
4
6
8
0
20
40
60
80
100
time in s

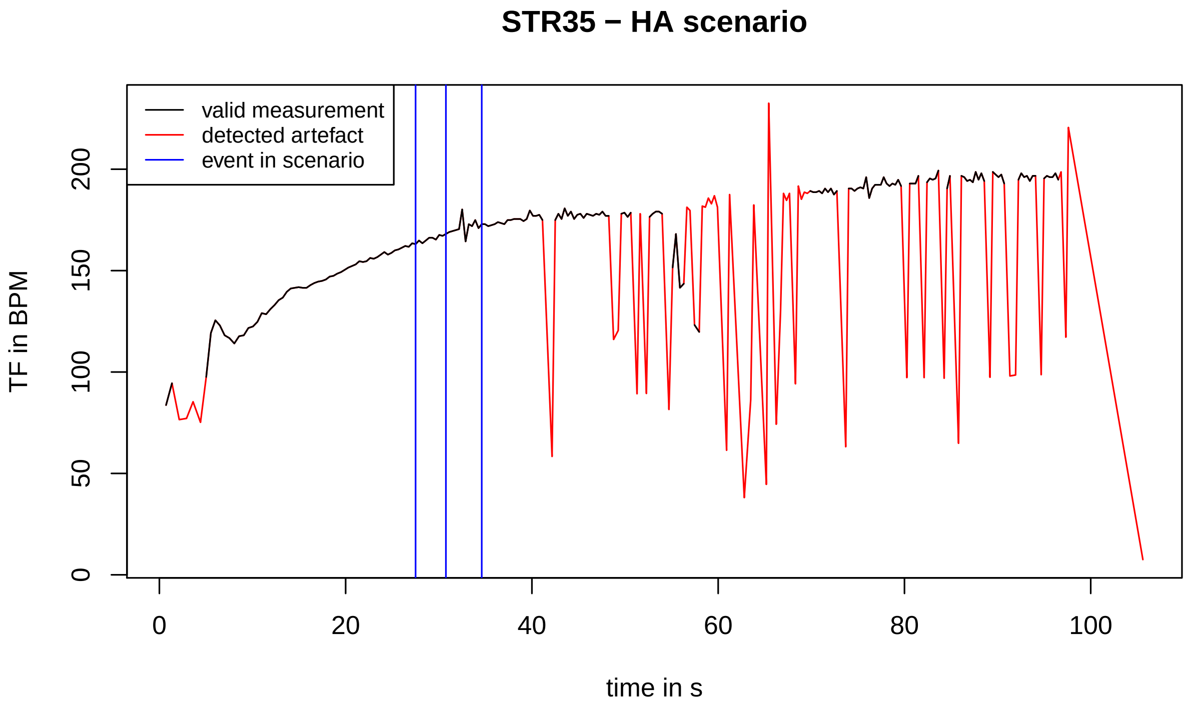
STR35 − HA scenario
valid measurement
detected artefact
event in scenario
TF in BPM
0
50
100
150
200
0
20
40
60
80
100
time in s

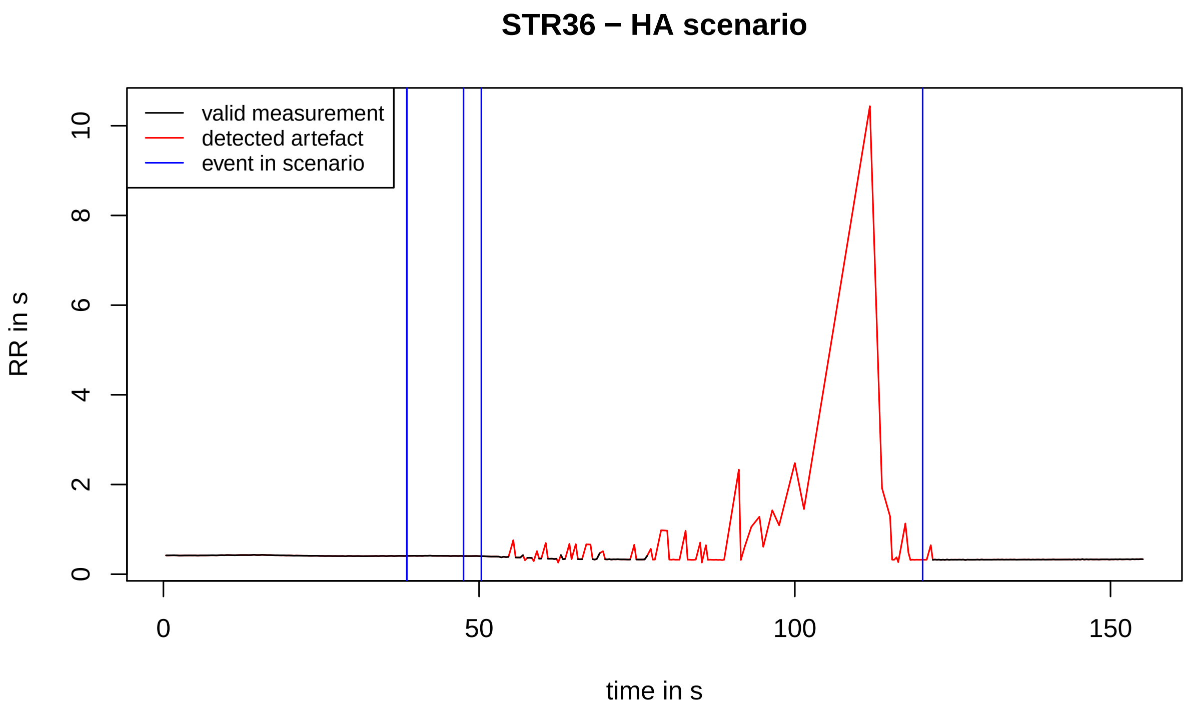
STR36 − HA scenario
valid measurement
detected artefact
event in scenario
RR in s
0
2
4
6
8
10
0
50
100
150
time in s

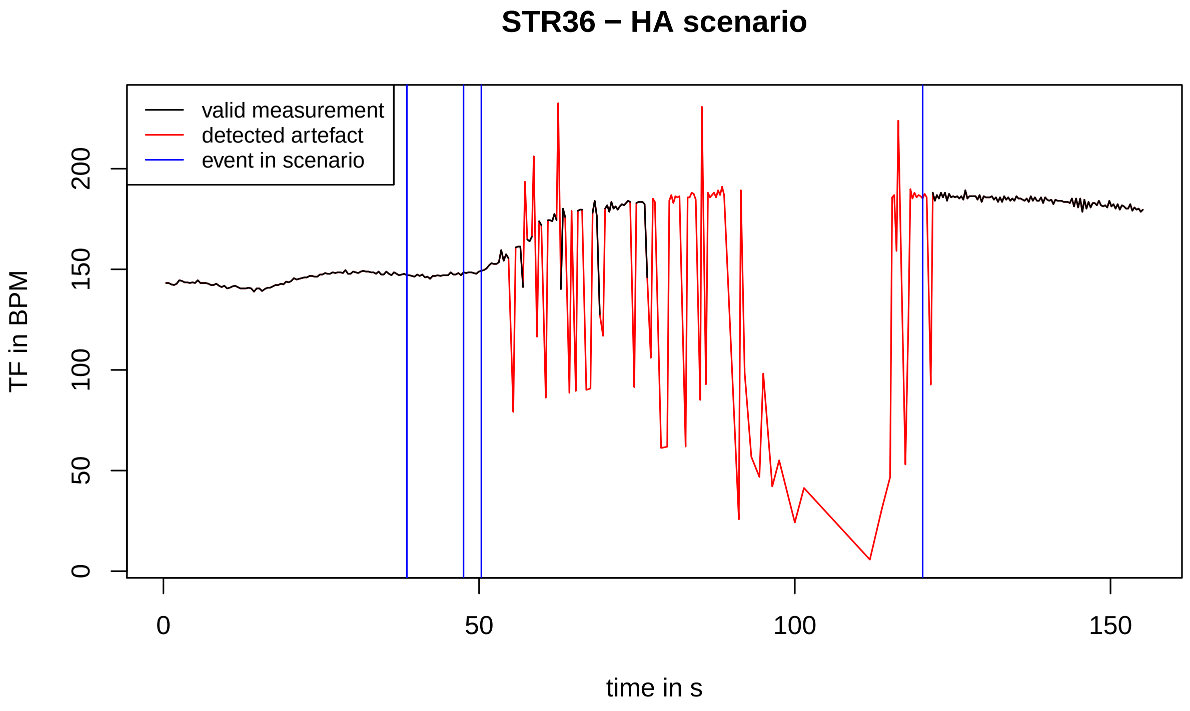
STR36 − HA scenario
valid measurement
detected artefact
event in scenario
TF in BPM
0
50
100
150
200
0
50
100
150
time in s

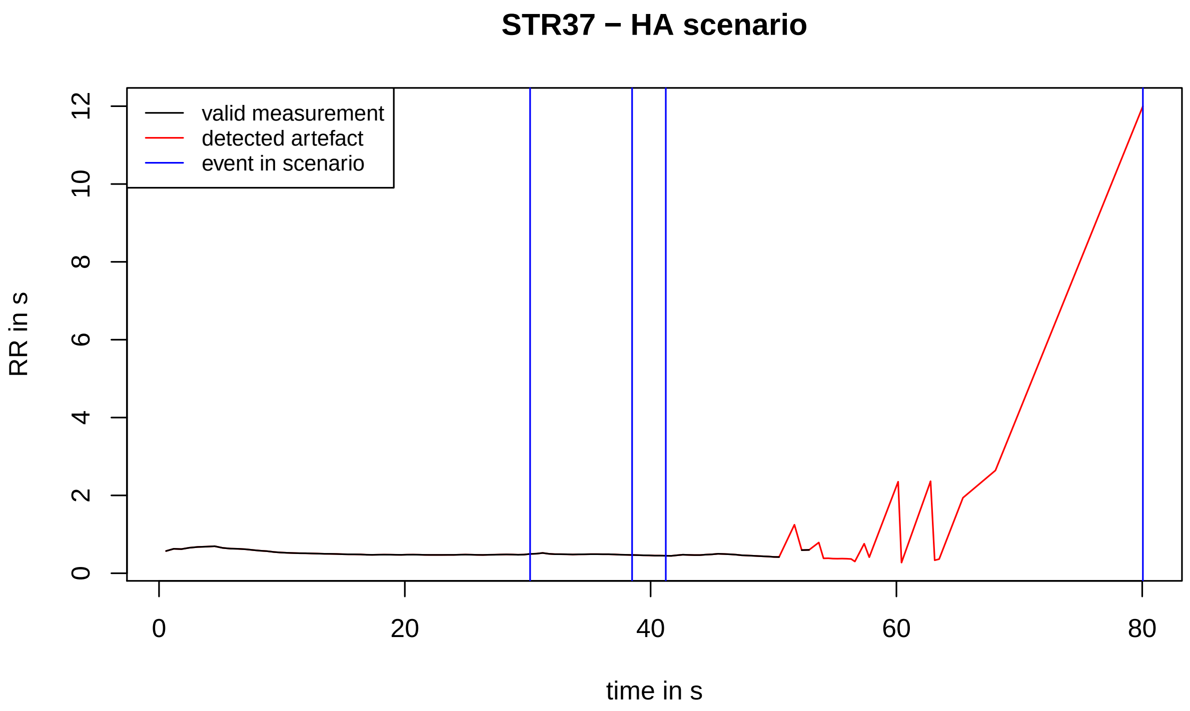
STR37 − HA scenario
valid measurement
detected artefact
event in scenario
RR in s
time in s

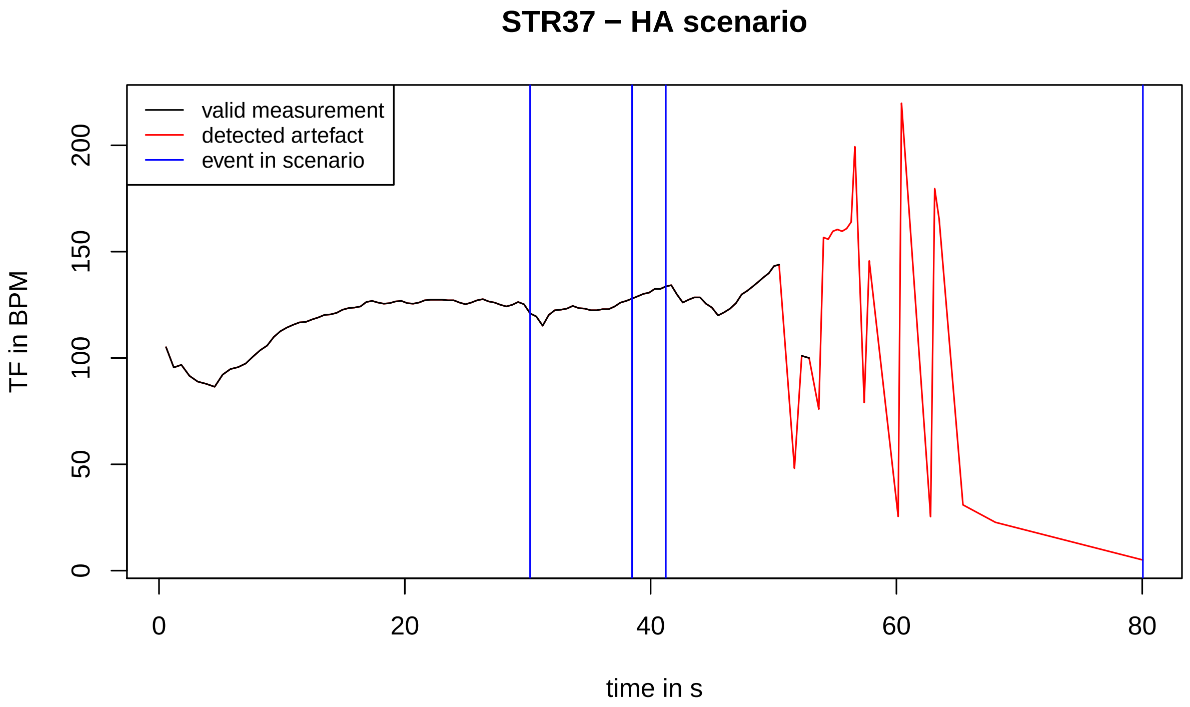
STR37 − HA scenario
valid measurement
detected artefact
event in scenario
TF in BPM
time in s

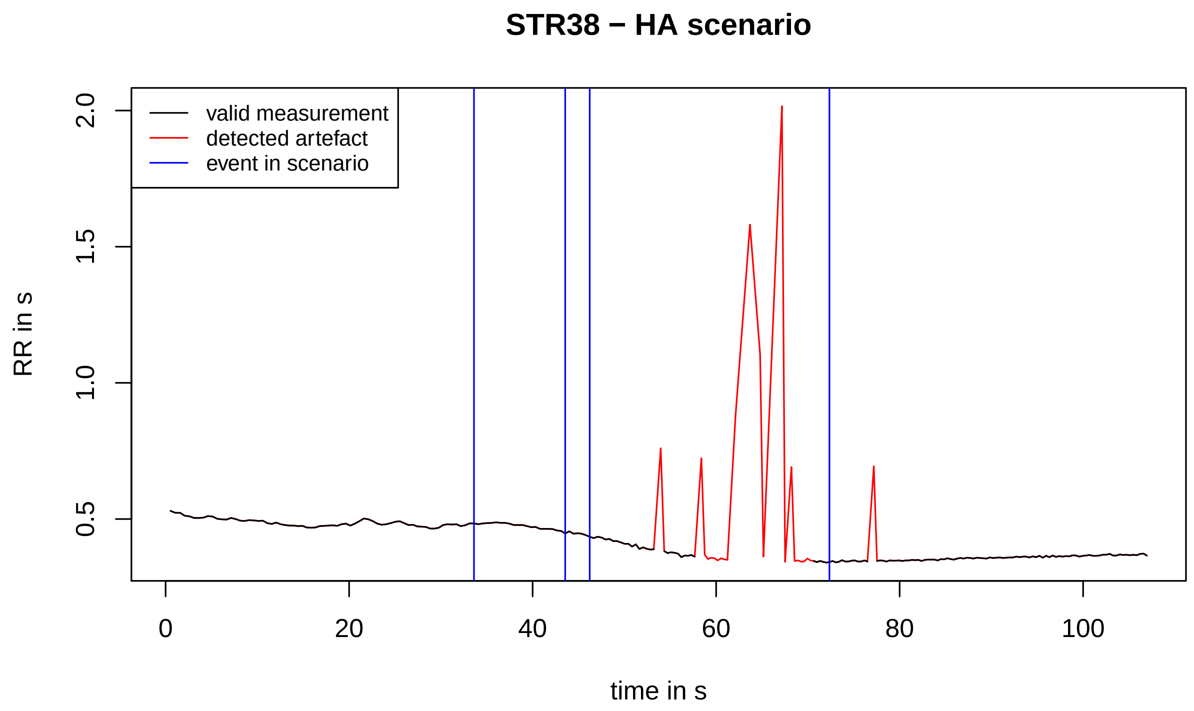
STR38 − HA scenario
valid measurement
detected artefact
event in scenario
RR in s
2.0
1.5
1.0
0.5
0
20
40
60
80
100
time in s

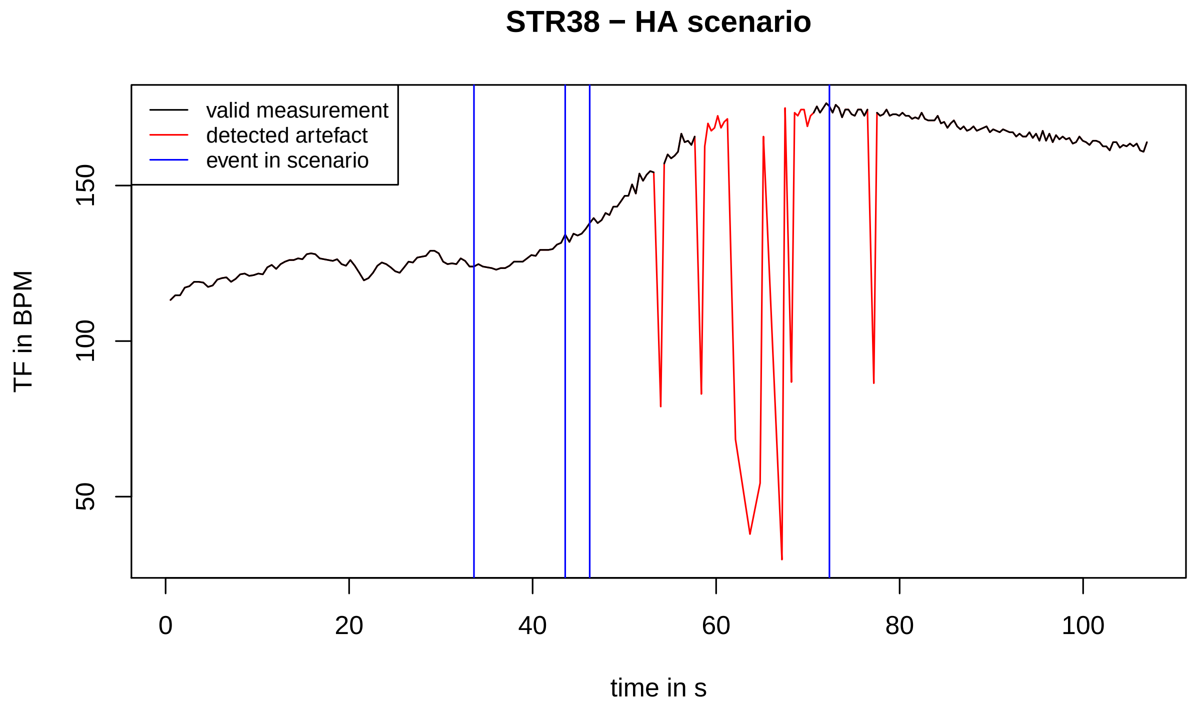
STR38 − HA scenario
valid measurement
detected artefact
event in scenario
TF in BPM
150
100
50
0
20
40
60
80
100
time in s

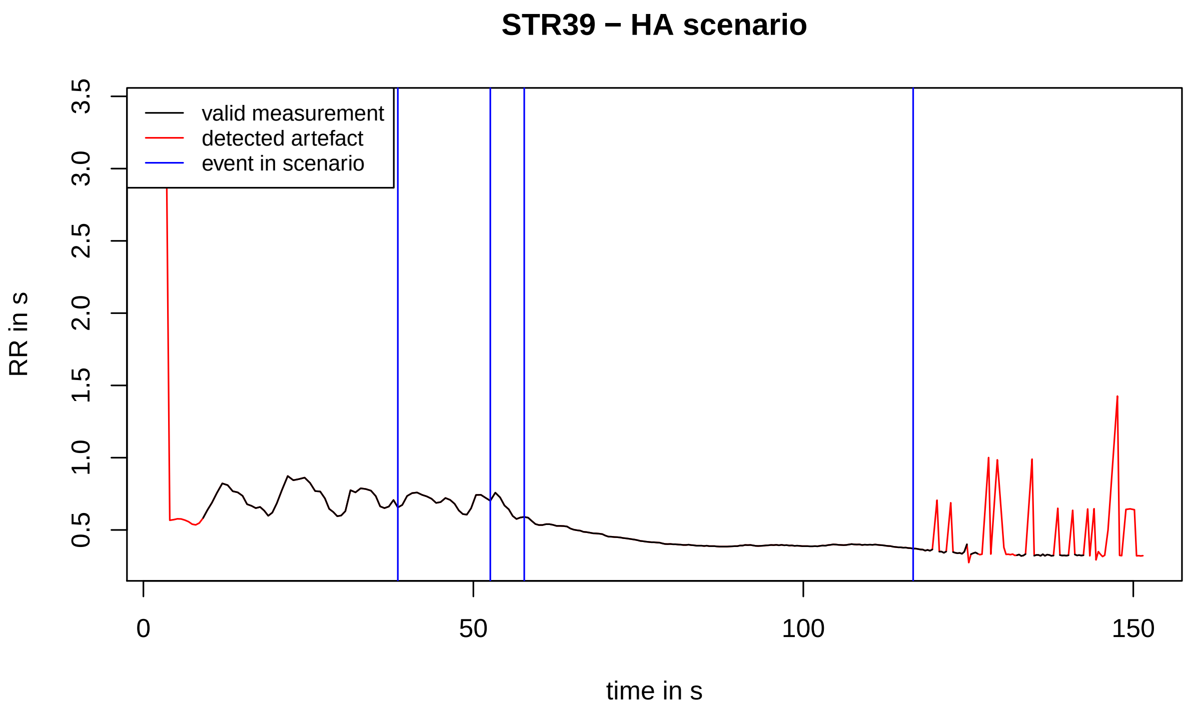
STR39 − HA scenario
valid measurement
detected artefact
event in scenario
RR in s
3.5
3.0
2.5
2.0
1.5
1.0
0.5
0
50
100
150
time in s

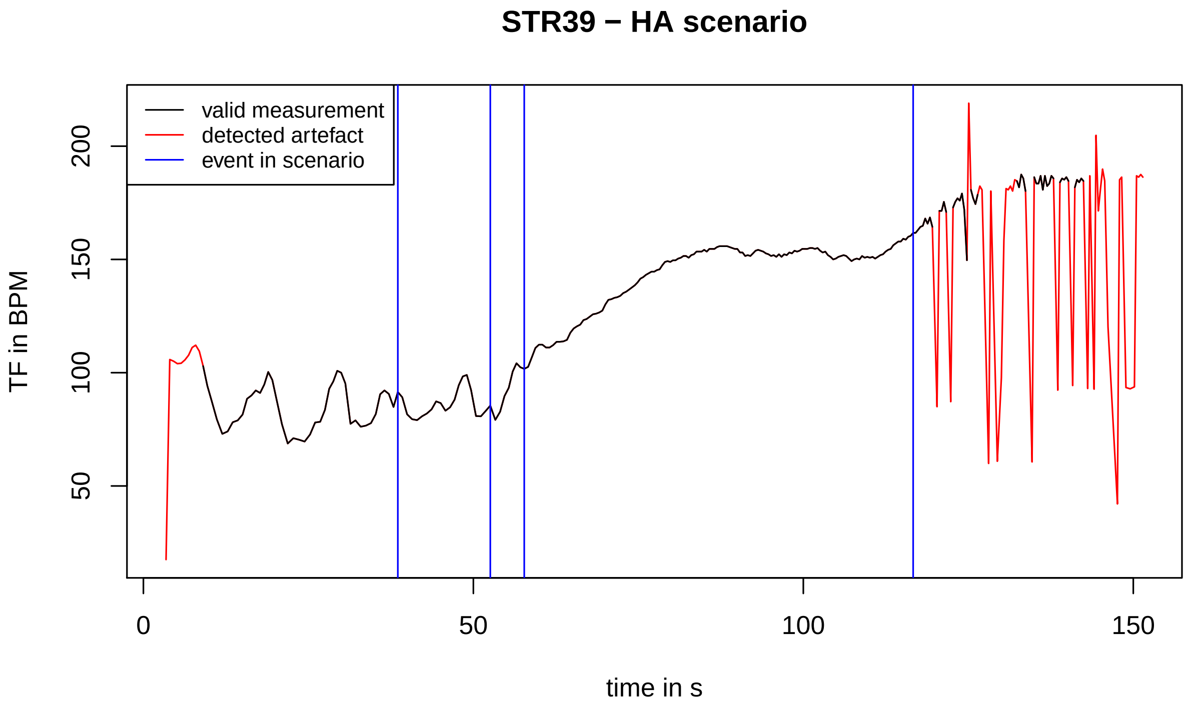
STR39 − HA scenario
valid measurement
detected artefact
event in scenario
TF in BPM
200
150
100
50
0
50
100
150
time in s

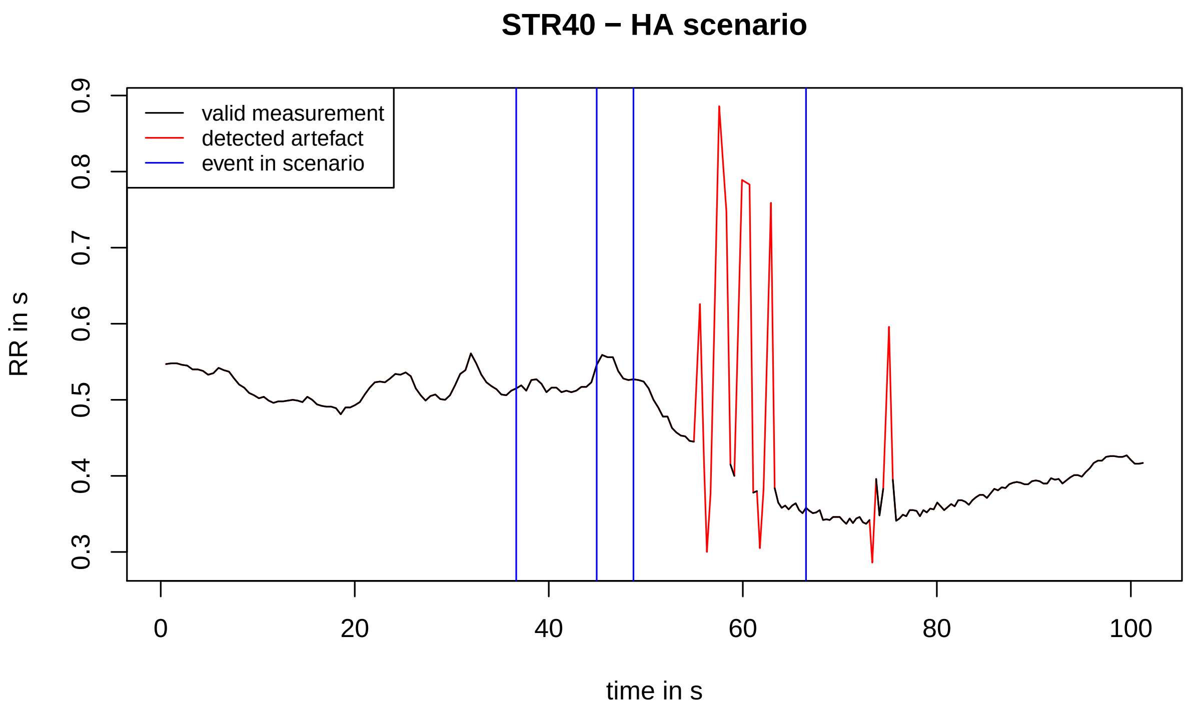
STR40 − HA scenario
valid measurement
detected artefact
event in scenario
RR in s
0.3
0.4
0.5
0.6
0.7
0.8
0.9
0
20
40
60
80
100
time in s

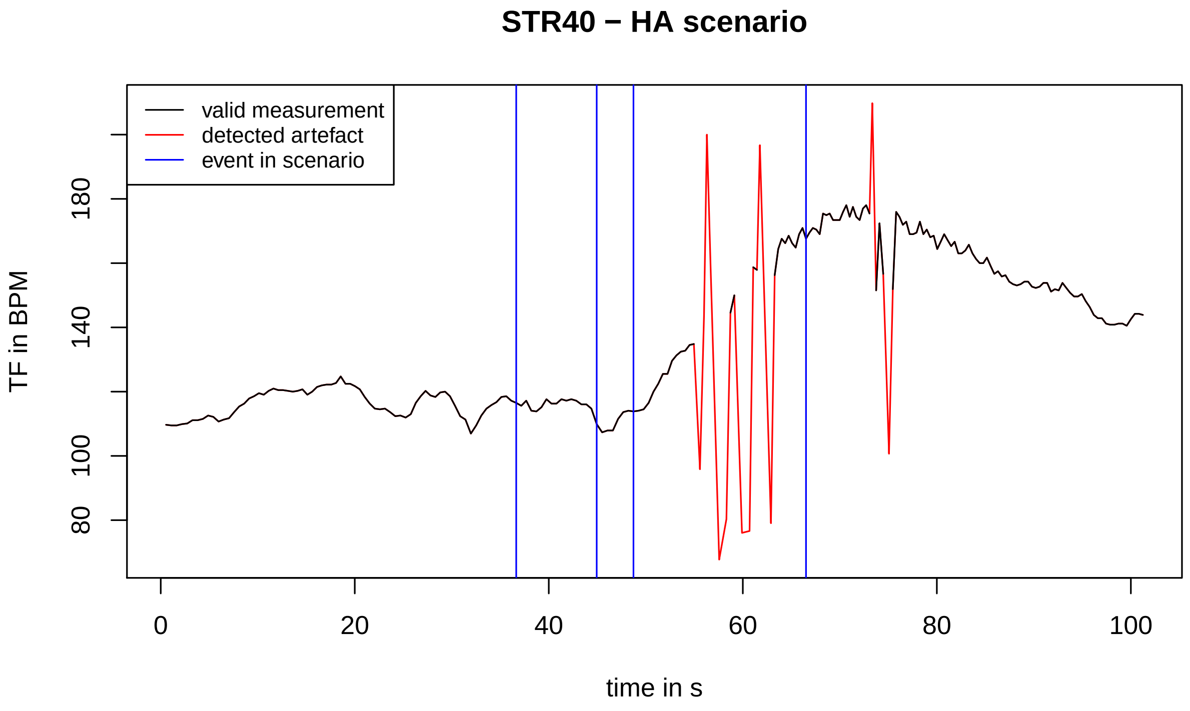
STR40 − HA scenario
valid measurement
detected artefact
event in scenario
TF in BPM
80
100
140
180
0
20
40
60
80
100
time in s

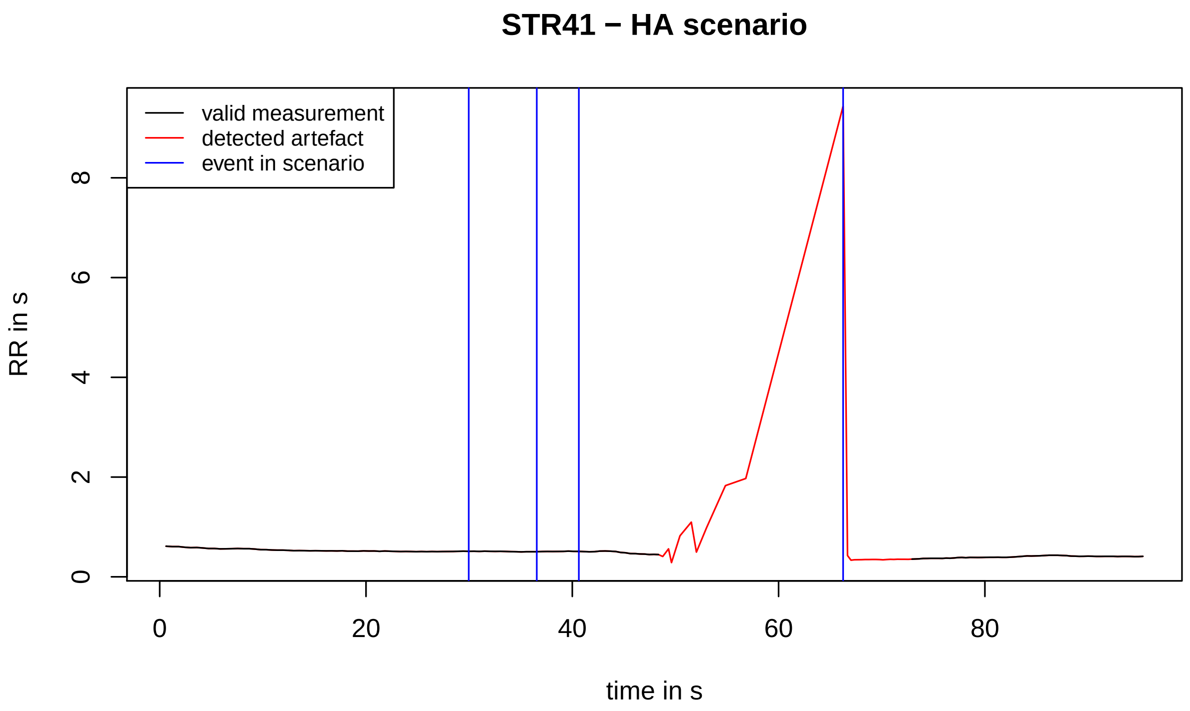
STR41 − HA scenario
valid measurement
detected artefact
event in scenario
RR in s
0
2
4
6
8
0
20
40
60
80
time in s

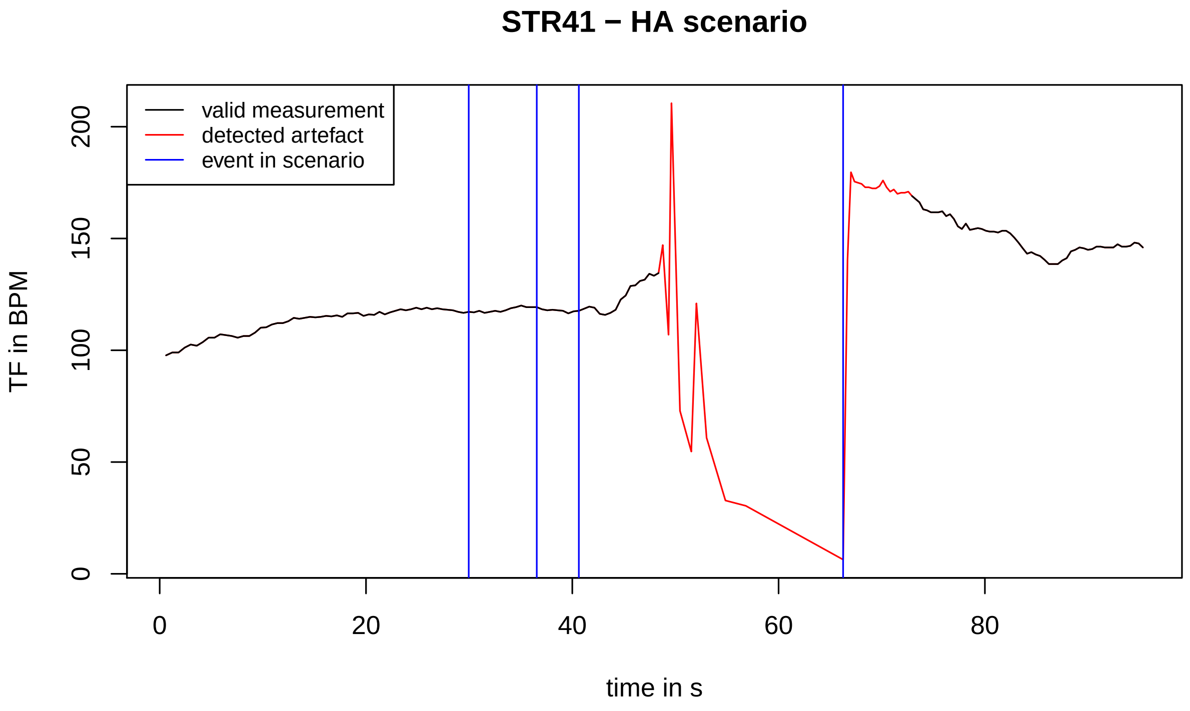
STR41 − HA scenario
valid measurement
detected artefact
event in scenario
TF in BPM
0
50
100
150
200
0
20
40
60
80
time in s

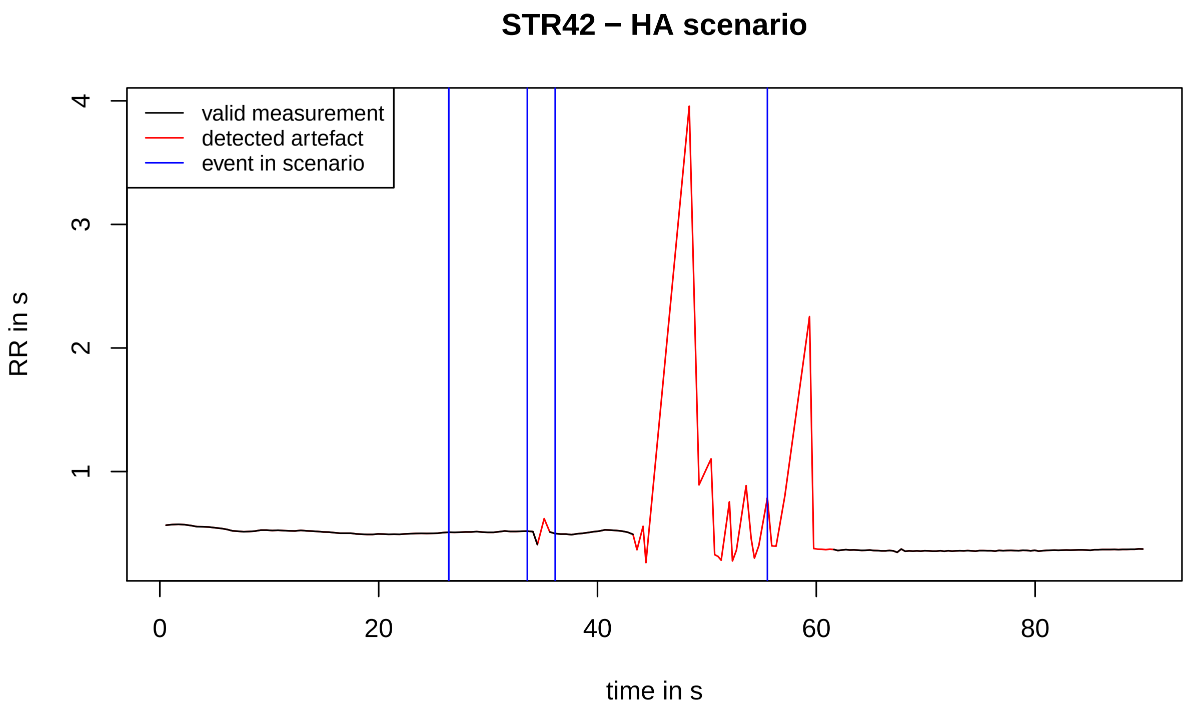
STR42 − HA scenario
valid measurement
detected artefact
event in scenario
RR in s
4
3
2
1
0
20
40
60
80
time in s

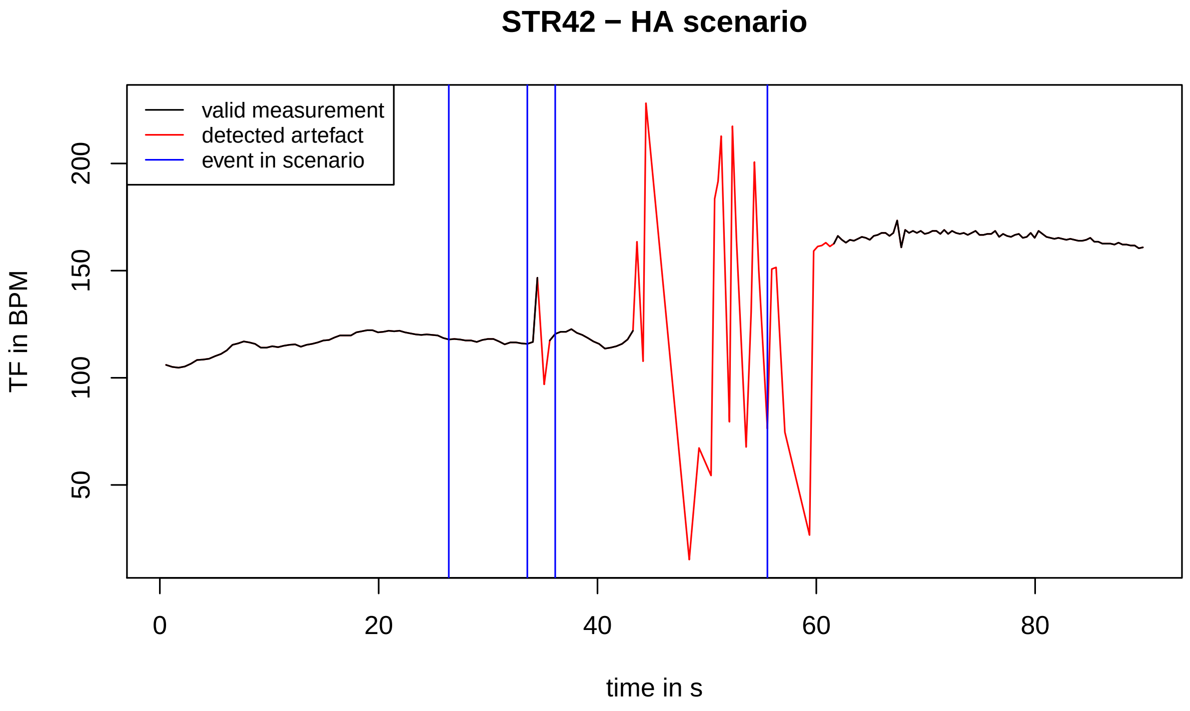
STR42 − HA scenario
valid measurement
detected artefact
event in scenario
TF in BPM
200
150
100
50
0
20
40
60
80
time in s

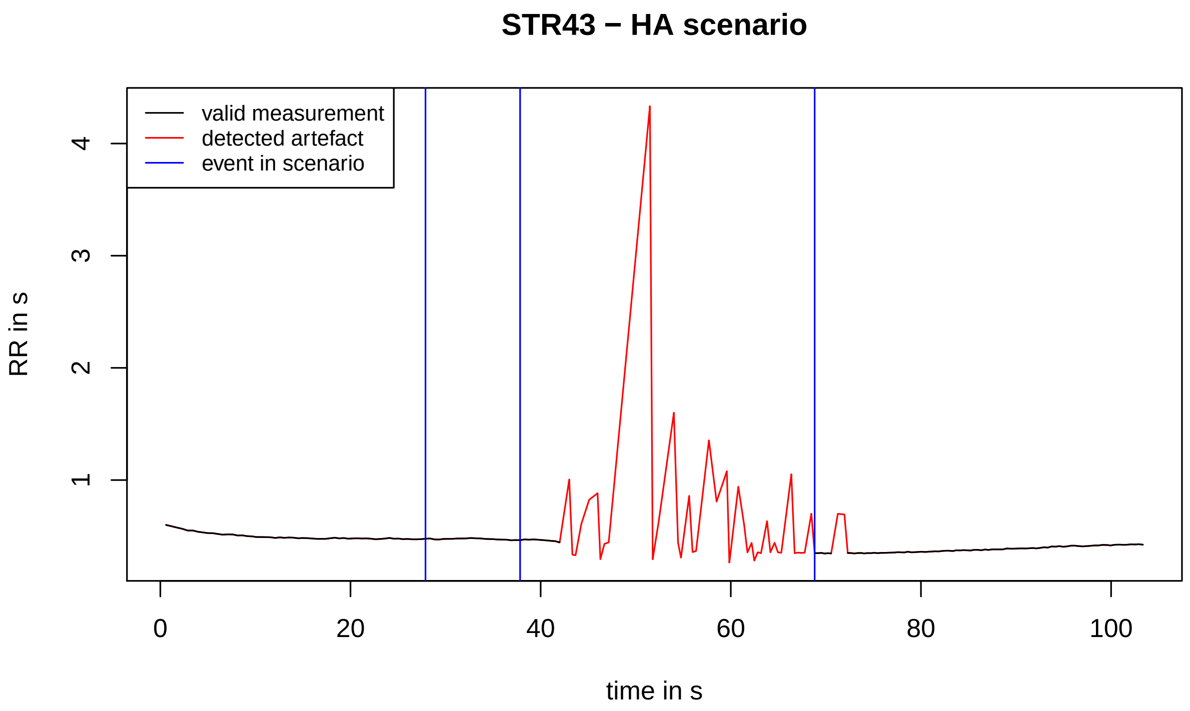
STR43 − HA scenario
valid measurement
detected artefact
event in scenario
RR in s
1
2
3
4
0
20
40
60
80
100
time in s

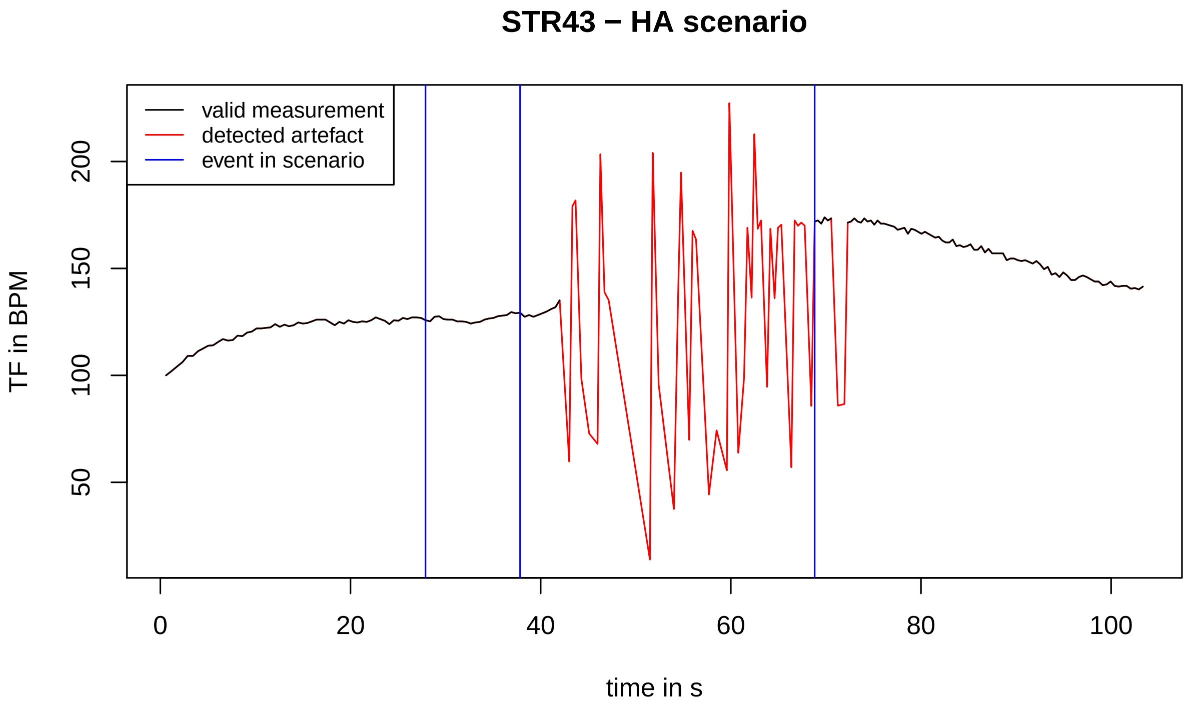
STR43 − HA scenario
valid measurement
detected artefact
event in scenario
TF in BPM
50
100
150
200
0
20
40
60
80
100
time in s

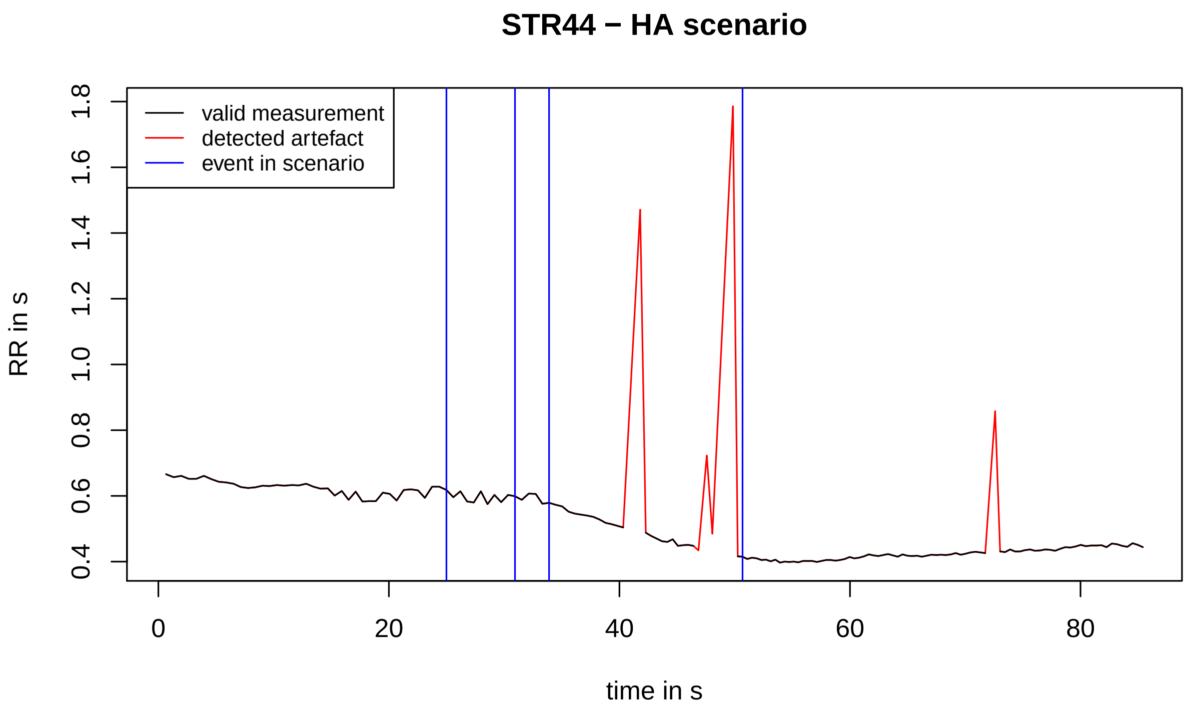
STR44 − HA scenario
valid measurement
detected artefact
event in scenario
RR in s
0.4
0.6
0.8
1.0
1.2
1.4
1.6
1.8
0
20
40
60
80
time in s

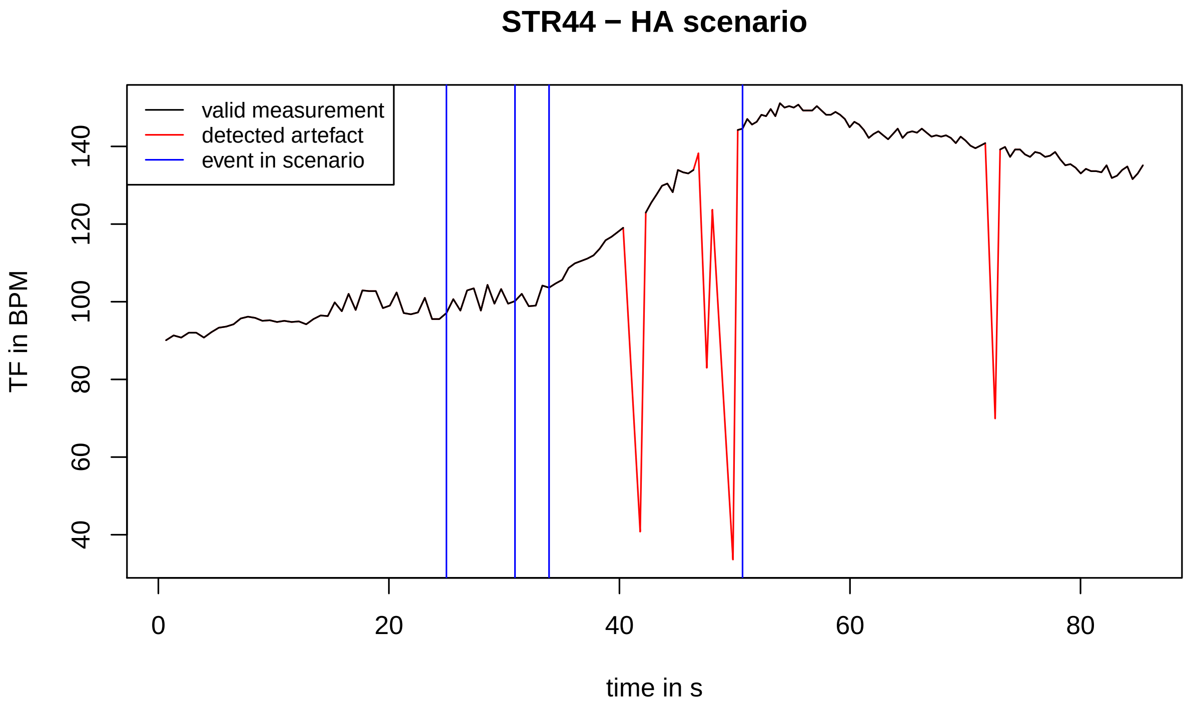
STR44 − HA scenario
valid measurement
detected artefact
event in scenario
TF in BPM
40
60
80
100
120
140
0
20
40
60
80
time in s

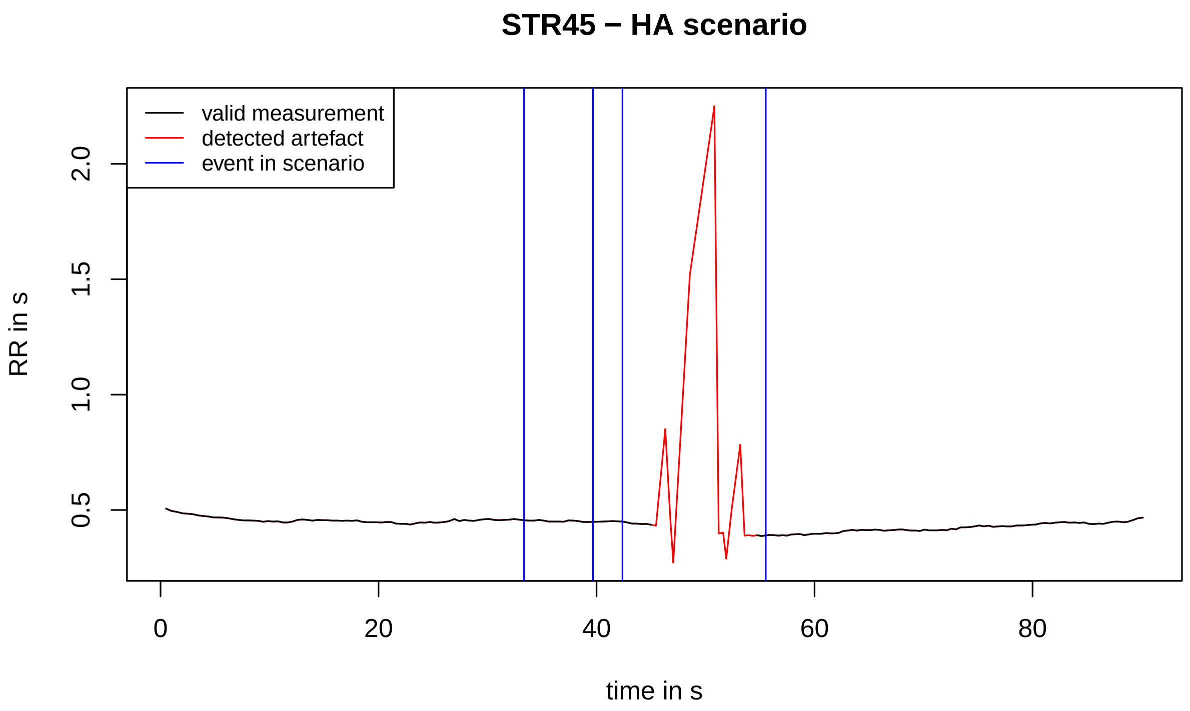
STR45 − HA scenario
valid measurement
detected artefact
event in scenario
RR in s
0.5
1.0
1.5
2.0
0
20
40
60
80
time in s

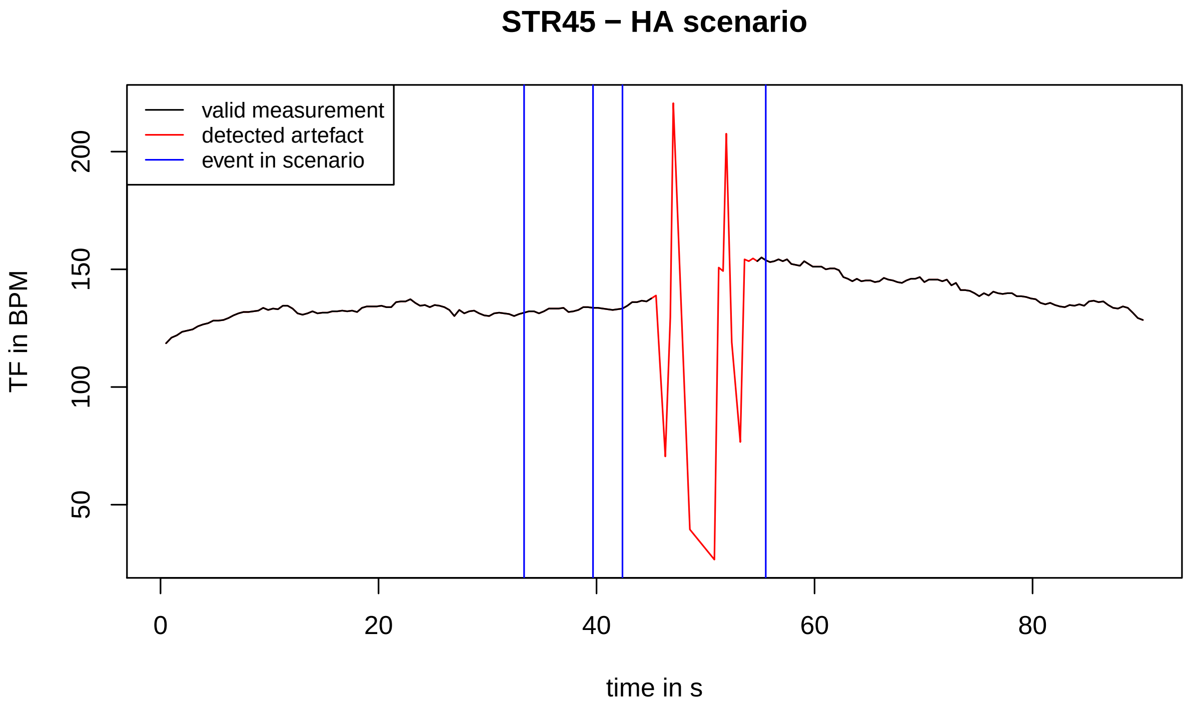
STR45 − HA scenario
valid measurement
detected artefact
event in scenario
TF in BPM
50
100
150
200
0
20
40
60
80
time in s

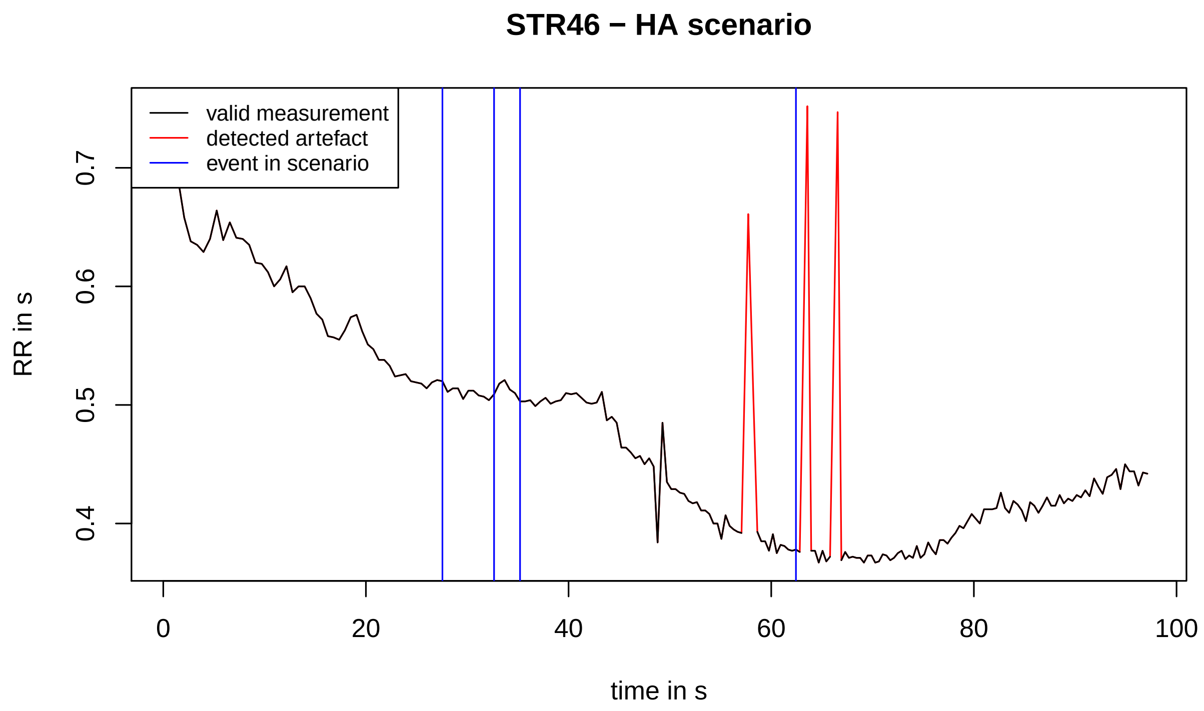
STR46 − HA scenario
valid measurement
detected artefact
event in scenario
RR in s
0.4
0.5
0.6
0.7
0
20
40
60
80
100
time in s

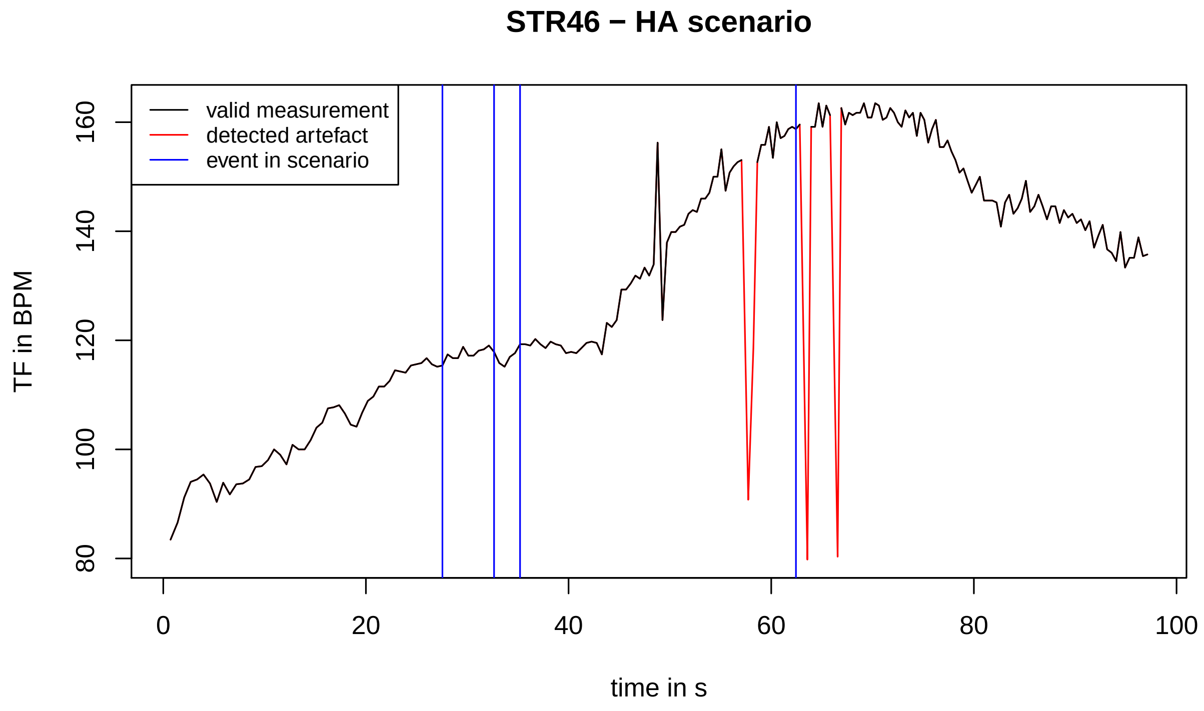
STR46 − HA scenario
valid measurement
detected artefact
event in scenario
TF in BPM
80
100
120
140
160
0
20
40
60
80
100
time in s

#### STR47 − HA scenario

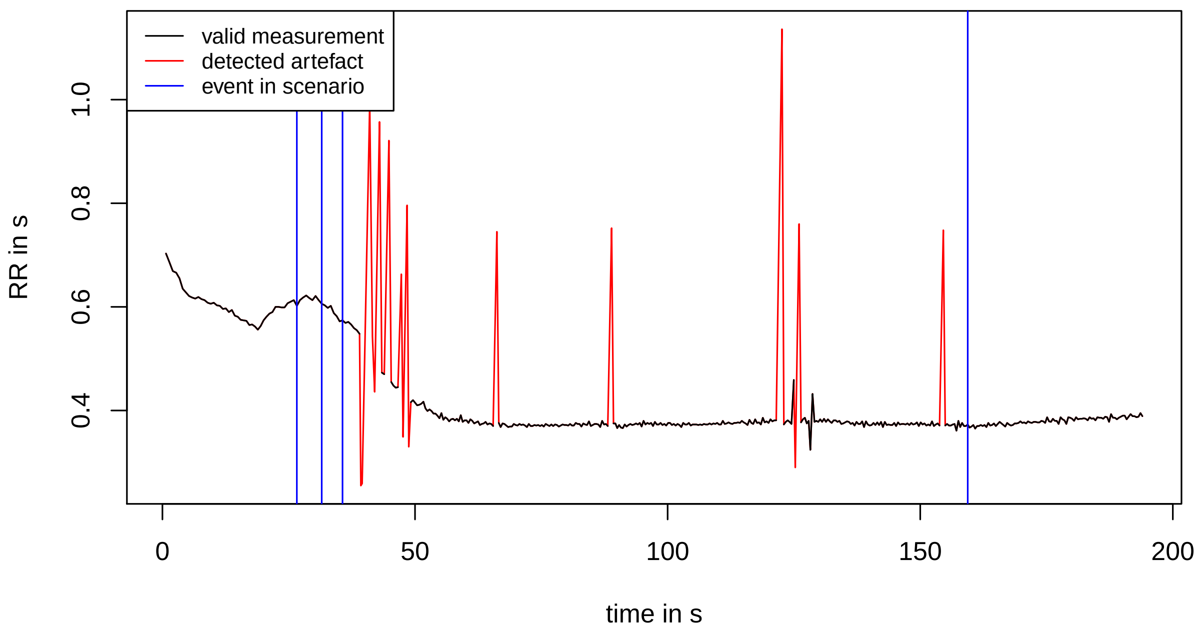


#### STR47 − HA scenario

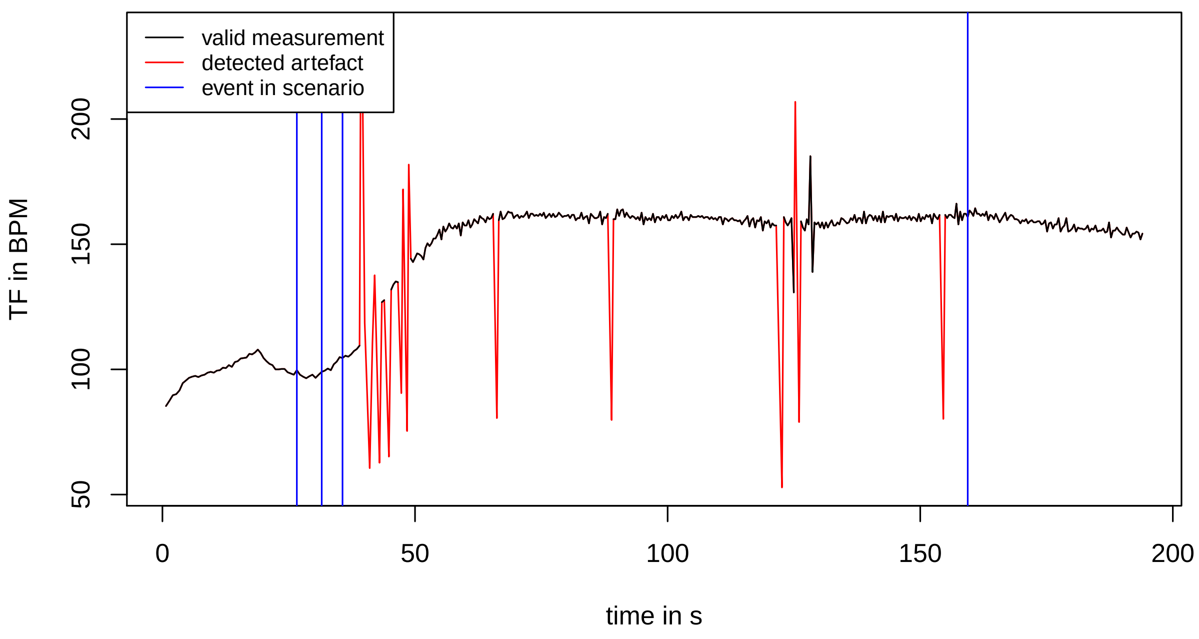

STR48 − HA scenario

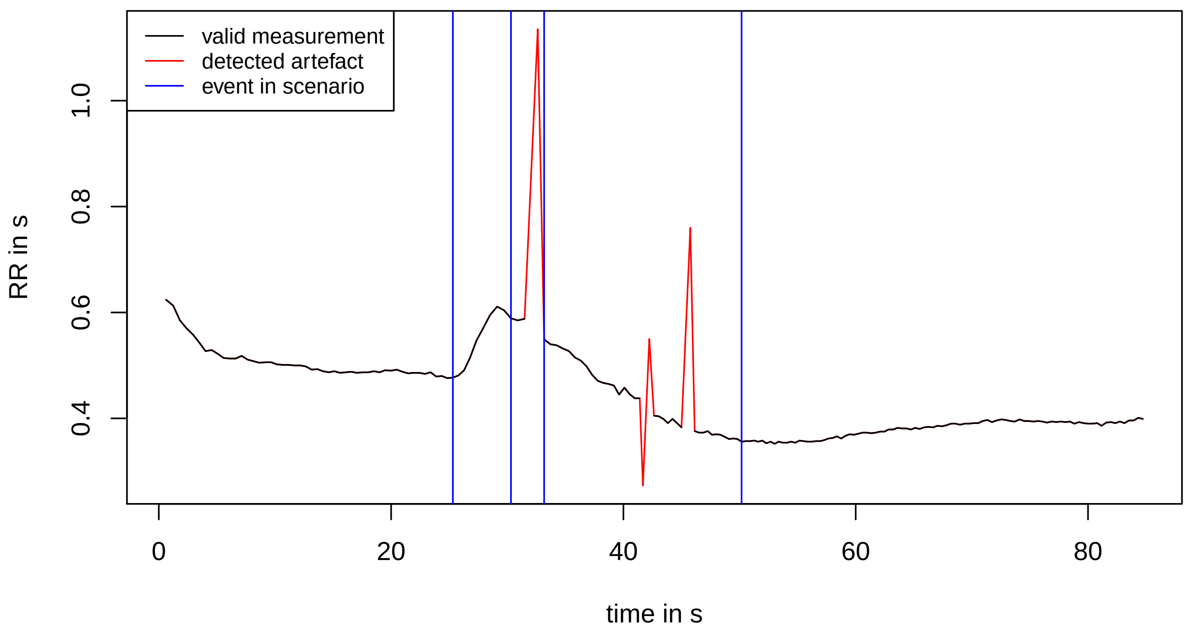


STR48 − HA scenario

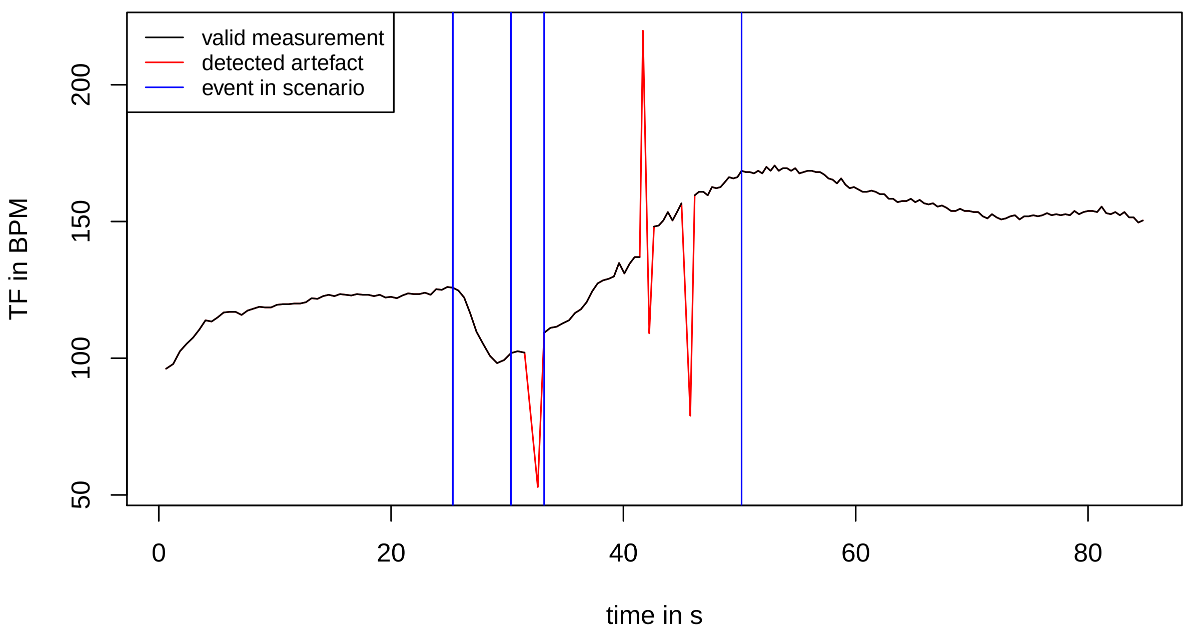

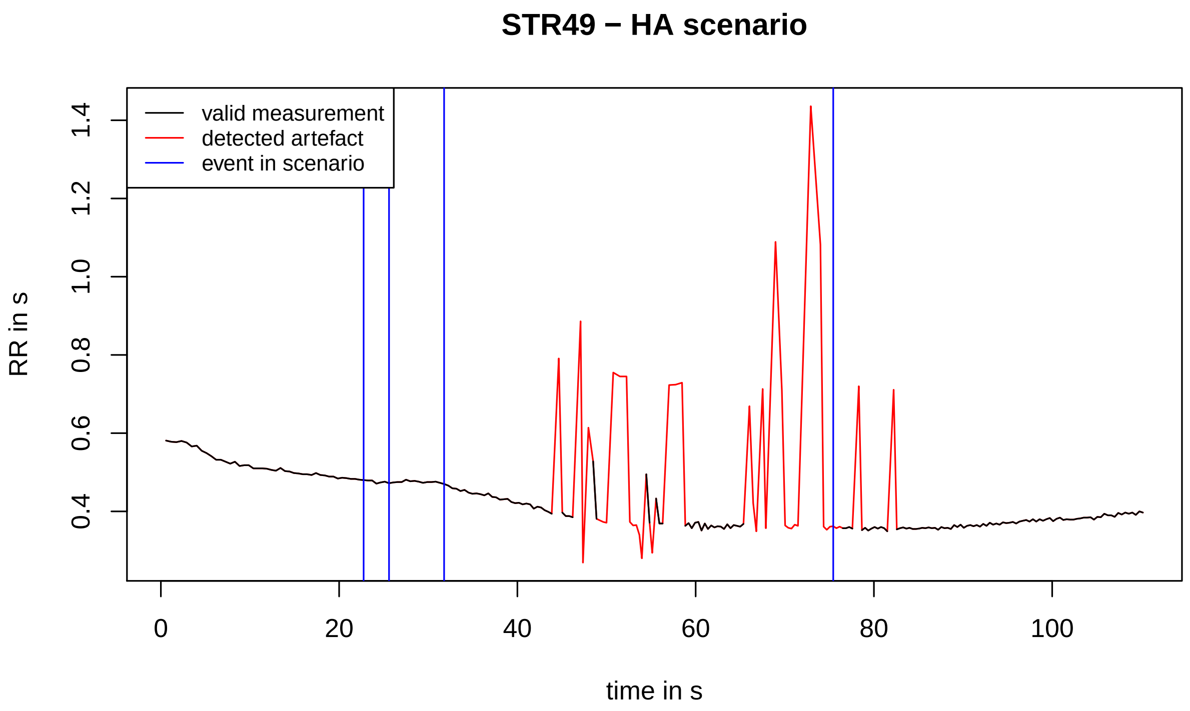

STR49 − HA scenario
valid measurement
detected artefact
event in scenario
RR in s
0.4
0.6
0.8
1.0
1.2
1.4
0
20
40
60
80
100
time in s


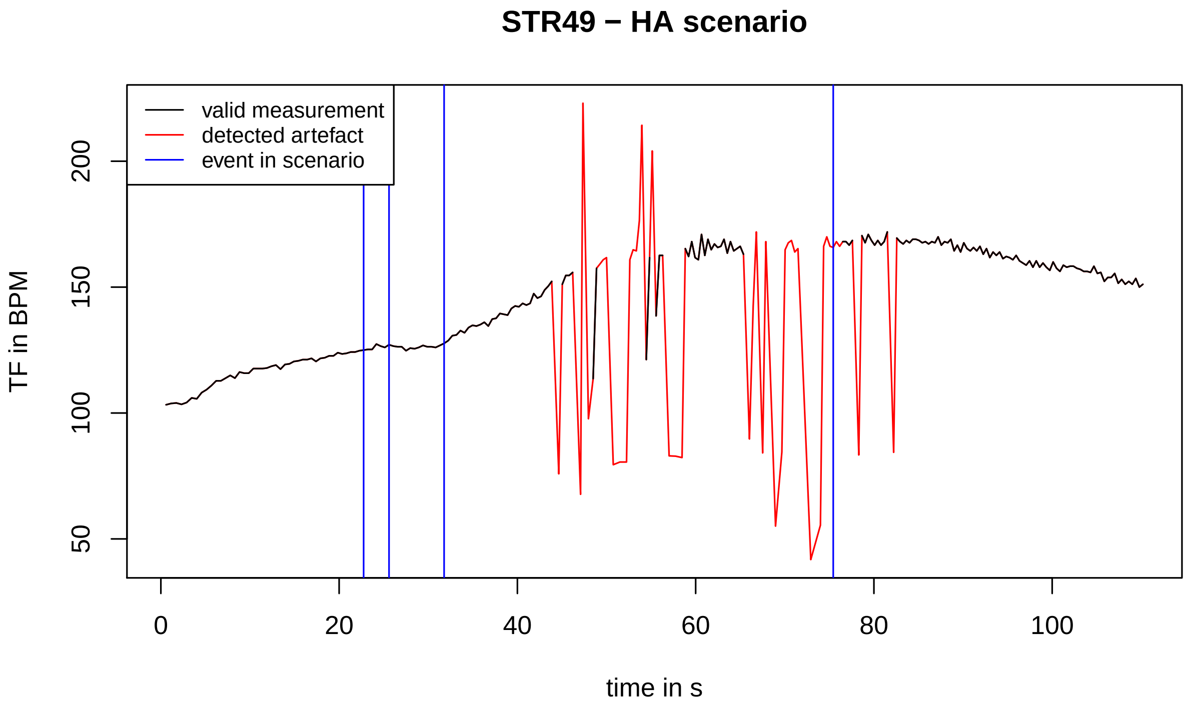

STR49 − HA scenario
valid measurement
detected artefact
event in scenario
TF in BPM
50
100
150
200
0
20
40
60
80
100
time in s

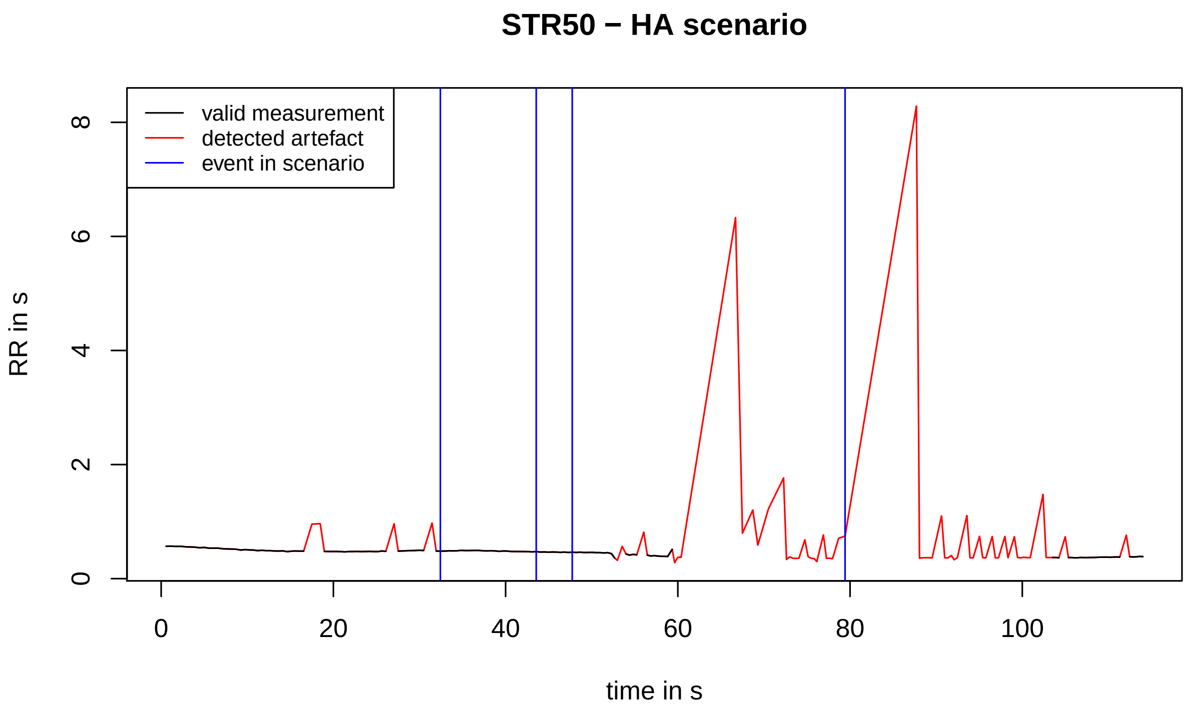
STR50 − HA scenario
valid measurement
detected artefact
event in scenario
RR in s
0
2
4
6
8
0
20
40
60
80
100
time in s

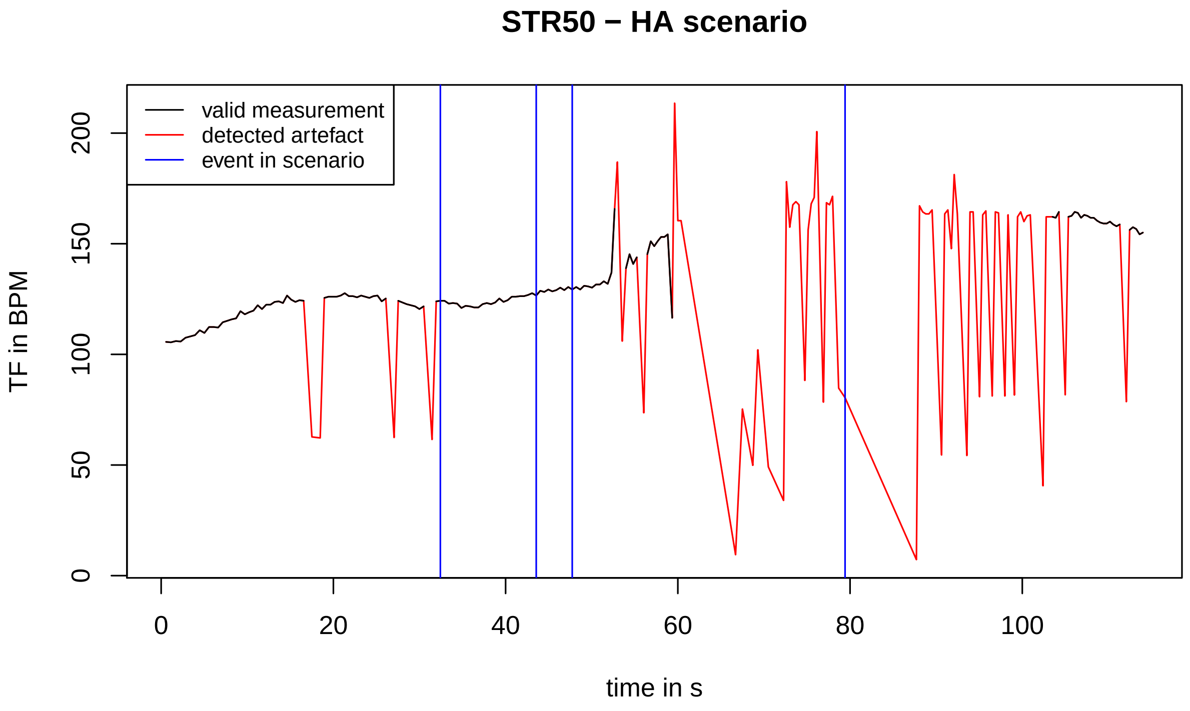
STR50 − HA scenario
valid measurement
detected artefact
event in scenario
TF in BPM
0
50
100
150
200
0
20
40
60
80
100
time in s

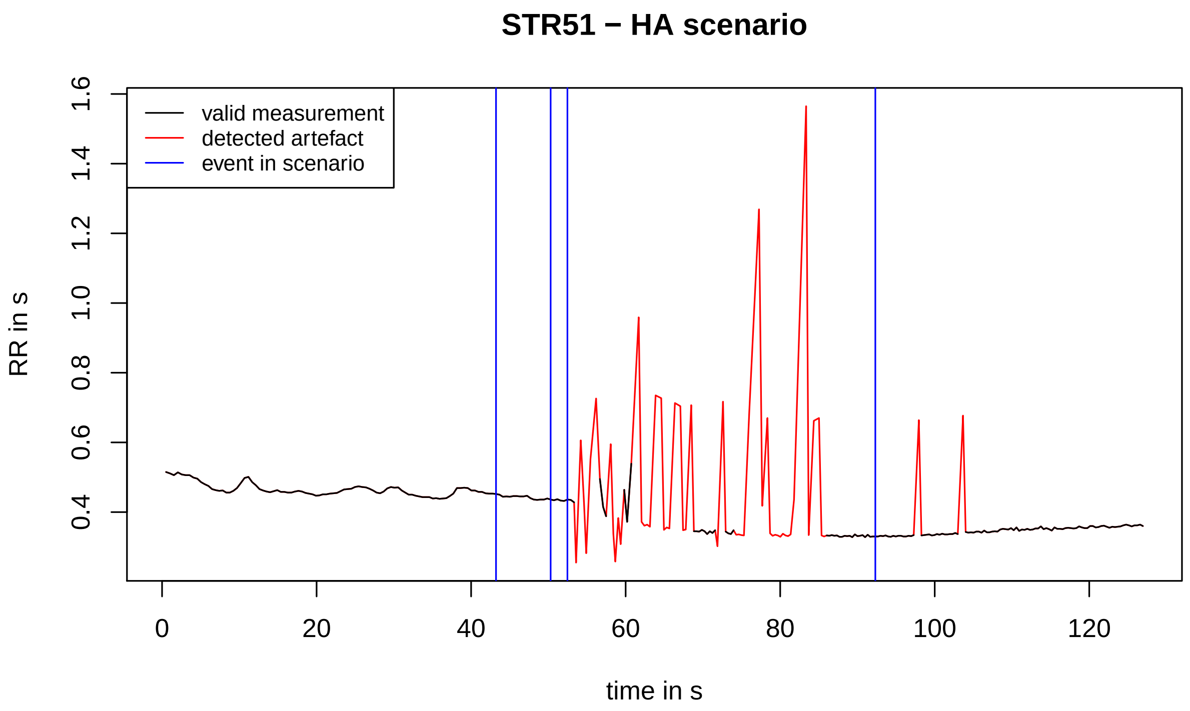
STR51 − HA scenario
valid measurement
detected artefact
event in scenario
RR in s
0.4
0.6
0.8
1.0
1.2
1.4
1.6
0
20
40
60
80
100
120
time in s

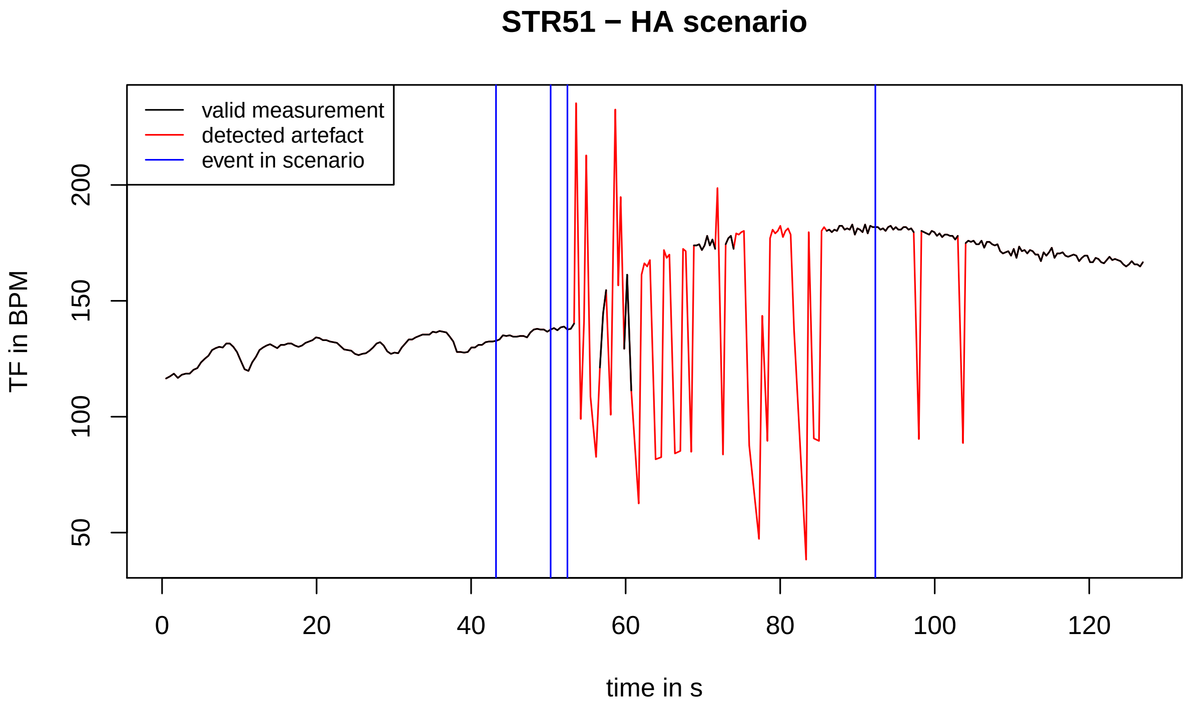
STR51 − HA scenario
valid measurement
detected artefact
event in scenario
TF in BPM
50
100
150
200
0
20
40
60
80
100
120
time in s

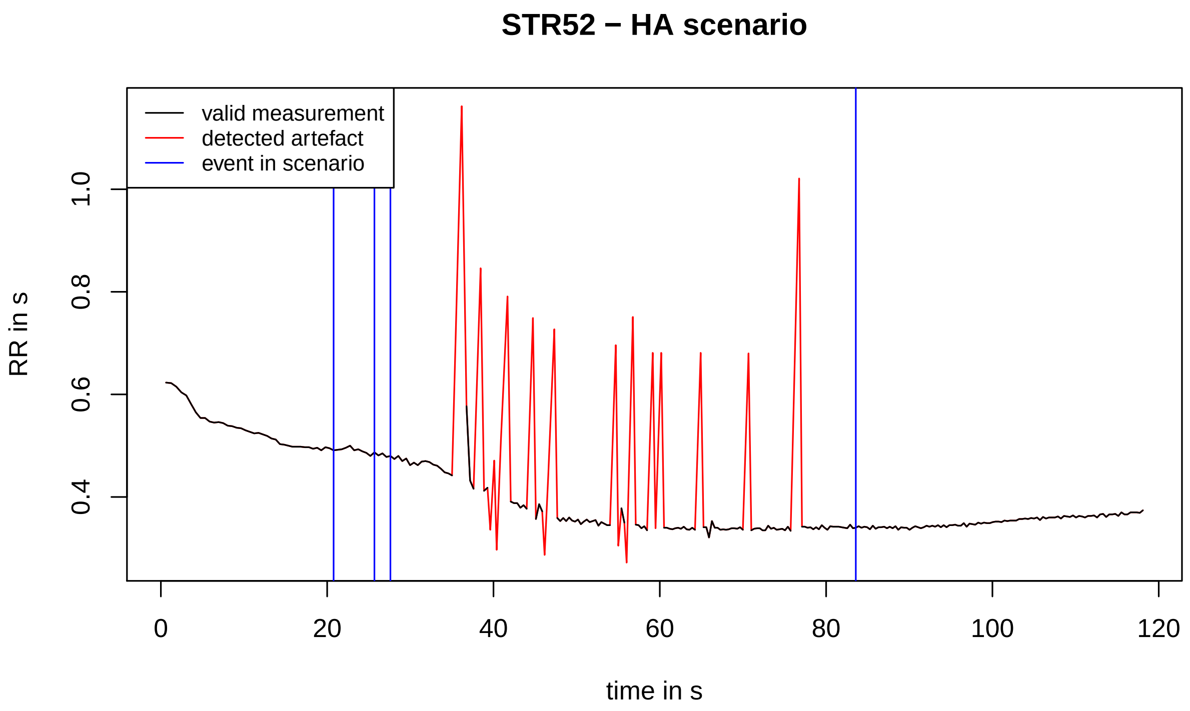
STR52 − HA scenario
valid measurement
detected artefact
event in scenario
RR in s
0.4
0.6
0.8
1.0
0
20
40
60
80
100
120
time in s

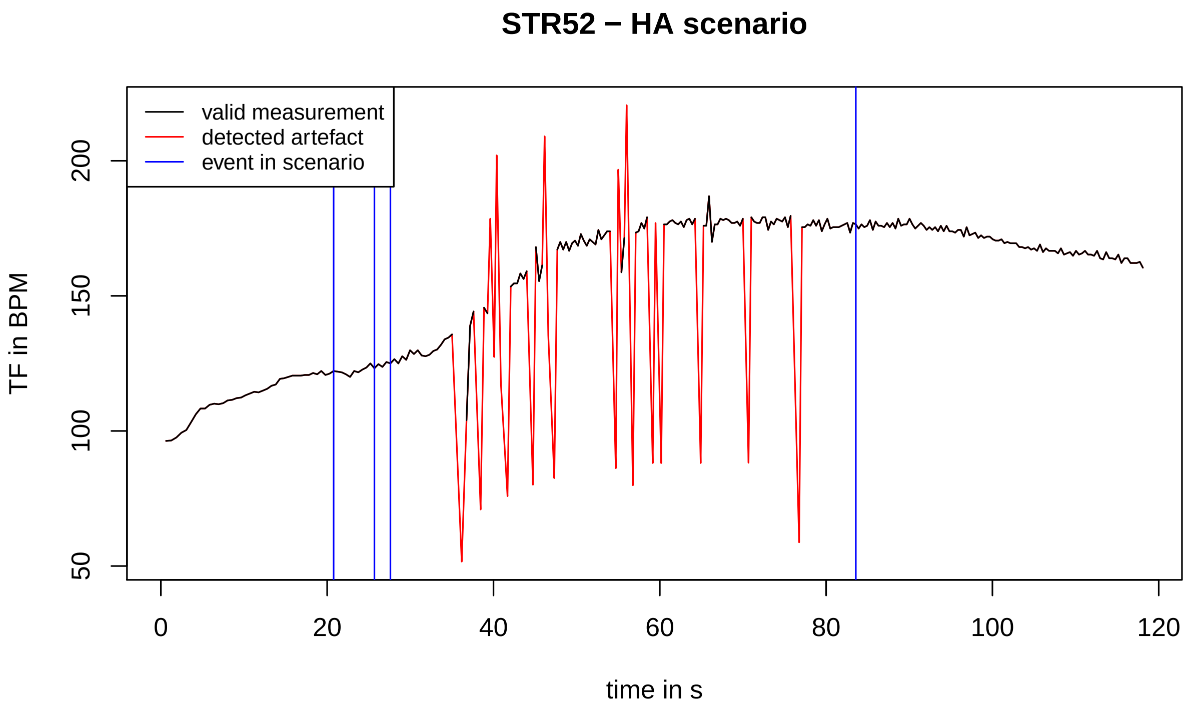
STR52 − HA scenario
valid measurement
detected artefact
event in scenario
TF in BPM
50
100
150
200
0
20
40
60
80
100
120
time in s

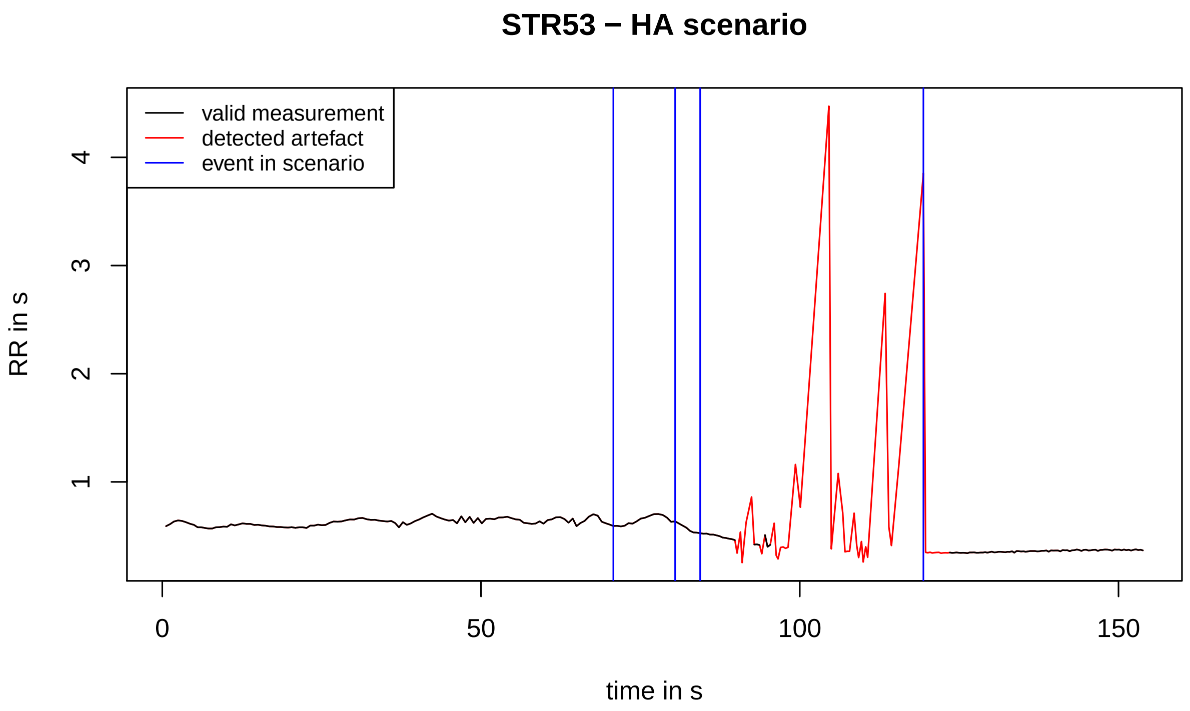
STR53 − HA scenario
valid measurement
detected artefact
event in scenario
RR in s
1
2
3
4
0
50
100
150
time in s

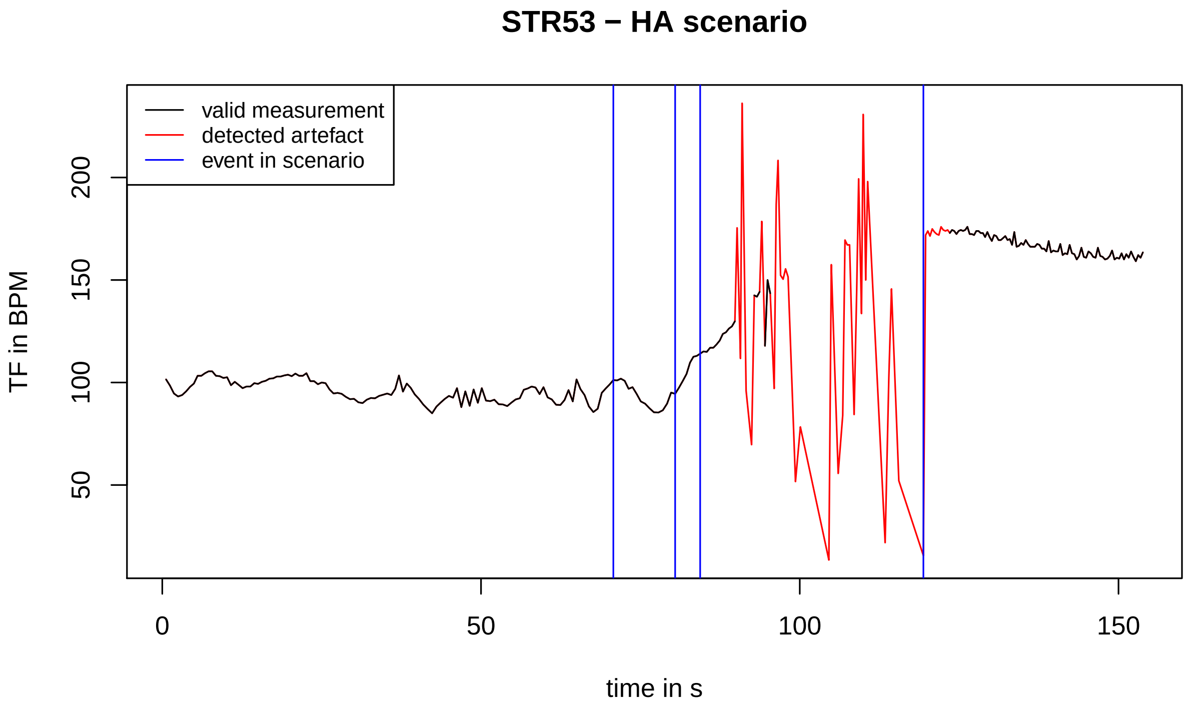
STR53 − HA scenario
valid measurement
detected artefact
event in scenario
TF in BPM
50
100
150
200
0
50
100
150
time in s

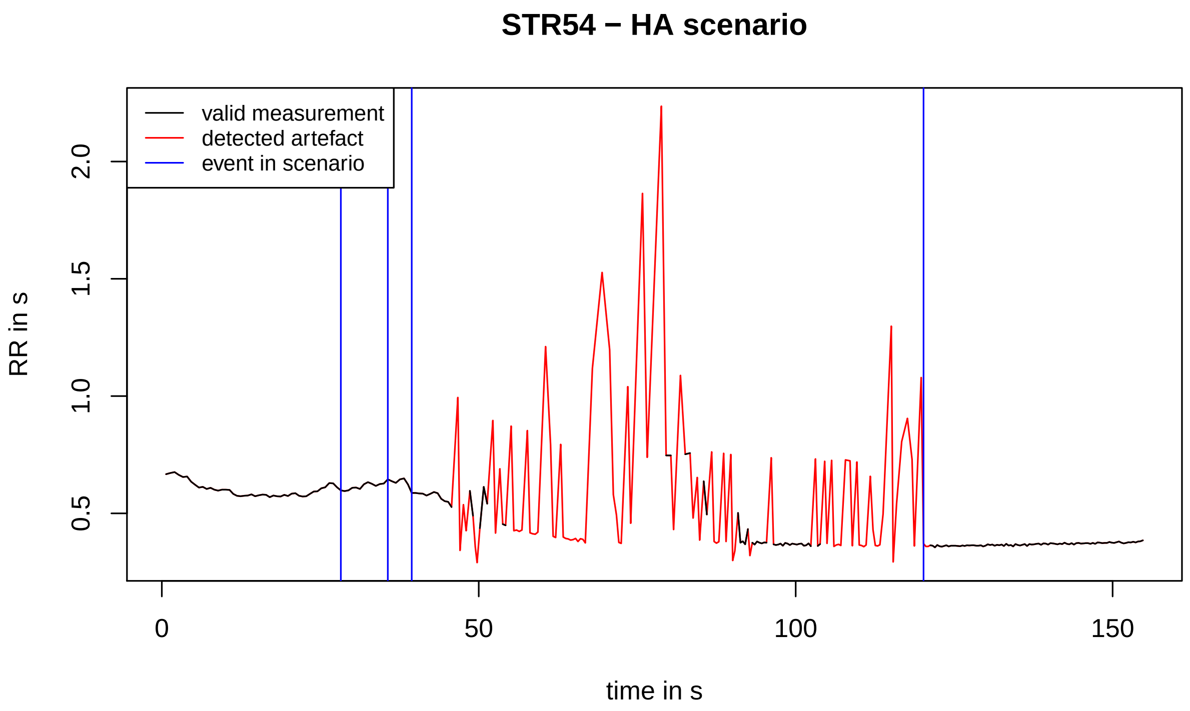
STR54 − HA scenario
valid measurement
detected artefact
event in scenario
RR in s
0.5
1.0
1.5
2.0
0
50
100
150
time in s

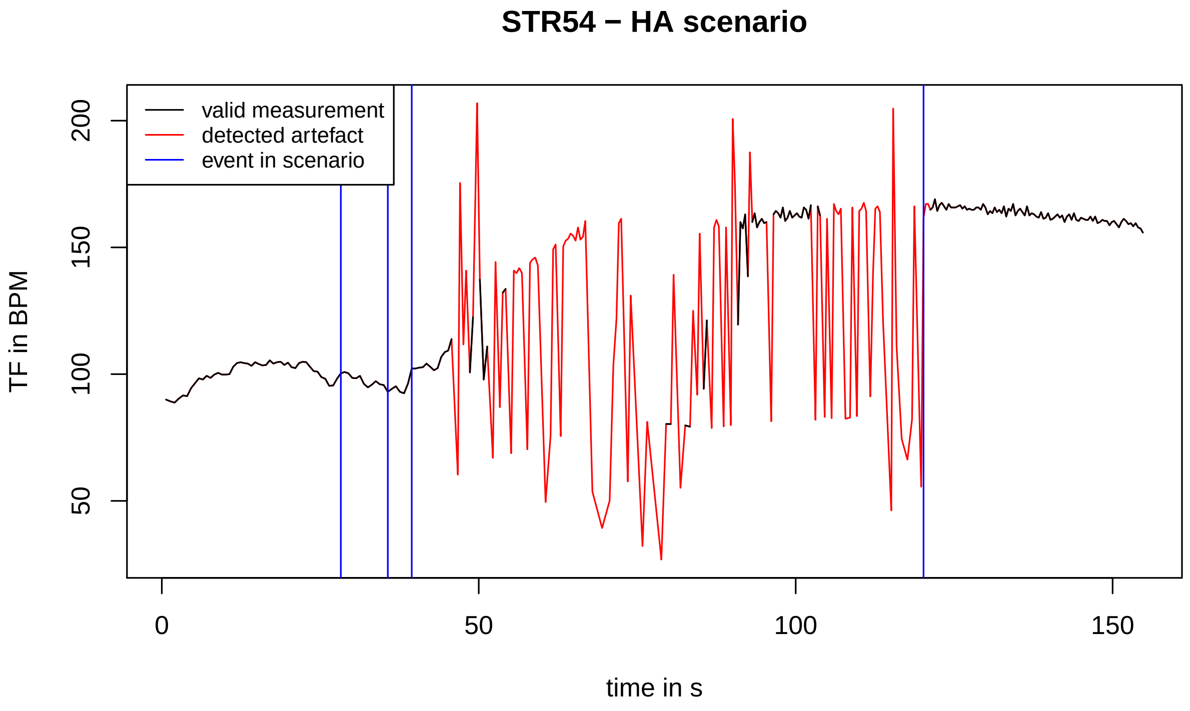
STR54 − HA scenario
valid measurement
detected artefact
event in scenario
TF in BPM
50
100
150
200
0
50
100
150
time in s

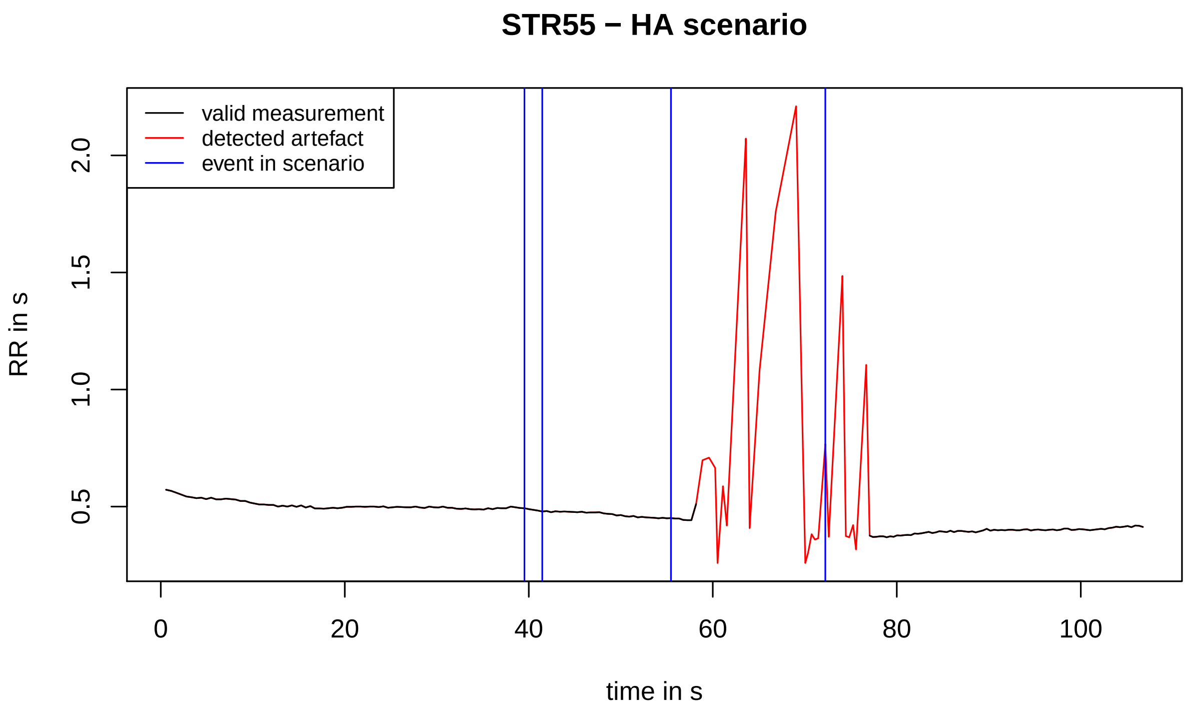
STR55 − HA scenario
valid measurement
detected artefact
event in scenario
RR in s
0.5
1.0
1.5
2.0
0
20
40
60
80
100
time in s

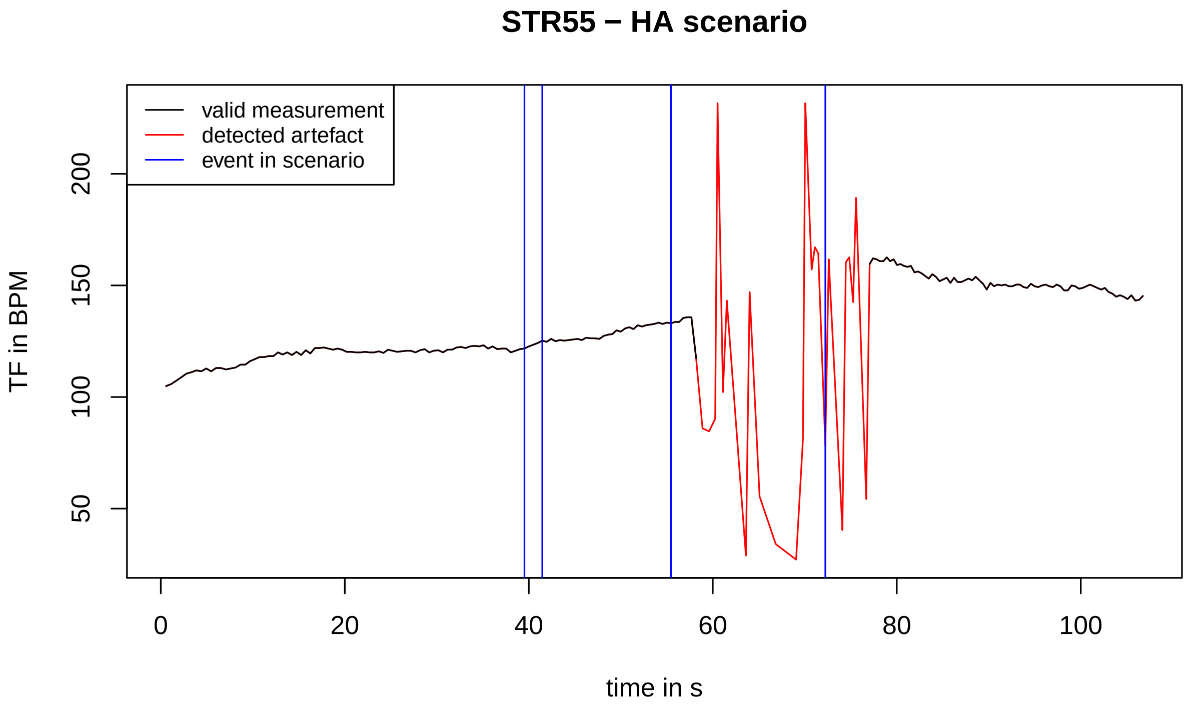
STR55 − HA scenario
valid measurement
detected artefact
event in scenario
TF in BPM
50
100
150
200
0
20
40
60
80
100
time in s

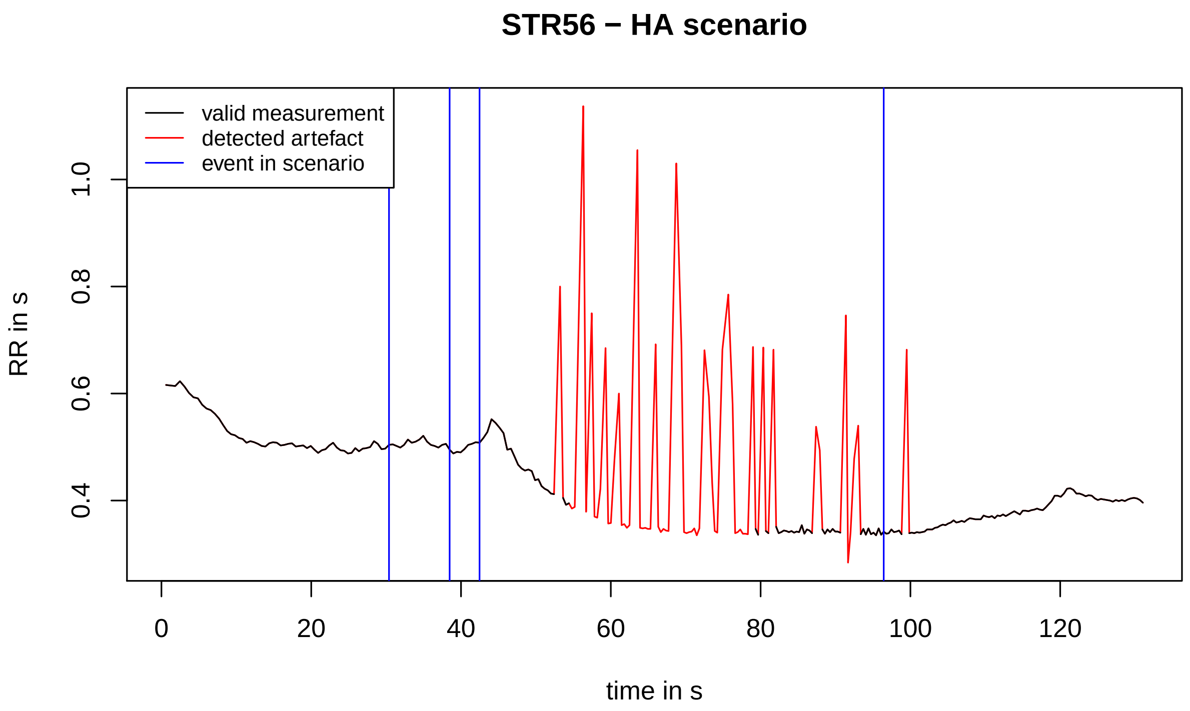
STR56 − HA scenario
valid measurement
detected artefact
event in scenario
RR in s
0.4
0.6
0.8
1.0
0
20
40
60
80
100
120
time in s

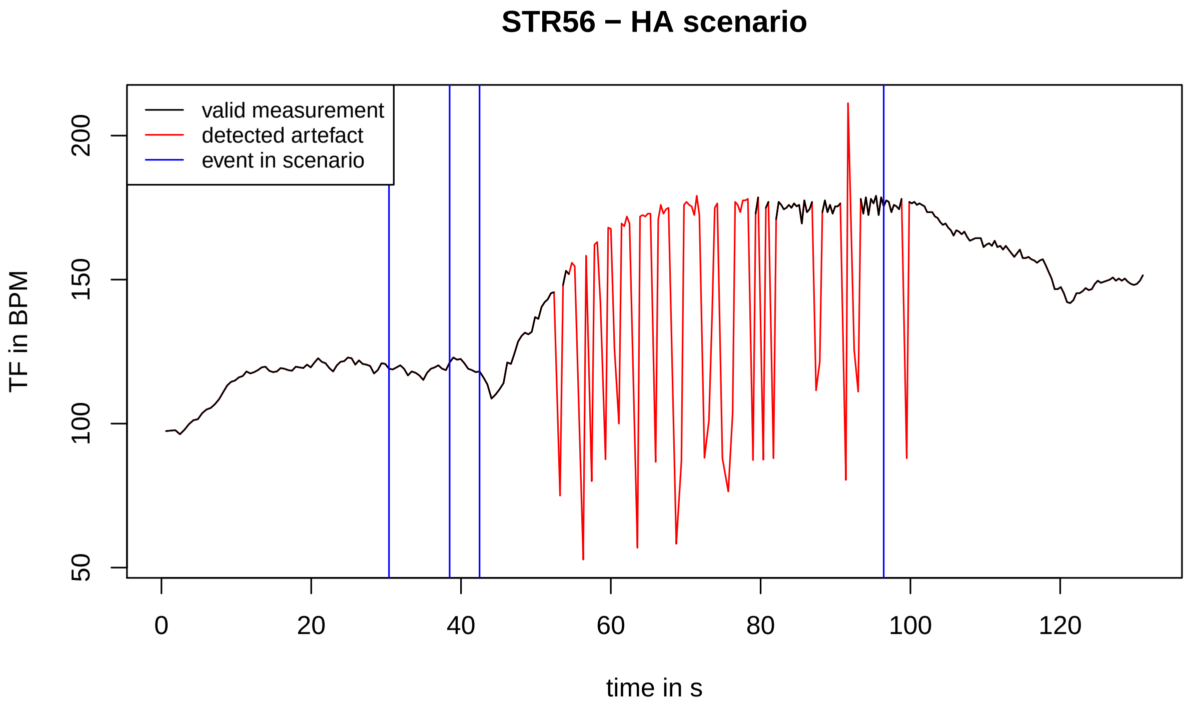
STR56 − HA scenario
valid measurement
detected artefact
event in scenario
TF in BPM
50
100
150
200
0
20
40
60
80
100
120
time in s

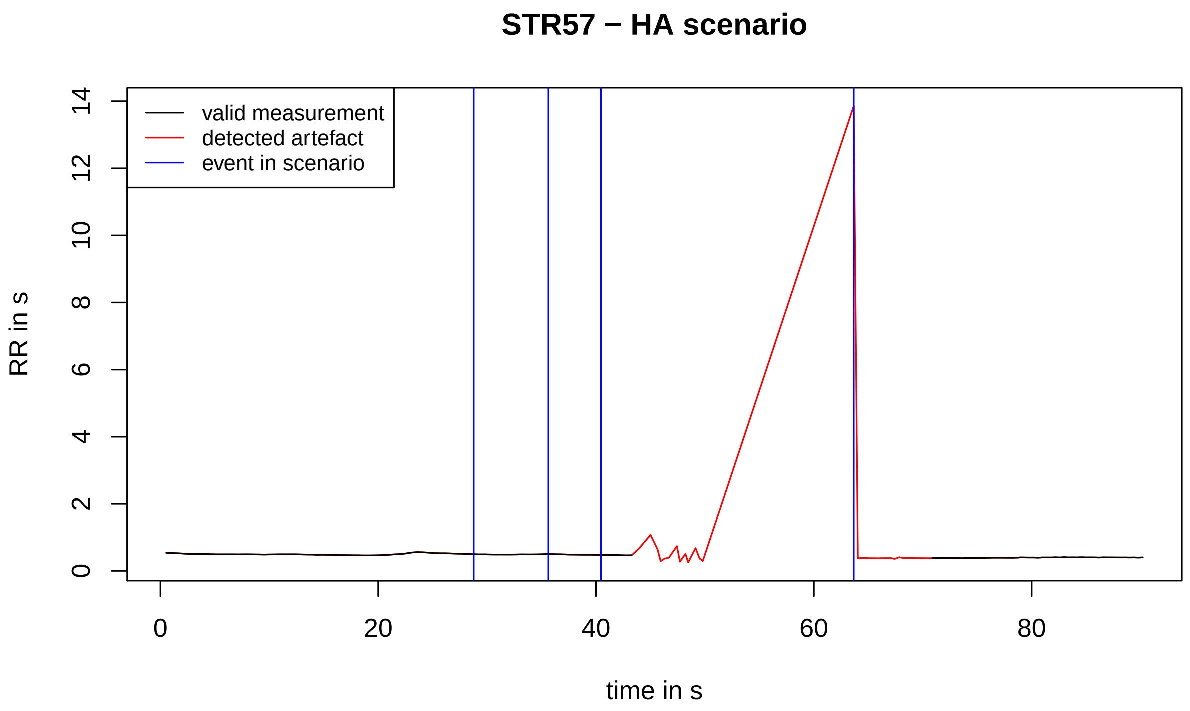
STR57 − HA scenario
valid measurement
detected artefact
event in scenario
RR in s
0
2
4
6
8
10
12
14
0
20
40
60
80
time in s

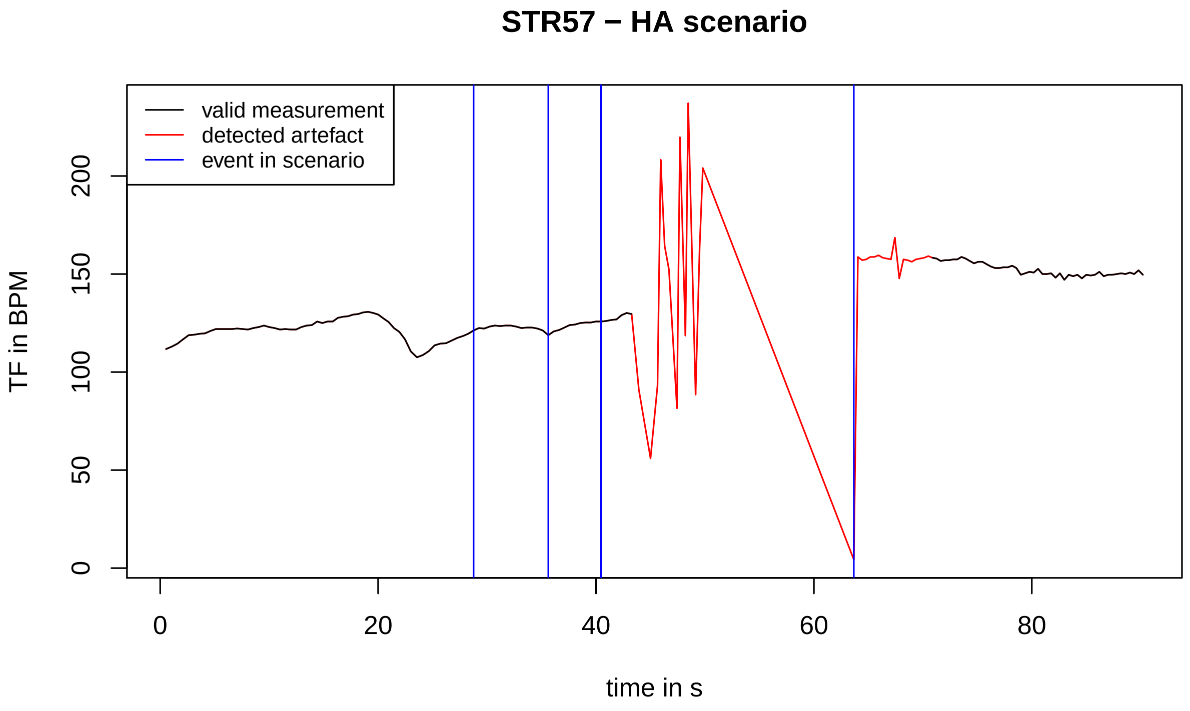
STR57 − HA scenario
valid measurement
detected artefact
event in scenario
TF in BPM
0
50
100
150
200
0
20
40
60
80
time in s

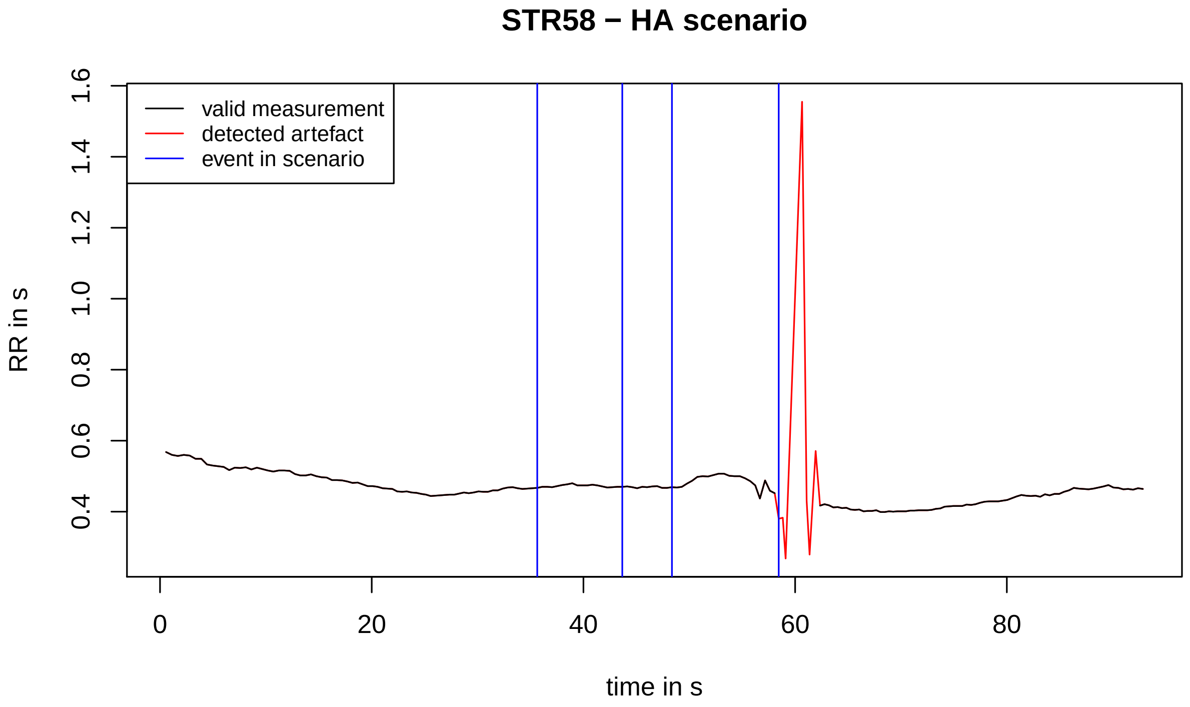
STR58 − HA scenario
valid measurement
detected artefact
event in scenario
RR in s
0.4
0.6
0.8
1.0
1.2
1.4
1.6
0
20
40
60
80
time in s

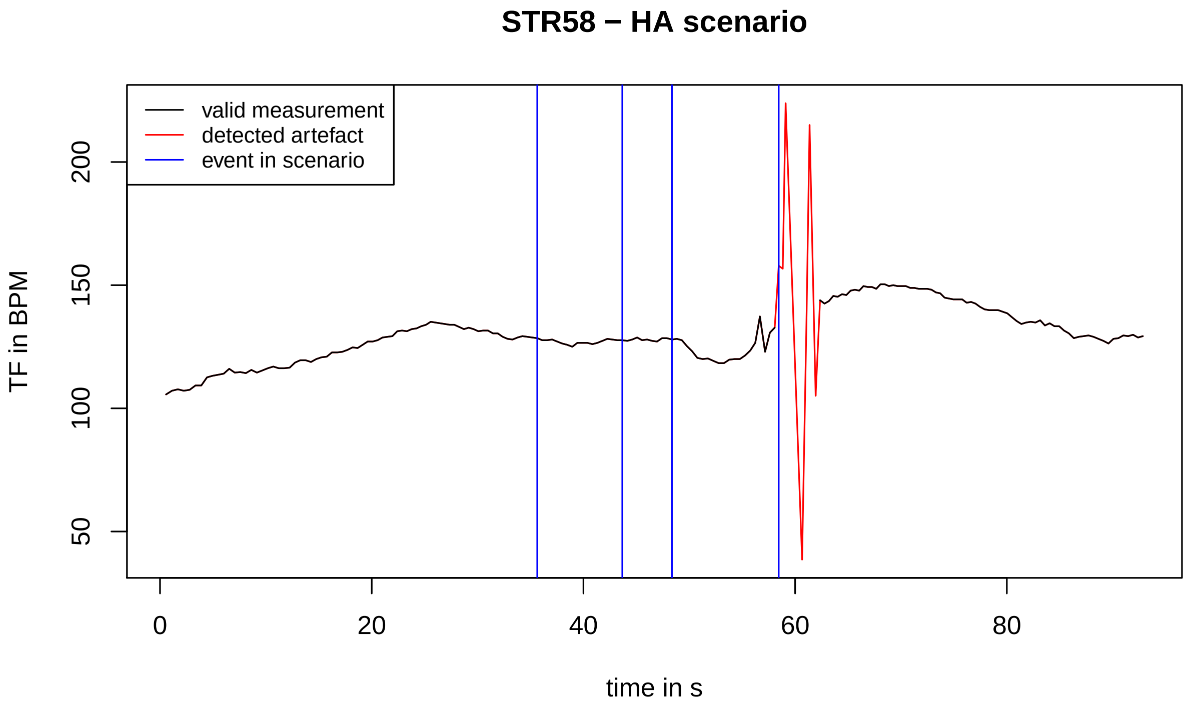
STR58 − HA scenario
valid measurement
detected artefact
event in scenario
TF in BPM
50
100
150
200
0
20
40
60
80
time in s

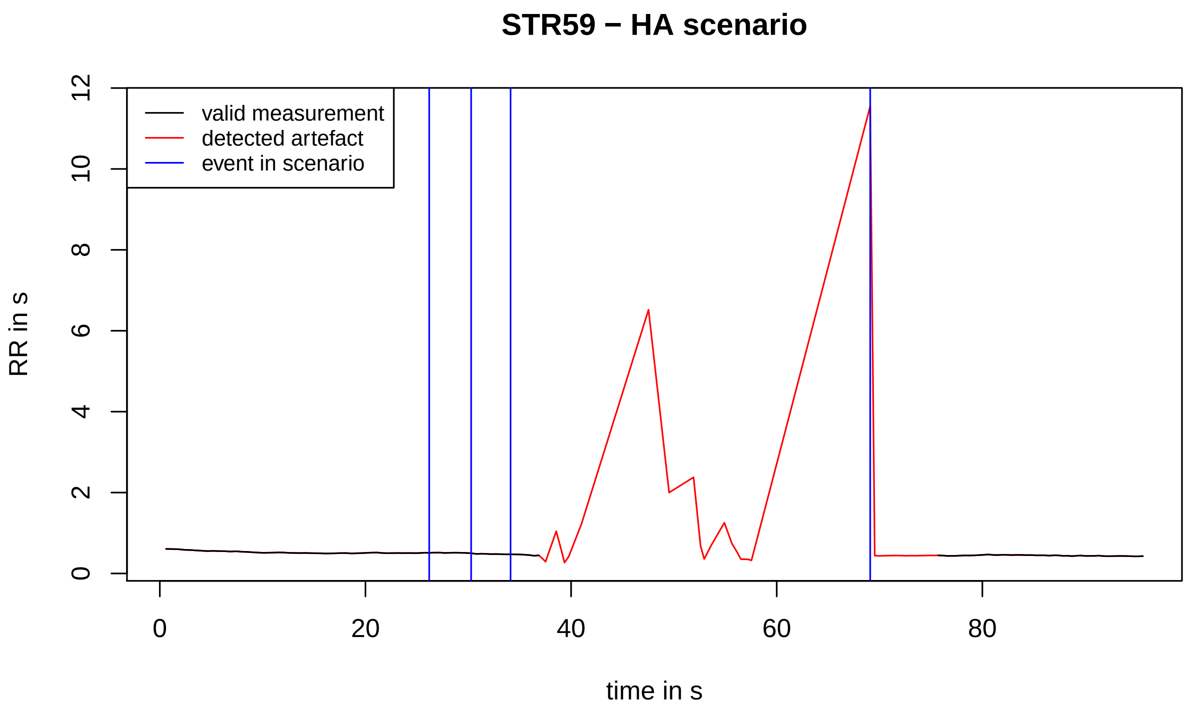
STR59 − HA scenario
valid measurement
detected artefact
event in scenario
RR in s
0
2
4
6
8
10
12
0
20
40
60
80
time in s

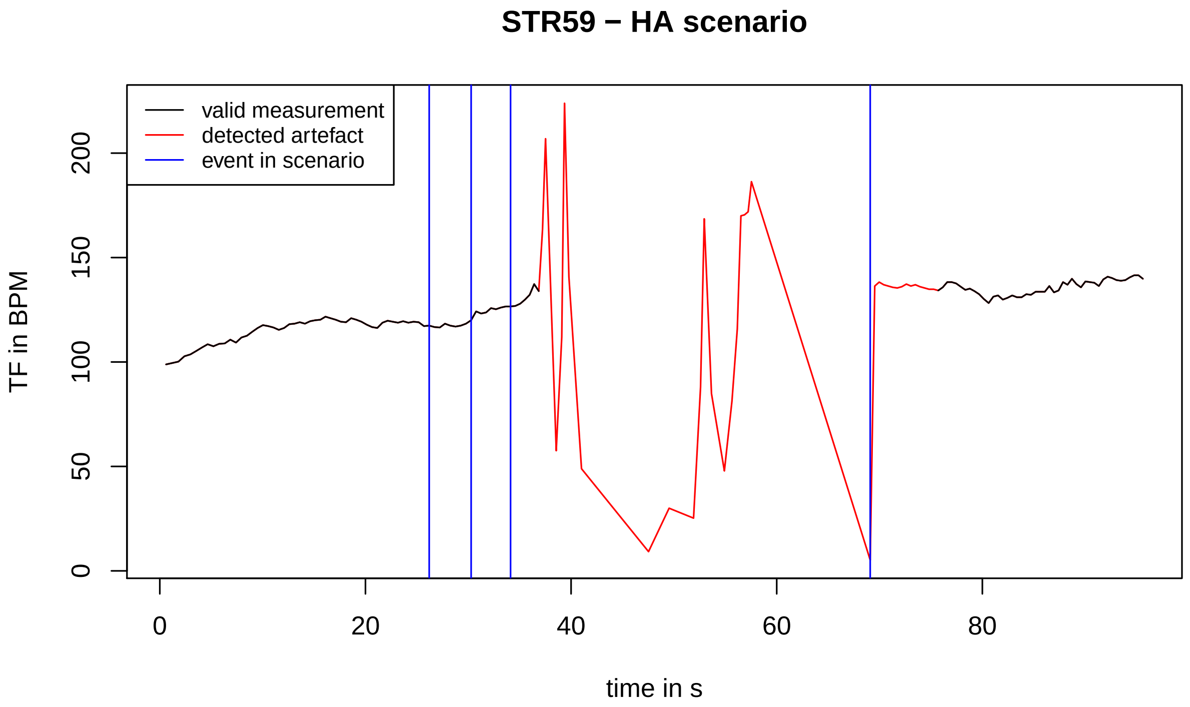
STR59 − HA scenario
valid measurement
detected artefact
event in scenario
TF in BPM
0
50
100
150
200
0
20
40
60
80
time in s

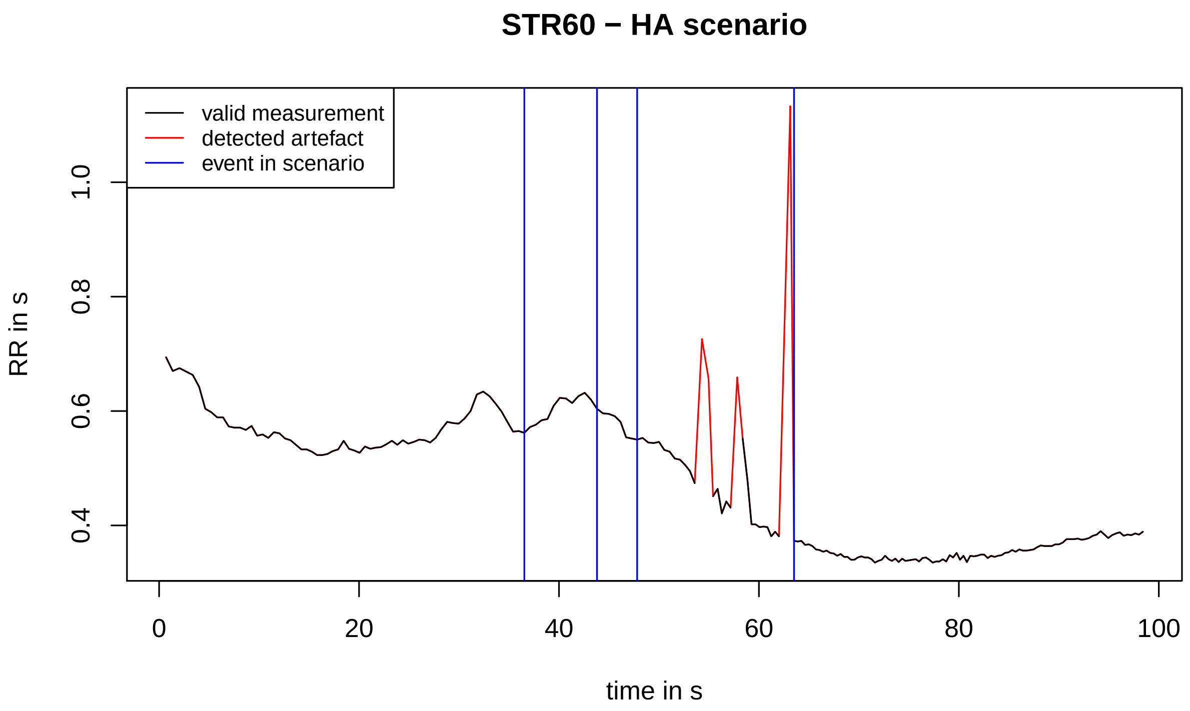

STR60 − HA scenario
valid measurement
detected artefact
event in scenario
RR in s
0.4
0.6
0.8
1.0
0
20
40
60
80
100
time in s


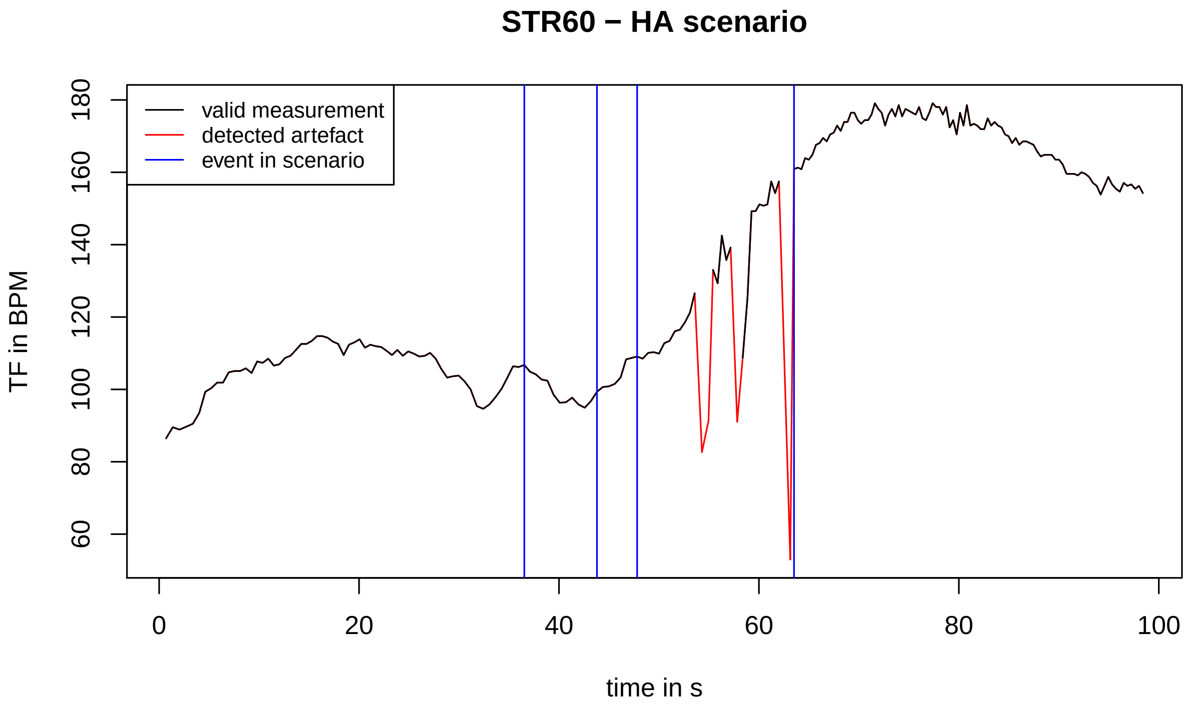

STR60 − HA scenario
valid measurement
detected artefact
event in scenario
TF in BPM
60
80
100
120
140
160
180
0
20
40
60
80
100
time in s

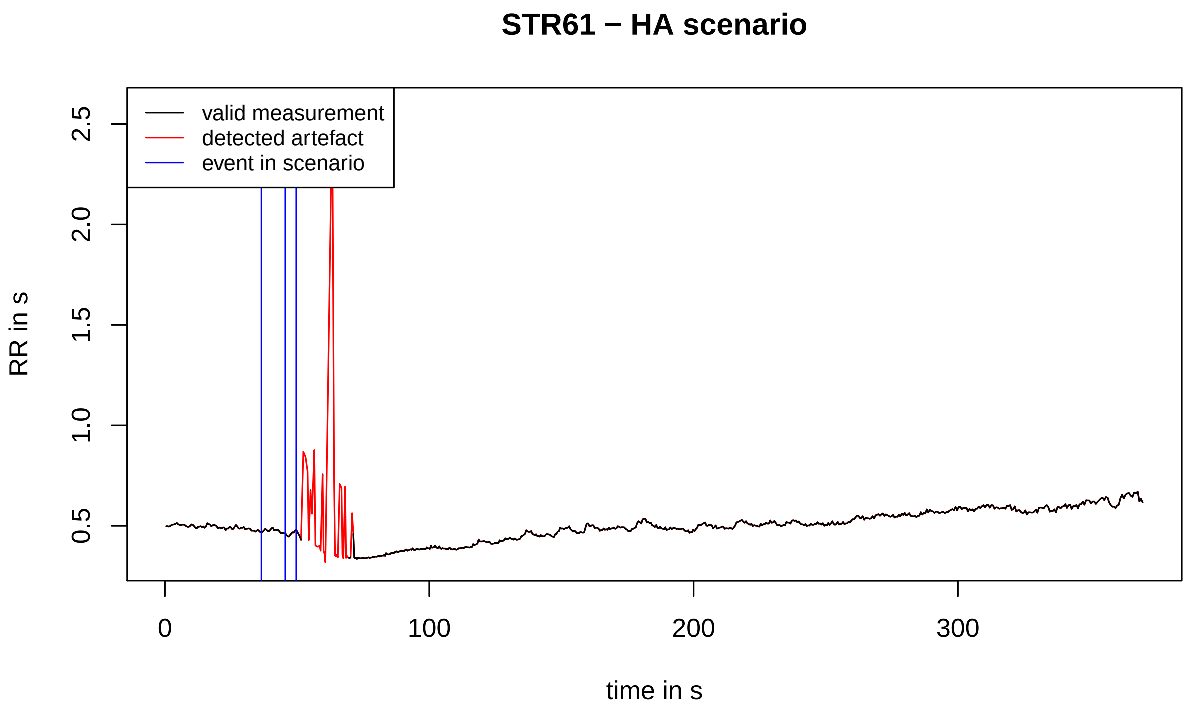
STR61 − HA scenario
valid measurement
detected artefact
event in scenario
RR in s
0.5
1.0
1.5
2.0
2.5
0
100
200
300
time in s

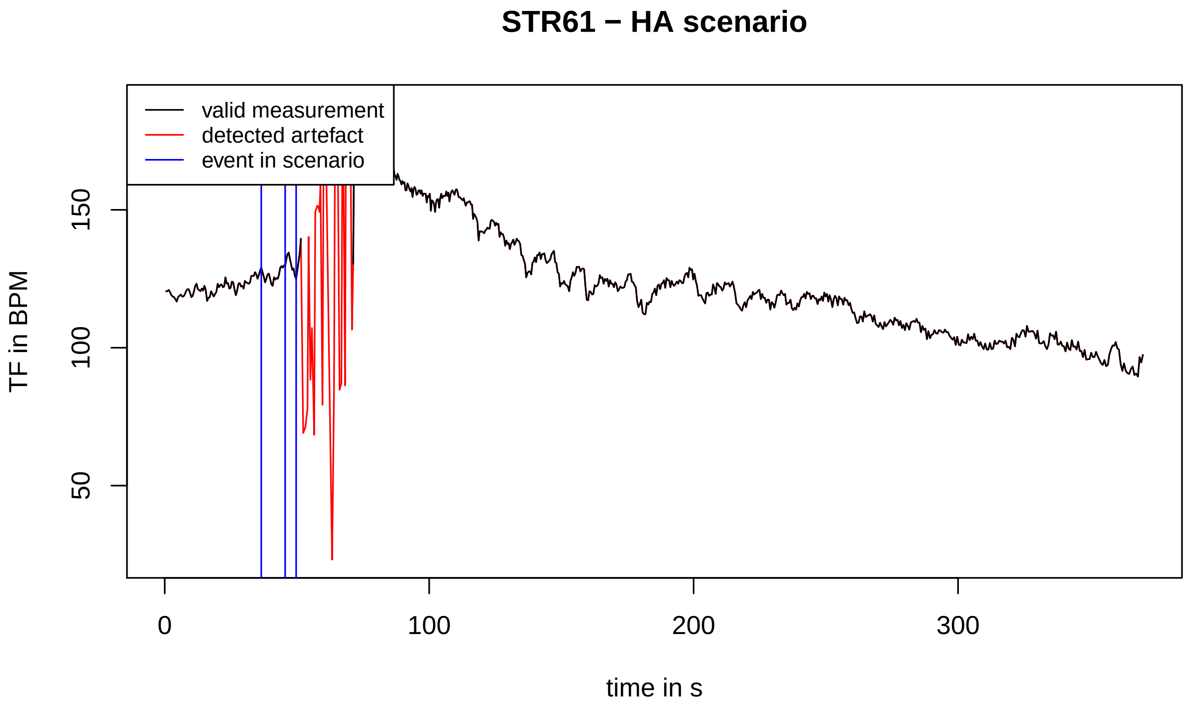
STR61 − HA scenario
valid measurement
detected artefact
event in scenario
TF in BPM
50
100
150
0
100
200
300
time in s

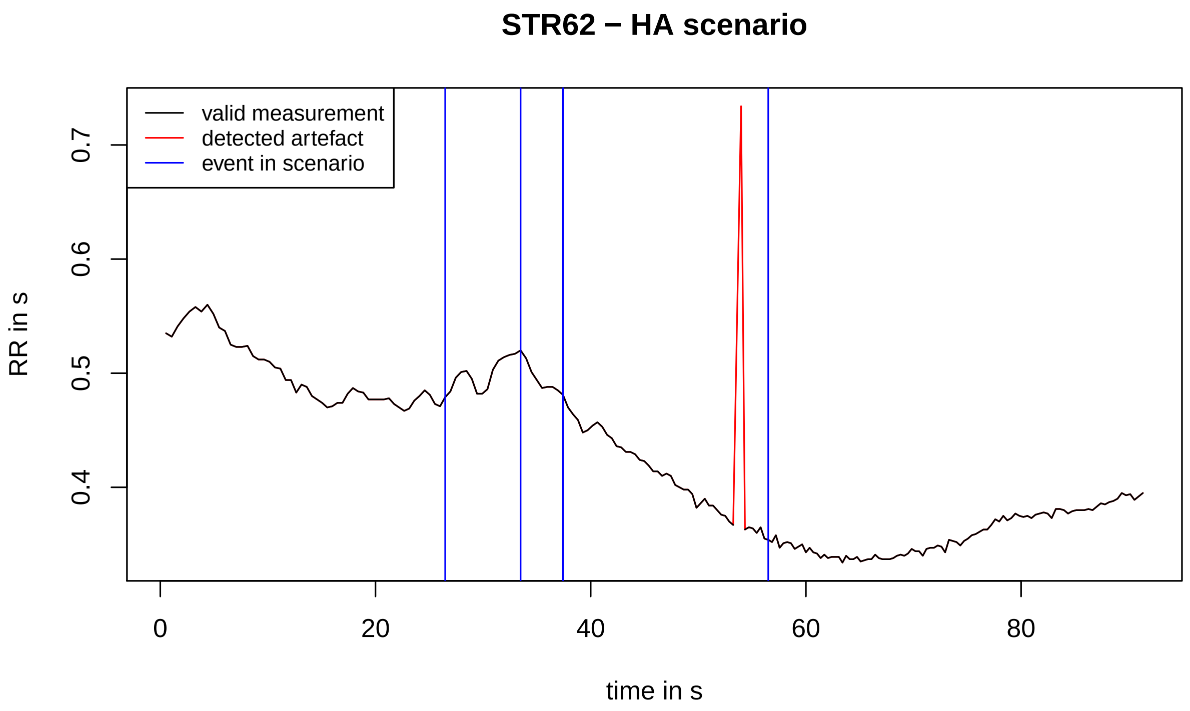
STR62 − HA scenario
valid measurement
detected artefact
event in scenario
RR in s
0.4
0.5
0.6
0.7
0
20
40
60
80
time in s

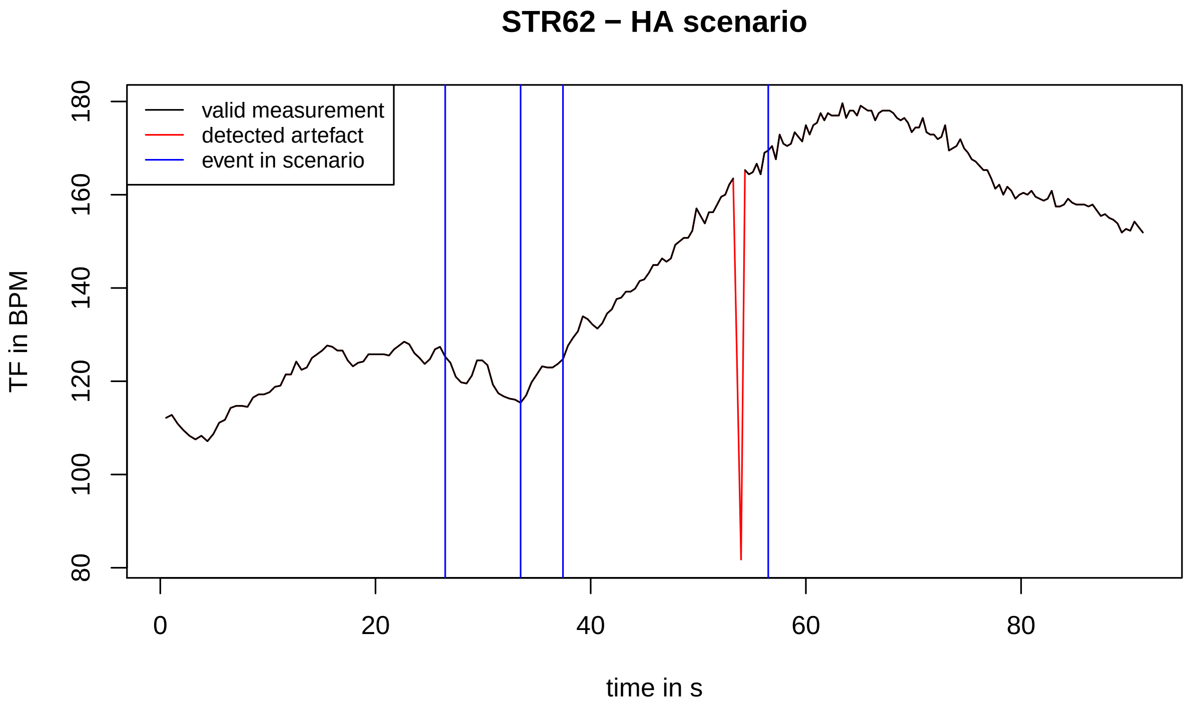
STR62 − HA scenario
valid measurement
detected artefact
event in scenario
TF in BPM
80
100
120
140
160
180
0
20
40
60
80
time in s
